# Highly Heritable and Functionally Relevant Breed Differences in Dog Behavior

**DOI:** 10.1101/509315

**Authors:** Evan L MacLeant, Noah Snyder-Mackler, Bridgett M. vonHoldt, James A. Serpell

## Abstract

Variation across dog breeds presents a unique opportunity for investigating the evolution and biological basis of complex behavioral traits. We integrated behavioral data from more than 17,000 dogs from 101 breeds with breed-averaged genotypic data (N = 5,697 dogs) from over 100,000 loci in the dog genome. Across 14 traits, we found that breed differences in behavior are highly heritable, and that clustering of breeds based on behavior accurately recapitulates genetic relationships. We identify 131 single nucleotide polymorphisms associated with breed differences in behavior, which are found in genes that are highly expressed in the brain and enriched for neurobiological functions and developmental processes. Our results provide insight into the heritability and genetic architecture of complex behavioral traits, and suggest that dogs provide a powerful model for these questions.

## Introduction

Variation across dog breeds provides a unique opportunity for investigating questions about the evolution and biological basis of complex traits. For example, studies of breed differences have led to major advances in our understanding of the genetics of diseases, including cancer, metabolic disorders, and blindness (1), as well as the genetic underpinnings of morphological traits, such as body mass, coat type, and coloration (2). Despite rapid progress in these areas, we still know little about the biological basis of breed differences in behavior. For example, although breed differences in behavior are well documented (reviewed in 3), it remains unknown to what extent these differences are heritable. Further, even less is known about the genetic architecture of these behavioral traits. Are they largely polygenic – as is the case for most complex traits – or instead predominantly influenced by a small set of loci (2, 4)?

Compared to humans, dogs provide a uniquely powerful model for questions about behavioral evolution, due to a simplified genetic architecture resulting from population bottlenecks during domestication and strong selection during subsequent breed diversification (5–7). The majority of variance among modern breeds has likely resulted from the repeated crossing of novel phenotypes, which – drawing on a limited genetic toolkit – has given rise to extraordinary phenotypic diversity. In addition to these practical advantages, dogs exhibit complex cognitive and behavioral phenotypes, with striking parallels to traits in humans (8–13). For example, common genetic mechanisms contribute to individual differences in social behavior in dogs and humans, with relevance to understanding behavioral syndromes such as hypersociability (14). However, research to date has been conducted with small sample sizes from a restricted number of breeds, limiting our ability to make inferences about the evolution and biological basis of behavioral diversity across breeds (15, 16).

Here, we investigated the genetic basis of breed differences in behavior by combining a behavioral dataset of more than 14,000 dogs from 101 breeds (Table S1) with breed-typical genotypic data from over 100,000 loci across the dog genome. Drawing on these data, we quantified, for the first time, the heritability of 14 behavioral traits across breeds, and identified key genetic variants implicated in biological pathways associated with breed differences in behavior.

## Results

A fundamental criterion for the evolution of a trait by natural or artificial selection is that variance in the trait is heritable. We assessed the heritability (h^2^) of 14 behavioral traits (Fig 1) measured by the Canine Behavioral Assessment and Research Questionnaire (C-BARQ), a well-validated instrument for quantifying diverse aspects of dog behavior (17–20), including aggression, fear, trainability, attachment, and predatory chasing behaviors. We combined behavioral data from 14,020 individual dogs with breed-level genetic identity-by-state (IBS) estimates from two independent studies (N = 5,697, *Materials and Methods*; Hayward et al. (4) and Parker et al. (21)), which allowed us to incorporate behavioral and genetic variance both within and across breeds. Using a mixed-effects modeling approach (Efficient Mixed-Model Association; EMMA) to control for relatedness between breeds, we found that a large proportion of variance in dog behavior is attributable to genetic factors (Fig 1). The mean heritability was 0.51 ± 0.12 (SD) across all 14 traits (range: *h^2^* 0.27-0.77), and significantly higher than the null expectation in all cases (permutation tests, p < 0.001).

**Fig 1.**
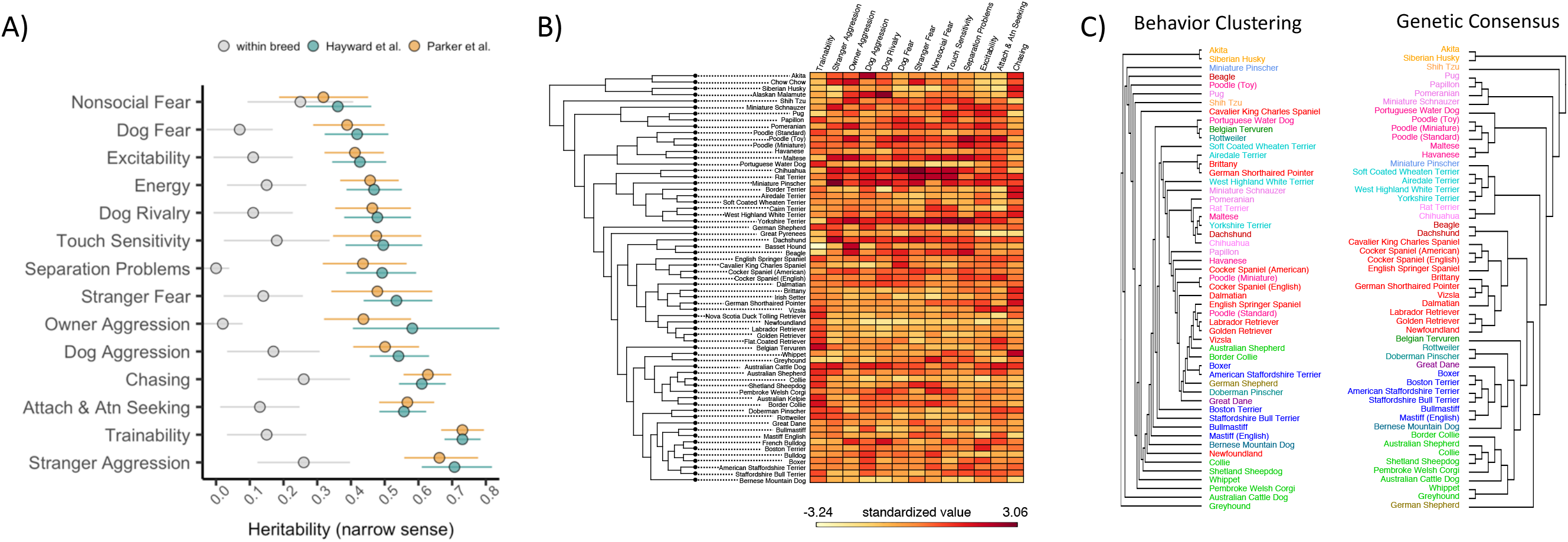
Heritability estimates, breed-level behavioral data, and clustering based on behavioral and genetic data. A) Heritability (h2) estimates (proportion of variance attributable to genetic factors) for 14 behavioral traits. Genotypic variation accounts for five times more variance in analyses across vs. within breeds (within-breed estimates compiled from llska et al., 2017). Points for Hayward et al. and Parker et al. reflect the results of analyses with independent genetic datasets. Error bars reflect the 95% confidence intervals. B) Heatmap of breed-average behavioral scores plotted alongside a cladogram of breed relatedness from Parker et al., 2017. C) Breed dendrograms from clustering based on behavioral (left panel) and genetic (right panel) similarity. Colors correspond to clades from Parker et al., 2017.

These estimates are also significantly higher than those identified in previous studies assessing heritability of these traits in large within-breed samples (t13 = -12.25, p < 0.001; 22, but see 23). Estimating between-breed variance thus yields h^2^ estimates that are on average, five times higher (range = 1.3-25.5 times higher), which is likely due to more variance among, compared to within breeds. Interestingly, the traits with the highest heritability were trainability (h^2^ = 0.73), stranger-directed aggression (h^2^ = 0.68), chasing (h^2^ = 0.62) and attachment and attention seeking (h^2^ = 0.56), which is consistent with the hypothesis that these behaviors have been important targets of selection during the cultivation of modern breeds (3). Hierarchical clustering of breeds based on behavioral traits recapitulated the genetic similarities with striking accuracy (Fig 1; cophenetic correlation coefficient = 0.32; p = 0.01).

To identify specific loci associated with breed differences in behavior, we conducted a Genome-Wide Association Study across breeds. Specifically, we modeled breed-average behavioral scores as a function of breed-average allele frequencies, controlling for relatedness among breeds using EMMA (24). To further control for possible inflation of p values due to cryptic population stratification, we corrected p values using the approach described by Amin et al. (25) Each trait was modeled twice: once with each of the independent genetic datasets. The effects of the same single-nucleotide polymorphisms (SNPs) on each behavior were strongly correlated across the two datasets (median r = 0.77, range: r = 0.68-0.82, Fig S1), and the strength of this association increased (median r = 0.93) when we included SNPs significantly associated with a trait at a nominal p ≤ 0.01 (Fig S2). We therefore used Fisher’s combined probability test to combine the p values for each shared SNP across the two datasets (*Materials and Methods*).

Overall, we identified 131 unique SNPs that were significantly associated with at least one of the 14 behavioral traits (Bonferroni p ≤ 0.05, Fig 2). Forty percent of these SNPs (n= 52) were located within a gene – none of which encoded for changes in the amino acid sequence of the protein (see *SI* for analyses using other distance thresholds for mapping SNPs to genes). On average, the top SNP explained 15% of variance in the behavioral trait (range, PVE = 0.06-0.25, Fig 2, Table S2). Thus, while we identify multiple variants with moderately large effects, the variance explained by individual SNPs is far less than that explained by additive variation across the genome (heritability), suggesting that as in humans, behavioral traits in dogs are highly polygenic. However, the variance explained by the top SNPs in our analysis across breeds was, on average, more than 5 times higher than that from within-breed association studies using the C-BARQ (22).

**Fig 2.**
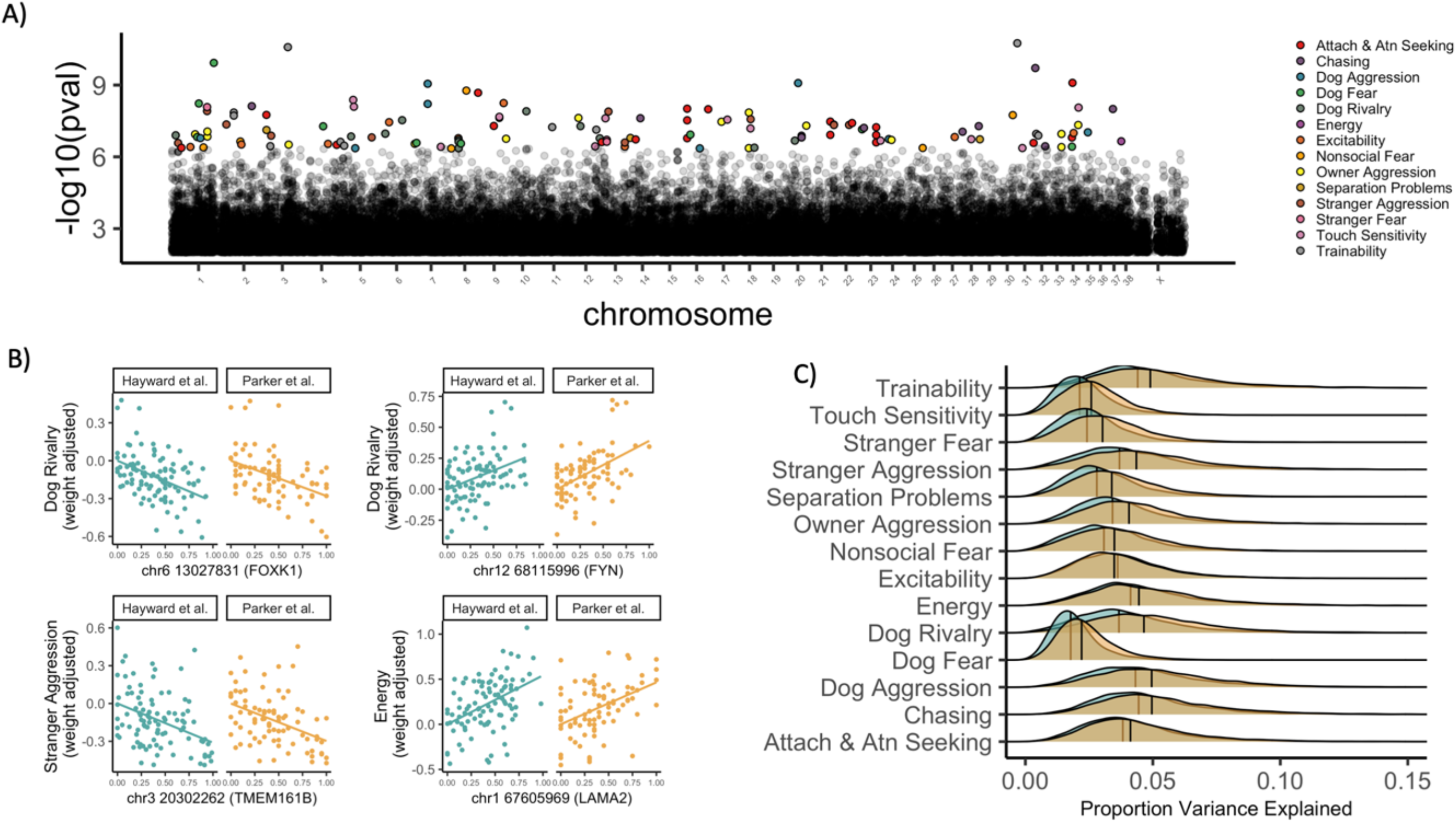
Genetic associations with breed differences in behavior. A) Manhattan plot showing SNPs associated with behavioral traits (color coded) after Bonferroni correction. B) Behavioral trait values, corrected for body weight, as a function of allele frequency. Plots show data for 4 SNPs in genes with sequence or expression differences between foxes artificially bred for tameness or aggression. C) Distributions of the proportion variance explained (PVE) by SNPs associated with behavioral traits at p ≤ 0.05, after correction for the false discovery rate. Turquoise points and distributions: Hayward et al. genetic data; Yellow points and distributions: Parker et al. genetic data.

To examine if these variants may be linked to behavior-relevant genes, we further derived gene-level associations using a meta-analytic approach (*Materials and Methods*). Many of the gene-level associations with dog behavioral traits (Table S3) include (i) candidate domestication genes, (ii) genes mapped to phenotypes implicated in domestication, (iii) genes implicated in behavioral differences between foxes bred for tameness or aggression, and (iv) genes that underwent positive selection in both human evolution and dog domestication (Table S4). For example, *PDE7B*, which is differentially expressed in the brains of tame and aggressive foxes (26) has been identified as a target of selection during domestication, and is highly expressed in the brain (27) where it functions in dopaminergic pathways (28). In our analyses, SNPs in this gene were associated with breed differences in aggression, which is consistent with data from experimentally bred foxes, as well as hypotheses that selection against aggression was the primary evolutionary pressure during initial domestication events (29–31).

The gene-trait associations identified in our study also align closely with similar associations in human populations (Table S5). For example, breed differences in aggression are associated with multiple genes that have been linked to aggressive behavior in humans. Molecular associations with breed differences in energy include genes previously linked to resting heart rate, daytime rest, and sleep duration in humans. Lastly, breed differences in fear were associated with genes linked with temperament and startle response in humans, and several of the genes implicated in breed differences in trainability have been previously associated with intelligence and information processing speed in humans.

If the variants in genes identified in our analyses make major contributions to behavior and cognition, then the associated genes should be (i) involved in biological processes related to nervous system development and function, and (ii) primarily expressed in the brain. Indeed, we found that behavior-associated genes (as identified through meta-analysis) were enriched for numerous nervous system processes (Fig 3, Table S6). These processes include neurogenesis, neuron migration and differentiation, axon and dendrite development, and regulation of neurotransmitter transport and release. We also identified gene ontology (GO) terms relating to key developmental processes implicated in domestication (32, 33), including neural tube development, neural crest cell differentiation and migration, and the development and organization of brain structures including the forebrain, cerebral cortex, and cerebellum (Table S6). On a trait-specific level, we identified several interesting links between dog behavior and putative biological mechanisms implicated in the regulation of these behaviors. For example, dopaminergic pathways play important roles in mammalian social attachment (34, 35), and GO terms relating to dopamine transport were significantly enriched in genes associated with breed differences in attachment and attention seeking.

**Fig 3.**
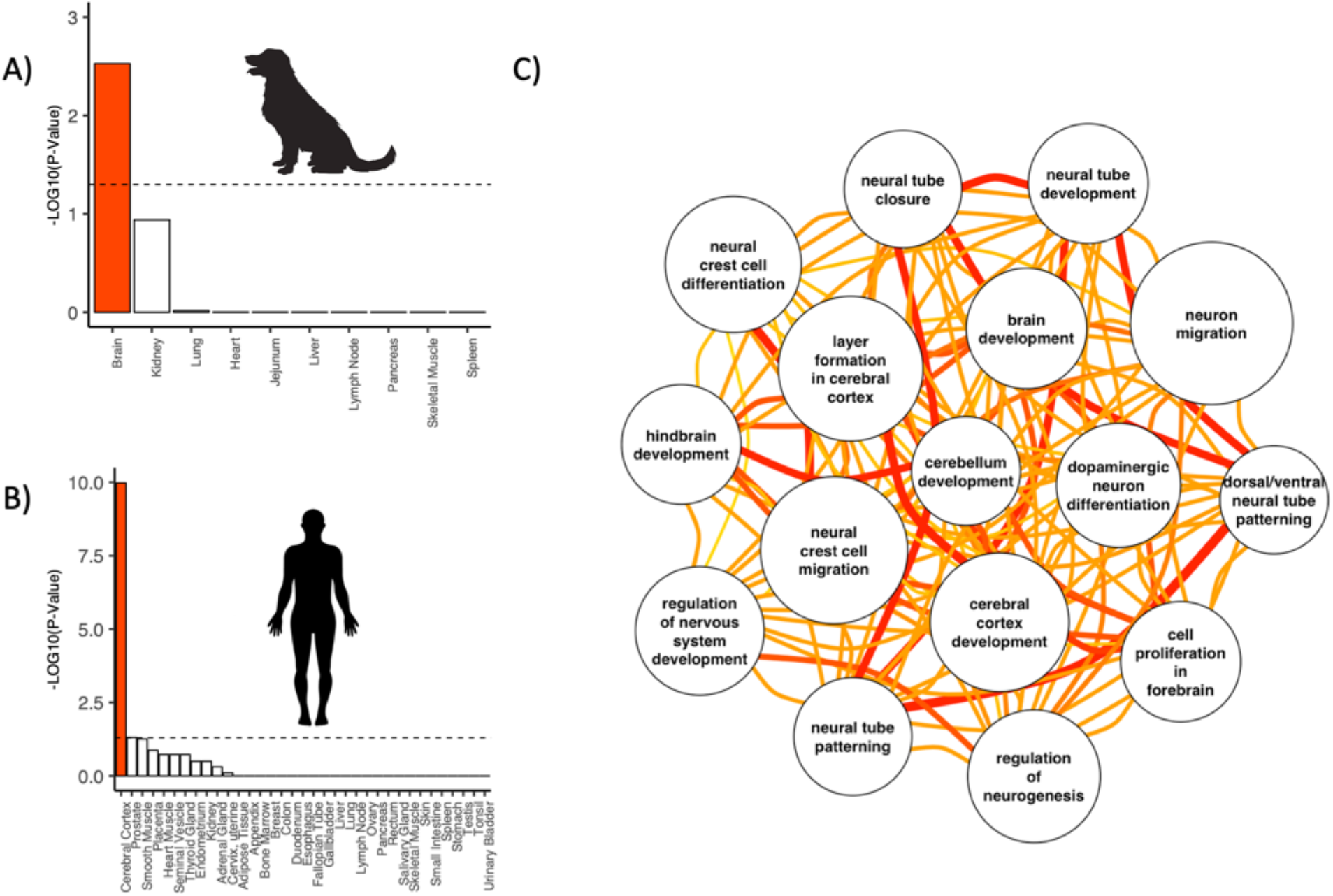
Genes containing behavior-associated SNPs are highly expressed in the brain and associated with gene ontology terms relating to brain development and function. (A) Enrichment tests using dog gene expression to identify tissue-specific genes. (B) Enrichment tests using human gene expression to identify tissue-specific genes. Bars reflect the -log 10 p value from a hypergeometric test for tissue-specific gene enrichment, corrected for multiple comparisons. The dashed line indicates -log10(p = 0.05) and the results for brain tissue are highlighted in red. (C) Network plot for a subset of gene ontology terms relating to brain development and function that were associated with breed differences in behavior. Edge colors and line widths reflect Resnik’s similarity scores between GO terms. Wider and redder lines reflect greater similarity between nodes. Node sizes are inversely proportional to p values from enrichment tests.

To examine if the genes identified through GWAS are predominantly expressed in the brain, we analyzed tissue-specific enrichment for the set of genes with SNPs that were significantly associated with any of the 14 behavioral traits. We used two datasets of tissue-specific expression: (i) gene expression across 10 tissues from a sample of 4 dogs (36), and (ii) human gene expression from larger samples of individuals across a greater range of tissues (N = 35) (37). In both datasets, we found that the genes containing SNPs associated with dog behavior are significantly more likely to be expressed in the brain (Fig 3, hypergeometric test: dog tissue: p < 0.001, human tissue: p < 0.001; see *SI* and Figure S3 for similar results including SNPs nearby, but not necessarily in, genes). Together, this suggests that the SNPs identified in our GWAS may affect behavioral processes by altering expression levels in genes that are highly expressed in the brain.

## Discussion

The vast phenotypic diversity, and simplified genetic architecture of dog breeds has led to major advances in our understanding of complex traits relevant to morphology and disease. Our findings suggest that dog breeds also provide a powerful and highly tractable model for questions about the evolution and genetic basis of behavioral traits. Breed differences in behavior covary strongly with relatedness between breeds, and for several traits, genotype accounts for more than 50% of behavioral variation across breeds – up to 25x higher than heritability estimates from genetic studies within breeds. Individual SNPs that are associated with behaviors tend to fall in genes that are disproportionately expressed in the brain, and are involved in pathways related to the development and expression of behavior and cognition. In addition, the variants associated with breed differences in behavior are found in genes with sequence or brain-expression differences in foxes artificially bred for tameness or aggression, and are implicated in human behavioral genetics, suggesting that these genes may play important roles in modulating behavior across species.

One limitation of this study is that genotypic and phenotypic data were not collected from the same subjects, but rather aggregated across independent datasets. This limitation is mitigated by the fact that both the genetic and behavioral datasets were collected from large representative samples, and that our findings were robust across resampling and independent genetic datasets. Nonetheless, future work incorporating genotypic and phenotypic data from the same subjects will be important for finer-resolution trait-mapping.

More so than most model organisms, dogs exhibit a suite of cognitive and behavioral traits that make them a unique model for complex aspects of human social behavior and cognition (11, 38). These similarities are hypothesized to result from convergent evolution, due to similar selective pressures in human evolution and dog domestication (8, 39). Thus, the combination of phenotypic diversity across breeds, the expression of complex traits shared with humans, and the comparative simplicity of trait-mapping in this species, make dogs an invaluable organism for questions about the genetic bases of complex behaviors.

## Materials and Methods

### Genetic data

We estimated genetic relationships among breeds using identity-by-state (IBS) matrices generated in PLINK (40) with data from two large-scale dog genotyping analyses: Hayward et al. (4) and Parker et al. (21). Both datasets were generated using the Illumina Canine HD SNP chip (Illumina, San Diego, CA), with 12,143 custom markers added in the study by Hayward et al. The Hayward et al. dataset included 160,727 SNPs after phasing with a minor allele frequency filter of > 0.01 (https://doi.org/10.5061/dryad.266k4). The Parker et al. dataset included 150,131 SNPs, and was obtained directly from the authors. The dataset from Parker et al. (21) contains individuals sampled in Hayward et al. (4), and these subjects were removed from the Parker et al. data in order to obtain independent genetic datasets for analysis. The resulting datasets included 4,342 dogs from Hayward et al. (4) and 1,355 dogs from Parker et al. (21). Median inter-SNP distances were 5,829 bases for the Parker et al. data and 6,606 bases for the Hayward et al. data. For heritability analyses, breed-level IBS matrices were multiplied by an individual-level incidence matrix in order to generate an individual-level IBS matrix. To avoid the assumption that members of the same breed were clonal, pairwise within-breed IBS values were set to the average IBS value between members of that breed.

### Behavioral data

Behavioral data were obtained from the Canine Behavioral Assessment and Research Questionnaire (C-BARQ), a well-validated instrument for the assessment of diverse aspects of dog behavior (18, 41, 42). This dataset included all C-BARQ entries for pet dogs collected between 2005-2016, and contained 29,656 dogs. These data were filtered to include only pure-bred dogs (owner report), and further restricted to breeds represented in the genetic data sources described above (N = 14,020, Table S1). To ensure representative samples, we included only breeds with at least 25 observations per breed for analysis. Our final behavioral dataset linked to the Parker et al. (21) molecular data included 12,806 dogs from 86 different breeds. Our final behavioral dataset linked to the Hayward et al. (4) molecular data included 13,907 dogs from 98 different breeds.

### Heritability analyses

Heritability was estimated using Efficient Mixed Model Association (EMMA) (24, 43). For comparison to a more conventional approach, we conducted supplementary analyses implemented with Markov chain Monte Carlo (MCMC) linear mixed models (44). Although both approaches yield estimates of the phenotypic variance attributable to (additive) genetic and environmental effects, they differ in the computations through which variance components are estimated. Specifically, EMMA employs restricted maximum likelihood (REML) to efficiently estimate variance components whereas variance components are numerically optimized in the MCMC framework. EMMA models were fit using the NAM R package (45), and MCMC models were fit using the MCMCglmm R package (44).

To estimate heritability for each behavioral trait we employed a resampling procedure as follows: (i) randomly sample 25 individuals from each breed without replacement; (ii) estimate heritability in this subsample. The mean value across resampling was used as the final heritability estimate. For MCMC modeling, each model used a 1000 iteration burn-in, followed by a 10-iteration thinning interval across 9,000 subsequent iterations. For each MCMC model, we retained the mean heritability value from the posterior distribution.

Heritability estimates from EMMA and MCMC were highly correlated within each of the two genetic datasets (Fig S4; Hayward et al., R = 0.91; Parker et al., R = 0.92), with no significant difference in the magnitude of heritability estimates between methods (Hayward et al., t13= 1.04, p = 0.32; Parker et al., t13 = -1.35, p = 0.20). Across genetic datasets, the heritability estimates for each trait were also highly correlated using both statistical approaches (MCMC: R = 0.81; EMMA: R = 0.94). The distributions of heritability estimates with the Hayward et al. and Parker et al. datasets were largely overlapping (Fig S5). To assess statistical significance of heritability estimates, we performed the resampling procedure but randomly permuted the trait values across breeds at each iteration. Permutation tests revealed that the observed heritability estimate was higher than that from the permuted data in all iterations of this procedure (p < 0.001 for all traits in both datasets).

### Clustering

Hierarchical clustering was performed using breed average scores for all individual C-BARQ items (raw questionnaire item scores, rather than factor scores), including breeds with at least 100 individuals of complete C-BARQ data. We performed hierarchical clustering on the behavioral distance matrix using the median-linkage method, with the resulting dendrogram aligned against the cladogram of breeds from Parker et al. (21) using a stepwise branch rotation procedure in the dendextend R package (46). We compared the structure of the genetic and behavioral dendrograms using cophenetic correlation. We assessed the significance of this correlation using a permutation test (1k iterations) to simulate the null distribution.

### Genome-wide association study (GWAS)

We identified single nucleotide polymorphisms (SNPs) associated with each trait using EMMA (24) predicting breed-average behavioral traits as a function of breed-average allele frequency and a polynomial term for the log of breed-average body weight. EMMA models were fit using the EMMREML R package (47). We included a 2^nd^ order polynomial term for log body weight because preliminary analyses revealed non-linear associations between breed-average weight and behavior that were best described using a polynomial term. We only included SNPs with a median minor allele frequency across breeds ≥ 0. 05 (Hayward et al.: 127,970 SNPs; Parker et al. 110,096 SNPs). GWAS analyses were conducted separately with each genetic dataset, and the resulting p values were combined across datasets using meta-analysis (Fisher’s method), first at the level of the SNP (109,780 overlapping SNPs), and then at the level of the gene. Combined p values were Bonferroni corrected for identification of genes associated with behavioral traits, and false discovery rate (FDR) corrected for enrichment analyses (48). Our primary analyses were restricted to SNPs located in genes (Table S3), but we report supplemental analyses that associated SNPs with the nearest gene within 20kb (Table S7 (49)). To derive gene-level p values from the GWAS, we combined (FDR corrected) p values from multiple SNPs associated with the same gene using meta-analysis (Fisher’s method).

### Gene Ontology Enrichment Analyses

Enrichment analyses were conducted using gene ontology (GO), and tests of tissue-specific gene expression. GO enrichment analyses were conducted using a Fisher Exact Test and the ‘weight01’ algorithm in the topGO R package (50), with ENSEMBL gene identifiers mapped to GO terms using the biomaRt R package (51, 52). Genes were included as significant if the FDR corrected p value for the gene was ≤ 0.05. Our primary GO analyses were restricted to gene-level p values derived from meta-analysis of SNPs in the gene (Table S6). The results of complementary analyses using SNPs within 20kb of the nearest gene are shown in Table S8. The network structure for GO terms in Figure 3 was generated using Resnik’s similarity measure, as implemented in NaviGO (53).

### Tissue enrichment

Tissue enrichment was assessed using the TissueEnrich R package (54). Data on tissue-specific gene expression in dogs were compiled from a study of gene expression in 10 tissues from a sample of four dogs (36). Microarray expression data was averaged across dogs, and across probes in cases where genes were mapped to more than one probe. Tissue-specific genes were defined using the algorithm from Uhlén et al. (37), and a threshold of 3-fold higher expression in a particular tissue as a cutoff for classification as tissue-specific. Enrichment tests were corrected for multiple hypothesis testing using the method described by Benjamini & Hochberg (48). The results of tissue-specific enrichment analyses using SNPs mapped to the nearest gene within 20kb are shown in Figure S6.

## Acknowledgements

We thank X. Zhou for helpful comments regarding analysis and A. Boyko and H. Parker for discussion of data from their labs.

## Author contributions

ELM and NSM analyzed the data. ELM, NSM, BMV and JAS wrote the paper. The authors declare no competing interests. C-BARQ data are available from and genetic data used in these analyses are available from https://datadryad.org/resource/doi:10.5061/dryad.266k4 and GEO accession numbers: GSE90441, GSE83160, GSE70454, and GSE96736.

