## supplemental information for "Highly Heritable and Functionally Relevant Breed Differences in Dog Behavior"

### Supplementary Materials

Supplemental Figures S1-S5

Supplemental Tables S1-S8

Supplemental References

Fig S1. Scatterplot of regression coefficients for each SNP across genetic datasets, by trait.

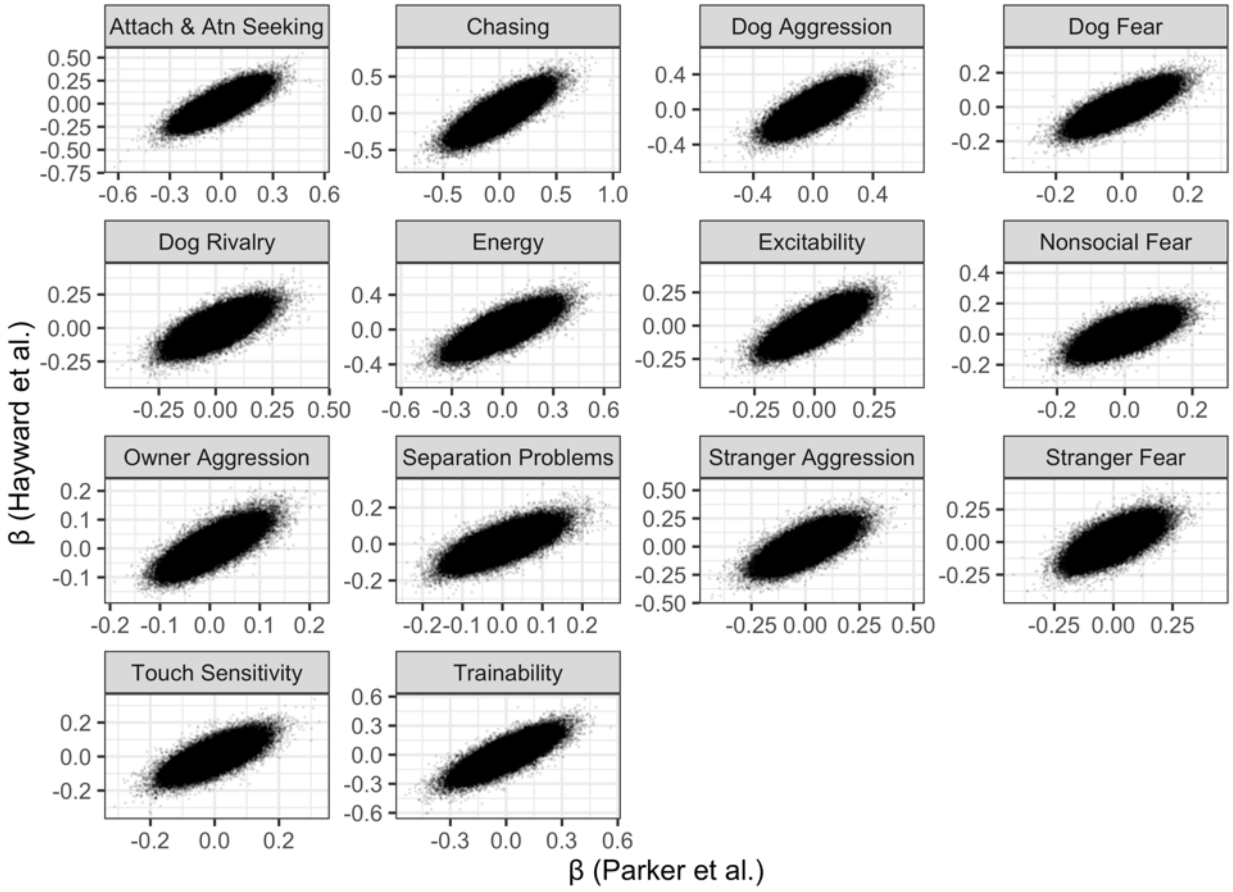

Fig S2. Correlation of SNP regression coefficients across genetic datasets as a function of p value threshold (from meta-analysis of p values) for inclusion. Points reflect the mean and error bars extend to the minimum and maximum values across traits.

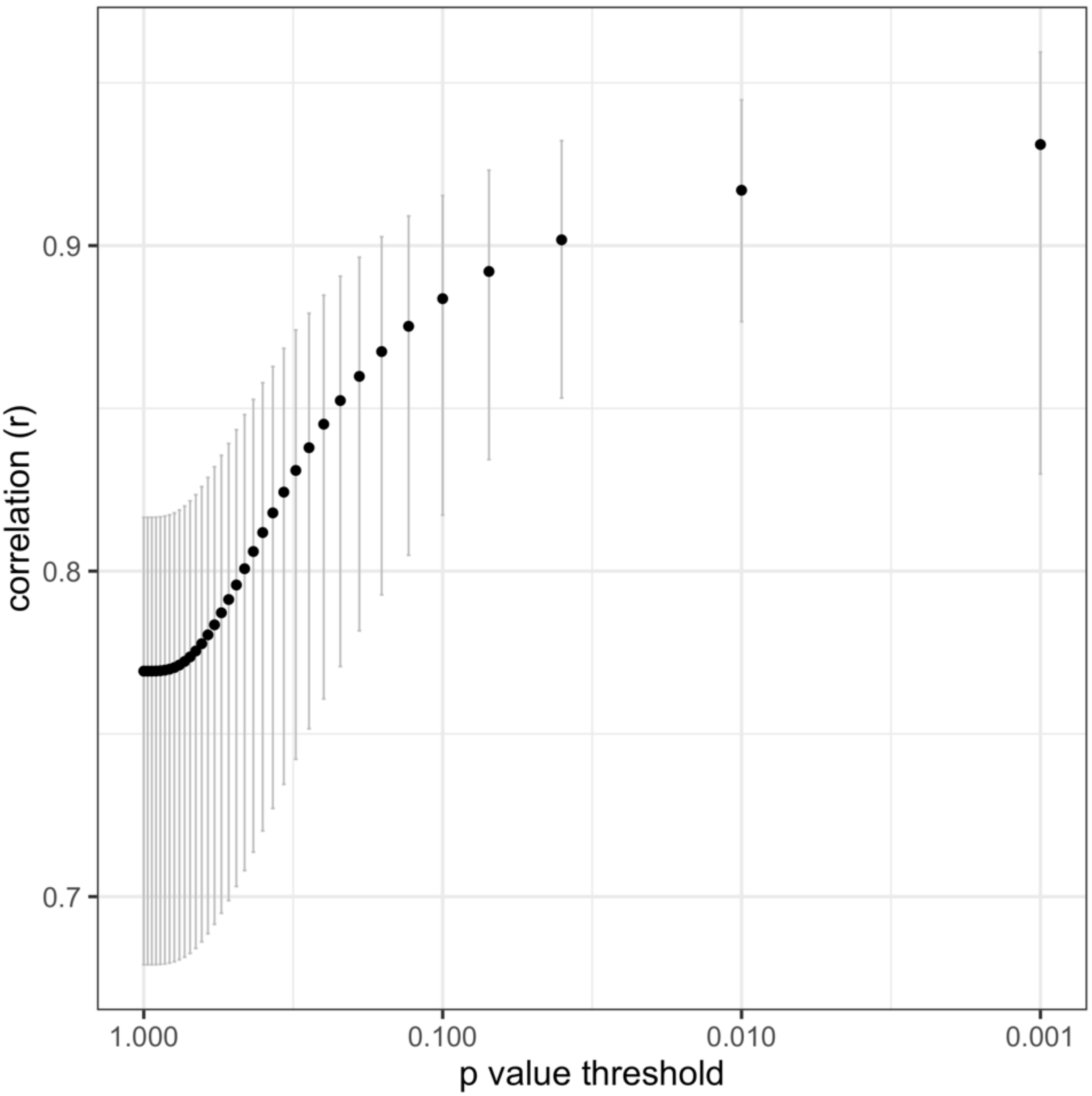

**Figure S3.** Tissue-specific enrichment for the collective set of genes associated with breed differences in dog behavior, mapping SNPs to the nearest gene within 20kb. (A) Enrichment tests using dog gene expression to identify tissue-specific genes. (B) Enrichment tests using human gene expression to identify tissue-specific genes. Bars reflect the -log10 p value from a hypergeometric test for tissue-specific gene enrichment, corrected for multiple comparisons. The dashed line indicates -log10(p = 0.05) and the results for brain tissue are highlighted in red.

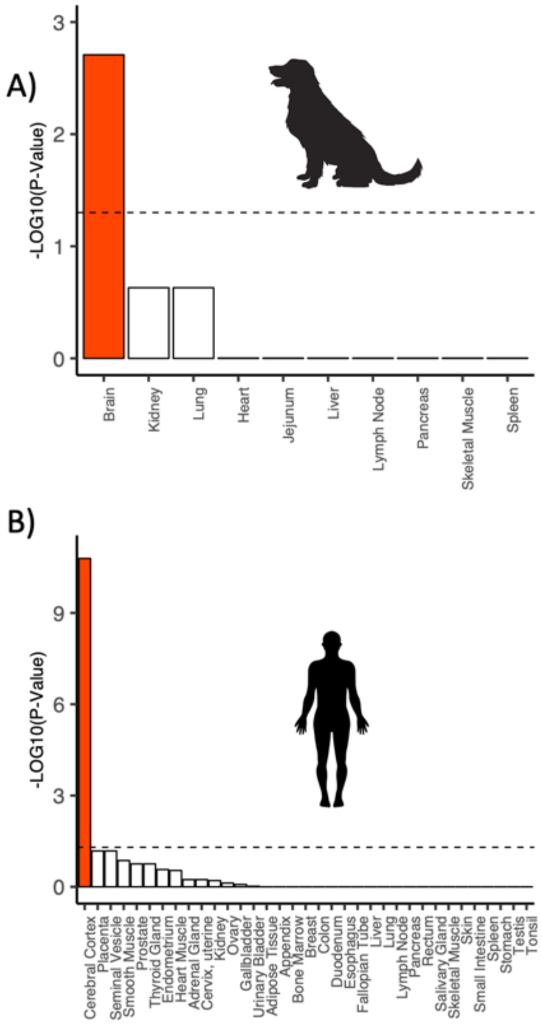

**Figure S4.** Trait heritability estimates using Efficient Mixed Model Association (EMMA) and a Bayesian implementation of the 'Animal Model' (MCMC). Heritability estimates from EMMA and MCMC were highly correlated within each of the two genetic datasets (Hayward et al., Parker et al.). Across genetic datasets, the heritability estimates for each trait were also highly correlated using both statistical approaches (MCMC: R = 0.81; EMMA: R = 0.94).

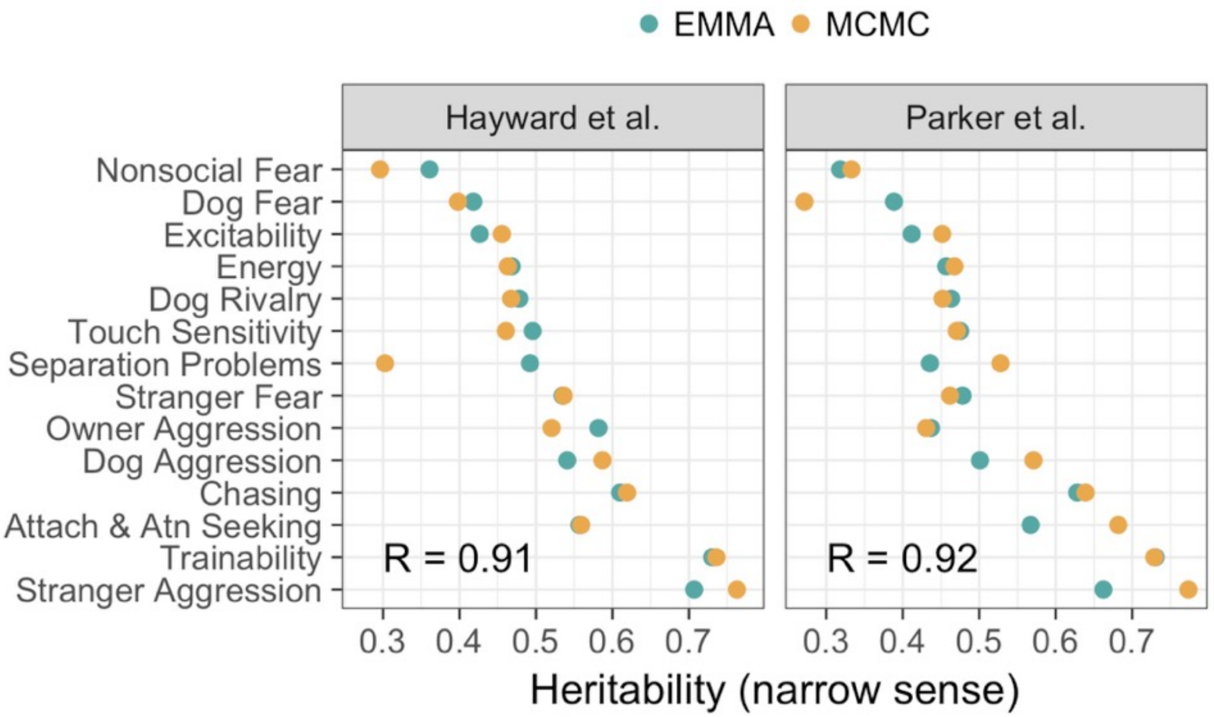

**Figure S5.** Distributions of heritability estimates from resampling using Efficient Mixed-Model Association (EMMA).

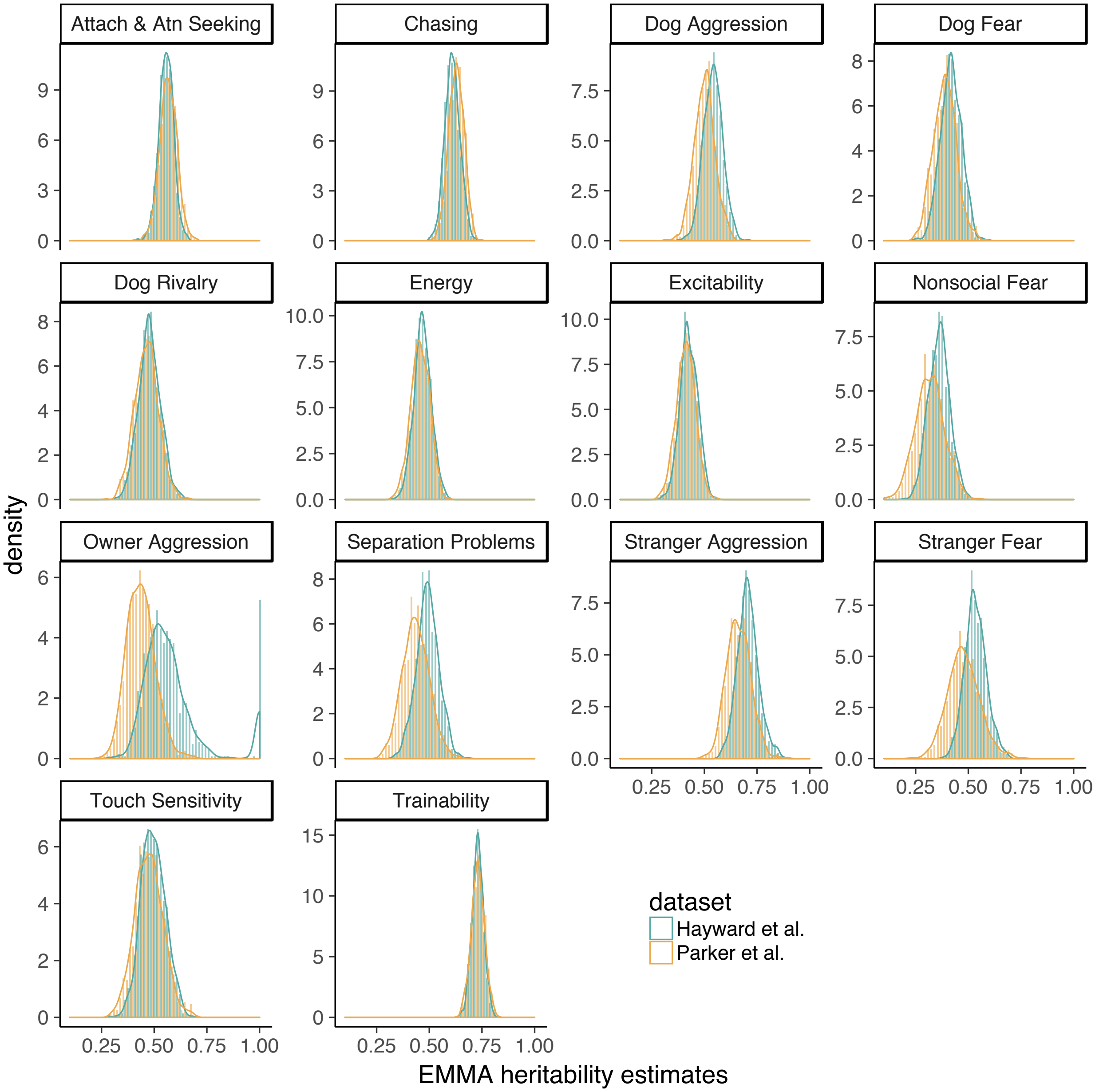

Table S1. Number of individuals per breed in the behavioral data.

| *Breed* | *N* |
|---|---|
| Airedale Terrier | 110 |
| Akita | 130 |
| Alaskan Malamute | 67 |
| American Eskimo Dog | 34 |
| American Pit Bull Terrier | 288 |
| American Staffordshire Terrier | 101 |
| Australian Cattle Dog | 276 |
| Australian Shepherd | 488 |
| Basenji | 33 |
| Basset Hound | 63 |
| Beagle | 192 |
| Bearded Collie | 35 |
| Belgian Malinois | 120 |
| Belgian Sheepdog | 46 |
| Belgian Tervuren | 114 |
| Bernese Mountain Dog | 174 |
| Bichon Frise | 107 |
| Border Collie | 597 |
| Border Terrier | 61 |
| Borzoi | 43 |
| Boston Terrier | 90 |
| Bouvier des Flandres | 32 |
| Boxer | 231 |
| Brittany | 99 |
| Bull Terrier | 40 |
| Bulldog | 52 |

| *Breed* | *N* |
|---|---|
| Bullmastiff | 76 |
| Cairn Terrier | 60 |
| Cardigan Welsh Corgi | 40 |
| Cavalier King Charles Spaniel | 93 |
| Chesapeake Bay Retriever | 35 |
| Chihuahua | 252 |
| Chinese Crested | 32 |
| Chinook | 64 |
| Chow Chow | 54 |
| Cocker Spaniel (American) | 160 |
| Cocker Spaniel (English) | 136 |
| Collie | 205 |
| Dachshund | 150 |
| Dachshund (Miniature) | 85 |
| Dalmatian | 85 |
| Doberman Pinscher | 297 |
| English Setter | 66 |
| English Springer Spaniel | 153 |
| Eurasier | 40 |
| Flat-Coated Retriever | 63 |
| French Bulldog | 56 |
| German Shepherd | 829 |
| German Shorthaired Pointer | 86 |
| German Wirehaired Pointer | 27 |
| Golden Retriever | 701 |
| Gordon Setter | 26 |
| Great Dane | 145 |

| *Breed* | *N* |
|---:|---|
| Great Pyrenees | 71 |
| Greyhound | 112 |
| Havanese | 105 |
| Irish Setter | 55 |
| Irish Wolfhound | 44 |
| Italian Greyhound | 32 |
| Jack Russell Terrier | 252 |
| Keeshond | 29 |
| Labrador Retriever | 1300 |
| Lhasa Apso | 35 |
| Maltese | 99 |
| Mastiff (English) | 152 |
| Miniature Pinscher | 82 |
| Miniature Schnauzer | 134 |
| Newfoundland | 236 |
| Norwegian Elkhound | 27 |
| Nova Scotia Duck Tolling Retriever | 67 |
| Papillon | 85 |
| Pekingese | 30 |
| Pembroke Welsh Corgi | 109 |
| Pomeranian | 112 |
| Poodle (Miniature) | 115 |
| Poodle (Standard) | 300 |
| Poodle (Toy) | 58 |
| Portuguese Water Dog | 104 |
| Pug | 121 |
| Rat Terrier | 87 |

| *Breed* | *N* |
|---:|---:|
| Redbone Coonhound | 30 |
| Rhodesian Ridgeback | 141 |
| Rottweiler | 392 |
| Saint Bernard | 41 |
| Samoyed | 37 |
| Shetland Sheepdog | 257 |
| Shiba Inu | 149 |
| Shih Tzu | 133 |
| Siberian Husky | 200 |
| Soft Coated Wheaten Terrier | 232 |
| Staffordshire Bull Terrier | 103 |
| Standard Schnauzer | 44 |
| Tibetan Terrier | 29 |
| Vizsla | 76 |
| Weimaraner | 99 |
| West Highland White Terrier | 79 |
| Whippet | 150 |
| Yorkshire Terrier | 123 |
| Anatolian Shepherd | 26 |
| Australian Kelpie | 57 |
| Parson Russell Terrier | 30 |

**Table S2.** Number of significant SNP associations (Bonferroni p ≤ 0.05), the top SNP and associated gene, and proportion variance explained (PVE) by the top SNP from the GWAS of dog behavioral traits.

| *Trait* | *Total Associations* | *Top SNP* | *Gene (top SNP)* | *Ensembl gene (top SNP)* | *PVE (Hayward et al.)* | *PVE (Parker et al.)* |
|---|---|---|---|---|---|---|
| Attach & Atn Seeking | 34 | chr34_19778169 | *MASP1* | ENSCAFG00000013838 | 0.19 | 0.18 |
| Chasing | 60 | chr32_4513202 | *FGF5* | ENSCAFG00000008886 | 0.20 | 0.21 |
| Dog Aggression | 27 | chr20_29700107 | *[novel gene]* | ENSCAFG00000035599 | 0.18 | 0.25 |
| Dog Fear | 20 | chr1_96469867 | *[novel gene]* | ENSCAFG00000036506 | 0.06 | 0.11 |
| Dog Rivalry | 23 | chr10_43493767 | *[novel gene]* | ENSCAFG00000031628 | 0.11 | 0.21 |
| Energy | 19 | chr38_1286873 | *[novel gene]* | ENSCAFG00000033514 | 0.06 | 0.20 |
| Excitability | 23 | chr9_52449334 | *RAPGEF1* | ENSCAFG00000019902 | 0.13 | 0.13 |
| Nonsocial Fear | 6 | chr8_40424847 | *[novel gene]* | ENSCAFG00000035815 | 0.17 | 0.10 |
| Owner Aggression | 32 | chr18_26274094 | *TTC17* | ENSCAFG00000006730 | 0.15 | 0.14 |
| Separation Problems | 4 | chr3_9599516 | *[novel gene]* | ENSCAFG00000039437 | 0.12 | 0.09 |
| Stranger Aggression | 22 | chr1_81065940 | *GNA14* | ENSCAFG00000031429 | 0.18 | 0.12 |
| Stranger Fear | 7 | chr1_81065940 | *GNA14* | ENSCAFG00000031429 | 0.10 | 0.12 |
| Touch Sensitivity | 20 | chr5_28225323 | *[novel gene]* | ENSCAFG00000040905 | 0.07 | 0.11 |
| Trainability | 39 | chr31_2974937 | *[novel gene]* | ENSCAFG00000033559 | 0.19 | 0.25 |

Table S3. Genes associated with behavioral traits at $p \leq 0.05$ after Bonferroni correction, using SNPs in the gene to derive gene-level p values (meta-analysis, Fisher's method).

| *Behavior* | *Ensemble ID* | *Gene Name* | *p* |
|---|---|---|---|
| Attach & Atn Seeking | ENSCAFG00000013221 | | < 0.00001 |
| Attach & Atn Seeking | ENSCAFG00000023562 | *DMD* | < 0.00001 |
| Attach & Atn Seeking | ENSCAFG00000000353 | *STXBP5* | < 0.00001 |
| Attach & Atn Seeking | ENSCAFG00000018100 | *SCAPER* | < 0.00001 |
| Attach & Atn Seeking | ENSCAFG00000013297 | *IGF2BP2* | < 0.00001 |
| Attach & Atn Seeking | ENSCAFG00000005914 | *TMTC2* | < 0.00001 |
| Attach & Atn Seeking | ENSCAFG00000028388 | *RF00322* | < 0.00001 |
| Attach & Atn Seeking | ENSCAFG00000018577 | *XIAP* | < 0.00001 |
| Attach & Atn Seeking | ENSCAFG00000006625 | *SFSWAP* | < 0.00001 |
| Attach & Atn Seeking | ENSCAFG00000000678 | *ZFPM2* | < 0.00001 |
| Attach & Atn Seeking | ENSCAFG00000000345 | *SRGAP1* | < 0.00001 |
| Attach & Atn Seeking | ENSCAFG00000007301 | *CWC27* | < 0.00001 |
| Attach & Atn Seeking | ENSCAFG00000000677 | *LRP12* | < 0.00001 |
| Attach & Atn Seeking | ENSCAFG00000002271 | *PHF14* | < 0.00001 |
| Attach & Atn Seeking | ENSCAFG00000007500 | *SBF2* | < 0.00001 |
| Attach & Atn Seeking | ENSCAFG00000007380 | *STAG1* | < 0.00001 |
| Attach & Atn Seeking | ENSCAFG00000026031 | *RF00026* | < 0.00001 |
| Attach & Atn Seeking | ENSCAFG00000003988 | *SPATA5* | < 0.00001 |
| Attach & Atn Seeking | ENSCAFG00000033690 | | < 0.00001 |
| Attach & Atn Seeking | ENSCAFG00000012413 | *RPS6KC1* | < 0.00001 |
| Attach & Atn Seeking | ENSCAFG00000033993 | | < 0.00001 |
| Attach & Atn Seeking | ENSCAFG00000000945 | *MAN1A1* | < 0.00001 |
| Attach & Atn Seeking | ENSCAFG00000001859 | *TEX47* | < 0.00001 |
| Attach & Atn Seeking | ENSCAFG00000010384 | *DNAH7* | < 0.00001 |
| Attach & Atn Seeking | ENSCAFG00000019030 | *ABAT* | < 0.00001 |

| | | | |
|---|---|---|---|
| Attach & Atn Seeking | ENSCAFG00000003678 | *CCNY* | < 0.00001 |
| Attach & Atn Seeking | ENSCAFG00000038165 | | < 0.00001 |
| Attach & Atn Seeking | ENSCAFG00000013823 | *SPECC1L* | < 0.00001 |
| Attach & Atn Seeking | ENSCAFG00000027967 | *RF00009* | < 0.00001 |
| Attach & Atn Seeking | ENSCAFG00000006743 | *PCMTD1* | < 0.00001 |
| Attach & Atn Seeking | ENSCAFG00000017716 | *EEF2K* | < 0.00001 |
| Attach & Atn Seeking | ENSCAFG00000000090 | *MC4R* | < 0.00001 |
| Attach & Atn Seeking | ENSCAFG00000018858 | *TARS* | < 0.00001 |
| Attach & Atn Seeking | ENSCAFG00000000157 | *DCC* | < 0.00001 |
| Attach & Atn Seeking | ENSCAFG00000031499 | *GLIS3* | < 0.00001 |
| Attach & Atn Seeking | ENSCAFG00000034328 | | < 0.00001 |
| Attach & Atn Seeking | ENSCAFG00000027973 | *RF00100* | 0.00001 |
| Attach & Atn Seeking | ENSCAFG00000003570 | *FBXW2* | 0.00001 |
| Attach & Atn Seeking | ENSCAFG00000016354 | *STIM2* | 0.00001 |
| Attach & Atn Seeking | ENSCAFG00000003322 | *IFRD1* | 0.00001 |
| Attach & Atn Seeking | ENSCAFG00000013852 | *MATN2* | 0.00001 |
| Attach & Atn Seeking | ENSCAFG00000040856 | | 0.00001 |
| Attach & Atn Seeking | ENSCAFG00000001700 | *SND1* | 0.00001 |
| Attach & Atn Seeking | ENSCAFG00000029666 | *HNRNPLL* | 0.00003 |
| Attach & Atn Seeking | ENSCAFG00000038450 | | 0.00004 |
| Attach & Atn Seeking | ENSCAFG00000000279 | *REPS1* | 0.00006 |
| Attach & Atn Seeking | ENSCAFG00000009207 | *AP3B1* | 0.00006 |
| Attach & Atn Seeking | ENSCAFG00000034522 | | 0.00010 |
| Attach & Atn Seeking | ENSCAFG00000005192 | *ACER3* | 0.00014 |
| Attach & Atn Seeking | ENSCAFG00000003701 | *ATG5* | 0.00015 |
| Attach & Atn Seeking | ENSCAFG00000009390 | *PPP3CC* | 0.00017 |
| Attach & Atn Seeking | ENSCAFG00000032548 | *KCNH5* | 0.00026 |
| Attach & Atn Seeking | ENSCAFG00000000938 | *MCM9* | 0.00028 |
| Attach & Atn Seeking | ENSCAFG00000005326 | *UVRAG* | 0.00031 |

| | | | |
|---|---|---|---|
| Attach & Atn Seeking | ENSCAFG00000002152 | *IL1R2* | 0.00038 |
| Attach & Atn Seeking | ENSCAFG00000000361 | *TBC1D30* | 0.00049 |
| Attach & Atn Seeking | ENSCAFG00000001889 | *GABRB1* | 0.00059 |
| Attach & Atn Seeking | ENSCAFG00000008056 | *RYR3* | 0.00071 |
| Attach & Atn Seeking | ENSCAFG00000017419 | *DACH2* | 0.00076 |
| Attach & Atn Seeking | ENSCAFG00000018564 | *GRIA3* | 0.00116 |
| Attach & Atn Seeking | ENSCAFG00000034953 | | 0.00128 |
| Attach & Atn Seeking | ENSCAFG00000030771 | *LCLAT1* | 0.00132 |
| Attach & Atn Seeking | ENSCAFG00000008525 | *CACNA1D* | 0.00135 |
| Attach & Atn Seeking | ENSCAFG00000018267 | *TOP3A* | 0.00149 |
| Attach & Atn Seeking | ENSCAFG00000033697 | | 0.00149 |
| Attach & Atn Seeking | ENSCAFG00000009088 | *NUP93* | 0.00179 |
| Attach & Atn Seeking | ENSCAFG00000001852 | *ADAM22* | 0.00203 |
| Attach & Atn Seeking | ENSCAFG00000029926 | | 0.00212 |
| Attach & Atn Seeking | ENSCAFG00000034741 | | 0.00214 |
| Attach & Atn Seeking | ENSCAFG00000007963 | *BLK* | 0.00219 |
| Attach & Atn Seeking | ENSCAFG00000006266 | *ARHGAP26* | 0.00253 |
| Attach & Atn Seeking | ENSCAFG00000033544 | | 0.00271 |
| Attach & Atn Seeking | ENSCAFG00000003737 | *PARD3* | 0.00278 |
| Attach & Atn Seeking | ENSCAFG00000008568 | *AGPAT5* | 0.00328 |
| Attach & Atn Seeking | ENSCAFG00000036531 | | 0.00344 |
| Attach & Atn Seeking | ENSCAFG00000030012 | | 0.00354 |
| Attach & Atn Seeking | ENSCAFG00000004358 | *IQSEC1* | 0.00363 |
| Attach & Atn Seeking | ENSCAFG00000032853 | | 0.00397 |
| Attach & Atn Seeking | ENSCAFG00000000935 | *CEP85L* | 0.00428 |
| Attach & Atn Seeking | ENSCAFG00000009609 | *TMEM117* | 0.00477 |
| Attach & Atn Seeking | ENSCAFG00000005766 | *SYT1* | 0.00482 |
| Attach & Atn Seeking | ENSCAFG00000013838 | *MASP1* | 0.00555 |
| Attach & Atn Seeking | ENSCAFG00000010140 | *BICDL1* | 0.00638 |

| | | | |
|---|---|---|---|
| Attach & Atn Seeking | ENSCAFG00000034873 | | 0.00661 |
| Attach & Atn Seeking | ENSCAFG00000033071 | | 0.00680 |
| Attach & Atn Seeking | ENSCAFG00000011125 | *RAB3GAP2* | 0.00705 |
| Attach & Atn Seeking | ENSCAFG00000004182 | *ZC3H4* | 0.00729 |
| Attach & Atn Seeking | ENSCAFG00000027573 | | 0.00816 |
| Attach & Atn Seeking | ENSCAFG00000039830 | | 0.00931 |
| Attach & Atn Seeking | ENSCAFG00000000460 | *LAPTM4B* | 0.01114 |
| Attach & Atn Seeking | ENSCAFG00000039541 | | 0.01243 |
| Attach & Atn Seeking | ENSCAFG00000005720 | *CLPB* | 0.01280 |
| Attach & Atn Seeking | ENSCAFG00000003996 | *FGF2* | 0.01447 |
| Attach & Atn Seeking | ENSCAFG00000000828 | *RPS6KA2* | 0.01483 |
| Attach & Atn Seeking | ENSCAFG00000035717 | | 0.01493 |
| Attach & Atn Seeking | ENSCAFG00000025075 | *DEFB1* | 0.01533 |
| Attach & Atn Seeking | ENSCAFG00000003651 | *CSMD2* | 0.01736 |
| Attach & Atn Seeking | ENSCAFG00000039575 | | 0.01821 |
| Attach & Atn Seeking | ENSCAFG00000014707 | *CDC42* | 0.01971 |
| Attach & Atn Seeking | ENSCAFG00000005054 | *ZRANB3* | 0.02275 |
| Attach & Atn Seeking | ENSCAFG00000029250 | *CDK13* | 0.02615 |
| Attach & Atn Seeking | ENSCAFG00000005870 | *RASGRP3* | 0.02720 |
| Attach & Atn Seeking | ENSCAFG00000007780 | *HTR1F* | 0.02916 |
| Attach & Atn Seeking | ENSCAFG00000006326 | *KCTD16* | 0.03037 |
| Attach & Atn Seeking | ENSCAFG00000003334 | *ABCA13* | 0.03099 |
| Attach & Atn Seeking | ENSCAFG00000004254 | *FAAH* | 0.03211 |
| Attach & Atn Seeking | ENSCAFG00000040232 | | 0.03219 |
| Attach & Atn Seeking | ENSCAFG00000001930 | *PGM5* | 0.03302 |
| Attach & Atn Seeking | ENSCAFG00000033154 | | 0.03426 |
| Attach & Atn Seeking | ENSCAFG00000007770 | *LNPEP* | 0.03439 |
| Attach & Atn Seeking | ENSCAFG00000004406 | *FBLN2* | 0.03490 |
| Attach & Atn Seeking | ENSCAFG00000008501 | *PPEF2* | 0.03495 |

| | | | |
|---|---|---|---|
| Attach & Atn Seeking | ENSCAFG00000009840 | *FAM13A* | 0.03527 |
| Attach & Atn Seeking | ENSCAFG00000035923 | | 0.03853 |
| Attach & Atn Seeking | ENSCAFG00000010811 | *LRRC28* | 0.04488 |
| Chasing | ENSCAFG00000008141 | *FBXW7* | < 0.00001 |
| Chasing | ENSCAFG00000003554 | *ASCC3* | < 0.00001 |
| Chasing | ENSCAFG00000005998 | *LHFPL6* | < 0.00001 |
| Chasing | ENSCAFG00000000670 | *RIMS2* | < 0.00001 |
| Chasing | ENSCAFG00000018720 | *ATG4C* | < 0.00001 |
| Chasing | ENSCAFG00000005849 | *TBC1D5* | < 0.00001 |
| Chasing | ENSCAFG00000006568 | *ARHGAP22* | < 0.00001 |
| Chasing | ENSCAFG00000014845 | *FAM193A* | < 0.00001 |
| Chasing | ENSCAFG00000039364 | | < 0.00001 |
| Chasing | ENSCAFG00000008886 | *FGF5* | < 0.00001 |
| Chasing | ENSCAFG00000002155 | | < 0.00001 |
| Chasing | ENSCAFG00000030067 | | < 0.00001 |
| Chasing | ENSCAFG00000033197 | | < 0.00001 |
| Chasing | ENSCAFG00000003553 | *CADPS2* | < 0.00001 |
| Chasing | ENSCAFG00000035873 | | < 0.00001 |
| Chasing | ENSCAFG00000018081 | *SPECC1* | < 0.00001 |
| Chasing | ENSCAFG00000011773 | *C6H7orf50* | < 0.00001 |
| Chasing | ENSCAFG00000010354 | *NT5C2* | < 0.00001 |
| Chasing | ENSCAFG00000032853 | | < 0.00001 |
| Chasing | ENSCAFG00000008160 | | < 0.00001 |
| Chasing | ENSCAFG00000000677 | *LRP12* | < 0.00001 |
| Chasing | ENSCAFG00000034837 | | < 0.00001 |
| Chasing | ENSCAFG00000034827 | | < 0.00001 |
| Chasing | ENSCAFG00000003580 | *GRIK2* | < 0.00001 |
| Chasing | ENSCAFG00000002316 | *MLIP* | < 0.00001 |
| Chasing | ENSCAFG00000030939 | | < 0.00001 |

| | | | |
|---|---|---|---|
| Chasing | ENSCAFG00000018181 | | < 0.00001 |
| Chasing | ENSCAFG00000002083 | | < 0.00001 |
| Chasing | ENSCAFG00000002562 | *COL19A1* | < 0.00001 |
| Chasing | ENSCAFG00000026776 | | < 0.00001 |
| Chasing | ENSCAFG00000014682 | *HTT* | < 0.00001 |
| Chasing | ENSCAFG00000014403 | *SORCS2* | < 0.00001 |
| Chasing | ENSCAFG00000014975 | *EIF4G3* | < 0.00001 |
| Chasing | ENSCAFG00000004986 | *GDPD4* | 0.00001 |
| Chasing | ENSCAFG00000000770 | | 0.00001 |
| Chasing | ENSCAFG00000031643 | | 0.00001 |
| Chasing | ENSCAFG00000037340 | | 0.00002 |
| Chasing | ENSCAFG00000026119 | *RF00026* | 0.00002 |
| Chasing | ENSCAFG00000018570 | *SGIP1* | 0.00003 |
| Chasing | ENSCAFG00000006920 | *KCNIP3* | 0.00003 |
| Chasing | ENSCAFG00000038858 | | 0.00004 |
| Chasing | ENSCAFG00000009831 | *PRMT3* | 0.00005 |
| Chasing | ENSCAFG00000019237 | *ADCY9* | 0.00008 |
| Chasing | ENSCAFG00000004817 | *ELF1* | 0.00009 |
| Chasing | ENSCAFG00000026661 | | 0.00010 |
| Chasing | ENSCAFG00000015345 | *CC2D2A* | 0.00011 |
| Chasing | ENSCAFG00000035608 | | 0.00011 |
| Chasing | ENSCAFG00000001801 | *TMC1* | 0.00012 |
| Chasing | ENSCAFG00000010034 | *RYR2* | 0.00015 |
| Chasing | ENSCAFG00000034984 | | 0.00017 |
| Chasing | ENSCAFG00000031499 | *GLIS3* | 0.00018 |
| Chasing | ENSCAFG00000031408 | *CFAP299* | 0.00020 |
| Chasing | ENSCAFG00000007246 | *CYP7B1* | 0.00023 |
| Chasing | ENSCAFG00000038312 | | 0.00024 |
| Chasing | ENSCAFG00000009924 | *TAOK3* | 0.00028 |

| | | | |
|---|---|---|---|
| Chasing | ENSCAFG00000013779 | | 0.00030 |
| Chasing | ENSCAFG00000018858 | *TARS* | 0.00031 |
| Chasing | ENSCAFG00000010744 | *NAV1* | 0.00036 |
| Chasing | ENSCAFG00000034717 | | 0.00038 |
| Chasing | ENSCAFG00000000459 | *TBC1D15* | 0.00041 |
| Chasing | ENSCAFG00000035147 | | 0.00042 |
| Chasing | ENSCAFG00000020373 | *USP33* | 0.00049 |
| Chasing | ENSCAFG00000039762 | | 0.00052 |
| Chasing | ENSCAFG00000040770 | | 0.00057 |
| Chasing | ENSCAFG00000009828 | *SPATA19* | 0.00058 |
| Chasing | ENSCAFG00000003176 | *DPY19L2* | 0.00061 |
| Chasing | ENSCAFG00000006694 | *STX2* | 0.00066 |
| Chasing | ENSCAFG00000018418 | *RHBDL3* | 0.00073 |
| Chasing | ENSCAFG00000007152 | *PTPRG* | 0.00077 |
| Chasing | ENSCAFG00000033124 | | 0.00091 |
| Chasing | ENSCAFG00000034423 | | 0.00096 |
| Chasing | ENSCAFG00000000353 | *STXBP5* | 0.00103 |
| Chasing | ENSCAFG00000006691 | *SLC46A3* | 0.00111 |
| Chasing | ENSCAFG00000029135 | *GDNF* | 0.00143 |
| Chasing | ENSCAFG00000007657 | *ATOH8* | 0.00146 |
| Chasing | ENSCAFG00000022461 | *RF00100* | 0.00151 |
| Chasing | ENSCAFG00000002081 | *CDC37L1* | 0.00159 |
| Chasing | ENSCAFG00000003927 | *ELAVL4* | 0.00189 |
| Chasing | ENSCAFG00000000011 | *NFATC1* | 0.00204 |
| Chasing | ENSCAFG00000007009 | *TMEM68* | 0.00241 |
| Chasing | ENSCAFG00000023764 | | 0.00252 |
| Chasing | ENSCAFG00000004406 | *FBLN2* | 0.00254 |
| Chasing | ENSCAFG00000010822 | *ACTN2* | 0.00271 |
| Chasing | ENSCAFG00000015800 | *MYO5A* | 0.00272 |

| | | | |
|---|---|---|---|
| Chasing | ENSCAFG00000028798 | | 0.00285 |
| Chasing | ENSCAFG00000004535 | *NR2C2* | 0.00315 |
| Chasing | ENSCAFG00000030105 | *IL33* | 0.00321 |
| Chasing | ENSCAFG00000002941 | *ME1* | 0.00356 |
| Chasing | ENSCAFG00000006266 | *ARHGAP26* | 0.00359 |
| Chasing | ENSCAFG00000039479 | | 0.00491 |
| Chasing | ENSCAFG00000038915 | | 0.00513 |
| Chasing | ENSCAFG00000002271 | *PHF14* | 0.00515 |
| Chasing | ENSCAFG00000006293 | *NR3C1* | 0.00527 |
| Chasing | ENSCAFG00000016428 | *ARG2* | 0.00566 |
| Chasing | ENSCAFG00000008185 | *ARFIP1* | 0.00614 |
| Chasing | ENSCAFG00000013247 | *PRR14L* | 0.00785 |
| Chasing | ENSCAFG00000002300 | *KLHL31* | 0.00828 |
| Chasing | ENSCAFG00000008123 | *TENM3* | 0.00962 |
| Chasing | ENSCAFG00000009984 | *TBR1* | 0.00990 |
| Chasing | ENSCAFG00000038024 | | 0.01016 |
| Chasing | ENSCAFG00000017519 | *TNMD* | 0.01023 |
| Chasing | ENSCAFG00000034066 | | 0.01085 |
| Chasing | ENSCAFG00000007130 | *NSMAF* | 0.01224 |
| Chasing | ENSCAFG00000010000 | *C27H12orf40* | 0.01234 |
| Chasing | ENSCAFG00000031727 | *STC2* | 0.01277 |
| Chasing | ENSCAFG00000035216 | | 0.01302 |
| Chasing | ENSCAFG00000007822 | *BDP1* | 0.01428 |
| Chasing | ENSCAFG00000027923 | *RF01169* | 0.01457 |
| Chasing | ENSCAFG00000009881 | *ALAS1* | 0.01557 |
| Chasing | ENSCAFG00000011155 | *STARD9* | 0.01712 |
| Chasing | ENSCAFG00000026513 | | 0.01789 |
| Chasing | ENSCAFG00000039166 | | 0.01794 |
| Chasing | ENSCAFG00000012781 | *XPR1* | 0.01799 |

| | | | |
|---|---|---|---|
| Chasing | ENSCAFG00000003663 | *GSN* | 0.01802 |
| Chasing | ENSCAFG00000003778 | *AFG1L* | 0.01891 |
| Chasing | ENSCAFG00000006255 | *NBEA* | 0.02035 |
| Chasing | ENSCAFG00000034549 | | 0.02044 |
| Chasing | ENSCAFG00000017167 | *KCNN3* | 0.02259 |
| Chasing | ENSCAFG00000032263 | | 0.02359 |
| Chasing | ENSCAFG00000035717 | | 0.02446 |
| Chasing | ENSCAFG00000010114 | *CIT* | 0.03010 |
| Chasing | ENSCAFG00000013067 | *PAK2* | 0.03623 |
| Chasing | ENSCAFG00000040446 | | 0.03682 |
| Chasing | ENSCAFG00000033649 | | 0.03862 |
| Chasing | ENSCAFG00000006612 | *SUCLG2* | 0.03937 |
| Chasing | ENSCAFG00000040257 | | 0.03981 |
| Chasing | ENSCAFG00000010735 | *GRAMD1C* | 0.04285 |
| Chasing | ENSCAFG00000011688 | *TRUB1* | 0.04841 |
| Dog Aggression | ENSCAFG00000009924 | *TAOK3* | < 0.00001 |
| Dog Aggression | ENSCAFG00000023804 | *HERC3* | < 0.00001 |
| Dog Aggression | ENSCAFG00000009911 | *FBXW4* | < 0.00001 |
| Dog Aggression | ENSCAFG00000002155 | | < 0.00001 |
| Dog Aggression | ENSCAFG00000031429 | *GNA14* | < 0.00001 |
| Dog Aggression | ENSCAFG00000036640 | | < 0.00001 |
| Dog Aggression | ENSCAFG00000021629 | *RF00026* | < 0.00001 |
| Dog Aggression | ENSCAFG00000023460 | *FBRSL1* | < 0.00001 |
| Dog Aggression | ENSCAFG00000018100 | *SCAPER* | < 0.00001 |
| Dog Aggression | ENSCAFG00000018864 | *GPC3* | < 0.00001 |
| Dog Aggression | ENSCAFG00000024350 | *CENPP* | < 0.00001 |
| Dog Aggression | ENSCAFG00000001010 | *PPP2CA* | < 0.00001 |
| Dog Aggression | ENSCAFG00000039364 | | < 0.00001 |
| Dog Aggression | ENSCAFG00000001016 | *TRDN* | < 0.00001 |

| | | | |
|---|---|---|---|
| Dog Aggression | ENSCAFG00000020112 | *ABCD3* | < 0.00001 |
| Dog Aggression | ENSCAFG00000018850 | *HS6ST2* | < 0.00001 |
| Dog Aggression | ENSCAFG00000005998 | *LHFPL6* | < 0.00001 |
| Dog Aggression | ENSCAFG00000014222 | *IFT80* | < 0.00001 |
| Dog Aggression | ENSCAFG00000013004 | *CACNA1E* | < 0.00001 |
| Dog Aggression | ENSCAFG00000011936 | *TRPM8* | 0.00001 |
| Dog Aggression | ENSCAFG00000037915 | | 0.00001 |
| Dog Aggression | ENSCAFG00000018811 | | 0.00001 |
| Dog Aggression | ENSCAFG00000015679 | *RGS7* | 0.00001 |
| Dog Aggression | ENSCAFG00000000678 | *ZFPM2* | 0.00001 |
| Dog Aggression | ENSCAFG00000008431 | *C30H15orf41* | 0.00001 |
| Dog Aggression | ENSCAFG00000001935 | *AKAP9* | 0.00002 |
| Dog Aggression | ENSCAFG00000006196 | *CPNE4* | 0.00002 |
| Dog Aggression | ENSCAFG00000002364 | *ADGRL3* | 0.00003 |
| Dog Aggression | ENSCAFG00000010941 | *SLC24A4* | 0.00003 |
| Dog Aggression | ENSCAFG00000015803 | *SDCCAG8* | 0.00003 |
| Dog Aggression | ENSCAFG00000004872 | *PTPN4* | 0.00003 |
| Dog Aggression | ENSCAFG00000009842 | *OPCML* | 0.00010 |
| Dog Aggression | ENSCAFG00000006766 | *DLC1* | 0.00011 |
| Dog Aggression | ENSCAFG00000039974 | | 0.00013 |
| Dog Aggression | ENSCAFG00000010772 | *GAL* | 0.00024 |
| Dog Aggression | ENSCAFG00000000557 | *CSNK1G3* | 0.00029 |
| Dog Aggression | ENSCAFG00000007822 | *BDP1* | 0.00047 |
| Dog Aggression | ENSCAFG00000022438 | *RF00026* | 0.00061 |
| Dog Aggression | ENSCAFG00000014864 | *CASP12* | 0.00069 |
| Dog Aggression | ENSCAFG00000009771 | *TTC3* | 0.00071 |
| Dog Aggression | ENSCAFG00000014169 | | 0.00074 |
| Dog Aggression | ENSCAFG00000008557 | *APP* | 0.00090 |
| Dog Aggression | ENSCAFG00000015700 | *STX18* | 0.00113 |

| | | | |
|---|---|---|---|
| Dog Aggression | ENSCAFG00000027910 | *RF00026* | 0.00139 |
| Dog Aggression | ENSCAFG00000005809 | *TM9SF2* | 0.00152 |
| Dog Aggression | ENSCAFG00000031950 | *CARTPT* | 0.00166 |
| Dog Aggression | ENSCAFG00000007803 | *GALNTL6* | 0.00170 |
| Dog Aggression | ENSCAFG00000014845 | *FAM193A* | 0.00173 |
| Dog Aggression | ENSCAFG00000007364 | *KLKB1* | 0.00173 |
| Dog Aggression | ENSCAFG00000028102 | *RF00088* | 0.00179 |
| Dog Aggression | ENSCAFG00000033673 | | 0.00209 |
| Dog Aggression | ENSCAFG00000009831 | *PRMT3* | 0.00218 |
| Dog Aggression | ENSCAFG00000034987 | | 0.00256 |
| Dog Aggression | ENSCAFG00000018799 | *IGSF1* | 0.00260 |
| Dog Aggression | ENSCAFG00000019194 | *SLC43A2* | 0.00291 |
| Dog Aggression | ENSCAFG00000002390 | *FAM178B* | 0.00292 |
| Dog Aggression | ENSCAFG00000034449 | | 0.00335 |
| Dog Aggression | ENSCAFG00000003792 | | 0.00347 |
| Dog Aggression | ENSCAFG00000000079 | *RELCH* | 0.00350 |
| Dog Aggression | ENSCAFG00000028659 | | 0.00363 |
| Dog Aggression | ENSCAFG00000007857 | *SCRG1* | 0.00380 |
| Dog Aggression | ENSCAFG00000000239 | *PDE7B* | 0.00382 |
| Dog Aggression | ENSCAFG00000000279 | *REPS1* | 0.00459 |
| Dog Aggression | ENSCAFG00000009327 | *ADGRG7* | 0.00470 |
| Dog Aggression | ENSCAFG00000011544 | *AOX4* | 0.00552 |
| Dog Aggression | ENSCAFG00000030515 | *YPEL2* | 0.00625 |
| Dog Aggression | ENSCAFG00000004731 | *TENM4* | 0.00679 |
| Dog Aggression | ENSCAFG00000033796 | | 0.00682 |
| Dog Aggression | ENSCAFG00000034165 | | 0.00751 |
| Dog Aggression | ENSCAFG00000004282 | *WDFY2* | 0.00790 |
| Dog Aggression | ENSCAFG00000028870 | *RAB6A* | 0.00864 |
| Dog Aggression | ENSCAFG00000013733 | *GATM* | 0.00877 |

| | | | |
|---|---|---|---|
| Dog Aggression | ENSCAFG00000000935 | *CEP85L* | 0.00901 |
| Dog Aggression | ENSCAFG00000039775 | | 0.00909 |
| Dog Aggression | ENSCAFG00000035815 | | 0.01182 |
| Dog Aggression | ENSCAFG00000007251 | *KRT5* | 0.01230 |
| Dog Aggression | ENSCAFG00000040105 | | 0.01457 |
| Dog Aggression | ENSCAFG00000006952 | *DHX37* | 0.01766 |
| Dog Aggression | ENSCAFG00000004436 | *RB1* | 0.02029 |
| Dog Aggression | ENSCAFG00000016949 | *ALOXE3* | 0.02079 |
| Dog Aggression | ENSCAFG00000033281 | | 0.02096 |
| Dog Aggression | ENSCAFG00000011430 | *STXBP5L* | 0.02247 |
| Dog Aggression | ENSCAFG00000039623 | | 0.02262 |
| Dog Aggression | ENSCAFG00000020012 | *PKD1L2* | 0.02437 |
| Dog Aggression | ENSCAFG00000039115 | | 0.02559 |
| Dog Aggression | ENSCAFG00000015343 | *DCAF6* | 0.03217 |
| Dog Aggression | ENSCAFG00000033300 | | 0.03269 |
| Dog Aggression | ENSCAFG00000001355 | *KIAA2026* | 0.03358 |
| Dog Aggression | ENSCAFG00000023463 | *ITPR1* | 0.03564 |
| Dog Aggression | ENSCAFG00000005423 | *STT3B* | 0.04004 |
| Dog Aggression | ENSCAFG00000002323 | *MGAT4A* | 0.04155 |
| Dog Aggression | ENSCAFG00000026645 | | 0.04384 |
| Dog Aggression | ENSCAFG00000010766 | *TESMIN* | 0.04425 |
| Dog Aggression | ENSCAFG00000039224 | | 0.04808 |
| Dog Fear | ENSCAFG00000013297 | *IGF2BP2* | < 0.00001 |
| Dog Fear | ENSCAFG00000008827 | *ALMS1* | < 0.00001 |
| Dog Fear | ENSCAFG00000007218 | | < 0.00001 |
| Dog Fear | ENSCAFG00000002667 | | < 0.00001 |
| Dog Fear | ENSCAFG00000011430 | *STXBP5L* | < 0.00001 |
| Dog Fear | ENSCAFG00000009207 | *AP3B1* | < 0.00001 |
| Dog Fear | ENSCAFG00000005692 | *RPTOR* | < 0.00001 |

| | | | |
|---|---|---|---|
| Dog Fear | ENSCAFG00000002257 | *ICA1* | < 0.00001 |
| Dog Fear | ENSCAFG00000009242 | *FILIP1L* | < 0.00001 |
| Dog Fear | ENSCAFG00000028659 | | < 0.00001 |
| Dog Fear | ENSCAFG00000009390 | *PPP3CC* | < 0.00001 |
| Dog Fear | ENSCAFG00000040856 | | < 0.00001 |
| Dog Fear | ENSCAFG00000029740 | *MSRB3* | < 0.00001 |
| Dog Fear | ENSCAFG00000001002 | *SMPDL3A* | < 0.00001 |
| Dog Fear | ENSCAFG00000018100 | *SCAPER* | < 0.00001 |
| Dog Fear | ENSCAFG00000006592 | *STK32A* | < 0.00001 |
| Dog Fear | ENSCAFG00000005699 | *DNAJC18* | < 0.00001 |
| Dog Fear | ENSCAFG00000000334 | *ADAMTS2* | < 0.00001 |
| Dog Fear | ENSCAFG00000031408 | *CFAP299* | < 0.00001 |
| Dog Fear | ENSCAFG00000039641 | | < 0.00001 |
| Dog Fear | ENSCAFG00000002848 | *BCKDHB* | < 0.00001 |
| Dog Fear | ENSCAFG00000006248 | *CHL1* | < 0.00001 |
| Dog Fear | ENSCAFG00000013228 | *CNKSR2* | < 0.00001 |
| Dog Fear | ENSCAFG00000000945 | *MAN1A1* | < 0.00001 |
| Dog Fear | ENSCAFG00000003657 | *NBAS* | < 0.00001 |
| Dog Fear | ENSCAFG00000004436 | *RB1* | < 0.00001 |
| Dog Fear | ENSCAFG00000036994 | | < 0.00001 |
| Dog Fear | ENSCAFG00000012967 | *PCYT1A* | < 0.00001 |
| Dog Fear | ENSCAFG00000004586 | *RSU1* | < 0.00001 |
| Dog Fear | ENSCAFG00000012344 | *FASTKD1* | 0.00001 |
| Dog Fear | ENSCAFG00000001879 | *NUAK1* | 0.00001 |
| Dog Fear | ENSCAFG00000003373 | *DGKI* | 0.00002 |
| Dog Fear | ENSCAFG00000006743 | *PCMTD1* | 0.00004 |
| Dog Fear | ENSCAFG00000017453 | *INTS3* | 0.00004 |
| Dog Fear | ENSCAFG00000000932 | *SLC35F1* | 0.00006 |
| Dog Fear | ENSCAFG00000001105 | *KCNQ3* | 0.00006 |

| | | | |
|---|---|---|---|
| Dog Fear | ENSCAFG00000001277 | *ANKS1A* | 0.00013 |
| Dog Fear | ENSCAFG00000010811 | *LRRC28* | 0.00013 |
| Dog Fear | ENSCAFG00000016811 | *RHBG* | 0.00014 |
| Dog Fear | ENSCAFG00000035064 | | 0.00016 |
| Dog Fear | ENSCAFG00000009270 | *PEBP4* | 0.00019 |
| Dog Fear | ENSCAFG00000000557 | *CSNK1G3* | 0.00020 |
| Dog Fear | ENSCAFG00000035994 | | 0.00021 |
| Dog Fear | ENSCAFG00000008279 | *PTPRJ* | 0.00039 |
| Dog Fear | ENSCAFG00000037560 | | 0.00043 |
| Dog Fear | ENSCAFG00000004799 | *GBX1* | 0.00047 |
| Dog Fear | ENSCAFG00000015705 | *MSX1* | 0.00050 |
| Dog Fear | ENSCAFG00000001312 | *CHCHD3* | 0.00059 |
| Dog Fear | ENSCAFG00000015695 | *PRKCH* | 0.00060 |
| Dog Fear | ENSCAFG00000017298 | *MAP2K1* | 0.00061 |
| Dog Fear | ENSCAFG00000003334 | *ABCA13* | 0.00068 |
| Dog Fear | ENSCAFG00000017932 | *ARID3B* | 0.00082 |
| Dog Fear | ENSCAFG00000010127 | *SNRPN* | 0.00084 |
| Dog Fear | ENSCAFG00000033988 | | 0.00096 |
| Dog Fear | ENSCAFG00000015422 | *KIAA0753* | 0.00100 |
| Dog Fear | ENSCAFG00000008739 | *BOLA3* | 0.00105 |
| Dog Fear | ENSCAFG00000030592 | *RF00026* | 0.00106 |
| Dog Fear | ENSCAFG00000001646 | *PRUNE2* | 0.00109 |
| Dog Fear | ENSCAFG00000011756 | *PLXNA2* | 0.00121 |
| Dog Fear | ENSCAFG00000033910 | | 0.00122 |
| Dog Fear | ENSCAFG00000033403 | | 0.00126 |
| Dog Fear | ENSCAFG00000007691 | *GALNT18* | 0.00149 |
| Dog Fear | ENSCAFG00000032976 | | 0.00181 |
| Dog Fear | ENSCAFG00000013995 | *CABIN1* | 0.00219 |
| Dog Fear | ENSCAFG00000008370 | *RNF13* | 0.00254 |

| | | | |
|---|---|---|---|
| Dog Fear | ENSCAFG00000007780 | *HTR1F* | 0.00267 |
| Dog Fear | ENSCAFG00000017318 | *EBF1* | 0.00288 |
| Dog Fear | ENSCAFG00000015787 | *PPP2R5E* | 0.00333 |
| Dog Fear | ENSCAFG00000033761 | | 0.00335 |
| Dog Fear | ENSCAFG00000014670 | | 0.00383 |
| Dog Fear | ENSCAFG00000021668 | *RF00003* | 0.00399 |
| Dog Fear | ENSCAFG00000011155 | *STARD9* | 0.00422 |
| Dog Fear | ENSCAFG00000000167 | *DMXL1* | 0.00445 |
| Dog Fear | ENSCAFG00000002096 | *UXS1* | 0.00478 |
| Dog Fear | ENSCAFG00000011680 | *ATP11B* | 0.00490 |
| Dog Fear | ENSCAFG00000008551 | | 0.00532 |
| Dog Fear | ENSCAFG00000010766 | *TESMIN* | 0.00583 |
| Dog Fear | ENSCAFG00000027110 | *RF00026* | 0.00637 |
| Dog Fear | ENSCAFG00000005061 | *LMO7* | 0.00640 |
| Dog Fear | ENSCAFG00000017847 | | 0.00686 |
| Dog Fear | ENSCAFG00000038217 | | 0.00695 |
| Dog Fear | ENSCAFG00000006264 | *PCLO* | 0.00712 |
| Dog Fear | ENSCAFG00000005814 | *RAB5A* | 0.00713 |
| Dog Fear | ENSCAFG00000014989 | *WDHD1* | 0.00734 |
| Dog Fear | ENSCAFG00000000345 | *SRGAP1* | 0.00749 |
| Dog Fear | ENSCAFG00000008101 | *PLOD2* | 0.00767 |
| Dog Fear | ENSCAFG00000015257 | *ANO2* | 0.00791 |
| Dog Fear | ENSCAFG00000004369 | *NOX4* | 0.00793 |
| Dog Fear | ENSCAFG00000033944 | | 0.00823 |
| Dog Fear | ENSCAFG00000006370 | *HGF* | 0.00884 |
| Dog Fear | ENSCAFG00000023549 | *C9orf3* | 0.00896 |
| Dog Fear | ENSCAFG00000010536 | *CARMIL1* | 0.00899 |
| Dog Fear | ENSCAFG00000036543 | | 0.01037 |
| Dog Fear | ENSCAFG00000010102 | *SEMA5A* | 0.01101 |

| | | | |
|---|---|---|---|
| Dog Fear | ENSCAFG00000039324 | | 0.01189 |
| Dog Fear | ENSCAFG00000036086 | | 0.01376 |
| Dog Fear | ENSCAFG00000000430 | *ESR1* | 0.01432 |
| Dog Fear | ENSCAFG00000028434 | *RF01210* | 0.01490 |
| Dog Fear | ENSCAFG00000007533 | *ANKRD27* | 0.01574 |
| Dog Fear | ENSCAFG00000013177 | *NPAS3* | 0.01742 |
| Dog Fear | ENSCAFG00000018996 | *CDH10* | 0.01914 |
| Dog Fear | ENSCAFG00000004250 | *RAB10* | 0.01986 |
| Dog Fear | ENSCAFG00000015499 | *DLG5* | 0.01996 |
| Dog Fear | ENSCAFG00000010745 | *IGHMBP2* | 0.02311 |
| Dog Fear | ENSCAFG00000015679 | *RGS7* | 0.02337 |
| Dog Fear | ENSCAFG00000014255 | | 0.02537 |
| Dog Fear | ENSCAFG00000027709 | *RF00001* | 0.02654 |
| Dog Fear | ENSCAFG00000035441 | | 0.02842 |
| Dog Fear | ENSCAFG00000020135 | *FNBP1L* | 0.02850 |
| Dog Fear | ENSCAFG00000003323 | *KIDINS220* | 0.03284 |
| Dog Fear | ENSCAFG00000037826 | | 0.03366 |
| Dog Fear | ENSCAFG00000004568 | *CUBN* | 0.03498 |
| Dog Fear | ENSCAFG00000013246 | *LIPH* | 0.03582 |
| Dog Fear | ENSCAFG00000038695 | | 0.03670 |
| Dog Fear | ENSCAFG00000002238 | *MIOS* | 0.03837 |
| Dog Fear | ENSCAFG00000000361 | *TBC1D30* | 0.04357 |
| Dog Fear | ENSCAFG00000008276 | *HLTF* | 0.04436 |
| Dog Fear | ENSCAFG00000002372 | *ZNF135* | 0.04468 |
| Dog Fear | ENSCAFG00000003571 | *POU6F2* | 0.04603 |
| Dog Fear | ENSCAFG00000005700 | *BAIAP2* | 0.04660 |
| Dog Rivalry | ENSCAFG00000014093 | | < 0.00001 |
| Dog Rivalry | ENSCAFG00000039058 | | < 0.00001 |
| Dog Rivalry | ENSCAFG00000008909 | *CNBD1* | < 0.00001 |

| | | | |
|---|---|---|---|
| Dog Rivalry | ENSCAFG00000038113 | | < 0.00001 |
| Dog Rivalry | ENSCAFG00000037915 | | < 0.00001 |
| Dog Rivalry | ENSCAFG00000034875 | | < 0.00001 |
| Dog Rivalry | ENSCAFG00000015903 | | < 0.00001 |
| Dog Rivalry | ENSCAFG00000012545 | *CARF* | < 0.00001 |
| Dog Rivalry | ENSCAFG00000034281 | | < 0.00001 |
| Dog Rivalry | ENSCAFG00000028659 | | < 0.00001 |
| Dog Rivalry | ENSCAFG00000016111 | *FOXK1* | < 0.00001 |
| Dog Rivalry | ENSCAFG00000038128 | | < 0.00001 |
| Dog Rivalry | ENSCAFG00000020112 | *ABCD3* | < 0.00001 |
| Dog Rivalry | ENSCAFG00000005809 | *TM9SF2* | < 0.00001 |
| Dog Rivalry | ENSCAFG00000005040 | *KLF12* | < 0.00001 |
| Dog Rivalry | ENSCAFG00000017651 | *DNM2* | < 0.00001 |
| Dog Rivalry | ENSCAFG00000003005 | *CREB5* | < 0.00001 |
| Dog Rivalry | ENSCAFG00000017996 | *ACACA* | < 0.00001 |
| Dog Rivalry | ENSCAFG00000019514 | *NPHP4* | < 0.00001 |
| Dog Rivalry | ENSCAFG00000021629 | *RF00026* | < 0.00001 |
| Dog Rivalry | ENSCAFG00000001358 | *AGTPBP1* | < 0.00001 |
| Dog Rivalry | ENSCAFG00000016200 | *TCF12* | 0.00001 |
| Dog Rivalry | ENSCAFG00000039648 | | 0.00001 |
| Dog Rivalry | ENSCAFG00000030938 | | 0.00001 |
| Dog Rivalry | ENSCAFG00000003467 | *FBXL4* | 0.00001 |
| Dog Rivalry | ENSCAFG00000038668 | | 0.00001 |
| Dog Rivalry | ENSCAFG00000001630 | *MYH9* | 0.00002 |
| Dog Rivalry | ENSCAFG00000030539 | | 0.00003 |
| Dog Rivalry | ENSCAFG00000004343 | *EFCC1* | 0.00003 |
| Dog Rivalry | ENSCAFG00000031241 | *LRRC55* | 0.00004 |
| Dog Rivalry | ENSCAFG00000004243 | *SLC36A4* | 0.00004 |
| Dog Rivalry | ENSCAFG00000001646 | *PRUNE2* | 0.00004 |

| | | | |
|---|---|---|---|
| Dog Rivalry | ENSCAFG00000017293 | *CEP128* | 0.00005 |
| Dog Rivalry | ENSCAFG00000040702 | | 0.00006 |
| Dog Rivalry | ENSCAFG00000002413 | *DST* | 0.00006 |
| Dog Rivalry | ENSCAFG00000005862 | *PLCL2* | 0.00007 |
| Dog Rivalry | ENSCAFG00000000086 | *CDH20* | 0.00008 |
| Dog Rivalry | ENSCAFG00000004855 | *EPB41L5* | 0.00010 |
| Dog Rivalry | ENSCAFG00000033119 | | 0.00018 |
| Dog Rivalry | ENSCAFG00000014222 | *IFT80* | 0.00020 |
| Dog Rivalry | ENSCAFG00000031628 | | 0.00023 |
| Dog Rivalry | ENSCAFG00000005549 | *UCP3* | 0.00024 |
| Dog Rivalry | ENSCAFG00000035258 | | 0.00030 |
| Dog Rivalry | ENSCAFG00000009831 | *PRMT3* | 0.00031 |
| Dog Rivalry | ENSCAFG00000019133 | *MTMR1* | 0.00032 |
| Dog Rivalry | ENSCAFG00000011176 | *NR5A2* | 0.00032 |
| Dog Rivalry | ENSCAFG00000012638 | *DPP3* | 0.00041 |
| Dog Rivalry | ENSCAFG00000033142 | | 0.00054 |
| Dog Rivalry | ENSCAFG00000010905 | *B4GALT4* | 0.00075 |
| Dog Rivalry | ENSCAFG00000007803 | *GALNTL6* | 0.00076 |
| Dog Rivalry | ENSCAFG00000016354 | *STIM2* | 0.00080 |
| Dog Rivalry | ENSCAFG00000002390 | *FAM178B* | 0.00082 |
| Dog Rivalry | ENSCAFG00000002663 | *CD109* | 0.00085 |
| Dog Rivalry | ENSCAFG00000001166 | *KHDRBS3* | 0.00088 |
| Dog Rivalry | ENSCAFG00000025956 | *RF00001* | 0.00090 |
| Dog Rivalry | ENSCAFG00000018890 | *TXNDC11* | 0.00096 |
| Dog Rivalry | ENSCAFG00000009954 | | 0.00119 |
| Dog Rivalry | ENSCAFG00000010677 | *EYA2* | 0.00141 |
| Dog Rivalry | ENSCAFG00000036020 | | 0.00150 |
| Dog Rivalry | ENSCAFG00000002271 | *PHF14* | 0.00156 |
| Dog Rivalry | ENSCAFG00000034449 | | 0.00197 |

| | | | |
|---|---|---|---|
| Dog Rivalry | ENSCAFG00000039491 | | 0.00204 |
| Dog Rivalry | ENSCAFG00000012804 | *MACO1* | 0.00207 |
| Dog Rivalry | ENSCAFG00000012413 | *RPS6KC1* | 0.00298 |
| Dog Rivalry | ENSCAFG00000024975 | *GRXCR1* | 0.00304 |
| Dog Rivalry | ENSCAFG00000034165 | | 0.00363 |
| Dog Rivalry | ENSCAFG00000011544 | *AOX4* | 0.00397 |
| Dog Rivalry | ENSCAFG00000000696 | *RSPO2* | 0.00426 |
| Dog Rivalry | ENSCAFG00000015877 | *SH2D4B* | 0.00544 |
| Dog Rivalry | ENSCAFG00000040595 | | 0.00620 |
| Dog Rivalry | ENSCAFG00000005405 | *SEC23B* | 0.00626 |
| Dog Rivalry | ENSCAFG00000010224 | | 0.00648 |
| Dog Rivalry | ENSCAFG00000003420 | *VRK3* | 0.00655 |
| Dog Rivalry | ENSCAFG00000003043 | *KIAA1958* | 0.00666 |
| Dog Rivalry | ENSCAFG00000006462 | *UBXN8* | 0.00697 |
| Dog Rivalry | ENSCAFG00000015255 | *SLC35F4* | 0.00771 |
| Dog Rivalry | ENSCAFG00000033172 | | 0.00820 |
| Dog Rivalry | ENSCAFG00000002640 | *RIMS1* | 0.00840 |
| Dog Rivalry | ENSCAFG00000000719 | *NUDCD1* | 0.00861 |
| Dog Rivalry | ENSCAFG00000023912 | *LYG2* | 0.00867 |
| Dog Rivalry | ENSCAFG00000036254 | | 0.00998 |
| Dog Rivalry | ENSCAFG00000033155 | | 0.01008 |
| Dog Rivalry | ENSCAFG00000013057 | *MAP3K15* | 0.01083 |
| Dog Rivalry | ENSCAFG00000007575 | *DPY19L3* | 0.01123 |
| Dog Rivalry | ENSCAFG00000000655 | *TMEM181* | 0.01193 |
| Dog Rivalry | ENSCAFG00000010267 | *RAP1GDS1* | 0.01295 |
| Dog Rivalry | ENSCAFG00000011983 | *HSPBAP1* | 0.01398 |
| Dog Rivalry | ENSCAFG00000024350 | *CENPP* | 0.01519 |
| Dog Rivalry | ENSCAFG00000033819 | | 0.01877 |
| Dog Rivalry | ENSCAFG00000035757 | | 0.01897 |

| | | | |
|---|---|---|---|
| Dog Rivalry | ENSCAFG00000023463 | *ITPR1* | 0.02310 |
| Dog Rivalry | ENSCAFG00000010942 | *CCDC91* | 0.02447 |
| Dog Rivalry | ENSCAFG00000005121 | *SFMBT2* | 0.02707 |
| Dog Rivalry | ENSCAFG00000036961 | | 0.03569 |
| Dog Rivalry | ENSCAFG00000039974 | | 0.03705 |
| Dog Rivalry | ENSCAFG00000008407 | *EIF2A* | 0.03825 |
| Dog Rivalry | ENSCAFG00000005873 | *RFTN1* | 0.03840 |
| Dog Rivalry | ENSCAFG00000008185 | *ARFIP1* | 0.03915 |
| Dog Rivalry | ENSCAFG00000034736 | | 0.04064 |
| Dog Rivalry | ENSCAFG00000018204 | *TRPC5* | 0.04234 |
| Dog Rivalry | ENSCAFG00000002641 | *TTC7A* | 0.04277 |
| Dog Rivalry | ENSCAFG00000028804 | *RNF150* | 0.04413 |
| Dog Rivalry | ENSCAFG00000007494 | *SGTB* | 0.04576 |
| Dog Rivalry | ENSCAFG00000014062 | *FANCM* | 0.04626 |
| Energy | ENSCAFG00000003737 | *PARD3* | < 0.00001 |
| Energy | ENSCAFG00000001106 | *LAMA2* | < 0.00001 |
| Energy | ENSCAFG00000001700 | *SND1* | < 0.00001 |
| Energy | ENSCAFG00000009699 | *CHD9* | < 0.00001 |
| Energy | ENSCAFG00000008056 | *RYR3* | < 0.00001 |
| Energy | ENSCAFG00000007520 | *CBR4* | < 0.00001 |
| Energy | ENSCAFG00000023267 | *PRIM2* | < 0.00001 |
| Energy | ENSCAFG00000018849 | *SNX29* | < 0.00001 |
| Energy | ENSCAFG00000007301 | *CWC27* | < 0.00001 |
| Energy | ENSCAFG00000013221 | | < 0.00001 |
| Energy | ENSCAFG00000000557 | *CSNK1G3* | < 0.00001 |
| Energy | ENSCAFG00000008199 | *FMN1* | < 0.00001 |
| Energy | ENSCAFG00000032608 | *LURAP1L* | < 0.00001 |
| Energy | ENSCAFG00000009971 | *ARMH3* | < 0.00001 |
| Energy | ENSCAFG00000018857 | *GPC4* | < 0.00001 |

| | | | |
|---|---|---|---|
| Energy | ENSCAFG00000025529 | *TPK1* | < 0.00001 |
| Energy | ENSCAFG00000000430 | *ESR1* | < 0.00001 |
| Energy | ENSCAFG00000008790 | *CTNNBL1* | < 0.00001 |
| Energy | ENSCAFG00000006625 | *SFSWAP* | < 0.00001 |
| Energy | ENSCAFG00000001154 | *MRTFA* | < 0.00001 |
| Energy | ENSCAFG00000006812 | *TMEM132D* | < 0.00001 |
| Energy | ENSCAFG00000004589 | | < 0.00001 |
| Energy | ENSCAFG00000039364 | | < 0.00001 |
| Energy | ENSCAFG00000039407 | | < 0.00001 |
| Energy | ENSCAFG00000038915 | | < 0.00001 |
| Energy | ENSCAFG00000018850 | *HS6ST2* | < 0.00001 |
| Energy | ENSCAFG00000018564 | *GRIA3* | 0.00001 |
| Energy | ENSCAFG00000035898 | | 0.00001 |
| Energy | ENSCAFG00000008211 | *CACNA2D3* | 0.00002 |
| Energy | ENSCAFG00000018952 | *ARHGEF6* | 0.00002 |
| Energy | ENSCAFG00000030512 | | 0.00002 |
| Energy | ENSCAFG00000013228 | *CNKSR2* | 0.00002 |
| Energy | ENSCAFG00000033823 | | 0.00002 |
| Energy | ENSCAFG00000018024 | *TRAPPC8* | 0.00002 |
| Energy | ENSCAFG00000002364 | *ADGRL3* | 0.00002 |
| Energy | ENSCAFG00000019051 | | 0.00003 |
| Energy | ENSCAFG00000040039 | | 0.00004 |
| Energy | ENSCAFG00000007586 | *MAST4* | 0.00005 |
| Energy | ENSCAFG00000014257 | *PPP2R2C* | 0.00006 |
| Energy | ENSCAFG00000028405 | *RF00026* | 0.00006 |
| Energy | ENSCAFG00000012360 | *ALS2* | 0.00006 |
| Energy | ENSCAFG00000014916 | *FAM227B* | 0.00006 |
| Energy | ENSCAFG00000018858 | *TARS* | 0.00007 |
| Energy | ENSCAFG00000011082 | *CAMSAP2* | 0.00008 |

| | | | |
|---|---|---|---|
| Energy | ENSCAFG00000035434 | | 0.00008 |
| Energy | ENSCAFG00000033989 | | 0.00010 |
| Energy | ENSCAFG00000022240 | *RF00410* | 0.00010 |
| Energy | ENSCAFG00000040529 | | 0.00012 |
| Energy | ENSCAFG00000006764 | *PARG* | 0.00014 |
| Energy | ENSCAFG00000001234 | *UHRF1BP1* | 0.00018 |
| Energy | ENSCAFG00000028137 | *RF00100* | 0.00019 |
| Energy | ENSCAFG00000039300 | | 0.00022 |
| Energy | ENSCAFG00000008551 | | 0.00022 |
| Energy | ENSCAFG00000003322 | *IFRD1* | 0.00023 |
| Energy | ENSCAFG00000000011 | *NFATC1* | 0.00028 |
| Energy | ENSCAFG00000008684 | *SLC4A5* | 0.00034 |
| Energy | ENSCAFG00000003663 | *GSN* | 0.00035 |
| Energy | ENSCAFG00000029157 | *NOVA1* | 0.00043 |
| Energy | ENSCAFG00000013749 | *CUX1* | 0.00048 |
| Energy | ENSCAFG00000016531 | *ADGRA3* | 0.00055 |
| Energy | ENSCAFG00000000279 | *REPS1* | 0.00066 |
| Energy | ENSCAFG00000030012 | | 0.00070 |
| Energy | ENSCAFG00000002028 | *VLDLR* | 0.00074 |
| Energy | ENSCAFG00000039266 | | 0.00082 |
| Energy | ENSCAFG00000001425 | *GKAP1* | 0.00082 |
| Energy | ENSCAFG00000007750 | *CASP3* | 0.00084 |
| Energy | ENSCAFG00000038450 | | 0.00085 |
| Energy | ENSCAFG00000036209 | | 0.00097 |
| Energy | ENSCAFG00000036778 | | 0.00124 |
| Energy | ENSCAFG00000014454 | *ATM* | 0.00124 |
| Energy | ENSCAFG00000038201 | | 0.00184 |
| Energy | ENSCAFG00000012098 | *PGBD5* | 0.00190 |
| Energy | ENSCAFG00000004290 | *SAP130* | 0.00229 |

| | | | |
|---|---|---|---|
| Energy | ENSCAFG00000012544 | *COPA* | 0.00236 |
| Energy | ENSCAFG00000002414 | *AGMO* | 0.00268 |
| Energy | ENSCAFG00000003161 | *BBS9* | 0.00288 |
| Energy | ENSCAFG00000032743 | | 0.00309 |
| Energy | ENSCAFG00000019082 | *RBFOX1* | 0.00324 |
| Energy | ENSCAFG00000039058 | | 0.00327 |
| Energy | ENSCAFG00000009486 | *RPGRIP1L* | 0.00401 |
| Energy | ENSCAFG00000001852 | *ADAM22* | 0.00416 |
| Energy | ENSCAFG00000002640 | *RIMS1* | 0.00455 |
| Energy | ENSCAFG00000015135 | *LDB2* | 0.00480 |
| Energy | ENSCAFG00000015422 | *KIAA0753* | 0.00494 |
| Energy | ENSCAFG00000018204 | *TRPC5* | 0.00496 |
| Energy | ENSCAFG00000039092 | | 0.00501 |
| Energy | ENSCAFG00000009903 | *MYLIP* | 0.00530 |
| Energy | ENSCAFG00000015604 | *ILDR2* | 0.00557 |
| Energy | ENSCAFG00000013114 | *RGSL1* | 0.00560 |
| Energy | ENSCAFG00000014349 | *TBC1D14* | 0.00585 |
| Energy | ENSCAFG00000036051 | | 0.00620 |
| Energy | ENSCAFG00000018996 | *CDH10* | 0.00639 |
| Energy | ENSCAFG00000003370 | *ASAP2* | 0.00776 |
| Energy | ENSCAFG00000038846 | | 0.00792 |
| Energy | ENSCAFG00000013852 | *MATN2* | 0.00906 |
| Energy | ENSCAFG00000004479 | *ANO10* | 0.00950 |
| Energy | ENSCAFG00000018853 | *USP26* | 0.00957 |
| Energy | ENSCAFG00000009991 | | 0.00969 |
| Energy | ENSCAFG00000033361 | | 0.01042 |
| Energy | ENSCAFG00000005870 | *RASGRP3* | 0.01043 |
| Energy | ENSCAFG00000018083 | *TMEM164* | 0.01089 |
| Energy | ENSCAFG00000033131 | | 0.01108 |

| | | | |
|---|---|---|---|
| Energy | ENSCAFG00000014169 | | 0.01150 |
| Energy | ENSCAFG00000008284 | *EXTL3* | 0.01178 |
| Energy | ENSCAFG00000007907 | *LRBA* | 0.01197 |
| Energy | ENSCAFG00000037023 | | 0.01209 |
| Energy | ENSCAFG00000029142 | *C4H5orf51* | 0.01313 |
| Energy | ENSCAFG00000002341 | *HMGCLL1* | 0.01379 |
| Energy | ENSCAFG00000008039 | *DIP2B* | 0.01522 |
| Energy | ENSCAFG00000015351 | *BAIAP2L1* | 0.01535 |
| Energy | ENSCAFG00000037362 | | 0.01601 |
| Energy | ENSCAFG00000036039 | | 0.01678 |
| Energy | ENSCAFG00000006229 | *PCDH1* | 0.01925 |
| Energy | ENSCAFG00000038646 | | 0.01983 |
| Energy | ENSCAFG00000032548 | *KCNH5* | 0.02030 |
| Energy | ENSCAFG00000002045 | | 0.02171 |
| Energy | ENSCAFG00000005330 | *SH2D4A* | 0.02285 |
| Energy | ENSCAFG00000011150 | *GALNT17* | 0.02560 |
| Energy | ENSCAFG00000018351 | *ENOSF1* | 0.02601 |
| Energy | ENSCAFG00000028579 | *RTN4RL1* | 0.02693 |
| Energy | ENSCAFG00000001906 | *FZD1* | 0.03125 |
| Energy | ENSCAFG00000008293 | *FZD3* | 0.03140 |
| Energy | ENSCAFG00000026050 | *RF00026* | 0.03177 |
| Energy | ENSCAFG00000037039 | | 0.03274 |
| Energy | ENSCAFG00000038167 | | 0.03353 |
| Energy | ENSCAFG00000024967 | *ASIP* | 0.03577 |
| Energy | ENSCAFG00000003432 | *CTTNBP2* | 0.03770 |
| Energy | ENSCAFG00000035899 | | 0.03996 |
| Energy | ENSCAFG00000038463 | | 0.04014 |
| Energy | ENSCAFG00000009839 | *CD101* | 0.04215 |
| Energy | ENSCAFG00000037990 | | 0.04223 |

| | | | |
|---|---|---|---|
| Energy | ENSCAFG00000023072 | *POC1A* | 0.04327 |
| Energy | ENSCAFG00000035285 | | 0.04419 |
| Energy | ENSCAFG00000001434 | *KDM4C* | 0.04440 |
| Energy | ENSCAFG00000019455 | *REXO1* | 0.04543 |
| Energy | ENSCAFG00000002025 | *GCC2* | 0.04646 |
| Energy | ENSCAFG00000017349 | *STXBP4* | 0.04753 |
| Energy | ENSCAFG00000033190 | | 0.04809 |
| Energy | ENSCAFG00000018175 | *CAPN6* | 0.04825 |
| Energy | ENSCAFG00000000070 | *PHLPP1* | 0.04880 |
| Excitability | ENSCAFG00000017252 | *ATRX* | < 0.00001 |
| Excitability | ENSCAFG00000007650 | *NEK1* | < 0.00001 |
| Excitability | ENSCAFG00000005914 | *TMTC2* | < 0.00001 |
| Excitability | ENSCAFG00000006732 | *PXDNL* | < 0.00001 |
| Excitability | ENSCAFG00000038256 | | < 0.00001 |
| Excitability | ENSCAFG00000007301 | *CWC27* | < 0.00001 |
| Excitability | ENSCAFG00000000070 | *PHLPP1* | < 0.00001 |
| Excitability | ENSCAFG00000006743 | *PCMTD1* | < 0.00001 |
| Excitability | ENSCAFG00000004087 | *MAML2* | < 0.00001 |
| Excitability | ENSCAFG00000031499 | *GLIS3* | < 0.00001 |
| Excitability | ENSCAFG00000009849 | *SPTBN5* | < 0.00001 |
| Excitability | ENSCAFG00000013607 | *PDE11A* | < 0.00001 |
| Excitability | ENSCAFG00000039072 | | < 0.00001 |
| Excitability | ENSCAFG00000008352 | *PHF20* | < 0.00001 |
| Excitability | ENSCAFG00000006764 | *PARG* | < 0.00001 |
| Excitability | ENSCAFG00000027967 | *RF00009* | < 0.00001 |
| Excitability | ENSCAFG00000023079 | *AFF3* | < 0.00001 |
| Excitability | ENSCAFG00000027835 | *RF00026* | < 0.00001 |
| Excitability | ENSCAFG00000007007 | *ATP8A2* | < 0.00001 |
| Excitability | ENSCAFG00000001314 | *CACNA1I* | < 0.00001 |

| | | | |
|---|---|---|---|
| Excitability | ENSCAFG00000013221 | | 0.00001 |
| Excitability | ENSCAFG00000009662 | *MGA* | 0.00001 |
| Excitability | ENSCAFG00000033690 | | 0.00001 |
| Excitability | ENSCAFG00000010194 | *ABCC12* | 0.00001 |
| Excitability | ENSCAFG00000014916 | *FAM227B* | 0.00002 |
| Excitability | ENSCAFG00000004160 | *TNIP3* | 0.00002 |
| Excitability | ENSCAFG00000024647 | *SERPINB5* | 0.00002 |
| Excitability | ENSCAFG00000033155 | | 0.00003 |
| Excitability | ENSCAFG00000038695 | | 0.00003 |
| Excitability | ENSCAFG00000001557 | *CCDC171* | 0.00003 |
| Excitability | ENSCAFG00000033688 | | 0.00003 |
| Excitability | ENSCAFG00000007379 | *CAMK4* | 0.00005 |
| Excitability | ENSCAFG00000004535 | *NR2C2* | 0.00005 |
| Excitability | ENSCAFG00000000935 | *CEP85L* | 0.00007 |
| Excitability | ENSCAFG00000006625 | *SFSWAP* | 0.00007 |
| Excitability | ENSCAFG00000007963 | *BLK* | 0.00007 |
| Excitability | ENSCAFG00000003465 | *EGFR* | 0.00007 |
| Excitability | ENSCAFG00000006729 | *OGDHL* | 0.00017 |
| Excitability | ENSCAFG00000005373 | *SLC24A3* | 0.00018 |
| Excitability | ENSCAFG00000004589 | | 0.00018 |
| Excitability | ENSCAFG00000011212 | *PIK3CA* | 0.00022 |
| Excitability | ENSCAFG00000005054 | *ZRANB3* | 0.00024 |
| Excitability | ENSCAFG00000000756 | *GRAMD4* | 0.00026 |
| Excitability | ENSCAFG00000002155 | | 0.00028 |
| Excitability | ENSCAFG00000000337 | *PPM1H* | 0.00039 |
| Excitability | ENSCAFG00000002671 | *SP4* | 0.00047 |
| Excitability | ENSCAFG00000010242 | *SLC4A10* | 0.00052 |
| Excitability | ENSCAFG00000016732 | *SIPA1L1* | 0.00052 |
| Excitability | ENSCAFG00000039058 | | 0.00060 |

| | | | |
|---|---|---|---|
| Excitability | ENSCAFG00000026031 | *RF00026* | 0.00060 |
| Excitability | ENSCAFG00000000786 | *CSMD3* | 0.00061 |
| Excitability | ENSCAFG00000002323 | *MGAT4A* | 0.00068 |
| Excitability | ENSCAFG00000031780 | | 0.00089 |
| Excitability | ENSCAFG00000015721 | *LIPA* | 0.00096 |
| Excitability | ENSCAFG00000036709 | | 0.00105 |
| Excitability | ENSCAFG00000013228 | *CNKSR2* | 0.00108 |
| Excitability | ENSCAFG00000001930 | *PGM5* | 0.00118 |
| Excitability | ENSCAFG00000039585 | | 0.00146 |
| Excitability | ENSCAFG00000008761 | *CX3CL1* | 0.00148 |
| Excitability | ENSCAFG00000018100 | *SCAPER* | 0.00175 |
| Excitability | ENSCAFG00000012890 | *HAT1* | 0.00188 |
| Excitability | ENSCAFG00000012478 | *PLEKHA5* | 0.00224 |
| Excitability | ENSCAFG00000022264 | *RF00015* | 0.00330 |
| Excitability | ENSCAFG00000037577 | | 0.00372 |
| Excitability | ENSCAFG00000040724 | | 0.00394 |
| Excitability | ENSCAFG00000019051 | | 0.00420 |
| Excitability | ENSCAFG00000029961 | *APCDD1L* | 0.00494 |
| Excitability | ENSCAFG00000010221 | *SUFU* | 0.00494 |
| Excitability | ENSCAFG00000015648 | | 0.00621 |
| Excitability | ENSCAFG00000008293 | *FZD3* | 0.00766 |
| Excitability | ENSCAFG00000003710 | *CRYBG1* | 0.00766 |
| Excitability | ENSCAFG00000031721 | *CPNE1* | 0.00826 |
| Excitability | ENSCAFG00000032470 | | 0.00854 |
| Excitability | ENSCAFG00000035101 | | 0.00869 |
| Excitability | ENSCAFG00000036231 | | 0.00973 |
| Excitability | ENSCAFG00000013838 | *MASP1* | 0.01039 |
| Excitability | ENSCAFG00000029156 | | 0.01043 |
| Excitability | ENSCAFG00000031045 | | 0.01112 |

| | | | |
|---|---|---|---|
| Excitability | ENSCAFG00000040631 | | 0.01148 |
| Excitability | ENSCAFG00000015943 | *PRTG* | 0.01253 |
| Excitability | ENSCAFG00000007361 | *EPB41L4A* | 0.01256 |
| Excitability | ENSCAFG00000005271 | *C22H16orf87* | 0.01259 |
| Excitability | ENSCAFG00000029313 | *CLDN1* | 0.01340 |
| Excitability | ENSCAFG00000007079 | | 0.01406 |
| Excitability | ENSCAFG00000033410 | | 0.01419 |
| Excitability | ENSCAFG00000008058 | *MTMR9* | 0.01502 |
| Excitability | ENSCAFG00000008568 | *AGPAT5* | 0.01550 |
| Excitability | ENSCAFG00000039209 | | 0.01560 |
| Excitability | ENSCAFG00000007970 | *FAM167A* | 0.01563 |
| Excitability | ENSCAFG00000012179 | *UNC45A* | 0.01579 |
| Excitability | ENSCAFG00000005140 | *VHL* | 0.01581 |
| Excitability | ENSCAFG00000014454 | *ATM* | 0.01629 |
| Excitability | ENSCAFG00000001106 | *LAMA2* | 0.01708 |
| Excitability | ENSCAFG00000002364 | *ADGRL3* | 0.01824 |
| Excitability | ENSCAFG00000007789 | *ARHGAP10* | 0.01874 |
| Excitability | ENSCAFG00000006555 | *PRKDC* | 0.01911 |
| Excitability | ENSCAFG00000007912 | *TRPC4AP* | 0.02049 |
| Excitability | ENSCAFG00000033823 | | 0.02049 |
| Excitability | ENSCAFG00000013925 | *SEMA6D* | 0.02319 |
| Excitability | ENSCAFG00000038846 | | 0.02326 |
| Excitability | ENSCAFG00000039828 | | 0.02339 |
| Excitability | ENSCAFG00000018123 | *ARSI* | 0.02376 |
| Excitability | ENSCAFG00000009371 | *SORBS3* | 0.02486 |
| Excitability | ENSCAFG00000010988 | *CFAP410* | 0.02856 |
| Excitability | ENSCAFG00000009367 | *HPS1* | 0.02942 |
| Excitability | ENSCAFG00000017958 | *COL5A3* | 0.02959 |
| Excitability | ENSCAFG00000022350 | *RF00026* | 0.03231 |

| | | | |
|---|---|---|---|
| Excitability | ENSCAFG00000029157 | *NOVA1* | 0.03279 |
| Excitability | ENSCAFG00000015932 | *DNAAF4* | 0.03324 |
| Excitability | ENSCAFG00000011934 | *ARHGEF12* | 0.03560 |
| Excitability | ENSCAFG00000036595 | | 0.03576 |
| Excitability | ENSCAFG00000006746 | *PDX1* | 0.03741 |
| Excitability | ENSCAFG00000034875 | | 0.03862 |
| Excitability | ENSCAFG00000020376 | *ZZZ3* | 0.03936 |
| Excitability | ENSCAFG00000006809 | *NCOA4* | 0.04165 |
| Excitability | ENSCAFG00000034407 | | 0.04620 |
| Excitability | ENSCAFG00000007302 | *TNKS2* | 0.04716 |
| Nonsocial Fear | ENSCAFG00000010026 | *SMARCAD1* | < 0.00001 |
| Nonsocial Fear | ENSCAFG00000005775 | *BIRC6* | < 0.00001 |
| Nonsocial Fear | ENSCAFG00000009386 | *HPSE2* | < 0.00001 |
| Nonsocial Fear | ENSCAFG00000038400 | | < 0.00001 |
| Nonsocial Fear | ENSCAFG00000001460 | *PTPRD* | < 0.00001 |
| Nonsocial Fear | ENSCAFG00000009227 | *WDR41* | < 0.00001 |
| Nonsocial Fear | ENSCAFG00000000955 | *TBC1D32* | < 0.00001 |
| Nonsocial Fear | ENSCAFG00000033142 | | < 0.00001 |
| Nonsocial Fear | ENSCAFG00000034838 | | < 0.00001 |
| Nonsocial Fear | ENSCAFG00000035815 | | < 0.00001 |
| Nonsocial Fear | ENSCAFG00000017256 | | < 0.00001 |
| Nonsocial Fear | ENSCAFG00000011680 | *ATP11B* | < 0.00001 |
| Nonsocial Fear | ENSCAFG00000017297 | *GPR174* | < 0.00001 |
| Nonsocial Fear | ENSCAFG00000017327 | *BRWD3* | < 0.00001 |
| Nonsocial Fear | ENSCAFG00000034053 | | < 0.00001 |
| Nonsocial Fear | ENSCAFG00000030358 | *RF00026* | < 0.00001 |
| Nonsocial Fear | ENSCAFG00000006592 | *STK32A* | < 0.00001 |
| Nonsocial Fear | ENSCAFG00000002160 | *SPTLC1* | 0.00001 |
| Nonsocial Fear | ENSCAFG00000014245 | *PDE1A* | 0.00001 |

| | | | |
|---|---|---|---|
| Nonsocial Fear | ENSCAFG00000033300 | | 0.00001 |
| Nonsocial Fear | ENSCAFG00000018635 | *ROR1* | 0.00001 |
| Nonsocial Fear | ENSCAFG00000003161 | *BBS9* | 0.00001 |
| Nonsocial Fear | ENSCAFG00000009764 | *PPHLN1* | 0.00003 |
| Nonsocial Fear | ENSCAFG00000040257 | | 0.00004 |
| Nonsocial Fear | ENSCAFG00000012098 | *PGBD5* | 0.00004 |
| Nonsocial Fear | ENSCAFG00000011936 | *TRPM8* | 0.00005 |
| Nonsocial Fear | ENSCAFG00000006278 | *ATRN* | 0.00011 |
| Nonsocial Fear | ENSCAFG00000003479 | | 0.00011 |
| Nonsocial Fear | ENSCAFG00000004568 | *CUBN* | 0.00017 |
| Nonsocial Fear | ENSCAFG00000018580 | | 0.00027 |
| Nonsocial Fear | ENSCAFG00000007420 | *CBFA2T2* | 0.00028 |
| Nonsocial Fear | ENSCAFG00000017294 | *P2RY10* | 0.00034 |
| Nonsocial Fear | ENSCAFG00000004186 | | 0.00039 |
| Nonsocial Fear | ENSCAFG00000029154 | *EFCAB2* | 0.00050 |
| Nonsocial Fear | ENSCAFG00000001106 | *LAMA2* | 0.00056 |
| Nonsocial Fear | ENSCAFG00000039818 | | 0.00056 |
| Nonsocial Fear | ENSCAFG00000008882 | *LEKR1* | 0.00081 |
| Nonsocial Fear | ENSCAFG00000002527 | *LRPPRC* | 0.00085 |
| Nonsocial Fear | ENSCAFG00000037837 | | 0.00093 |
| Nonsocial Fear | ENSCAFG00000002390 | *FAM178B* | 0.00103 |
| Nonsocial Fear | ENSCAFG00000028102 | *RF00088* | 0.00114 |
| Nonsocial Fear | ENSCAFG00000027105 | *RF00026* | 0.00124 |
| Nonsocial Fear | ENSCAFG00000036236 | | 0.00131 |
| Nonsocial Fear | ENSCAFG00000033248 | | 0.00148 |
| Nonsocial Fear | ENSCAFG00000039648 | | 0.00152 |
| Nonsocial Fear | ENSCAFG00000023549 | *C9orf3* | 0.00191 |
| Nonsocial Fear | ENSCAFG00000036778 | | 0.00208 |
| Nonsocial Fear | ENSCAFG00000018100 | *SCAPER* | 0.00275 |

| | | | |
|---|---|---|---|
| Nonsocial Fear | ENSCAFG00000027810 | *RF00026* | 0.00325 |
| Nonsocial Fear | ENSCAFG00000009417 | *CDYL* | 0.00373 |
| Nonsocial Fear | ENSCAFG00000019384 | *DIRAS1* | 0.00393 |
| Nonsocial Fear | ENSCAFG00000002667 | | 0.00415 |
| Nonsocial Fear | ENSCAFG00000004377 | *ACBD5* | 0.00416 |
| Nonsocial Fear | ENSCAFG00000005704 | *TMEM173* | 0.00422 |
| Nonsocial Fear | ENSCAFG00000017645 | *MYO9A* | 0.00422 |
| Nonsocial Fear | ENSCAFG00000008945 | *SLIT1* | 0.00438 |
| Nonsocial Fear | ENSCAFG00000037340 | | 0.00470 |
| Nonsocial Fear | ENSCAFG00000022548 | *RF00001* | 0.00474 |
| Nonsocial Fear | ENSCAFG00000017502 | *ANP32A* | 0.00487 |
| Nonsocial Fear | ENSCAFG00000001697 | *UBR2* | 0.00538 |
| Nonsocial Fear | ENSCAFG00000005720 | *CLPB* | 0.00668 |
| Nonsocial Fear | ENSCAFG00000016051 | *CACNA1C* | 0.00851 |
| Nonsocial Fear | ENSCAFG00000019831 | *SYPL2* | 0.00853 |
| Nonsocial Fear | ENSCAFG00000017252 | *ATRX* | 0.00941 |
| Nonsocial Fear | ENSCAFG00000040591 | | 0.01188 |
| Nonsocial Fear | ENSCAFG00000038749 | | 0.01205 |
| Nonsocial Fear | ENSCAFG00000030067 | | 0.01257 |
| Nonsocial Fear | ENSCAFG00000009885 | *JARID2* | 0.01368 |
| Nonsocial Fear | ENSCAFG00000034343 | | 0.01529 |
| Nonsocial Fear | ENSCAFG00000016982 | *DOCK2* | 0.01578 |
| Nonsocial Fear | ENSCAFG00000008630 | *KIFC3* | 0.01644 |
| Nonsocial Fear | ENSCAFG00000033801 | | 0.01669 |
| Nonsocial Fear | ENSCAFG00000030771 | *LCLAT1* | 0.01825 |
| Nonsocial Fear | ENSCAFG00000034943 | | 0.01924 |
| Nonsocial Fear | ENSCAFG00000001091 | *TGFBI* | 0.01991 |
| Nonsocial Fear | ENSCAFG00000037802 | | 0.02090 |
| Nonsocial Fear | ENSCAFG00000009624 | *CWF19L1* | 0.02266 |

| | | | |
|---|---|---|---|
| Nonsocial Fear | ENSCAFG00000001362 | *PLXNA4* | 0.02492 |
| Nonsocial Fear | ENSCAFG00000006663 | | 0.02578 |
| Nonsocial Fear | ENSCAFG00000011430 | *STXBP5L* | 0.02793 |
| Nonsocial Fear | ENSCAFG00000004432 | *RCBTB2* | 0.02975 |
| Nonsocial Fear | ENSCAFG00000027910 | *RF00026* | 0.03166 |
| Nonsocial Fear | ENSCAFG00000006691 | *SLC46A3* | 0.03168 |
| Nonsocial Fear | ENSCAFG00000040307 | | 0.03204 |
| Nonsocial Fear | ENSCAFG00000040668 | | 0.03236 |
| Nonsocial Fear | ENSCAFG00000031017 | *SMIM33* | 0.03311 |
| Nonsocial Fear | ENSCAFG00000014030 | *LARS2* | 0.03417 |
| Nonsocial Fear | ENSCAFG00000039348 | | 0.03497 |
| Nonsocial Fear | ENSCAFG00000030316 | *RF00026* | 0.04945 |
| Owner Aggression | ENSCAFG00000033361 | | < 0.00001 |
| Owner Aggression | ENSCAFG00000018100 | *SCAPER* | < 0.00001 |
| Owner Aggression | ENSCAFG00000011983 | *HSPBAP1* | < 0.00001 |
| Owner Aggression | ENSCAFG00000001646 | *PRUNE2* | < 0.00001 |
| Owner Aggression | ENSCAFG00000015639 | *MNAT1* | < 0.00001 |
| Owner Aggression | ENSCAFG00000018811 | | < 0.00001 |
| Owner Aggression | ENSCAFG00000003749 | *SLC7A11* | < 0.00001 |
| Owner Aggression | ENSCAFG00000006730 | *TTC17* | < 0.00001 |
| Owner Aggression | ENSCAFG00000013288 | *ADAMTSL3* | < 0.00001 |
| Owner Aggression | ENSCAFG00000036018 | | < 0.00001 |
| Owner Aggression | ENSCAFG00000039058 | | < 0.00001 |
| Owner Aggression | ENSCAFG00000038128 | | < 0.00001 |
| Owner Aggression | ENSCAFG00000035215 | | < 0.00001 |
| Owner Aggression | ENSCAFG00000013779 | | < 0.00001 |
| Owner Aggression | ENSCAFG00000001557 | *CCDC171* | < 0.00001 |
| Owner Aggression | ENSCAFG00000034411 | | < 0.00001 |
| Owner Aggression | ENSCAFG00000005121 | *SFMBT2* | < 0.00001 |

| | | | |
|---|---|---|---|
| Owner Aggression | ENSCAFG00000027188 | *RF00026* | < 0.00001 |
| Owner Aggression | ENSCAFG00000014257 | *PPP2R2C* | < 0.00001 |
| Owner Aggression | ENSCAFG00000010120 | *RAD54L2* | < 0.00001 |
| Owner Aggression | ENSCAFG00000016511 | *RFX1* | < 0.00001 |
| Owner Aggression | ENSCAFG00000023938 | *GPR158* | < 0.00001 |
| Owner Aggression | ENSCAFG00000002271 | *PHF14* | < 0.00001 |
| Owner Aggression | ENSCAFG00000000770 | | 0.00001 |
| Owner Aggression | ENSCAFG00000001700 | *SND1* | 0.00001 |
| Owner Aggression | ENSCAFG00000001630 | *MYH9* | 0.00001 |
| Owner Aggression | ENSCAFG00000011940 | *PARP9* | 0.00001 |
| Owner Aggression | ENSCAFG00000018971 | | 0.00002 |
| Owner Aggression | ENSCAFG00000007420 | *CBFA2T2* | 0.00002 |
| Owner Aggression | ENSCAFG00000013840 | | 0.00002 |
| Owner Aggression | ENSCAFG00000038859 | | 0.00002 |
| Owner Aggression | ENSCAFG00000018799 | *IGSF1* | 0.00003 |
| Owner Aggression | ENSCAFG00000006492 | *B4GALNT4* | 0.00003 |
| Owner Aggression | ENSCAFG00000004157 | *RAB7A* | 0.00004 |
| Owner Aggression | ENSCAFG00000006989 | *DEFB119* | 0.00004 |
| Owner Aggression | ENSCAFG00000007107 | *TTLL9* | 0.00009 |
| Owner Aggression | ENSCAFG00000033119 | | 0.00009 |
| Owner Aggression | ENSCAFG00000003580 | *GRIK2* | 0.00018 |
| Owner Aggression | ENSCAFG00000018777 | *ENOX2* | 0.00021 |
| Owner Aggression | ENSCAFG00000019958 | *NTNG1* | 0.00023 |
| Owner Aggression | ENSCAFG00000025956 | *RF00001* | 0.00029 |
| Owner Aggression | ENSCAFG00000011176 | *NR5A2* | 0.00031 |
| Owner Aggression | ENSCAFG00000034449 | | 0.00037 |
| Owner Aggression | ENSCAFG00000015618 | *TULP3* | 0.00041 |
| Owner Aggression | ENSCAFG00000016982 | *DOCK2* | 0.00042 |
| Owner Aggression | ENSCAFG00000006462 | *UBXN8* | 0.00051 |

| | | | |
|---|---|---|---|
| Owner Aggression | ENSCAFG00000003705 | *CUL2* | 0.00061 |
| Owner Aggression | ENSCAFG00000003106 | *EHBP1* | 0.00062 |
| Owner Aggression | ENSCAFG00000002949 | *GC* | 0.00062 |
| Owner Aggression | ENSCAFG00000035525 | | 0.00064 |
| Owner Aggression | ENSCAFG00000037915 | | 0.00064 |
| Owner Aggression | ENSCAFG00000006573 | *FAM19A4* | 0.00079 |
| Owner Aggression | ENSCAFG00000001166 | *KHDRBS3* | 0.00081 |
| Owner Aggression | ENSCAFG00000014579 | *PRDX6* | 0.00093 |
| Owner Aggression | ENSCAFG00000002714 | *EPHA5* | 0.00096 |
| Owner Aggression | ENSCAFG00000004764 | *VWA8* | 0.00102 |
| Owner Aggression | ENSCAFG00000028804 | *RNF150* | 0.00230 |
| Owner Aggression | ENSCAFG00000004343 | *EFCC1* | 0.00293 |
| Owner Aggression | ENSCAFG00000031347 | | 0.00317 |
| Owner Aggression | ENSCAFG00000017676 | *SETBP1* | 0.00354 |
| Owner Aggression | ENSCAFG00000039028 | | 0.00373 |
| Owner Aggression | ENSCAFG00000002796 | *CCDC88A* | 0.00462 |
| Owner Aggression | ENSCAFG00000008237 | *HMBOX1* | 0.00521 |
| Owner Aggression | ENSCAFG00000012545 | *CARF* | 0.00576 |
| Owner Aggression | ENSCAFG00000038496 | | 0.00596 |
| Owner Aggression | ENSCAFG00000031408 | *CFAP299* | 0.00636 |
| Owner Aggression | ENSCAFG00000023322 | *RASSF9* | 0.00644 |
| Owner Aggression | ENSCAFG00000035968 | | 0.00659 |
| Owner Aggression | ENSCAFG00000009669 | *MDM4* | 0.00672 |
| Owner Aggression | ENSCAFG00000014718 | *CWF19L2* | 0.00760 |
| Owner Aggression | ENSCAFG00000030440 | *NOV* | 0.00859 |
| Owner Aggression | ENSCAFG00000031628 | | 0.00860 |
| Owner Aggression | ENSCAFG00000038140 | | 0.00877 |
| Owner Aggression | ENSCAFG00000000086 | *CDH20* | 0.00895 |
| Owner Aggression | ENSCAFG00000039505 | | 0.00979 |

| | | | |
|---|---|---|---|
| Owner Aggression | ENSCAFG00000040675 | | 0.01019 |
| Owner Aggression | ENSCAFG00000015877 | *SH2D4B* | 0.01102 |
| Owner Aggression | ENSCAFG00000012804 | *MACO1* | 0.01156 |
| Owner Aggression | ENSCAFG00000004224 | *PLEKHB2* | 0.01188 |
| Owner Aggression | ENSCAFG00000013410 | *RGL1* | 0.01217 |
| Owner Aggression | ENSCAFG00000038122 | | 0.01223 |
| Owner Aggression | ENSCAFG00000000828 | *RPS6KA2* | 0.01492 |
| Owner Aggression | ENSCAFG00000026119 | *RF00026* | 0.01517 |
| Owner Aggression | ENSCAFG00000016111 | *FOXK1* | 0.01740 |
| Owner Aggression | ENSCAFG00000010857 | *MMAB* | 0.02052 |
| Owner Aggression | ENSCAFG00000012413 | *RPS6KC1* | 0.02066 |
| Owner Aggression | ENSCAFG00000018136 | | 0.02168 |
| Owner Aggression | ENSCAFG00000039151 | | 0.02474 |
| Owner Aggression | ENSCAFG00000032993 | | 0.02878 |
| Owner Aggression | ENSCAFG00000014093 | | 0.02920 |
| Owner Aggression | ENSCAFG00000011672 | *SLC15A2* | 0.03073 |
| Owner Aggression | ENSCAFG00000008467 | *EDIL3* | 0.03589 |
| Owner Aggression | ENSCAFG00000000795 | *PDE10A* | 0.03990 |
| Owner Aggression | ENSCAFG00000009318 | *CCDC148* | 0.04004 |
| Owner Aggression | ENSCAFG00000032160 | | 0.04078 |
| Owner Aggression | ENSCAFG00000005607 | *SPTLC3* | 0.04210 |
| Owner Aggression | ENSCAFG00000005862 | *PLCL2* | 0.04222 |
| Owner Aggression | ENSCAFG00000014599 | *GOLIM4* | 0.04459 |
| Owner Aggression | ENSCAFG00000007557 | *HHIP* | 0.04758 |
| Separation Problems | ENSCAFG00000028659 | | < 0.00001 |
| Separation Problems | ENSCAFG00000035085 | | < 0.00001 |
| Separation Problems | ENSCAFG00000013221 | | < 0.00001 |
| Separation Problems | ENSCAFG00000011820 | *PRPF3* | < 0.00001 |
| Separation Problems | ENSCAFG00000031499 | *GLIS3* | < 0.00001 |

| | | | |
|---|---|---|---|
| Separation Problems | ENSCAFG00000005746 | *NAV3* | < 0.00001 |
| Separation Problems | ENSCAFG00000039076 | | < 0.00001 |
| Separation Problems | ENSCAFG00000040856 | | < 0.00001 |
| Separation Problems | ENSCAFG00000004087 | *MAML2* | < 0.00001 |
| Separation Problems | ENSCAFG00000002096 | *UXS1* | < 0.00001 |
| Separation Problems | ENSCAFG00000015459 | | < 0.00001 |
| Separation Problems | ENSCAFG00000017685 | *RIT2* | < 0.00001 |
| Separation Problems | ENSCAFG00000039762 | | < 0.00001 |
| Separation Problems | ENSCAFG00000000745 | *TBC1D22A* | < 0.00001 |
| Separation Problems | ENSCAFG00000015477 | | < 0.00001 |
| Separation Problems | ENSCAFG00000036551 | | < 0.00001 |
| Separation Problems | ENSCAFG00000000756 | *GRAMD4* | < 0.00001 |
| Separation Problems | ENSCAFG00000002413 | *DST* | 0.00001 |
| Separation Problems | ENSCAFG00000019132 | *RPH3AL* | 0.00001 |
| Separation Problems | ENSCAFG00000012545 | *CARF* | 0.00001 |
| Separation Problems | ENSCAFG00000002667 | | 0.00001 |
| Separation Problems | ENSCAFG00000009318 | *CCDC148* | 0.00002 |
| Separation Problems | ENSCAFG00000001002 | *SMPDL3A* | 0.00002 |
| Separation Problems | ENSCAFG00000014975 | *EIF4G3* | 0.00003 |
| Separation Problems | ENSCAFG00000039187 | | 0.00003 |
| Separation Problems | ENSCAFG00000010120 | *RAD54L2* | 0.00003 |
| Separation Problems | ENSCAFG00000003570 | *FBXW2* | 0.00004 |
| Separation Problems | ENSCAFG00000026031 | *RF00026* | 0.00004 |
| Separation Problems | ENSCAFG00000033314 | | 0.00007 |
| Separation Problems | ENSCAFG00000001251 | | 0.00007 |
| Separation Problems | ENSCAFG00000013926 | *FBXO33* | 0.00009 |
| Separation Problems | ENSCAFG00000016354 | *STIM2* | 0.00012 |
| Separation Problems | ENSCAFG00000016568 | *ADAM10* | 0.00013 |
| Separation Problems | ENSCAFG00000030067 | | 0.00014 |

| | | | |
|---|---|---|---|
| Separation Problems | ENSCAFG00000008827 | *ALMS1* | 0.00014 |
| Separation Problems | ENSCAFG00000000334 | *ADAMTS2* | 0.00018 |
| Separation Problems | ENSCAFG00000036684 | | 0.00020 |
| Separation Problems | ENSCAFG00000000767 | *CELSR1* | 0.00025 |
| Separation Problems | ENSCAFG00000005270 | *CFAP61* | 0.00027 |
| Separation Problems | ENSCAFG00000018100 | *SCAPER* | 0.00031 |
| Separation Problems | ENSCAFG00000001161 | *OBSCN* | 0.00040 |
| Separation Problems | ENSCAFG00000010377 | | 0.00041 |
| Separation Problems | ENSCAFG00000031802 | *ISPD* | 0.00057 |
| Separation Problems | ENSCAFG00000004545 | *TRDMT1* | 0.00064 |
| Separation Problems | ENSCAFG00000005720 | *CLPB* | 0.00071 |
| Separation Problems | ENSCAFG00000017765 | *GRIA1* | 0.00078 |
| Separation Problems | ENSCAFG00000013410 | *RGL1* | 0.00086 |
| Separation Problems | ENSCAFG00000002762 | *EML6* | 0.00088 |
| Separation Problems | ENSCAFG00000038028 | | 0.00089 |
| Separation Problems | ENSCAFG00000038198 | | 0.00096 |
| Separation Problems | ENSCAFG00000015993 | *DCP1B* | 0.00159 |
| Separation Problems | ENSCAFG00000000164 | *SMAD4* | 0.00179 |
| Separation Problems | ENSCAFG00000010026 | *SMARCAD1* | 0.00226 |
| Separation Problems | ENSCAFG00000003752 | | 0.00237 |
| Separation Problems | ENSCAFG00000006103 | *NEK11* | 0.00274 |
| Separation Problems | ENSCAFG00000039072 | | 0.00286 |
| Separation Problems | ENSCAFG00000022418 | *RF00100* | 0.00293 |
| Separation Problems | ENSCAFG00000035474 | | 0.00299 |
| Separation Problems | ENSCAFG00000009831 | *PRMT3* | 0.00317 |
| Separation Problems | ENSCAFG00000016511 | *RFX1* | 0.00352 |
| Separation Problems | ENSCAFG00000032668 | *SPOCK1* | 0.00362 |
| Separation Problems | ENSCAFG00000023562 | *DMD* | 0.00368 |
| Separation Problems | ENSCAFG00000009799 | *NCAPD3* | 0.00420 |

| | | | |
|---|---|---|---|
| Separation Problems | ENSCAFG00000006042 | *SLC23A2* | 0.00447 |
| Separation Problems | ENSCAFG00000000169 | *MRO* | 0.00510 |
| Separation Problems | ENSCAFG00000006762 | *ST18* | 0.00663 |
| Separation Problems | ENSCAFG00000016588 | *C20H19orf57* | 0.00667 |
| Separation Problems | ENSCAFG00000008413 | *EPB41L1* | 0.00685 |
| Separation Problems | ENSCAFG00000015742 | | 0.00731 |
| Separation Problems | ENSCAFG00000000945 | *MAN1A1* | 0.00776 |
| Separation Problems | ENSCAFG00000004092 | *MTMR2* | 0.00787 |
| Separation Problems | ENSCAFG00000029321 | *WNT5A* | 0.00804 |
| Separation Problems | ENSCAFG00000015686 | | 0.00805 |
| Separation Problems | ENSCAFG00000013052 | *TANC2* | 0.00888 |
| Separation Problems | ENSCAFG00000017787 | | 0.00890 |
| Separation Problems | ENSCAFG00000039437 | | 0.00912 |
| Separation Problems | ENSCAFG00000013503 | *DGKG* | 0.00932 |
| Separation Problems | ENSCAFG00000012301 | *TNS3* | 0.01061 |
| Separation Problems | ENSCAFG00000034552 | | 0.01130 |
| Separation Problems | ENSCAFG00000007098 | *CYP7A1* | 0.01166 |
| Separation Problems | ENSCAFG00000007568 | *CPA6* | 0.01218 |
| Separation Problems | ENSCAFG00000030145 | *AUH* | 0.01299 |
| Separation Problems | ENSCAFG00000015862 | *FAM214A* | 0.01412 |
| Separation Problems | ENSCAFG00000002091 | *RCL1* | 0.01687 |
| Separation Problems | ENSCAFG00000033196 | | 0.01753 |
| Separation Problems | ENSCAFG00000014146 | *ZNF385B* | 0.01926 |
| Separation Problems | ENSCAFG00000001312 | *CHCHD3* | 0.01958 |
| Separation Problems | ENSCAFG00000005450 | *KAT14* | 0.02136 |
| Separation Problems | ENSCAFG00000014112 | *KIF15* | 0.02200 |
| Separation Problems | ENSCAFG00000008340 | *RACGAP1* | 0.02247 |
| Separation Problems | ENSCAFG00000016310 | *IKZF3* | 0.02598 |
| Separation Problems | ENSCAFG00000008846 | *FBXO41* | 0.02818 |

| | | | |
|---|---|---|---|
| Separation Problems | ENSCAFG00000006520 | *KCNG3* | 0.02862 |
| Separation Problems | ENSCAFG00000006812 | *TMEM132D* | 0.02977 |
| Separation Problems | ENSCAFG00000016449 | *FCRL1* | 0.03228 |
| Separation Problems | ENSCAFG00000004057 | *SRPK2* | 0.03257 |
| Separation Problems | ENSCAFG00000011585 | *PCNX2* | 0.03289 |
| Separation Problems | ENSCAFG00000024996 | | 0.03645 |
| Separation Problems | ENSCAFG00000031998 | *RF00026* | 0.03646 |
| Separation Problems | ENSCAFG00000006668 | *KBTBD8* | 0.03722 |
| Separation Problems | ENSCAFG00000032790 | | 0.03817 |
| Separation Problems | ENSCAFG00000040189 | | 0.03987 |
| Separation Problems | ENSCAFG00000001474 | | 0.04021 |
| Separation Problems | ENSCAFG00000015800 | *MYO5A* | 0.04120 |
| Separation Problems | ENSCAFG00000000557 | *CSNK1G3* | 0.04168 |
| Separation Problems | ENSCAFG00000034267 | | 0.04483 |
| Separation Problems | ENSCAFG00000033275 | | 0.04548 |
| Separation Problems | ENSCAFG00000010881 | *IGF1R* | 0.04783 |
| Separation Problems | ENSCAFG00000006743 | *PCMTD1* | 0.04858 |
| Stranger Aggression | ENSCAFG00000000079 | *RELCH* | < 0.00001 |
| Stranger Aggression | ENSCAFG00000003678 | *CCNY* | < 0.00001 |
| Stranger Aggression | ENSCAFG00000004408 | *CAB39L* | < 0.00001 |
| Stranger Aggression | ENSCAFG00000039115 | | < 0.00001 |
| Stranger Aggression | ENSCAFG00000009971 | *ARMH3* | < 0.00001 |
| Stranger Aggression | ENSCAFG00000004341 | *SETDB2* | < 0.00001 |
| Stranger Aggression | ENSCAFG00000010619 | *BICD1* | < 0.00001 |
| Stranger Aggression | ENSCAFG00000004325 | *KPNA3* | < 0.00001 |
| Stranger Aggression | ENSCAFG00000031429 | *GNA14* | < 0.00001 |
| Stranger Aggression | ENSCAFG00000011415 | *MCUB* | < 0.00001 |
| Stranger Aggression | ENSCAFG00000004379 | *FNDC3A* | < 0.00001 |
| Stranger Aggression | ENSCAFG00000004478 | *LRCH1* | < 0.00001 |

| | | | |
|---|---|---|---|
| Stranger Aggression | ENSCAFG00000002257 | *ICA1* | < 0.00001 |
| Stranger Aggression | ENSCAFG00000004436 | *RB1* | < 0.00001 |
| Stranger Aggression | ENSCAFG00000003737 | *PARD3* | < 0.00001 |
| Stranger Aggression | ENSCAFG00000005667 | *NEK10* | < 0.00001 |
| Stranger Aggression | ENSCAFG00000003705 | *CUL2* | < 0.00001 |
| Stranger Aggression | ENSCAFG00000001560 | *KIF6* | < 0.00001 |
| Stranger Aggression | ENSCAFG00000013410 | *RGL1* | < 0.00001 |
| Stranger Aggression | ENSCAFG00000005954 | *SUMF1* | < 0.00001 |
| Stranger Aggression | ENSCAFG00000005326 | *UVRAG* | < 0.00001 |
| Stranger Aggression | ENSCAFG00000000353 | *STXBP5* | < 0.00001 |
| Stranger Aggression | ENSCAFG00000005707 | *OSBPL8* | < 0.00001 |
| Stranger Aggression | ENSCAFG00000001557 | *CCDC171* | < 0.00001 |
| Stranger Aggression | ENSCAFG00000035101 | | 0.00001 |
| Stranger Aggression | ENSCAFG00000000637 | *SLC12A2* | 0.00001 |
| Stranger Aggression | ENSCAFG00000004345 | *RCBTB1* | 0.00003 |
| Stranger Aggression | ENSCAFG00000002364 | *ADGRL3* | 0.00003 |
| Stranger Aggression | ENSCAFG00000014030 | *LARS2* | 0.00005 |
| Stranger Aggression | ENSCAFG00000005040 | *KLF12* | 0.00006 |
| Stranger Aggression | ENSCAFG00000034987 | | 0.00006 |
| Stranger Aggression | ENSCAFG00000020058 | *SNX7* | 0.00009 |
| Stranger Aggression | ENSCAFG00000005175 | *SCEL* | 0.00013 |
| Stranger Aggression | ENSCAFG00000036142 | | 0.00015 |
| Stranger Aggression | ENSCAFG00000034005 | | 0.00025 |
| Stranger Aggression | ENSCAFG00000040913 | | 0.00028 |
| Stranger Aggression | ENSCAFG00000006103 | *NEK11* | 0.00031 |
| Stranger Aggression | ENSCAFG00000036701 | | 0.00061 |
| Stranger Aggression | ENSCAFG00000006496 | *MITF* | 0.00062 |
| Stranger Aggression | ENSCAFG00000019027 | *LDOC1* | 0.00132 |
| Stranger Aggression | ENSCAFG00000006263 | | 0.00173 |

| | | | |
|---|---|---|---|
| Stranger Aggression | ENSCAFG00000014864 | *CASP12* | 0.00191 |
| Stranger Aggression | ENSCAFG00000001312 | *CHCHD3* | 0.00196 |
| Stranger Aggression | ENSCAFG00000024856 | *KPNA4* | 0.00199 |
| Stranger Aggression | ENSCAFG00000018920 | *CLEC16A* | 0.00229 |
| Stranger Aggression | ENSCAFG00000027303 | *RF00100* | 0.00243 |
| Stranger Aggression | ENSCAFG00000033999 | | 0.00268 |
| Stranger Aggression | ENSCAFG00000010984 | *VTI1A* | 0.00316 |
| Stranger Aggression | ENSCAFG00000038361 | | 0.00335 |
| Stranger Aggression | ENSCAFG00000007235 | *HIPK3* | 0.00370 |
| Stranger Aggression | ENSCAFG00000000507 | *VPS13B* | 0.00382 |
| Stranger Aggression | ENSCAFG00000013966 | *CPS1* | 0.00386 |
| Stranger Aggression | ENSCAFG00000034814 | | 0.00394 |
| Stranger Aggression | ENSCAFG00000001286 | *FANCC* | 0.00442 |
| Stranger Aggression | ENSCAFG00000031367 | | 0.00488 |
| Stranger Aggression | ENSCAFG00000004337 | *EBPL* | 0.00499 |
| Stranger Aggression | ENSCAFG00000004764 | *VWA8* | 0.00695 |
| Stranger Aggression | ENSCAFG00000038544 | | 0.00725 |
| Stranger Aggression | ENSCAFG00000003860 | *MFSD8* | 0.00826 |
| Stranger Aggression | ENSCAFG00000034539 | | 0.00979 |
| Stranger Aggression | ENSCAFG00000005838 | *PCCA* | 0.01052 |
| Stranger Aggression | ENSCAFG00000002335 | | 0.01067 |
| Stranger Aggression | ENSCAFG00000036116 | | 0.01085 |
| Stranger Aggression | ENSCAFG00000014191 | *PBX4* | 0.01242 |
| Stranger Aggression | ENSCAFG00000017685 | *RIT2* | 0.01286 |
| Stranger Aggression | ENSCAFG00000017571 | *CACNG3* | 0.01421 |
| Stranger Aggression | ENSCAFG00000000157 | *DCC* | 0.01453 |
| Stranger Aggression | ENSCAFG00000010942 | *CCDC91* | 0.01637 |
| Stranger Aggression | ENSCAFG00000027991 | *RF00026* | 0.01691 |
| Stranger Aggression | ENSCAFG00000006431 | *LARS* | 0.01740 |

| | | | |
|---|---|---|---|
| Stranger Aggression | ENSCAFG00000039466 | | 0.01988 |
| Stranger Aggression | ENSCAFG00000006879 | *MARCH8* | 0.02081 |
| Stranger Aggression | ENSCAFG00000037402 | | 0.02103 |
| Stranger Aggression | ENSCAFG00000024350 | *CENPP* | 0.02472 |
| Stranger Aggression | ENSCAFG00000009506 | *ABI3BP* | 0.03005 |
| Stranger Aggression | ENSCAFG00000035111 | | 0.03077 |
| Stranger Aggression | ENSCAFG00000001635 | *MLLT3* | 0.03118 |
| Stranger Aggression | ENSCAFG00000002397 | *SHB* | 0.03402 |
| Stranger Aggression | ENSCAFG00000028102 | *RF00088* | 0.03734 |
| Stranger Aggression | ENSCAFG00000037818 | | 0.03787 |
| Stranger Aggression | ENSCAFG00000035993 | | 0.03838 |
| Stranger Aggression | ENSCAFG00000002671 | *SP4* | 0.04034 |
| Stranger Aggression | ENSCAFG00000007943 | *FAM81B* | 0.04258 |
| Stranger Aggression | ENSCAFG00000035622 | | 0.04310 |
| Stranger Aggression | ENSCAFG00000026189 | *RF00091* | 0.04436 |
| Stranger Aggression | ENSCAFG00000016732 | *SIPA1L1* | 0.04922 |
| Stranger Fear | ENSCAFG00000003929 | *COG5* | < 0.00001 |
| Stranger Fear | ENSCAFG00000011082 | *CAMSAP2* | < 0.00001 |
| Stranger Fear | ENSCAFG00000001434 | *KDM4C* | < 0.00001 |
| Stranger Fear | ENSCAFG00000031429 | *GNA14* | < 0.00001 |
| Stranger Fear | ENSCAFG00000002257 | *ICA1* | < 0.00001 |
| Stranger Fear | ENSCAFG00000000693 | *ANGPT1* | < 0.00001 |
| Stranger Fear | ENSCAFG00000033485 | | < 0.00001 |
| Stranger Fear | ENSCAFG00000018024 | *TRAPPC8* | < 0.00001 |
| Stranger Fear | ENSCAFG00000006462 | *UBXN8* | 0.00001 |
| Stranger Fear | ENSCAFG00000002667 | | 0.00002 |
| Stranger Fear | ENSCAFG00000023580 | | 0.00002 |
| Stranger Fear | ENSCAFG00000015395 | | 0.00002 |
| Stranger Fear | ENSCAFG00000034027 | | 0.00004 |

| | | | |
|---|---|---|---|
| Stranger Fear | ENSCAFG00000015131 | *CEP126* | 0.00004 |
| Stranger Fear | ENSCAFG00000034629 | | 0.00004 |
| Stranger Fear | ENSCAFG00000013221 | | 0.00014 |
| Stranger Fear | ENSCAFG00000035111 | | 0.00024 |
| Stranger Fear | ENSCAFG00000004568 | *CUBN* | 0.00030 |
| Stranger Fear | ENSCAFG00000009280 | *PTPRT* | 0.00034 |
| Stranger Fear | ENSCAFG00000023938 | *GPR158* | 0.00036 |
| Stranger Fear | ENSCAFG00000015679 | *RGS7* | 0.00038 |
| Stranger Fear | ENSCAFG00000003153 | *VPS54* | 0.00040 |
| Stranger Fear | ENSCAFG00000010811 | *LRRC28* | 0.00045 |
| Stranger Fear | ENSCAFG00000002546 | *CAMKMT* | 0.00052 |
| Stranger Fear | ENSCAFG00000036261 | | 0.00064 |
| Stranger Fear | ENSCAFG00000001988 | *DMRT2* | 0.00066 |
| Stranger Fear | ENSCAFG00000008669 | *TMEM144* | 0.00067 |
| Stranger Fear | ENSCAFG00000037968 | | 0.00074 |
| Stranger Fear | ENSCAFG00000017227 | *GABRG2* | 0.00075 |
| Stranger Fear | ENSCAFG00000009159 | *TRIO* | 0.00099 |
| Stranger Fear | ENSCAFG00000014030 | *LARS2* | 0.00125 |
| Stranger Fear | ENSCAFG00000013671 | *EFL1* | 0.00133 |
| Stranger Fear | ENSCAFG00000004942 | | 0.00133 |
| Stranger Fear | ENSCAFG00000015967 | *MTHFD1* | 0.00170 |
| Stranger Fear | ENSCAFG00000019296 | *METTL16* | 0.00209 |
| Stranger Fear | ENSCAFG00000006103 | *NEK11* | 0.00238 |
| Stranger Fear | ENSCAFG00000000337 | *PPM1H* | 0.00274 |
| Stranger Fear | ENSCAFG00000009173 | *EXOC2* | 0.00287 |
| Stranger Fear | ENSCAFG00000016732 | *SIPA1L1* | 0.00430 |
| Stranger Fear | ENSCAFG00000010766 | *TESMIN* | 0.00431 |
| Stranger Fear | ENSCAFG00000036396 | | 0.00601 |
| Stranger Fear | ENSCAFG00000004408 | *CAB39L* | 0.00625 |

| | | | |
|---|---|---|---|
| Stranger Fear | ENSCAFG00000023546 | *TCTN2* | 0.00747 |
| Stranger Fear | ENSCAFG00000036050 | | 0.00755 |
| Stranger Fear | ENSCAFG00000035085 | | 0.00778 |
| Stranger Fear | ENSCAFG00000032631 | | 0.00796 |
| Stranger Fear | ENSCAFG00000000696 | *RSPO2* | 0.00796 |
| Stranger Fear | ENSCAFG00000006326 | *KCTD16* | 0.00832 |
| Stranger Fear | ENSCAFG00000009140 | *GALNT13* | 0.00901 |
| Stranger Fear | ENSCAFG00000002552 | *INVS* | 0.01058 |
| Stranger Fear | ENSCAFG00000037907 | | 0.01203 |
| Stranger Fear | ENSCAFG00000006666 | | 0.01503 |
| Stranger Fear | ENSCAFG00000004243 | *SLC36A4* | 0.01576 |
| Stranger Fear | ENSCAFG00000038028 | | 0.01613 |
| Stranger Fear | ENSCAFG00000035420 | | 0.01631 |
| Stranger Fear | ENSCAFG00000008738 | *TM9SF3* | 0.01636 |
| Stranger Fear | ENSCAFG00000010619 | *BICD1* | 0.01716 |
| Stranger Fear | ENSCAFG00000012632 | *OR51S1* | 0.02099 |
| Stranger Fear | ENSCAFG00000014618 | *ARPC2* | 0.02939 |
| Stranger Fear | ENSCAFG00000000974 | *DERL1* | 0.03457 |
| Stranger Fear | ENSCAFG00000017314 | *MBTD1* | 0.03882 |
| Stranger Fear | ENSCAFG00000003582 | *VPS41* | 0.03909 |
| Stranger Fear | ENSCAFG00000018267 | *TOP3A* | 0.03910 |
| Stranger Fear | ENSCAFG00000009893 | *DTNBP1* | 0.04464 |
| Stranger Fear | ENSCAFG00000025134 | | 0.04558 |
| Stranger Fear | ENSCAFG00000003995 | *FYN* | 0.04670 |
| Stranger Fear | ENSCAFG00000005720 | *CLPB* | 0.04898 |
| Stranger Fear | ENSCAFG00000039313 | | 0.04959 |
| Stranger Fear | ENSCAFG00000035776 | | 0.04968 |
| Touch Sensitivity | ENSCAFG00000006013 | *CNTN4* | < 0.00001 |
| Touch Sensitivity | ENSCAFG00000017362 | *NUP210L* | < 0.00001 |

| | | | |
|---|---|---|---|
| Touch Sensitivity | ENSCAFG00000002796 | *CCDC88A* | < 0.00001 |
| Touch Sensitivity | ENSCAFG00000033155 | | < 0.00001 |
| Touch Sensitivity | ENSCAFG00000005998 | *LHFPL6* | < 0.00001 |
| Touch Sensitivity | ENSCAFG00000006791 | | < 0.00001 |
| Touch Sensitivity | ENSCAFG00000000696 | *RSPO2* | < 0.00001 |
| Touch Sensitivity | ENSCAFG00000015625 | *FMN2* | < 0.00001 |
| Touch Sensitivity | ENSCAFG00000028659 | | < 0.00001 |
| Touch Sensitivity | ENSCAFG00000000090 | *MC4R* | < 0.00001 |
| Touch Sensitivity | ENSCAFG00000033127 | | < 0.00001 |
| Touch Sensitivity | ENSCAFG00000036746 | | < 0.00001 |
| Touch Sensitivity | ENSCAFG00000002762 | *EML6* | < 0.00001 |
| Touch Sensitivity | ENSCAFG00000034355 | | < 0.00001 |
| Touch Sensitivity | ENSCAFG00000035449 | | < 0.00001 |
| Touch Sensitivity | ENSCAFG00000001754 | *POT1* | < 0.00001 |
| Touch Sensitivity | ENSCAFG00000003479 | | < 0.00001 |
| Touch Sensitivity | ENSCAFG00000020187 | *HFM1* | < 0.00001 |
| Touch Sensitivity | ENSCAFG00000010355 | *ICE1* | < 0.00001 |
| Touch Sensitivity | ENSCAFG00000005720 | *CLPB* | < 0.00001 |
| Touch Sensitivity | ENSCAFG00000017248 | *UBAP2L* | < 0.00001 |
| Touch Sensitivity | ENSCAFG00000002661 | *SMC2* | 0.00002 |
| Touch Sensitivity | ENSCAFG00000005916 | *ANKRD28* | 0.00002 |
| Touch Sensitivity | ENSCAFG00000032632 | *WNT7A* | 0.00003 |
| Touch Sensitivity | ENSCAFG00000017645 | *MYO9A* | 0.00003 |
| Touch Sensitivity | ENSCAFG00000007822 | *BDP1* | 0.00004 |
| Touch Sensitivity | ENSCAFG00000008185 | *ARFIP1* | 0.00005 |
| Touch Sensitivity | ENSCAFG00000000737 | *CDC42SE2* | 0.00005 |
| Touch Sensitivity | ENSCAFG00000015877 | *SH2D4B* | 0.00005 |
| Touch Sensitivity | ENSCAFG00000011585 | *PCNX2* | 0.00006 |
| Touch Sensitivity | ENSCAFG00000016568 | *ADAM10* | 0.00008 |

| | | | |
|---|---|---|---|
| Touch Sensitivity | ENSCAFG00000004454 | *SUCLA2* | 0.00016 |
| Touch Sensitivity | ENSCAFG00000000569 | *ZNF608* | 0.00019 |
| Touch Sensitivity | ENSCAFG00000000884 | *WDR27* | 0.00025 |
| Touch Sensitivity | ENSCAFG00000035409 | | 0.00029 |
| Touch Sensitivity | ENSCAFG00000018665 | *POLDIP2* | 0.00039 |
| Touch Sensitivity | ENSCAFG00000005775 | *BIRC6* | 0.00043 |
| Touch Sensitivity | ENSCAFG00000017651 | *DNM2* | 0.00046 |
| Touch Sensitivity | ENSCAFG00000033639 | | 0.00048 |
| Touch Sensitivity | ENSCAFG00000018663 | *TNFAIP1* | 0.00049 |
| Touch Sensitivity | ENSCAFG00000017598 | *SHISA6* | 0.00058 |
| Touch Sensitivity | ENSCAFG00000006520 | *KCNG3* | 0.00072 |
| Touch Sensitivity | ENSCAFG00000014275 | *SOS2* | 0.00086 |
| Touch Sensitivity | ENSCAFG00000033474 | | 0.00087 |
| Touch Sensitivity | ENSCAFG00000030928 | | 0.00099 |
| Touch Sensitivity | ENSCAFG00000018564 | *GRIA3* | 0.00104 |
| Touch Sensitivity | ENSCAFG00000025531 | | 0.00118 |
| Touch Sensitivity | ENSCAFG00000037655 | | 0.00124 |
| Touch Sensitivity | ENSCAFG00000038140 | | 0.00136 |
| Touch Sensitivity | ENSCAFG00000017606 | *UNC79* | 0.00149 |
| Touch Sensitivity | ENSCAFG00000023460 | *FBRSL1* | 0.00164 |
| Touch Sensitivity | ENSCAFG00000005600 | *TASP1* | 0.00175 |
| Touch Sensitivity | ENSCAFG00000015383 | *TNFSF10* | 0.00187 |
| Touch Sensitivity | ENSCAFG00000040274 | | 0.00205 |
| Touch Sensitivity | ENSCAFG00000008319 | *CHODL* | 0.00230 |
| Touch Sensitivity | ENSCAFG00000035085 | | 0.00278 |
| Touch Sensitivity | ENSCAFG00000018366 | *KIAA1210* | 0.00299 |
| Touch Sensitivity | ENSCAFG00000038398 | | 0.00388 |
| Touch Sensitivity | ENSCAFG00000038034 | | 0.00501 |
| Touch Sensitivity | ENSCAFG00000020112 | *ABCD3* | 0.00568 |

| | | | |
|---|---|---|---|
| Touch Sensitivity | ENSCAFG00000004130 | *NEBL* | 0.00572 |
| Touch Sensitivity | ENSCAFG00000018591 | *ARHGAP28* | 0.00583 |
| Touch Sensitivity | ENSCAFG00000002753 | *GPNMB* | 0.00634 |
| Touch Sensitivity | ENSCAFG00000034887 | | 0.00656 |
| Touch Sensitivity | ENSCAFG00000003705 | *CUL2* | 0.00671 |
| Touch Sensitivity | ENSCAFG00000000828 | *RPS6KA2* | 0.00844 |
| Touch Sensitivity | ENSCAFG00000035826 | | 0.00869 |
| Touch Sensitivity | ENSCAFG00000018890 | *TXNDC11* | 0.00873 |
| Touch Sensitivity | ENSCAFG00000039677 | | 0.00917 |
| Touch Sensitivity | ENSCAFG00000031628 | | 0.01009 |
| Touch Sensitivity | ENSCAFG00000016310 | *IKZF3* | 0.01127 |
| Touch Sensitivity | ENSCAFG00000010897 | *ARHGEF38* | 0.01138 |
| Touch Sensitivity | ENSCAFG00000005403 | *LRP1B* | 0.01183 |
| Touch Sensitivity | ENSCAFG00000022906 | *RF00026* | 0.01333 |
| Touch Sensitivity | ENSCAFG00000024183 | | 0.01515 |
| Touch Sensitivity | ENSCAFG00000033119 | | 0.01546 |
| Touch Sensitivity | ENSCAFG00000001173 | *FAM135B* | 0.01767 |
| Touch Sensitivity | ENSCAFG00000017369 | *PCTP* | 0.02012 |
| Touch Sensitivity | ENSCAFG00000017197 | *ATP8B2* | 0.02366 |
| Touch Sensitivity | ENSCAFG00000005061 | *LMO7* | 0.02381 |
| Touch Sensitivity | ENSCAFG00000000719 | *NUDCD1* | 0.02398 |
| Touch Sensitivity | ENSCAFG00000030884 | *APOPT1* | 0.02658 |
| Touch Sensitivity | ENSCAFG00000000068 | *BCL2* | 0.02676 |
| Touch Sensitivity | ENSCAFG00000010682 | *SP110* | 0.02734 |
| Touch Sensitivity | ENSCAFG00000001700 | *SND1* | 0.02783 |
| Touch Sensitivity | ENSCAFG00000006050 | *MYO16* | 0.02814 |
| Touch Sensitivity | ENSCAFG00000038632 | | 0.03090 |
| Touch Sensitivity | ENSCAFG00000036101 | | 0.03116 |
| Touch Sensitivity | ENSCAFG00000008744 | *RASGRF2* | 0.03174 |

| | | | |
|---|---|---|---|
| Touch Sensitivity | ENSCAFG00000018100 | *SCAPER* | 0.03393 |
| Touch Sensitivity | ENSCAFG00000009227 | *WDR41* | 0.03573 |
| Touch Sensitivity | ENSCAFG00000009218 | *MORN4* | 0.03630 |
| Touch Sensitivity | ENSCAFG00000040835 | | 0.03944 |
| Touch Sensitivity | ENSCAFG00000011657 | *EAF2* | 0.04232 |
| Touch Sensitivity | ENSCAFG00000004249 | *ARHGAP21* | 0.04351 |
| Touch Sensitivity | ENSCAFG00000010229 | *DOCK3* | 0.04478 |
| Trainability | ENSCAFG00000018564 | *GRIA3* | < 0.00001 |
| Trainability | ENSCAFG00000000090 | *MC4R* | < 0.00001 |
| Trainability | ENSCAFG00000007420 | *CBFA2T2* | < 0.00001 |
| Trainability | ENSCAFG00000014257 | *PPP2R2C* | < 0.00001 |
| Trainability | ENSCAFG00000018858 | *TARS* | < 0.00001 |
| Trainability | ENSCAFG00000000011 | *NFATC1* | < 0.00001 |
| Trainability | ENSCAFG00000000133 | *WDR7* | < 0.00001 |
| Trainability | ENSCAFG00000029157 | *NOVA1* | < 0.00001 |
| Trainability | ENSCAFG00000035434 | | < 0.00001 |
| Trainability | ENSCAFG00000033089 | | < 0.00001 |
| Trainability | ENSCAFG00000018849 | *SNX29* | < 0.00001 |
| Trainability | ENSCAFG00000000353 | *STXBP5* | < 0.00001 |
| Trainability | ENSCAFG00000003373 | *DGKI* | < 0.00001 |
| Trainability | ENSCAFG00000004545 | *TRDMT1* | < 0.00001 |
| Trainability | ENSCAFG00000037352 | | < 0.00001 |
| Trainability | ENSCAFG00000019030 | *ABAT* | < 0.00001 |
| Trainability | ENSCAFG00000035215 | | < 0.00001 |
| Trainability | ENSCAFG00000009912 | *ERG* | < 0.00001 |
| Trainability | ENSCAFG00000003580 | *GRIK2* | < 0.00001 |
| Trainability | ENSCAFG00000001700 | *SND1* | < 0.00001 |
| Trainability | ENSCAFG00000000522 | *SNCAIP* | < 0.00001 |
| Trainability | ENSCAFG00000038771 | | < 0.00001 |

| | | | |
|---|---|---|---|
| Trainability | ENSCAFG00000032608 | *LURAP1L* | < 0.00001 |
| Trainability | ENSCAFG00000027467 | *RF00026* | < 0.00001 |
| Trainability | ENSCAFG00000038087 | | < 0.00001 |
| Trainability | ENSCAFG00000008333 | *TMEM161B* | < 0.00001 |
| Trainability | ENSCAFG00000039561 | | < 0.00001 |
| Trainability | ENSCAFG00000008211 | *CACNA2D3* | < 0.00001 |
| Trainability | ENSCAFG00000035520 | | < 0.00001 |
| Trainability | ENSCAFG00000035311 | | < 0.00001 |
| Trainability | ENSCAFG00000011725 | *ATRNL1* | 0.00001 |
| Trainability | ENSCAFG00000000582 | *GRAMD2B* | 0.00001 |
| Trainability | ENSCAFG00000033361 | | 0.00001 |
| Trainability | ENSCAFG00000028708 | *SCD5* | 0.00001 |
| Trainability | ENSCAFG00000018972 | *EFCAB5* | 0.00001 |
| Trainability | ENSCAFG00000028601 | | 0.00001 |
| Trainability | ENSCAFG00000028075 | | 0.00001 |
| Trainability | ENSCAFG00000015351 | *BAIAP2L1* | 0.00001 |
| Trainability | ENSCAFG00000033559 | | 0.00001 |
| Trainability | ENSCAFG00000013852 | *MATN2* | 0.00002 |
| Trainability | ENSCAFG00000010692 | *USF3* | 0.00002 |
| Trainability | ENSCAFG00000028239 | *RF00003* | 0.00002 |
| Trainability | ENSCAFG00000034669 | | 0.00003 |
| Trainability | ENSCAFG00000000012 | *ATP9B* | 0.00003 |
| Trainability | ENSCAFG00000006791 | | 0.00003 |
| Trainability | ENSCAFG00000009081 | *EPHA6* | 0.00003 |
| Trainability | ENSCAFG00000031001 | *RF00026* | 0.00004 |
| Trainability | ENSCAFG00000033886 | | 0.00007 |
| Trainability | ENSCAFG00000004157 | *RAB7A* | 0.00008 |
| Trainability | ENSCAFG00000005949 | *HACL1* | 0.00009 |
| Trainability | ENSCAFG00000014096 | | 0.00010 |

| | | | |
|---|---|---|---|
| Trainability | ENSCAFG00000000745 | *TBC1D22A* | 0.00011 |
| Trainability | ENSCAFG00000032743 | | 0.00027 |
| Trainability | ENSCAFG00000039328 | | 0.00032 |
| Trainability | ENSCAFG00000033071 | | 0.00034 |
| Trainability | ENSCAFG00000034355 | | 0.00037 |
| Trainability | ENSCAFG00000009052 | *LIN54* | 0.00040 |
| Trainability | ENSCAFG00000005849 | *TBC1D5* | 0.00077 |
| Trainability | ENSCAFG00000001116 | *SAMD3* | 0.00080 |
| Trainability | ENSCAFG00000011266 | *TDRD1* | 0.00081 |
| Trainability | ENSCAFG00000039318 | | 0.00091 |
| Trainability | ENSCAFG00000015877 | *SH2D4B* | 0.00104 |
| Trainability | ENSCAFG00000002364 | *ADGRL3* | 0.00109 |
| Trainability | ENSCAFG00000003651 | *CSMD2* | 0.00118 |
| Trainability | ENSCAFG00000027188 | *RF00026* | 0.00144 |
| Trainability | ENSCAFG00000005326 | *UVRAG* | 0.00149 |
| Trainability | ENSCAFG00000014755 | *SAMD7* | 0.00206 |
| Trainability | ENSCAFG00000008199 | *FMN1* | 0.00213 |
| Trainability | ENSCAFG00000039816 | | 0.00229 |
| Trainability | ENSCAFG00000009595 | *PKD2* | 0.00236 |
| Trainability | ENSCAFG00000034093 | | 0.00251 |
| Trainability | ENSCAFG00000030938 | | 0.00287 |
| Trainability | ENSCAFG00000006989 | *DEFB119* | 0.00337 |
| Trainability | ENSCAFG00000038450 | | 0.00408 |
| Trainability | ENSCAFG00000006046 | *COL6A5* | 0.00452 |
| Trainability | ENSCAFG00000001425 | *GKAP1* | 0.00460 |
| Trainability | ENSCAFG00000025531 | | 0.00478 |
| Trainability | ENSCAFG00000012544 | *COPA* | 0.00563 |
| Trainability | ENSCAFG00000015742 | | 0.00580 |
| Trainability | ENSCAFG00000013288 | *ADAMTSL3* | 0.00648 |

| | | | |
|---|---|---|---|
| Trainability | ENSCAFG00000038280 | | 0.00803 |
| Trainability | ENSCAFG00000017514 | *PCDH19* | 0.00828 |
| Trainability | ENSCAFG00000014358 | *CASK* | 0.00852 |
| Trainability | ENSCAFG00000030561 | | 0.00971 |
| Trainability | ENSCAFG00000039541 | | 0.00976 |
| Trainability | ENSCAFG00000009991 | | 0.01028 |
| Trainability | ENSCAFG00000030656 | | 0.01161 |
| Trainability | ENSCAFG00000025514 | *C28H10orf53* | 0.01225 |
| Trainability | ENSCAFG00000000637 | *SLC12A2* | 0.01375 |
| Trainability | ENSCAFG00000002023 | *RCAN2* | 0.01463 |
| Trainability | ENSCAFG00000008790 | *CTNNBL1* | 0.01486 |
| Trainability | ENSCAFG00000008851 | *HECTD4* | 0.01540 |
| Trainability | ENSCAFG00000016024 | *FBXL18* | 0.01563 |
| Trainability | ENSCAFG00000038274 | | 0.01620 |
| Trainability | ENSCAFG00000010235 | *ARHGAP32* | 0.01649 |
| Trainability | ENSCAFG00000010827 | *TTC23* | 0.01714 |
| Trainability | ENSCAFG00000018267 | *TOP3A* | 0.01888 |
| Trainability | ENSCAFG00000011571 | *SOX5* | 0.01969 |
| Trainability | ENSCAFG00000007837 | *KNTC1* | 0.02249 |
| Trainability | ENSCAFG00000017137 | *TENM2* | 0.02264 |
| Trainability | ENSCAFG00000005056 | *SCN10A* | 0.02269 |
| Trainability | ENSCAFG00000005839 | *SATB1* | 0.02311 |
| Trainability | ENSCAFG00000011796 | *MCF2L2* | 0.02318 |
| Trainability | ENSCAFG00000011940 | *PARP9* | 0.02325 |
| Trainability | ENSCAFG00000035103 | | 0.02432 |
| Trainability | ENSCAFG00000033801 | | 0.02483 |
| Trainability | ENSCAFG00000010996 | *CCDC15* | 0.02506 |
| Trainability | ENSCAFG00000005180 | *MRPL33* | 0.02591 |
| Trainability | ENSCAFG00000005090 | *AANAT* | 0.02730 |

| | | | |
|---|---|---|---|
| Trainability | ENSCAFG00000013218 | *EHHADH* | 0.02775 |
| Trainability | ENSCAFG00000032922 | | 0.03014 |
| Trainability | ENSCAFG00000014001 | *TOGARAM1* | 0.03024 |
| Trainability | ENSCAFG00000000507 | *VPS13B* | 0.03074 |
| Trainability | ENSCAFG00000023072 | *POC1A* | 0.03148 |
| Trainability | ENSCAFG00000019062 | *DYM* | 0.03159 |
| Trainability | ENSCAFG00000035015 | | 0.03209 |
| Trainability | ENSCAFG00000010120 | *RAD54L2* | 0.03325 |
| Trainability | ENSCAFG00000002025 | *GCC2* | 0.03546 |
| Trainability | ENSCAFG00000023015 | *SERPINB12* | 0.03693 |
| Trainability | ENSCAFG00000014045 | *LRRC20* | 0.03778 |
| Trainability | ENSCAFG00000032255 | *RNF175* | 0.03865 |
| Trainability | ENSCAFG00000014108 | *OSBPL1A* | 0.03873 |
| Trainability | ENSCAFG00000034449 | | 0.04114 |
| Trainability | ENSCAFG00000016511 | *RFX1* | 0.04238 |
| Trainability | ENSCAFG00000015716 | *RAC1* | 0.04262 |
| Trainability | ENSCAFG00000000867 | *RAD50* | 0.04359 |
| Trainability | ENSCAFG00000010879 | *IGSF11* | 0.04743 |
| Trainability | ENSCAFG00000034432 | | 0.04785 |

**Table S4. Genes associated with breed differences in dog behavior that have also been implicated in differences between tame and aggressive foxes, phenotypic changes during domestication, previous studies of dog behavior, or parallel evolution in dogs and humans.**

| Associations from Previous Studies | Genes |
|---|---|
| genes implicated in behavioral differences between foxes bred for tameness or aggression (1-3) | *ABI3BP, ACTN2, ADAMTS2, AKAP9, ALMS1, ANO2, ATOH8, ATRN, BIRC6, CACNA1C, CACNA1E, CAMK4, CCDC148, CD101, CEP85L, CUX1, DCP1B, DIP2B, DMXL1, EDIL3, ENSCAFG00000005180, FANCC, FBXL18, FBXO41, FBXW4, FCRL1, FOXK1, GALNT13, GRAMD1C, HAT1, HNRNPLL, IFRD1, IGF1R, IL1R2, KIAA1958, KLF12, LAMA2, LMO7, LRRC28, MCM9, MKL1, MLIP, MTMR1, NEBL, NID2, OPCML, PCNX2, PDE7B, PLEKHA5, PLXNA2, PRPF3, PTPRT, RAB5A, RASGRF2, RBFOX1, RYR2, SATB1, SH2D4A, SLC35F4, SLC4A10, SND1, SOX5, SPOCK1, SPTLC3, SRGAP2, STK32A, TASP1, TBR1, TMEM161B, TMEM164, TTC23, TTC7A, UNC45A, WNT5A, ZRANB3* |
| Morphological traits implicated in domestication, including pigmentation, coat coloration, tail curl, ear morphology, and snout length (4-6) | *CAPN6, CDC37L1, DMD, FGF5, GLIS3, MITF, MSRB3, RCL1, RSPO2, SRGAP1* |
| Candidate dog domestication genes (7-9) | *ABAT, APOPT1, ASIP, CADPS2, DEFB119, DOCK2, PDE7B, RYR3, SLC24A4, TMEM132D* |
| Genes under selection in both dog domestication and human evolution (10) | *CENPP, LRPPRC* |
| Dog behavioral traits (6, 11) | *ENOX2, GPC3, GPC4, MSRB3* |

**Table S5. Genes associated with dog behavioral traits and potentially related human traits previously associated with these genes.**

| Dog Behavioral Trait | Gene(s) | Human Phenotypes Associated with Same Genes |
|---|---|---|
| **Chasing** | GRIK2, CDH10 | Obsessive-compulsive disorder (12, 13) |
| | NT5C2, SORCS2, PTPRG, MLIP | Attention deficit hyperactivity disorder (14-20) |
| **Dog Aggression** | CPNE4, OPCML | Aggressive behavior (21, 22) |
| **Energy** | TMEM132D, AGMO | Heart rate (23, 24) |
| | SNX29 | Daytime rest (25) |
| | CACNA2D3 | Sleep duration (25) |
| **Excitability** | *PDE11A, ARHGAP10* | Resting heart rate (26, 27) |
| | CSMD3 | Temperament (28) |
| **Nonsocial Fear** | PTPRD | Temperament (28) |
| | CACNA1C | Startle response (29) |
| **Stranger Fear** | CAMKMT | Anxiety disorder (30) |
| **Trainability** | *ERG, SNX29*<br>*CSMD2* | Intelligence (31-34) |
| | *ATRNL1* | Information processing speed (35) |

Table S6. Significant Gene Ontology (GO) terms from enrichment analyses using SNPs in the gene to derive gene-level p values (meta-analysis, Fisher's method).

| *Behavioral Trait* | *GO ID* | *Term* | *p value* |
|---|---|---|---|
| Attach & Atn Seeking | GO:0032367 | intracellular cholesterol transport | 0.00150 |
| Attach & Atn Seeking | GO:0060047 | heart contraction | 0.00310 |
| Attach & Atn Seeking | GO:0007178 | transmembrane receptor protein serine/th... | 0.00360 |
| Attach & Atn Seeking | GO:0008608 | attachment of spindle microtubules to ki... | 0.00470 |
| Attach & Atn Seeking | GO:0002093 | auditory receptor cell morphogenesis | 0.00580 |
| Attach & Atn Seeking | GO:0021984 | adenohypophysis development | 0.00580 |
| Attach & Atn Seeking | GO:0048172 | regulation of short-term neuronal synapt... | 0.00580 |
| Attach & Atn Seeking | GO:1903861 | positive regulation of dendrite extensio... | 0.00580 |
| Attach & Atn Seeking | GO:0072599 | establishment of protein localization to... | 0.00710 |
| Attach & Atn Seeking | GO:0006914 | autophagy | 0.00730 |
| Attach & Atn Seeking | GO:0048791 | calcium ion-regulated exocytosis of neur... | 0.00760 |
| Attach & Atn Seeking | GO:0071786 | endoplasmic reticulum tubular network or... | 0.00760 |
| Attach & Atn Seeking | GO:0034198 | cellular response to amino acid starvati... | 0.00930 |
| Attach & Atn Seeking | GO:0071467 | cellular response to pH | 0.00960 |
| Attach & Atn Seeking | GO:0071634 | regulation of transforming growth factor... | 0.00960 |
| Attach & Atn Seeking | GO:0033173 | calcineurin-NFAT signaling cascade | 0.01140 |
| Attach & Atn Seeking | GO:0048678 | response to axon injury | 0.01150 |
| Attach & Atn Seeking | GO:0071322 | cellular response to carbohydrate stimul... | 0.01160 |
| Attach & Atn Seeking | GO:2000785 | regulation of autophagosome assembly | 0.01450 |
| Attach & Atn Seeking | GO:0010611 | regulation of cardiac muscle hypertrophy | 0.01680 |
| Attach & Atn Seeking | GO:0051491 | positive regulation of filopodium assemb... | 0.01730 |
| Attach & Atn Seeking | GO:0010955 | negative regulation of protein processin... | 0.02050 |
| Attach & Atn Seeking | GO:0042692 | muscle cell differentiation | 0.02260 |
| Attach & Atn Seeking | GO:0015872 | dopamine transport | 0.02300 |
| Attach & Atn Seeking | GO:0000380 | alternative mRNA splicing, via spliceoso... | 0.02320 |
| Attach & Atn Seeking | GO:0008219 | cell death | 0.02330 |

| | | | |
|---|---|---|---|
| Attach & Atn Seeking | GO:0006418 | tRNA aminoacylation for protein translat... | 0.02510 |
| Attach & Atn Seeking | GO:0050850 | positive regulation of calcium-mediated ... | 0.02770 |
| Attach & Atn Seeking | GO:0006623 | protein targeting to vacuole | 0.03010 |
| Attach & Atn Seeking | GO:0031349 | positive regulation of defense response | 0.03060 |
| Attach & Atn Seeking | GO:0031401 | positive regulation of protein modificat... | 0.03090 |
| Attach & Atn Seeking | GO:0043171 | peptide catabolic process | 0.03180 |
| Attach & Atn Seeking | GO:0043067 | regulation of programmed cell death | 0.03590 |
| Attach & Atn Seeking | GO:0030513 | positive regulation of BMP signaling pat... | 0.03620 |
| Attach & Atn Seeking | GO:0032467 | positive regulation of cytokinesis | 0.03620 |
| Attach & Atn Seeking | GO:0048488 | synaptic vesicle endocytosis | 0.03620 |
| Attach & Atn Seeking | GO:0060548 | negative regulation of cell death | 0.03670 |
| Attach & Atn Seeking | GO:1903363 | negative regulation of cellular protein ... | 0.03800 |
| Attach & Atn Seeking | GO:2000145 | regulation of cell motility | 0.03820 |
| Attach & Atn Seeking | GO:0042327 | positive regulation of phosphorylation | 0.03860 |
| Attach & Atn Seeking | GO:0043547 | positive regulation of GTPase activity | 0.04030 |
| Attach & Atn Seeking | GO:0090307 | mitotic spindle assembly | 0.04030 |
| Attach & Atn Seeking | GO:0031032 | actomyosin structure organization | 0.04070 |
| Attach & Atn Seeking | GO:0032024 | positive regulation of insulin secretion | 0.04080 |
| Attach & Atn Seeking | GO:0090630 | activation of GTPase activity | 0.04100 |
| Attach & Atn Seeking | GO:0031175 | neuron projection development | 0.04380 |
| Attach & Atn Seeking | GO:0070884 | regulation of calcineurin-NFAT signaling... | 0.04570 |
| Attach & Atn Seeking | GO:0002115 | store-operated calcium entry | 0.04630 |
| Attach & Atn Seeking | GO:0002374 | cytokine secretion involved in immune re... | 0.04630 |
| Attach & Atn Seeking | GO:0010447 | response to acidic pH | 0.04630 |
| Attach & Atn Seeking | GO:0014047 | glutamate secretion | 0.04630 |
| Attach & Atn Seeking | GO:0032438 | melanosome organization | 0.04630 |
| Attach & Atn Seeking | GO:0032692 | negative regulation of interleukin-1 pro... | 0.04630 |
| Attach & Atn Seeking | GO:0035994 | response to muscle stretch | 0.04630 |
| Attach & Atn Seeking | GO:0033674 | positive regulation of kinase activity | 0.04710 |

| | | | |
|---|---|---|---|
| Chasing | GO:0035335 | peptidyl-tyrosine dephosphorylation | 0.00140 |
| Chasing | GO:0071526 | semaphorin-plexin signaling pathway | 0.00250 |
| Chasing | GO:0021680 | cerebellar Purkinje cell layer developme... | 0.00310 |
| Chasing | GO:0060047 | heart contraction | 0.00310 |
| Chasing | GO:0001755 | neural crest cell migration | 0.00340 |
| Chasing | GO:0072283 | metanephric renal vesicle morphogenesis | 0.00360 |
| Chasing | GO:0006886 | intracellular protein transport | 0.00440 |
| Chasing | GO:0035268 | protein mannosylation | 0.00440 |
| Chasing | GO:0030858 | positive regulation of epithelial cell d... | 0.00470 |
| Chasing | GO:0048843 | negative regulation of axon extension in... | 0.00570 |
| Chasing | GO:0010646 | regulation of cell communication | 0.00650 |
| Chasing | GO:0046365 | monosaccharide catabolic process | 0.00760 |
| Chasing | GO:0048663 | neuron fate commitment | 0.00920 |
| Chasing | GO:0051056 | regulation of small GTPase mediated sign... | 0.00950 |
| Chasing | GO:0006887 | exocytosis | 0.01080 |
| Chasing | GO:0001759 | organ induction | 0.01200 |
| Chasing | GO:0035767 | endothelial cell chemotaxis | 0.01200 |
| Chasing | GO:0060390 | regulation of SMAD protein signal transd... | 0.01200 |
| Chasing | GO:0009593 | detection of chemical stimulus | 0.01250 |
| Chasing | GO:0031648 | protein destabilization | 0.01420 |
| Chasing | GO:0060602 | branch elongation of an epithelium | 0.01460 |
| Chasing | GO:0048701 | embryonic cranial skeleton morphogenesis | 0.01610 |
| Chasing | GO:0051150 | regulation of smooth muscle cell differe... | 0.01690 |
| Chasing | GO:0021516 | dorsal spinal cord development | 0.01700 |
| Chasing | GO:0048041 | focal adhesion assembly | 0.01700 |
| Chasing | GO:1903146 | regulation of autophagy of mitochondrion | 0.01700 |
| Chasing | GO:0060271 | cilium assembly | 0.01740 |
| Chasing | GO:0019318 | hexose metabolic process | 0.01750 |
| Chasing | GO:0045601 | regulation of endothelial cell different... | 0.01750 |

| | | | |
|---|---|---|---|
| Chasing | GO:0050974 | detection of mechanical stimulus involve... | 0.01750 |
| Chasing | GO:0043547 | positive regulation of GTPase activity | 0.01940 |
| Chasing | GO:0050919 | negative chemotaxis | 0.02280 |
| Chasing | GO:0008219 | cell death | 0.02300 |
| Chasing | GO:0010761 | fibroblast migration | 0.02320 |
| Chasing | GO:1901343 | negative regulation of vasculature devel... | 0.02330 |
| Chasing | GO:0008406 | gonad development | 0.02350 |
| Chasing | GO:0030902 | hindbrain development | 0.02370 |
| Chasing | GO:1900026 | positive regulation of substrate adhesio... | 0.02420 |
| Chasing | GO:0042147 | retrograde transport, endosome to Golgi | 0.02450 |
| Chasing | GO:0061098 | positive regulation of protein tyrosine ... | 0.02540 |
| Chasing | GO:0007044 | cell-substrate junction assembly | 0.03000 |
| Chasing | GO:0055074 | calcium ion homeostasis | 0.03020 |
| Chasing | GO:0030219 | megakaryocyte differentiation | 0.03200 |
| Chasing | GO:0034508 | centromere complex assembly | 0.03200 |
| Chasing | GO:0060441 | epithelial tube branching involved in lu... | 0.03200 |
| Chasing | GO:0060740 | prostate gland epithelium morphogenesis | 0.03200 |
| Chasing | GO:2001240 | negative regulation of extrinsic apoptot... | 0.03200 |
| Chasing | GO:0001704 | formation of primary germ layer | 0.03500 |
| Chasing | GO:1901184 | regulation of ERBB signaling pathway | 0.03530 |
| Chasing | GO:0009799 | specification of symmetry | 0.03550 |
| Chasing | GO:0040011 | locomotion | 0.03610 |
| Chasing | GO:0000186 | activation of MAPKK activity | 0.03630 |
| Chasing | GO:0032467 | positive regulation of cytokinesis | 0.03640 |
| Chasing | GO:0007165 | signal transduction | 0.03660 |
| Chasing | GO:0001676 | long-chain fatty acid metabolic process | 0.03720 |
| Chasing | GO:0007411 | axon guidance | 0.03740 |
| Chasing | GO:1903169 | regulation of calcium ion transmembrane ... | 0.03790 |
| Chasing | GO:0031076 | embryonic camera-type eye development | 0.03800 |

| | | | |
|---|---|---|---|
| Chasing | GO:0001569 | branching involved in blood vessel morph... | 0.04110 |
| Chasing | GO:0071805 | potassium ion transmembrane transport | 0.04120 |
| Chasing | GO:0048008 | platelet-derived growth factor receptor ... | 0.04590 |
| Chasing | GO:0043277 | apoptotic cell clearance | 0.04610 |
| Chasing | GO:0021904 | dorsal/ventral neural tube patterning | 0.04640 |
| Chasing | GO:1903672 | positive regulation of sprouting angioge... | 0.04640 |
| Chasing | GO:0003337 | mesenchymal to epithelial transition inv... | 0.04660 |
| Chasing | GO:0007600 | sensory perception | 0.04660 |
| Chasing | GO:0043923 | positive regulation by host of viral tra... | 0.04660 |
| Chasing | GO:0048012 | hepatocyte growth factor receptor signal... | 0.04660 |
| Chasing | GO:0048311 | mitochondrion distribution | 0.04660 |
| Chasing | GO:0051151 | negative regulation of smooth muscle cel... | 0.04660 |
| Chasing | GO:0070633 | transepithelial transport | 0.04660 |
| Chasing | GO:1905606 | regulation of presynapse assembly | 0.04660 |
| Dog Aggression | GO:0043393 | regulation of protein binding | 0.00005 |
| Dog Aggression | GO:0003157 | endocardium development | 0.00040 |
| Dog Aggression | GO:0001541 | ovarian follicle development | 0.00099 |
| Dog Aggression | GO:0030324 | lung development | 0.00195 |
| Dog Aggression | GO:0045737 | positive regulation of cyclin-dependent ... | 0.00215 |
| Dog Aggression | GO:0006487 | protein N-linked glycosylation | 0.00271 |
| Dog Aggression | GO:0090049 | regulation of cell migration involved in... | 0.00322 |
| Dog Aggression | GO:0000289 | nuclear-transcribed mRNA poly(A) tail sh... | 0.00434 |
| Dog Aggression | GO:0045880 | positive regulation of smoothened signal... | 0.00466 |
| Dog Aggression | GO:0007030 | Golgi organization | 0.00549 |
| Dog Aggression | GO:0046596 | regulation of viral entry into host cell | 0.00565 |
| Dog Aggression | GO:0051383 | kinetochore organization | 0.00565 |
| Dog Aggression | GO:0060628 | regulation of ER to Golgi vesicle-mediat... | 0.00565 |
| Dog Aggression | GO:1902117 | positive regulation of organelle assembl... | 0.00626 |
| Dog Aggression | GO:0006890 | retrograde vesicle-mediated transport, G... | 0.00747 |

| | | | |
|---|---|---|---|
| Dog Aggression | GO:0060045 | positive regulation of cardiac muscle ce... | 0.00892 |
| Dog Aggression | GO:0051057 | positive regulation of small GTPase medi... | 0.00938 |
| Dog Aggression | GO:0010799 | regulation of peptidyl-threonine phospho... | 0.00942 |
| Dog Aggression | GO:0032774 | RNA biosynthetic process | 0.00953 |
| Dog Aggression | GO:0060412 | ventricular septum morphogenesis | 0.00986 |
| Dog Aggression | GO:1903902 | positive regulation of viral life cycle | 0.01079 |
| Dog Aggression | GO:0032925 | regulation of activin receptor signaling... | 0.01089 |
| Dog Aggression | GO:1903671 | negative regulation of sprouting angioge... | 0.01089 |
| Dog Aggression | GO:0045666 | positive regulation of neuron differenti... | 0.01093 |
| Dog Aggression | GO:0050680 | negative regulation of epithelial cell p... | 0.01232 |
| Dog Aggression | GO:0003015 | heart process | 0.01377 |
| Dog Aggression | GO:0034333 | adherens junction assembly | 0.01383 |
| Dog Aggression | GO:0035195 | gene silencing by miRNA | 0.01388 |
| Dog Aggression | GO:0034067 | protein localization to Golgi apparatus | 0.01554 |
| Dog Aggression | GO:0045724 | positive regulation of cilium assembly | 0.01554 |
| Dog Aggression | GO:0000186 | activation of MAPKK activity | 0.01786 |
| Dog Aggression | GO:0043537 | negative regulation of blood vessel endo... | 0.01821 |
| Dog Aggression | GO:0045687 | positive regulation of glial cell differ... | 0.01821 |
| Dog Aggression | GO:0055003 | cardiac myofibril assembly | 0.01821 |
| Dog Aggression | GO:0035051 | cardiocyte differentiation | 0.01860 |
| Dog Aggression | GO:0060415 | muscle tissue morphogenesis | 0.01863 |
| Dog Aggression | GO:0035909 | aorta morphogenesis | 0.02113 |
| Dog Aggression | GO:0022607 | cellular component assembly | 0.02404 |
| Dog Aggression | GO:0008089 | anterograde axonal transport | 0.02428 |
| Dog Aggression | GO:2000147 | positive regulation of cell motility | 0.02464 |
| Dog Aggression | GO:0055010 | ventricular cardiac muscle tissue morpho... | 0.02732 |
| Dog Aggression | GO:0051642 | centrosome localization | 0.02768 |
| Dog Aggression | GO:0070588 | calcium ion transmembrane transport | 0.02832 |
| Dog Aggression | GO:0021987 | cerebral cortex development | 0.02903 |

| | | | |
|---|---|---|---|
| Dog Aggression | GO:0031032 | actomyosin structure organization | 0.03062 |
| Dog Aggression | GO:1903169 | regulation of calcium ion transmembrane ... | 0.03105 |
| Dog Aggression | GO:0043039 | tRNA aminoacylation | 0.03159 |
| Dog Aggression | GO:0030433 | ubiquitin-dependent ERAD pathway | 0.03163 |
| Dog Aggression | GO:0036474 | cell death in response to hydrogen perox... | 0.03172 |
| Dog Aggression | GO:2001057 | reactive nitrogen species metabolic proc... | 0.03179 |
| Dog Aggression | GO:0008652 | cellular amino acid biosynthetic process | 0.03180 |
| Dog Aggression | GO:0031397 | negative regulation of protein ubiquitin... | 0.03439 |
| Dog Aggression | GO:0035914 | skeletal muscle cell differentiation | 0.03439 |
| Dog Aggression | GO:0031146 | SCF-dependent proteasomal ubiquitin-depe... | 0.03519 |
| Dog Aggression | GO:0045747 | positive regulation of Notch signaling p... | 0.03519 |
| Dog Aggression | GO:0070206 | protein trimerization | 0.03792 |
| Dog Aggression | GO:0002043 | blood vessel endothelial cell proliferat... | 0.03829 |
| Dog Aggression | GO:0002115 | store-operated calcium entry | 0.03829 |
| Dog Aggression | GO:0006684 | sphingomyelin metabolic process | 0.03829 |
| Dog Aggression | GO:0006883 | cellular sodium ion homeostasis | 0.03829 |
| Dog Aggression | GO:0006901 | vesicle coating | 0.03829 |
| Dog Aggression | GO:0032897 | negative regulation of viral transcripti... | 0.03829 |
| Dog Aggression | GO:0047497 | mitochondrion transport along microtubul... | 0.03829 |
| Dog Aggression | GO:0070365 | hepatocyte differentiation | 0.03829 |
| Dog Aggression | GO:0070863 | positive regulation of protein exit from... | 0.03829 |
| Dog Aggression | GO:0098719 | sodium ion import across plasma membrane | 0.03829 |
| Dog Aggression | GO:0106074 | aminoacyl-tRNA metabolism involved in tr... | 0.03829 |
| Dog Aggression | GO:2001259 | positive regulation of cation channel ac... | 0.03930 |
| Dog Aggression | GO:0042177 | negative regulation of protein catabolic... | 0.03940 |
| Dog Aggression | GO:0022602 | ovulation cycle process | 0.04364 |
| Dog Aggression | GO:0008584 | male gonad development | 0.04369 |
| Dog Aggression | GO:0072384 | organelle transport along microtubule | 0.04527 |
| Dog Aggression | GO:0001946 | lymphangiogenesis | 0.04583 |

| | | | |
|---|---|---|---|
| Dog Aggression | GO:0010832 | negative regulation of myotube different... | 0.04583 |
| Dog Aggression | GO:0032516 | positive regulation of phosphoprotein ph... | 0.04583 |
| Dog Aggression | GO:0034104 | negative regulation of tissue remodeling | 0.04583 |
| Dog Aggression | GO:0051497 | negative regulation of stress fiber asse... | 0.04583 |
| Dog Aggression | GO:0060216 | definitive hemopoiesis | 0.04583 |
| Dog Aggression | GO:0070207 | protein homotrimerization | 0.04583 |
| Dog Aggression | GO:2000811 | negative regulation of anoikis | 0.04583 |
| Dog Fear | GO:0034138 | toll-like receptor 3 signaling pathway | 0.00040 |
| Dog Fear | GO:0042531 | positive regulation of tyrosine phosphor... | 0.00290 |
| Dog Fear | GO:0016358 | dendrite development | 0.00420 |
| Dog Fear | GO:0099518 | vesicle cytoskeletal trafficking | 0.00440 |
| Dog Fear | GO:0035556 | intracellular signal transduction | 0.00640 |
| Dog Fear | GO:0045022 | early endosome to late endosome transpor... | 0.00670 |
| Dog Fear | GO:0032770 | positive regulation of monooxygenase act... | 0.00770 |
| Dog Fear | GO:0048820 | hair follicle maturation | 0.00770 |
| Dog Fear | GO:0051450 | myoblast proliferation | 0.00770 |
| Dog Fear | GO:0010324 | membrane invagination | 0.00860 |
| Dog Fear | GO:0050775 | positive regulation of dendrite morphoge... | 0.00990 |
| Dog Fear | GO:0051303 | establishment of chromosome localization | 0.00990 |
| Dog Fear | GO:2000001 | regulation of DNA damage checkpoint | 0.00990 |
| Dog Fear | GO:0051187 | cofactor catabolic process | 0.01250 |
| Dog Fear | GO:0014068 | positive regulation of phosphatidylinosi... | 0.01370 |
| Dog Fear | GO:0048870 | cell motility | 0.01390 |
| Dog Fear | GO:0031398 | positive regulation of protein ubiquitin... | 0.01510 |
| Dog Fear | GO:0035265 | organ growth | 0.01560 |
| Dog Fear | GO:0050918 | positive chemotaxis | 0.01880 |
| Dog Fear | GO:0051043 | regulation of membrane protein ectodomai... | 0.01890 |
| Dog Fear | GO:0046834 | lipid phosphorylation | 0.02040 |
| Dog Fear | GO:0034333 | adherens junction assembly | 0.02050 |

| | | | |
|---|---|---|---|
| Dog Fear | GO:0003015 | heart process | 0.02060 |
| Dog Fear | GO:0050953 | sensory perception of light stimulus | 0.02070 |
| Dog Fear | GO:0006793 | phosphorus metabolic process | 0.02130 |
| Dog Fear | GO:0070266 | necroptotic process | 0.02260 |
| Dog Fear | GO:0070534 | protein K63-linked ubiquitination | 0.02500 |
| Dog Fear | GO:0071168 | protein localization to chromatin | 0.02660 |
| Dog Fear | GO:0010508 | positive regulation of autophagy | 0.02670 |
| Dog Fear | GO:0014855 | striated muscle cell proliferation | 0.02780 |
| Dog Fear | GO:0051224 | negative regulation of protein transport | 0.02780 |
| Dog Fear | GO:1903202 | negative regulation of oxidative stress-... | 0.02790 |
| Dog Fear | GO:0090263 | positive regulation of canonical Wnt sig... | 0.02870 |
| Dog Fear | GO:0043330 | response to exogenous dsRNA | 0.03100 |
| Dog Fear | GO:2000352 | negative regulation of endothelial cell ... | 0.03100 |
| Dog Fear | GO:0006513 | protein monoubiquitination | 0.03550 |
| Dog Fear | GO:0030517 | negative regulation of axon extension | 0.03610 |
| Dog Fear | GO:0032387 | negative regulation of intracellular tra... | 0.03630 |
| Dog Fear | GO:0009056 | catabolic process | 0.03720 |
| Dog Fear | GO:0042330 | taxis | 0.03840 |
| Dog Fear | GO:0043331 | response to dsRNA | 0.03870 |
| Dog Fear | GO:0051937 | catecholamine transport | 0.03880 |
| Dog Fear | GO:1903509 | liposaccharide metabolic process | 0.03890 |
| Dog Fear | GO:0002294 | CD4-positive, alpha-beta T cell differen... | 0.03900 |
| Dog Fear | GO:0015749 | monosaccharide transmembrane transport | 0.03900 |
| Dog Fear | GO:0034204 | lipid translocation | 0.03900 |
| Dog Fear | GO:0009145 | purine nucleoside triphosphate biosynthe... | 0.03910 |
| Dog Fear | GO:0007411 | axon guidance | 0.04040 |
| Dog Fear | GO:0014059 | regulation of dopamine secretion | 0.04100 |
| Dog Fear | GO:0032008 | positive regulation of TOR signaling | 0.04100 |
| Dog Fear | GO:0016042 | lipid catabolic process | 0.04380 |

| | | | |
|---|---|---|---|
| Dog Fear | GO:0043069 | negative regulation of programmed cell d... | 0.04540 |
| Dog Fear | GO:0006352 | DNA-templated transcription, initiation | 0.04550 |
| Dog Fear | GO:1903169 | regulation of calcium ion transmembrane ... | 0.04550 |
| Dog Fear | GO:0050796 | regulation of insulin secretion | 0.04580 |
| Dog Fear | GO:0042130 | negative regulation of T cell proliferat... | 0.04600 |
| Dog Fear | GO:0060997 | dendritic spine morphogenesis | 0.04640 |
| Dog Rivalry | GO:0048477 | oogenesis | 0.00096 |
| Dog Rivalry | GO:1903046 | meiotic cell cycle process | 0.00139 |
| Dog Rivalry | GO:0042100 | B cell proliferation | 0.00207 |
| Dog Rivalry | GO:0006896 | Golgi to vacuole transport | 0.00279 |
| Dog Rivalry | GO:0001541 | ovarian follicle development | 0.00431 |
| Dog Rivalry | GO:0060669 | embryonic placenta morphogenesis | 0.00464 |
| Dog Rivalry | GO:0032092 | positive regulation of protein binding | 0.00515 |
| Dog Rivalry | GO:0051443 | positive regulation of ubiquitin-protein... | 0.00579 |
| Dog Rivalry | GO:0051293 | establishment of spindle localization | 0.00690 |
| Dog Rivalry | GO:0006471 | protein ADP-ribosylation | 0.01019 |
| Dog Rivalry | GO:1900026 | positive regulation of substrate adhesio... | 0.01198 |
| Dog Rivalry | GO:0090101 | negative regulation of transmembrane rec... | 0.01388 |
| Dog Rivalry | GO:0070373 | negative regulation of ERK1 and ERK2 cas... | 0.01443 |
| Dog Rivalry | GO:0007214 | gamma-aminobutyric acid signaling pathwa... | 0.01609 |
| Dog Rivalry | GO:0006974 | cellular response to DNA damage stimulus | 0.01773 |
| Dog Rivalry | GO:0043393 | regulation of protein binding | 0.02026 |
| Dog Rivalry | GO:0044248 | cellular catabolic process | 0.02055 |
| Dog Rivalry | GO:0031297 | replication fork processing | 0.02088 |
| Dog Rivalry | GO:1901575 | organic substance catabolic process | 0.02298 |
| Dog Rivalry | GO:0051932 | synaptic transmission, GABAergic | 0.02327 |
| Dog Rivalry | GO:0010466 | negative regulation of peptidase activit... | 0.02373 |
| Dog Rivalry | GO:0090305 | nucleic acid phosphodiester bond hydroly... | 0.02461 |
| Dog Rivalry | GO:0099536 | synaptic signaling | 0.02704 |

| | | | |
|---|---|---|---|
| Dog Rivalry | GO:1902850 | microtubule cytoskeleton organization in... | 0.02717 |
| Dog Rivalry | GO:0045580 | regulation of T cell differentiation | 0.02861 |
| Dog Rivalry | GO:0015807 | L-amino acid transport | 0.02868 |
| Dog Rivalry | GO:0009226 | nucleotide-sugar biosynthetic process | 0.02869 |
| Dog Rivalry | GO:0031573 | intra-S DNA damage checkpoint | 0.02869 |
| Dog Rivalry | GO:0050855 | regulation of B cell receptor signaling ... | 0.02869 |
| Dog Rivalry | GO:1905939 | regulation of gonad development | 0.02869 |
| Dog Rivalry | GO:0007041 | lysosomal transport | 0.02934 |
| Dog Rivalry | GO:0016339 | calcium-dependent cell-cell adhesion via... | 0.02939 |
| Dog Rivalry | GO:0044260 | cellular macromolecule metabolic process | 0.03271 |
| Dog Rivalry | GO:0030036 | actin cytoskeleton organization | 0.03279 |
| Dog Rivalry | GO:0072384 | organelle transport along microtubule | 0.03439 |
| Dog Rivalry | GO:0010633 | negative regulation of epithelial cell m... | 0.03441 |
| Dog Rivalry | GO:0006027 | glycosaminoglycan catabolic process | 0.03445 |
| Dog Rivalry | GO:0006359 | regulation of transcription by RNA polym... | 0.03445 |
| Dog Rivalry | GO:0010832 | negative regulation of myotube different... | 0.03445 |
| Dog Rivalry | GO:0032516 | positive regulation of phosphoprotein ph... | 0.03445 |
| Dog Rivalry | GO:0022900 | electron transport chain | 0.03598 |
| Dog Rivalry | GO:0051496 | positive regulation of stress fiber asse... | 0.03945 |
| Dog Rivalry | GO:0002312 | B cell activation involved in immune res... | 0.04026 |
| Dog Rivalry | GO:0007140 | male meiotic nuclear division | 0.04043 |
| Dog Rivalry | GO:0032924 | activin receptor signaling pathway | 0.04045 |
| Dog Rivalry | GO:0051303 | establishment of chromosome localization | 0.04060 |
| Dog Rivalry | GO:0032012 | regulation of ARF protein signal transdu... | 0.04061 |
| Dog Rivalry | GO:0032469 | endoplasmic reticulum calcium ion homeos... | 0.04061 |
| Dog Rivalry | GO:0044782 | cilium organization | 0.04122 |
| Dog Rivalry | GO:0034332 | adherens junction organization | 0.04293 |
| Dog Rivalry | GO:0051056 | regulation of small GTPase mediated sign... | 0.04597 |
| Dog Rivalry | GO:0009880 | embryonic pattern specification | 0.04669 |

| | | | |
|---|---|---|---|
| Dog Rivalry | GO:0043547 | positive regulation of GTPase activity | 0.04690 |
| Dog Rivalry | GO:0010862 | positive regulation of pathway-restricte... | 0.04701 |
| Dog Rivalry | GO:0044331 | cell-cell adhesion mediated by cadherin | 0.04701 |
| Dog Rivalry | GO:0007031 | peroxisome organization | 0.04715 |
| Dog Rivalry | GO:0030728 | ovulation | 0.04715 |
| Dog Rivalry | GO:0051016 | barbed-end actin filament capping | 0.04715 |
| Energy | GO:0001764 | neuron migration | 0.00100 |
| Energy | GO:0070585 | protein localization to mitochondrion | 0.00110 |
| Energy | GO:0006366 | transcription by RNA polymerase II | 0.00190 |
| Energy | GO:0006091 | generation of precursor metabolites and ... | 0.00240 |
| Energy | GO:0006622 | protein targeting to lysosome | 0.00470 |
| Energy | GO:0090140 | regulation of mitochondrial fission | 0.00470 |
| Energy | GO:0050807 | regulation of synapse organization | 0.00610 |
| Energy | GO:0051571 | positive regulation of histone H3-K4 met... | 0.00610 |
| Energy | GO:0021987 | cerebral cortex development | 0.00640 |
| Energy | GO:1905515 | non-motile cilium assembly | 0.00750 |
| Energy | GO:0062009 | secondary palate development | 0.00780 |
| Energy | GO:0006890 | retrograde vesicle-mediated transport, G... | 0.00820 |
| Energy | GO:0048193 | Golgi vesicle transport | 0.00940 |
| Energy | GO:2000114 | regulation of establishment of cell pola... | 0.00960 |
| Energy | GO:1905475 | regulation of protein localization to me... | 0.01000 |
| Energy | GO:0060074 | synapse maturation | 0.01180 |
| Energy | GO:0070593 | dendrite self-avoidance | 0.01180 |
| Energy | GO:1901016 | regulation of potassium ion transmembran... | 0.01400 |
| Energy | GO:0001702 | gastrulation with mouth forming second | 0.01410 |
| Energy | GO:0050869 | negative regulation of B cell activation | 0.01410 |
| Energy | GO:0060039 | pericardium development | 0.01410 |
| Energy | GO:0045732 | positive regulation of protein catabolic... | 0.01500 |
| Energy | GO:0042981 | regulation of apoptotic process | 0.01940 |

| | | | |
|---|---|---|---|
| Energy | GO:0007411 | axon guidance | 0.01960 |
| Energy | GO:0048710 | regulation of astrocyte differentiation | 0.01960 |
| Energy | GO:0021799 | cerebral cortex radially oriented cell m... | 0.01990 |
| Energy | GO:0007030 | Golgi organization | 0.02090 |
| Energy | GO:0021846 | cell proliferation in forebrain | 0.02270 |
| Energy | GO:0032801 | receptor catabolic process | 0.02270 |
| Energy | GO:0000413 | protein peptidyl-prolyl isomerization | 0.02400 |
| Energy | GO:0030036 | actin cytoskeleton organization | 0.02470 |
| Energy | GO:0031023 | microtubule organizing center organizati... | 0.02570 |
| Energy | GO:0033673 | negative regulation of kinase activity | 0.02590 |
| Energy | GO:0021532 | neural tube patterning | 0.02600 |
| Energy | GO:0035249 | synaptic transmission, glutamatergic | 0.02610 |
| Energy | GO:0007156 | homophilic cell adhesion via plasma memb... | 0.02620 |
| Energy | GO:0046328 | regulation of JNK cascade | 0.02630 |
| Energy | GO:0060122 | inner ear receptor cell stereocilium org... | 0.02970 |
| Energy | GO:0065009 | regulation of molecular function | 0.03190 |
| Energy | GO:0001701 | in utero embryonic development | 0.03220 |
| Energy | GO:0032210 | regulation of telomere maintenance via t... | 0.03260 |
| Energy | GO:0048017 | inositol lipid-mediated signaling | 0.03260 |
| Energy | GO:0060512 | prostate gland morphogenesis | 0.03260 |
| Energy | GO:0071219 | cellular response to molecule of bacteri... | 0.03280 |
| Energy | GO:2000060 | positive regulation of ubiquitin-depende... | 0.03300 |
| Energy | GO:0009948 | anterior/posterior axis specification | 0.03360 |
| Energy | GO:0045786 | negative regulation of cell cycle | 0.03370 |
| Energy | GO:0032743 | positive regulation of interleukin-2 pro... | 0.03780 |
| Energy | GO:0010463 | mesenchymal cell proliferation | 0.04020 |
| Energy | GO:0002755 | MyD88-dependent toll-like receptor signa... | 0.04030 |
| Energy | GO:0006582 | melanin metabolic process | 0.04030 |
| Energy | GO:0008334 | histone mRNA metabolic process | 0.04030 |

| | | | |
|---|---|---|---|
| Energy | GO:0030888 | regulation of B cell proliferation | 0.04030 |
| Energy | GO:0032438 | melanosome organization | 0.04030 |
| Energy | GO:0036092 | phosphatidylinositol-3-phosphate biosynt... | 0.04030 |
| Energy | GO:0048311 | mitochondrion distribution | 0.04030 |
| Energy | GO:0051127 | positive regulation of actin nucleation | 0.04030 |
| Energy | GO:0055026 | negative regulation of cardiac muscle ti... | 0.04030 |
| Energy | GO:0060749 | mammary gland alveolus development | 0.04030 |
| Energy | GO:0072111 | cell proliferation involved in kidney de... | 0.04030 |
| Energy | GO:0072677 | eosinophil migration | 0.04030 |
| Energy | GO:1904262 | negative regulation of TORC1 signaling | 0.04030 |
| Energy | GO:0032722 | positive regulation of chemokine product... | 0.04210 |
| Energy | GO:0032941 | secretion by tissue | 0.04210 |
| Energy | GO:0035196 | production of miRNAs involved in gene si... | 0.04210 |
| Energy | GO:0090263 | positive regulation of canonical Wnt sig... | 0.04580 |
| Energy | GO:0021549 | cerebellum development | 0.04750 |
| Energy | GO:0002377 | immunoglobulin production | 0.04780 |
| Energy | GO:0090102 | cochlea development | 0.04800 |
| Energy | GO:0006090 | pyruvate metabolic process | 0.04820 |
| Energy | GO:0022038 | corpus callosum development | 0.04820 |
| Energy | GO:0032509 | endosome transport via multivesicular bo... | 0.04820 |
| Energy | GO:0048172 | regulation of short-term neuronal synapt... | 0.04820 |
| Energy | GO:0090075 | relaxation of muscle | 0.04820 |
| Energy | GO:1901673 | regulation of mitotic spindle assembly | 0.04820 |
| Energy | GO:1903861 | positive regulation of dendrite extensio... | 0.04820 |
| Energy | GO:2000251 | positive regulation of actin cytoskeleto... | 0.04820 |
| Energy | GO:2000811 | negative regulation of anoikis | 0.04820 |
| Energy | GO:0045087 | innate immune response | 0.04990 |
| Excitability | GO:0030890 | positive regulation of B cell proliferat... | 0.00017 |
| Excitability | GO:0072422 | signal transduction involved in DNA dama... | 0.00299 |

| | | | |
|---|---|---|---|
| Excitability | GO:0051606 | detection of stimulus | 0.00362 |
| Excitability | GO:0018105 | peptidyl-serine phosphorylation | 0.00364 |
| Excitability | GO:0002755 | MyD88-dependent toll-like receptor signa... | 0.00417 |
| Excitability | GO:0051291 | protein heterooligomerization | 0.00600 |
| Excitability | GO:0006644 | phospholipid metabolic process | 0.00632 |
| Excitability | GO:0016447 | somatic recombination of immunoglobulin ... | 0.00691 |
| Excitability | GO:0002385 | mucosal immune response | 0.00726 |
| Excitability | GO:0006336 | DNA replication-independent nucleosome a... | 0.00726 |
| Excitability | GO:0010613 | positive regulation of cardiac muscle hy... | 0.00726 |
| Excitability | GO:0007010 | cytoskeleton organization | 0.00877 |
| Excitability | GO:0051092 | positive regulation of NF-kappaB transcr... | 0.00910 |
| Excitability | GO:0051570 | regulation of histone H3-K9 methylation | 0.00920 |
| Excitability | GO:0003009 | skeletal muscle contraction | 0.01391 |
| Excitability | GO:0008045 | motor neuron axon guidance | 0.01391 |
| Excitability | GO:0019731 | antibacterial humoral response | 0.01391 |
| Excitability | GO:0045923 | positive regulation of fatty acid metabo... | 0.01391 |
| Excitability | GO:0006974 | cellular response to DNA damage stimulus | 0.01493 |
| Excitability | GO:0021517 | ventral spinal cord development | 0.01649 |
| Excitability | GO:0050869 | negative regulation of B cell activation | 0.01668 |
| Excitability | GO:0090263 | positive regulation of canonical Wnt sig... | 0.01751 |
| Excitability | GO:0006357 | regulation of transcription by RNA polym... | 0.01863 |
| Excitability | GO:0008104 | protein localization | 0.01964 |
| Excitability | GO:0046326 | positive regulation of glucose import | 0.01965 |
| Excitability | GO:0034142 | toll-like receptor 4 signaling pathway | 0.02248 |
| Excitability | GO:0031572 | G2 DNA damage checkpoint | 0.02249 |
| Excitability | GO:1902750 | negative regulation of cell cycle G2/M p... | 0.02254 |
| Excitability | GO:0010171 | body morphogenesis | 0.02255 |
| Excitability | GO:0043410 | positive regulation of MAPK cascade | 0.02631 |
| Excitability | GO:0071260 | cellular response to mechanical stimulus | 0.02658 |

| | | | |
|---|---|---|---|
| Excitability | GO:0050922 | negative regulation of chemotaxis | 0.02910 |
| Excitability | GO:0006767 | water-soluble vitamin metabolic process | 0.03066 |
| Excitability | GO:0042149 | cellular response to glucose starvation | 0.03066 |
| Excitability | GO:0009411 | response to UV | 0.03236 |
| Excitability | GO:0043331 | response to dsRNA | 0.03464 |
| Excitability | GO:0009649 | entrainment of circadian clock | 0.03472 |
| Excitability | GO:0060512 | prostate gland morphogenesis | 0.03475 |
| Excitability | GO:0099536 | synaptic signaling | 0.03479 |
| Excitability | GO:0045321 | leukocyte activation | 0.03482 |
| Excitability | GO:0016572 | histone phosphorylation | 0.03487 |
| Excitability | GO:0050663 | cytokine secretion | 0.03660 |
| Excitability | GO:0046839 | phospholipid dephosphorylation | 0.03689 |
| Excitability | GO:0043123 | positive regulation of I-kappaB kinase/N... | 0.04344 |
| Excitability | GO:0030888 | regulation of B cell proliferation | 0.04393 |
| Excitability | GO:0045332 | phospholipid translocation | 0.04414 |
| Excitability | GO:0007165 | signal transduction | 0.04499 |
| Excitability | GO:0000921 | septin ring assembly | 0.04513 |
| Excitability | GO:0007252 | I-kappaB phosphorylation | 0.04513 |
| Excitability | GO:0007263 | nitric oxide mediated signal transductio... | 0.04513 |
| Excitability | GO:0010447 | response to acidic pH | 0.04513 |
| Excitability | GO:0030033 | microvillus assembly | 0.04513 |
| Excitability | GO:0030150 | protein import into mitochondrial matrix | 0.04513 |
| Excitability | GO:0032438 | melanosome organization | 0.04513 |
| Excitability | GO:0035994 | response to muscle stretch | 0.04513 |
| Excitability | GO:0036092 | phosphatidylinositol-3-phosphate biosynt... | 0.04513 |
| Excitability | GO:0042559 | pteridine-containing compound biosynthet... | 0.04513 |
| Excitability | GO:0060218 | hematopoietic stem cell differentiation | 0.04513 |
| Excitability | GO:0061081 | positive regulation of myeloid leukocyte... | 0.04513 |
| Excitability | GO:0098719 | sodium ion import across plasma membrane | 0.04513 |

| | | | |
|---|---|---|---|
| Excitability | GO:1903205 | regulation of hydrogen peroxide-induced ... | 0.04513 |
| Excitability | GO:1904262 | negative regulation of TORC1 signaling | 0.04513 |
| Excitability | GO:0050770 | regulation of axonogenesis | 0.04850 |
| Excitability | GO:0051453 | regulation of intracellular pH | 0.04904 |
| Excitability | GO:0032722 | positive regulation of chemokine product... | 0.04919 |
| Excitability | GO:0042177 | negative regulation of protein catabolic... | 0.04935 |
| Nonsocial Fear | GO:0044260 | cellular macromolecule metabolic process | 0.00053 |
| Nonsocial Fear | GO:0042307 | positive regulation of protein import in... | 0.00296 |
| Nonsocial Fear | GO:0031290 | retinal ganglion cell axon guidance | 0.00344 |
| Nonsocial Fear | GO:0035235 | ionotropic glutamate receptor signaling ... | 0.00512 |
| Nonsocial Fear | GO:0034220 | ion transmembrane transport | 0.00523 |
| Nonsocial Fear | GO:0033522 | histone H2A ubiquitination | 0.00713 |
| Nonsocial Fear | GO:0035249 | synaptic transmission, glutamatergic | 0.00961 |
| Nonsocial Fear | GO:0010575 | positive regulation of vascular endothel... | 0.00976 |
| Nonsocial Fear | GO:0071453 | cellular response to oxygen levels | 0.00989 |
| Nonsocial Fear | GO:0110020 | regulation of actomyosin structure organ... | 0.01302 |
| Nonsocial Fear | GO:0060047 | heart contraction | 0.01639 |
| Nonsocial Fear | GO:0010658 | striated muscle cell apoptotic process | 0.02029 |
| Nonsocial Fear | GO:0032438 | melanosome organization | 0.02029 |
| Nonsocial Fear | GO:0034143 | regulation of toll-like receptor 4 signa... | 0.02029 |
| Nonsocial Fear | GO:0043248 | proteasome assembly | 0.02029 |
| Nonsocial Fear | GO:0106074 | aminoacyl-tRNA metabolism involved in tr... | 0.02029 |
| Nonsocial Fear | GO:0007600 | sensory perception | 0.02059 |
| Nonsocial Fear | GO:0042330 | taxis | 0.02239 |
| Nonsocial Fear | GO:0032608 | interferon-beta production | 0.02245 |
| Nonsocial Fear | GO:0015749 | monosaccharide transmembrane transport | 0.02252 |
| Nonsocial Fear | GO:0065009 | regulation of molecular function | 0.02268 |
| Nonsocial Fear | GO:0044419 | interspecies interaction between organis... | 0.02282 |
| Nonsocial Fear | GO:0006027 | glycosaminoglycan catabolic process | 0.02443 |

| | | | |
|---|---|---|---|
| Nonsocial Fear | GO:0048820 | hair follicle maturation | 0.02443 |
| Nonsocial Fear | GO:0070207 | protein homotrimerization | 0.02443 |
| Nonsocial Fear | GO:0002230 | positive regulation of defense response ... | 0.02888 |
| Nonsocial Fear | GO:0035025 | positive regulation of Rho protein signa... | 0.02888 |
| Nonsocial Fear | GO:0051482 | positive regulation of cytosolic calcium... | 0.02888 |
| Nonsocial Fear | GO:0051570 | regulation of histone H3-K9 methylation | 0.03364 |
| Nonsocial Fear | GO:2000136 | regulation of cell proliferation involve... | 0.03364 |
| Nonsocial Fear | GO:0046854 | phosphatidylinositol phosphorylation | 0.04072 |
| Nonsocial Fear | GO:0043547 | positive regulation of GTPase activity | 0.04170 |
| Nonsocial Fear | GO:0008045 | motor neuron axon guidance | 0.04397 |
| Nonsocial Fear | GO:0048011 | neurotrophin TRK receptor signaling path... | 0.04397 |
| Nonsocial Fear | GO:0051043 | regulation of membrane protein ectodomai... | 0.04397 |
| Nonsocial Fear | GO:0060421 | positive regulation of heart growth | 0.04454 |
| Nonsocial Fear | GO:0033048 | negative regulation of mitotic sister ch... | 0.04458 |
| Nonsocial Fear | GO:0086001 | cardiac muscle cell action potential | 0.04464 |
| Nonsocial Fear | GO:0007283 | spermatogenesis | 0.04659 |
| Nonsocial Fear | GO:0031330 | negative regulation of cellular cataboli... | 0.04878 |
| Nonsocial Fear | GO:0060343 | trabecula formation | 0.04951 |
| Owner Aggression | GO:0050806 | positive regulation of synaptic transmis... | 0.00062 |
| Owner Aggression | GO:0030030 | cell projection organization | 0.00231 |
| Owner Aggression | GO:0050798 | activated T cell proliferation | 0.00242 |
| Owner Aggression | GO:0042100 | B cell proliferation | 0.00244 |
| Owner Aggression | GO:0080134 | regulation of response to stress | 0.00755 |
| Owner Aggression | GO:0051293 | establishment of spindle localization | 0.00777 |
| Owner Aggression | GO:0032309 | icosanoid secretion | 0.01136 |
| Owner Aggression | GO:0015833 | peptide transport | 0.01552 |
| Owner Aggression | GO:0055123 | digestive system development | 0.01589 |
| Owner Aggression | GO:0003333 | amino acid transmembrane transport | 0.01625 |
| Owner Aggression | GO:0000165 | MAPK cascade | 0.01887 |

| | | | |
|---|---|---|---|
| Owner Aggression | GO:0007044 | cell-substrate junction assembly | 0.02038 |
| Owner Aggression | GO:0016042 | lipid catabolic process | 0.02537 |
| Owner Aggression | GO:1903046 | meiotic cell cycle process | 0.02587 |
| Owner Aggression | GO:0097194 | execution phase of apoptosis | 0.02594 |
| Owner Aggression | GO:0015918 | sterol transport | 0.02607 |
| Owner Aggression | GO:0060047 | heart contraction | 0.02619 |
| Owner Aggression | GO:0001843 | neural tube closure | 0.02698 |
| Owner Aggression | GO:0031146 | SCF-dependent proteasomal ubiquitin-depe... | 0.02731 |
| Owner Aggression | GO:0060548 | negative regulation of cell death | 0.02988 |
| Owner Aggression | GO:0010595 | positive regulation of endothelial cell ... | 0.03051 |
| Owner Aggression | GO:0072657 | protein localization to membrane | 0.03076 |
| Owner Aggression | GO:0015807 | L-amino acid transport | 0.03190 |
| Owner Aggression | GO:0050650 | chondroitin sulfate proteoglycan biosynt... | 0.03190 |
| Owner Aggression | GO:0015804 | neutral amino acid transport | 0.03401 |
| Owner Aggression | GO:0060541 | respiratory system development | 0.03760 |
| Owner Aggression | GO:0031098 | stress-activated protein kinase signalin... | 0.03763 |
| Owner Aggression | GO:0042531 | positive regulation of tyrosine phosphor... | 0.03765 |
| Owner Aggression | GO:0035456 | response to interferon-beta | 0.03826 |
| Owner Aggression | GO:0048172 | regulation of short-term neuronal synapt... | 0.03826 |
| Owner Aggression | GO:0035116 | embryonic hindlimb morphogenesis | 0.04148 |
| Owner Aggression | GO:0045070 | positive regulation of viral genome repl... | 0.04148 |
| Owner Aggression | GO:0071300 | cellular response to retinoic acid | 0.04148 |
| Owner Aggression | GO:0008361 | regulation of cell size | 0.04158 |
| Owner Aggression | GO:0032924 | activin receptor signaling pathway | 0.04489 |
| Owner Aggression | GO:0002230 | positive regulation of defense response ... | 0.04506 |
| Owner Aggression | GO:0009435 | NAD biosynthetic process | 0.04506 |
| Owner Aggression | GO:0010043 | response to zinc ion | 0.04506 |
| Owner Aggression | GO:0010613 | positive regulation of cardiac muscle hy... | 0.04506 |
| Owner Aggression | GO:0042430 | indole-containing compound metabolic pro... | 0.04506 |

| | | | |
|---|---|---|---|
| Owner Aggression | GO:0048791 | calcium ion-regulated exocytosis of neur... | 0.04506 |
| Owner Aggression | GO:0072520 | seminiferous tubule development | 0.04506 |
| Owner Aggression | GO:2001032 | regulation of double-strand break repair... | 0.04506 |
| Owner Aggression | GO:0046395 | carboxylic acid catabolic process | 0.04518 |
| Owner Aggression | GO:0048701 | embryonic cranial skeleton morphogenesis | 0.04550 |
| Owner Aggression | GO:0030010 | establishment of cell polarity | 0.04860 |
| Owner Aggression | GO:2000649 | regulation of sodium ion transmembrane t... | 0.04970 |
| Owner Aggression | GO:0007098 | centrosome cycle | 0.04993 |
| Separation Problems | GO:0032288 | myelin assembly | 0.00024 |
| Separation Problems | GO:0062009 | secondary palate development | 0.00051 |
| Separation Problems | GO:0090162 | establishment of epithelial cell polarit... | 0.00069 |
| Separation Problems | GO:0034329 | cell junction assembly | 0.00276 |
| Separation Problems | GO:0031498 | chromatin disassembly | 0.00397 |
| Separation Problems | GO:0042249 | establishment of planar polarity of embr... | 0.00397 |
| Separation Problems | GO:1905276 | regulation of epithelial tube formation | 0.00517 |
| Separation Problems | GO:0051304 | chromosome separation | 0.00542 |
| Separation Problems | GO:0060071 | Wnt signaling pathway, planar cell polar... | 0.00812 |
| Separation Problems | GO:0030866 | cortical actin cytoskeleton organization | 0.00884 |
| Separation Problems | GO:0051099 | positive regulation of binding | 0.00985 |
| Separation Problems | GO:0001702 | gastrulation with mouth forming second | 0.01201 |
| Separation Problems | GO:0035561 | regulation of chromatin binding | 0.01201 |
| Separation Problems | GO:0046834 | lipid phosphorylation | 0.01298 |
| Separation Problems | GO:0051960 | regulation of nervous system development | 0.01303 |
| Separation Problems | GO:0042176 | regulation of protein catabolic process | 0.01314 |
| Separation Problems | GO:0043537 | negative regulation of blood vessel endo... | 0.01673 |
| Separation Problems | GO:0010977 | negative regulation of neuron projection... | 0.01959 |
| Separation Problems | GO:0060255 | regulation of macromolecule metabolic pr... | 0.02469 |
| Separation Problems | GO:0032467 | positive regulation of cytokinesis | 0.02548 |
| Separation Problems | GO:0031032 | actomyosin structure organization | 0.02865 |

| | | | |
|---|---|---|---|
| Separation Problems | GO:0060047 | heart contraction | 0.02971 |
| Separation Problems | GO:0015749 | monosaccharide transmembrane transport | 0.03077 |
| Separation Problems | GO:0048017 | inositol lipid-mediated signaling | 0.03084 |
| Separation Problems | GO:0006892 | post-Golgi vesicle-mediated transport | 0.03592 |
| Separation Problems | GO:0040020 | regulation of meiotic nuclear division | 0.03596 |
| Separation Problems | GO:0006582 | melanin metabolic process | 0.03611 |
| Separation Problems | GO:0006744 | ubiquinone biosynthetic process | 0.03611 |
| Separation Problems | GO:0010820 | positive regulation of T cell chemotaxis | 0.03611 |
| Separation Problems | GO:0060601 | lateral sprouting from an epithelium | 0.03611 |
| Separation Problems | GO:0061162 | establishment of monopolar cell polarity | 0.03611 |
| Separation Problems | GO:0106074 | aminoacyl-tRNA metabolism involved in tr... | 0.03611 |
| Separation Problems | GO:2000052 | positive regulation of non-canonical Wnt... | 0.03611 |
| Separation Problems | GO:1903725 | regulation of phospholipid metabolic pro... | 0.03613 |
| Separation Problems | GO:0034612 | response to tumor necrosis factor | 0.03616 |
| Separation Problems | GO:0050772 | positive regulation of axonogenesis | 0.04015 |
| Separation Problems | GO:0001541 | ovarian follicle development | 0.04029 |
| Separation Problems | GO:0045089 | positive regulation of innate immune res... | 0.04049 |
| Separation Problems | GO:0071277 | cellular response to calcium ion | 0.04244 |
| Separation Problems | GO:0072132 | mesenchyme morphogenesis | 0.04292 |
| Separation Problems | GO:0051983 | regulation of chromosome segregation | 0.04314 |
| Separation Problems | GO:0002360 | T cell lineage commitment | 0.04325 |
| Separation Problems | GO:0007183 | SMAD protein complex assembly | 0.04325 |
| Separation Problems | GO:0030859 | polarized epithelial cell differentiatio... | 0.04325 |
| Separation Problems | GO:0031641 | regulation of myelination | 0.04325 |
| Separation Problems | GO:0048820 | hair follicle maturation | 0.04325 |
| Separation Problems | GO:0070102 | interleukin-6-mediated signaling pathway | 0.04325 |
| Separation Problems | GO:0007157 | heterophilic cell-cell adhesion via plas... | 0.04454 |
| Separation Problems | GO:0007163 | establishment or maintenance of cell pol... | 0.04814 |
| Separation Problems | GO:0060412 | ventricular septum morphogenesis | 0.04900 |

| | | | |
|---|---|---|---|
| Stranger Aggression | GO:0070498 | interleukin-1-mediated signaling pathway | 0.00051 |
| Stranger Aggression | GO:0014808 | release of sequestered calcium ion into ... | 0.00188 |
| Stranger Aggression | GO:0072111 | cell proliferation involved in kidney de... | 0.00188 |
| Stranger Aggression | GO:0015740 | C4-dicarboxylate transport | 0.00331 |
| Stranger Aggression | GO:0072012 | glomerulus vasculature development | 0.00422 |
| Stranger Aggression | GO:0072176 | nephric duct development | 0.00422 |
| Stranger Aggression | GO:0002089 | lens morphogenesis in camera-type eye | 0.00780 |
| Stranger Aggression | GO:0009112 | nucleobase metabolic process | 0.00958 |
| Stranger Aggression | GO:0010464 | regulation of mesenchymal cell prolifera... | 0.00959 |
| Stranger Aggression | GO:0051966 | regulation of synaptic transmission, glu... | 0.01086 |
| Stranger Aggression | GO:0043604 | amide biosynthetic process | 0.01090 |
| Stranger Aggression | GO:1903432 | regulation of TORC1 signaling | 0.01094 |
| Stranger Aggression | GO:0008104 | protein localization | 0.01242 |
| Stranger Aggression | GO:0051146 | striated muscle cell differentiation | 0.01332 |
| Stranger Aggression | GO:0008608 | attachment of spindle microtubules to ki... | 0.01470 |
| Stranger Aggression | GO:0032008 | positive regulation of TOR signaling | 0.01470 |
| Stranger Aggression | GO:0034453 | microtubule anchoring | 0.01470 |
| Stranger Aggression | GO:0021987 | cerebral cortex development | 0.01558 |
| Stranger Aggression | GO:0006749 | glutathione metabolic process | 0.01683 |
| Stranger Aggression | GO:0007080 | mitotic metaphase plate congression | 0.02156 |
| Stranger Aggression | GO:0050982 | detection of mechanical stimulus | 0.02185 |
| Stranger Aggression | GO:0071230 | cellular response to amino acid stimulus | 0.02206 |
| Stranger Aggression | GO:0010466 | negative regulation of peptidase activit... | 0.02212 |
| Stranger Aggression | GO:0007215 | glutamate receptor signaling pathway | 0.02613 |
| Stranger Aggression | GO:0008272 | sulfate transport | 0.02621 |
| Stranger Aggression | GO:2001057 | reactive nitrogen species metabolic proc... | 0.02627 |
| Stranger Aggression | GO:0009799 | specification of symmetry | 0.02628 |
| Stranger Aggression | GO:0040019 | positive regulation of embryonic develop... | 0.02695 |
| Stranger Aggression | GO:0061299 | retina vasculature morphogenesis in came... | 0.02695 |

| | | | |
|---|---|---|---|
| Stranger Aggression | GO:0070633 | transepithelial transport | 0.02695 |
| Stranger Aggression | GO:0071218 | cellular response to misfolded protein | 0.02695 |
| Stranger Aggression | GO:0099590 | neurotransmitter receptor internalizatio... | 0.02695 |
| Stranger Aggression | GO:0106074 | aminoacyl-tRNA metabolism involved in tr... | 0.02695 |
| Stranger Aggression | GO:1902285 | semaphorin-plexin signaling pathway invo... | 0.02695 |
| Stranger Aggression | GO:1905939 | regulation of gonad development | 0.02695 |
| Stranger Aggression | GO:2000052 | positive regulation of non-canonical Wnt... | 0.02695 |
| Stranger Aggression | GO:0031532 | actin cytoskeleton reorganization | 0.02768 |
| Stranger Aggression | GO:0050918 | positive chemotaxis | 0.02989 |
| Stranger Aggression | GO:0031346 | positive regulation of cell projection o... | 0.03155 |
| Stranger Aggression | GO:0006622 | protein targeting to lysosome | 0.03238 |
| Stranger Aggression | GO:0006896 | Golgi to vacuole transport | 0.03238 |
| Stranger Aggression | GO:0030859 | polarized epithelial cell differentiatio... | 0.03238 |
| Stranger Aggression | GO:0035493 | SNARE complex assembly | 0.03238 |
| Stranger Aggression | GO:0048172 | regulation of short-term neuronal synapt... | 0.03238 |
| Stranger Aggression | GO:1901881 | positive regulation of protein depolymer... | 0.03238 |
| Stranger Aggression | GO:1902894 | negative regulation of pri-miRNA transcr... | 0.03238 |
| Stranger Aggression | GO:2000095 | regulation of Wnt signaling pathway, pla... | 0.03238 |
| Stranger Aggression | GO:0045840 | positive regulation of mitotic nuclear d... | 0.03299 |
| Stranger Aggression | GO:0006914 | autophagy | 0.03781 |
| Stranger Aggression | GO:0050807 | regulation of synapse organization | 0.03791 |
| Stranger Aggression | GO:0050881 | musculoskeletal movement | 0.03803 |
| Stranger Aggression | GO:0016032 | viral process | 0.03810 |
| Stranger Aggression | GO:0007064 | mitotic sister chromatid cohesion | 0.03819 |
| Stranger Aggression | GO:0015813 | L-glutamate transmembrane transport | 0.03819 |
| Stranger Aggression | GO:0070233 | negative regulation of T cell apoptotic ... | 0.03819 |
| Stranger Aggression | GO:0017157 | regulation of exocytosis | 0.03846 |
| Stranger Aggression | GO:0043547 | positive regulation of GTPase activity | 0.03923 |
| Stranger Aggression | GO:0060976 | coronary vasculature development | 0.03967 |

| | | | |
|---|---|---|---|
| Stranger Aggression | GO:0048666 | neuron development | 0.04235 |
| Stranger Aggression | GO:0008333 | endosome to lysosome transport | 0.04325 |
| Stranger Aggression | GO:0007063 | regulation of sister chromatid cohesion | 0.04437 |
| Stranger Aggression | GO:0019433 | triglyceride catabolic process | 0.04437 |
| Stranger Aggression | GO:0030212 | hyaluronan metabolic process | 0.04437 |
| Stranger Aggression | GO:0032148 | activation of protein kinase B activity | 0.04437 |
| Stranger Aggression | GO:0006487 | protein N-linked glycosylation | 0.04698 |
| Stranger Aggression | GO:0032543 | mitochondrial translation | 0.04698 |
| Stranger Fear | GO:0007026 | negative regulation of microtubule depol... | 0.00047 |
| Stranger Fear | GO:0051897 | positive regulation of protein kinase B ... | 0.00245 |
| Stranger Fear | GO:1900182 | positive regulation of protein localizat... | 0.00347 |
| Stranger Fear | GO:0035556 | intracellular signal transduction | 0.00596 |
| Stranger Fear | GO:0008015 | blood circulation | 0.00613 |
| Stranger Fear | GO:0003015 | heart process | 0.00619 |
| Stranger Fear | GO:0040012 | regulation of locomotion | 0.00630 |
| Stranger Fear | GO:0007030 | Golgi organization | 0.00900 |
| Stranger Fear | GO:0044774 | mitotic DNA integrity checkpoint | 0.01126 |
| Stranger Fear | GO:0044818 | mitotic G2/M transition checkpoint | 0.01132 |
| Stranger Fear | GO:0030036 | actin cytoskeleton organization | 0.01377 |
| Stranger Fear | GO:1901990 | regulation of mitotic cell cycle phase t... | 0.01414 |
| Stranger Fear | GO:0042147 | retrograde transport, endosome to Golgi | 0.01690 |
| Stranger Fear | GO:0015807 | L-amino acid transport | 0.01761 |
| Stranger Fear | GO:0009225 | nucleotide-sugar metabolic process | 0.01769 |
| Stranger Fear | GO:0000076 | DNA replication checkpoint | 0.01782 |
| Stranger Fear | GO:0007379 | segment specification | 0.01782 |
| Stranger Fear | GO:0043968 | histone H2A acetylation | 0.01782 |
| Stranger Fear | GO:0071218 | cellular response to misfolded protein | 0.01782 |
| Stranger Fear | GO:0106074 | aminoacyl-tRNA metabolism involved in tr... | 0.01782 |
| Stranger Fear | GO:2000052 | positive regulation of non-canonical Wnt... | 0.01782 |

| | | | |
|---|---|---|---|
| Stranger Fear | GO:0035116 | embryonic hindlimb morphogenesis | 0.01863 |
| Stranger Fear | GO:0048813 | dendrite morphogenesis | 0.02028 |
| Stranger Fear | GO:1990748 | cellular detoxification | 0.02100 |
| Stranger Fear | GO:0008272 | sulfate transport | 0.02101 |
| Stranger Fear | GO:0051937 | catecholamine transport | 0.02106 |
| Stranger Fear | GO:0034219 | carbohydrate transmembrane transport | 0.02112 |
| Stranger Fear | GO:0042330 | taxis | 0.02126 |
| Stranger Fear | GO:0030859 | polarized epithelial cell differentiatio... | 0.02148 |
| Stranger Fear | GO:0031935 | regulation of chromatin silencing | 0.02148 |
| Stranger Fear | GO:2000095 | regulation of Wnt signaling pathway, pla... | 0.02148 |
| Stranger Fear | GO:0035335 | peptidyl-tyrosine dephosphorylation | 0.02195 |
| Stranger Fear | GO:0050804 | modulation of chemical synaptic transmis... | 0.02203 |
| Stranger Fear | GO:0071901 | negative regulation of protein serine/th... | 0.02248 |
| Stranger Fear | GO:0000186 | activation of MAPKK activity | 0.02927 |
| Stranger Fear | GO:0051187 | cofactor catabolic process | 0.02945 |
| Stranger Fear | GO:0001780 | neutrophil homeostasis | 0.02963 |
| Stranger Fear | GO:0009070 | serine family amino acid biosynthetic pr... | 0.02963 |
| Stranger Fear | GO:0010738 | regulation of protein kinase A signaling | 0.02963 |
| Stranger Fear | GO:0072012 | glomerulus vasculature development | 0.02963 |
| Stranger Fear | GO:0048146 | positive regulation of fibroblast prolif... | 0.03171 |
| Stranger Fear | GO:0006144 | purine nucleobase metabolic process | 0.03410 |
| Stranger Fear | GO:0019934 | cGMP-mediated signaling | 0.03410 |
| Stranger Fear | GO:0030540 | female genitalia development | 0.03410 |
| Stranger Fear | GO:0043114 | regulation of vascular permeability | 0.03410 |
| Stranger Fear | GO:0046653 | tetrahydrofolate metabolic process | 0.03410 |
| Stranger Fear | GO:0071417 | cellular response to organonitrogen comp... | 0.03543 |
| Stranger Fear | GO:0097237 | cellular response to toxic substance | 0.03771 |
| Stranger Fear | GO:0002040 | sprouting angiogenesis | 0.03881 |
| Stranger Fear | GO:0006515 | protein quality control for misfolded or... | 0.03881 |

| | | | |
|---|---|---|---|
| Stranger Fear | GO:0018230 | peptidyl-L-cysteine S-palmitoylation | 0.03881 |
| Stranger Fear | GO:0043068 | positive regulation of programmed cell d... | 0.04185 |
| Stranger Fear | GO:0016049 | cell growth | 0.04227 |
| Stranger Fear | GO:0016331 | morphogenesis of embryonic epithelium | 0.04366 |
| Stranger Fear | GO:0033363 | secretory granule organization | 0.04375 |
| Stranger Fear | GO:0050974 | detection of mechanical stimulus involve... | 0.04375 |
| Stranger Fear | GO:0051894 | positive regulation of focal adhesion as... | 0.04375 |
| Stranger Fear | GO:2000272 | negative regulation of signaling recepto... | 0.04375 |
| Stranger Fear | GO:1905515 | non-motile cilium assembly | 0.04545 |
| Stranger Fear | GO:0000578 | embryonic axis specification | 0.04891 |
| Stranger Fear | GO:0006730 | one-carbon metabolic process | 0.04891 |
| Stranger Fear | GO:0071168 | protein localization to chromatin | 0.04891 |
| Stranger Fear | GO:2000036 | regulation of stem cell population maint... | 0.04891 |
| Touch Sensitivity | GO:0007264 | small GTPase mediated signal transductio... | 0.00081 |
| Touch Sensitivity | GO:0099175 | regulation of postsynapse organization | 0.00251 |
| Touch Sensitivity | GO:0034143 | regulation of toll-like receptor 4 signa... | 0.00309 |
| Touch Sensitivity | GO:0051642 | centrosome localization | 0.00369 |
| Touch Sensitivity | GO:0006359 | regulation of transcription by RNA polym... | 0.00415 |
| Touch Sensitivity | GO:0021819 | layer formation in cerebral cortex | 0.00415 |
| Touch Sensitivity | GO:1903861 | positive regulation of dendrite extensio... | 0.00415 |
| Touch Sensitivity | GO:2001224 | positive regulation of neuron migration | 0.00415 |
| Touch Sensitivity | GO:0062009 | secondary palate development | 0.00687 |
| Touch Sensitivity | GO:0006998 | nuclear envelope organization | 0.00707 |
| Touch Sensitivity | GO:0045104 | intermediate filament cytoskeleton organ... | 0.00842 |
| Touch Sensitivity | GO:0000729 | DNA double-strand break processing | 0.00854 |
| Touch Sensitivity | GO:0090162 | establishment of epithelial cell polarit... | 0.00854 |
| Touch Sensitivity | GO:0098930 | axonal transport | 0.00914 |
| Touch Sensitivity | GO:0050767 | regulation of neurogenesis | 0.00993 |
| Touch Sensitivity | GO:0007097 | nuclear migration | 0.01043 |

| | | | |
|---|---|---|---|
| Touch Sensitivity | GO:0016242 | negative regulation of macroautophagy | 0.01043 |
| Touch Sensitivity | GO:0036342 | post-anal tail morphogenesis | 0.01043 |
| Touch Sensitivity | GO:0048339 | paraxial mesoderm development | 0.01043 |
| Touch Sensitivity | GO:0035235 | ionotropic glutamate receptor signaling ... | 0.01255 |
| Touch Sensitivity | GO:0048643 | positive regulation of skeletal muscle t... | 0.01255 |
| Touch Sensitivity | GO:0042733 | embryonic digit morphogenesis | 0.01273 |
| Touch Sensitivity | GO:0042176 | regulation of protein catabolic process | 0.01331 |
| Touch Sensitivity | GO:0070201 | regulation of establishment of protein l... | 0.01338 |
| Touch Sensitivity | GO:0090090 | negative regulation of canonical Wnt sig... | 0.01407 |
| Touch Sensitivity | GO:0000578 | embryonic axis specification | 0.01489 |
| Touch Sensitivity | GO:0034122 | negative regulation of toll-like recepto... | 0.01489 |
| Touch Sensitivity | GO:1902042 | negative regulation of extrinsic apoptot... | 0.01489 |
| Touch Sensitivity | GO:0010975 | regulation of neuron projection developm... | 0.01673 |
| Touch Sensitivity | GO:0072384 | organelle transport along microtubule | 0.01711 |
| Touch Sensitivity | GO:0071542 | dopaminergic neuron differentiation | 0.01746 |
| Touch Sensitivity | GO:0021801 | cerebral cortex radial glia guided migra... | 0.01817 |
| Touch Sensitivity | GO:0010033 | response to organic substance | 0.01895 |
| Touch Sensitivity | GO:0045806 | negative regulation of endocytosis | 0.02027 |
| Touch Sensitivity | GO:0090503 | RNA phosphodiester bond hydrolysis, exon... | 0.02027 |
| Touch Sensitivity | GO:0097352 | autophagosome maturation | 0.02027 |
| Touch Sensitivity | GO:0047496 | vesicle transport along microtubule | 0.02330 |
| Touch Sensitivity | GO:0042634 | regulation of hair cycle | 0.02403 |
| Touch Sensitivity | GO:0035023 | regulation of Rho protein signal transdu... | 0.02405 |
| Touch Sensitivity | GO:0043030 | regulation of macrophage activation | 0.02657 |
| Touch Sensitivity | GO:0007283 | spermatogenesis | 0.02746 |
| Touch Sensitivity | GO:1903046 | meiotic cell cycle process | 0.02937 |
| Touch Sensitivity | GO:0009948 | anterior/posterior axis specification | 0.03007 |
| Touch Sensitivity | GO:0030488 | tRNA methylation | 0.03007 |
| Touch Sensitivity | GO:0045197 | establishment or maintenance of epitheli... | 0.03007 |

| | | | |
|---|---|---|---|
| Touch Sensitivity | GO:0031032 | actomyosin structure organization | 0.03012 |
| Touch Sensitivity | GO:0031929 | TOR signaling | 0.03015 |
| Touch Sensitivity | GO:0033036 | macromolecule localization | 0.03020 |
| Touch Sensitivity | GO:1903018 | regulation of glycoprotein metabolic pro... | 0.03105 |
| Touch Sensitivity | GO:0070304 | positive regulation of stress-activated ... | 0.03107 |
| Touch Sensitivity | GO:0045599 | negative regulation of fat cell differen... | 0.03380 |
| Touch Sensitivity | GO:0071560 | cellular response to transforming growth... | 0.03405 |
| Touch Sensitivity | GO:0030888 | regulation of B cell proliferation | 0.03692 |
| Touch Sensitivity | GO:0006582 | melanin metabolic process | 0.03719 |
| Touch Sensitivity | GO:0010820 | positive regulation of T cell chemotaxis | 0.03719 |
| Touch Sensitivity | GO:0018216 | peptidyl-arginine methylation | 0.03719 |
| Touch Sensitivity | GO:0050650 | chondroitin sulfate proteoglycan biosynt... | 0.03719 |
| Touch Sensitivity | GO:0061162 | establishment of monopolar cell polarity | 0.03719 |
| Touch Sensitivity | GO:1902043 | positive regulation of extrinsic apoptot... | 0.03719 |
| Touch Sensitivity | GO:1905606 | regulation of presynapse assembly | 0.03719 |
| Touch Sensitivity | GO:2000178 | negative regulation of neural precursor ... | 0.03719 |
| Touch Sensitivity | GO:0034612 | response to tumor necrosis factor | 0.03747 |
| Touch Sensitivity | GO:0042177 | negative regulation of protein catabolic... | 0.03755 |
| Touch Sensitivity | GO:0071222 | cellular response to lipopolysaccharide | 0.04152 |
| Touch Sensitivity | GO:0016236 | macroautophagy | 0.04408 |
| Touch Sensitivity | GO:0021549 | cerebellum development | 0.04435 |
| Touch Sensitivity | GO:0090102 | cochlea development | 0.04436 |
| Touch Sensitivity | GO:1901215 | negative regulation of neuron death | 0.04443 |
| Touch Sensitivity | GO:0000712 | resolution of meiotic recombination inte... | 0.04453 |
| Touch Sensitivity | GO:0030859 | polarized epithelial cell differentiatio... | 0.04453 |
| Touch Sensitivity | GO:0034104 | negative regulation of tissue remodeling | 0.04453 |
| Touch Sensitivity | GO:0035024 | negative regulation of Rho protein signa... | 0.04453 |
| Touch Sensitivity | GO:0035456 | response to interferon-beta | 0.04453 |
| Touch Sensitivity | GO:0042249 | establishment of planar polarity of embr... | 0.04453 |

| | | | |
|---|---|---|---|
| Touch Sensitivity | GO:0045109 | intermediate filament organization | 0.04453 |
| Touch Sensitivity | GO:0050718 | positive regulation of interleukin-1 bet... | 0.04453 |
| Touch Sensitivity | GO:0051457 | maintenance of protein location in nucle... | 0.04453 |
| Touch Sensitivity | GO:2000134 | negative regulation of G1/S transition o... | 0.04635 |
| Touch Sensitivity | GO:0050680 | negative regulation of epithelial cell p... | 0.04773 |
| Trainability | GO:0032956 | regulation of actin cytoskeleton organiz... | 0.00130 |
| Trainability | GO:0048013 | ephrin receptor signaling pathway | 0.00130 |
| Trainability | GO:0051099 | positive regulation of binding | 0.00180 |
| Trainability | GO:0006890 | retrograde vesicle-mediated transport, G... | 0.00200 |
| Trainability | GO:0016236 | macroautophagy | 0.00650 |
| Trainability | GO:0001662 | behavioral fear response | 0.00670 |
| Trainability | GO:0048172 | regulation of short-term neuronal synapt... | 0.00670 |
| Trainability | GO:0048843 | negative regulation of axon extension in... | 0.00670 |
| Trainability | GO:0014706 | striated muscle tissue development | 0.00760 |
| Trainability | GO:0014033 | neural crest cell differentiation | 0.00780 |
| Trainability | GO:0046395 | carboxylic acid catabolic process | 0.00860 |
| Trainability | GO:0031116 | positive regulation of microtubule polym... | 0.00870 |
| Trainability | GO:0051893 | regulation of focal adhesion assembly | 0.01080 |
| Trainability | GO:0010765 | positive regulation of sodium ion transp... | 0.01360 |
| Trainability | GO:2000114 | regulation of establishment of cell pola... | 0.01360 |
| Trainability | GO:0051865 | protein autoubiquitination | 0.01390 |
| Trainability | GO:0006497 | protein lipidation | 0.01630 |
| Trainability | GO:0060536 | cartilage morphogenesis | 0.01860 |
| Trainability | GO:0051491 | positive regulation of filopodium assemb... | 0.01980 |
| Trainability | GO:1901021 | positive regulation of calcium ion trans... | 0.01980 |
| Trainability | GO:0018209 | peptidyl-serine modification | 0.02000 |
| Trainability | GO:0007420 | brain development | 0.02210 |
| Trainability | GO:0030036 | actin cytoskeleton organization | 0.02250 |
| Trainability | GO:0021915 | neural tube development | 0.02310 |

| | | | |
|---|---|---|---|
| Trainability | GO:0002062 | chondrocyte differentiation | 0.02330 |
| Trainability | GO:0043433 | negative regulation of DNA binding trans... | 0.02390 |
| Trainability | GO:0018210 | peptidyl-threonine modification | 0.02560 |
| Trainability | GO:0043547 | positive regulation of GTPase activity | 0.02640 |
| Trainability | GO:0050919 | negative chemotaxis | 0.02650 |
| Trainability | GO:0051279 | regulation of release of sequestered cal... | 0.02720 |
| Trainability | GO:0051966 | regulation of synaptic transmission, glu... | 0.02720 |
| Trainability | GO:0043537 | negative regulation of blood vessel endo... | 0.02730 |
| Trainability | GO:0043552 | positive regulation of phosphatidylinosi... | 0.02730 |
| Trainability | GO:0006888 | ER to Golgi vesicle-mediated transport | 0.02890 |
| Trainability | GO:0006406 | mRNA export from nucleus | 0.02950 |
| Trainability | GO:0055117 | regulation of cardiac muscle contraction | 0.03120 |
| Trainability | GO:0007029 | endoplasmic reticulum organization | 0.03140 |
| Trainability | GO:0032801 | receptor catabolic process | 0.03150 |
| Trainability | GO:0050850 | positive regulation of calcium-mediated ... | 0.03150 |
| Trainability | GO:0007044 | cell-substrate junction assembly | 0.03220 |
| Trainability | GO:0031023 | microtubule organizing center organizati... | 0.03250 |
| Trainability | GO:0097502 | mannosylation | 0.03300 |
| Trainability | GO:0035249 | synaptic transmission, glutamatergic | 0.03540 |
| Trainability | GO:0031122 | cytoplasmic microtubule organization | 0.03590 |
| Trainability | GO:0019228 | neuronal action potential | 0.03610 |
| Trainability | GO:0006886 | intracellular protein transport | 0.03670 |
| Trainability | GO:0002718 | regulation of cytokine production involv... | 0.03700 |
| Trainability | GO:0007030 | Golgi organization | 0.03800 |
| Trainability | GO:0055010 | ventricular cardiac muscle tissue morpho... | 0.04060 |
| Trainability | GO:0030032 | lamellipodium assembly | 0.04070 |
| Trainability | GO:0007416 | synapse assembly | 0.04080 |
| Trainability | GO:0098659 | inorganic cation import across plasma me... | 0.04080 |
| Trainability | GO:0060444 | branching involved in mammary gland duct... | 0.04100 |

| | | | |
|---|---|---|---|
| Trainability | GO:0050684 | regulation of mRNA processing | 0.04120 |
| Trainability | GO:0051129 | negative regulation of cellular componen... | 0.04120 |
| Trainability | GO:0031047 | gene silencing by RNA | 0.04130 |
| Trainability | GO:0042594 | response to starvation | 0.04130 |
| Trainability | GO:0001935 | endothelial cell proliferation | 0.04140 |
| Trainability | GO:0086019 | cell-cell signaling involved in cardiac ... | 0.04140 |
| Trainability | GO:0046164 | alcohol catabolic process | 0.04150 |
| Trainability | GO:0071216 | cellular response to biotic stimulus | 0.04160 |
| Trainability | GO:1904951 | positive regulation of establishment of ... | 0.04170 |
| Trainability | GO:0034260 | negative regulation of GTPase activity | 0.04620 |
| Trainability | GO:0000045 | autophagosome assembly | 0.04680 |
| Trainability | GO:0048705 | skeletal system morphogenesis | 0.04680 |

Table S7. Genes associated with behavioral traits at p ≤ 0.05 after Bonferroni correction, using SNPs mapped to the nearest gene within 20kb to derive gene-level p values (meta-analysis, Fisher's method).

| *Behavior* | *Ensemble ID* | *Gene Name* | *p* |
|---|---|---|---|
| Attach & Atn Seeking | ENSCAFG00000013221 | | < 0.00001 |
| Attach & Atn Seeking | ENSCAFG00000023562 | *DMD* | < 0.00001 |
| Attach & Atn Seeking | ENSCAFG00000000353 | *STXBP5* | < 0.00001 |
| Attach & Atn Seeking | ENSCAFG00000013297 | *IGF2BP2* | < 0.00001 |
| Attach & Atn Seeking | ENSCAFG00000018100 | *SCAPER* | < 0.00001 |
| Attach & Atn Seeking | ENSCAFG00000005914 | *TMTC2* | < 0.00001 |
| Attach & Atn Seeking | ENSCAFG00000000345 | *SRGAP1* | < 0.00001 |
| Attach & Atn Seeking | ENSCAFG00000028388 | *RF00322* | < 0.00001 |
| Attach & Atn Seeking | ENSCAFG00000018577 | *XIAP* | < 0.00001 |
| Attach & Atn Seeking | ENSCAFG00000006625 | *SFSWAP* | < 0.00001 |
| Attach & Atn Seeking | ENSCAFG00000000678 | *ZFPM2* | < 0.00001 |
| Attach & Atn Seeking | ENSCAFG00000000945 | *MAN1A1* | < 0.00001 |
| Attach & Atn Seeking | ENSCAFG00000007301 | *CWC27* | < 0.00001 |
| Attach & Atn Seeking | ENSCAFG00000002271 | *PHF14* | < 0.00001 |
| Attach & Atn Seeking | ENSCAFG00000000677 | *LRP12* | < 0.00001 |
| Attach & Atn Seeking | ENSCAFG00000007380 | *STAG1* | < 0.00001 |
| Attach & Atn Seeking | ENSCAFG00000027973 | *RF00100* | < 0.00001 |
| Attach & Atn Seeking | ENSCAFG00000007500 | *SBF2* | < 0.00001 |
| Attach & Atn Seeking | ENSCAFG00000003988 | *SPATA5* | < 0.00001 |
| Attach & Atn Seeking | ENSCAFG00000026031 | *RF00026* | < 0.00001 |
| Attach & Atn Seeking | ENSCAFG00000033690 | | < 0.00001 |
| Attach & Atn Seeking | ENSCAFG00000012413 | *RPS6KC1* | < 0.00001 |
| Attach & Atn Seeking | ENSCAFG00000033993 | | < 0.00001 |
| Attach & Atn Seeking | ENSCAFG00000000157 | *DCC* | < 0.00001 |
| Attach & Atn Seeking | ENSCAFG00000006743 | *PCMTD1* | < 0.00001 |

| | | | |
|---|---|---|---|
| Attach & Atn Seeking | ENSCAFG00000001859 | *TEX47* | < 0.00001 |
| Attach & Atn Seeking | ENSCAFG00000003678 | *CCNY* | < 0.00001 |
| Attach & Atn Seeking | ENSCAFG00000010384 | *DNAH7* | < 0.00001 |
| Attach & Atn Seeking | ENSCAFG00000019030 | *ABAT* | < 0.00001 |
| Attach & Atn Seeking | ENSCAFG00000000279 | *REPS1* | < 0.00001 |
| Attach & Atn Seeking | ENSCAFG00000029666 | *HNRNPLL* | < 0.00001 |
| Attach & Atn Seeking | ENSCAFG00000038165 | | < 0.00001 |
| Attach & Atn Seeking | ENSCAFG00000013823 | *SPECC1L* | < 0.00001 |
| Attach & Atn Seeking | ENSCAFG00000028628 | | < 0.00001 |
| Attach & Atn Seeking | ENSCAFG00000040856 | | < 0.00001 |
| Attach & Atn Seeking | ENSCAFG00000017716 | *EEF2K* | < 0.00001 |
| Attach & Atn Seeking | ENSCAFG00000016354 | *STIM2* | < 0.00001 |
| Attach & Atn Seeking | ENSCAFG00000027967 | *RF00009* | < 0.00001 |
| Attach & Atn Seeking | ENSCAFG00000033623 | | < 0.00001 |
| Attach & Atn Seeking | ENSCAFG00000034328 | | 0.00001 |
| Attach & Atn Seeking | ENSCAFG00000003322 | *IFRD1* | 0.00001 |
| Attach & Atn Seeking | ENSCAFG00000003570 | *FBXW2* | 0.00001 |
| Attach & Atn Seeking | ENSCAFG00000013852 | *MATN2* | 0.00001 |
| Attach & Atn Seeking | ENSCAFG00000000090 | *MC4R* | 0.00001 |
| Attach & Atn Seeking | ENSCAFG00000015859 | *JAKMIP1* | 0.00002 |
| Attach & Atn Seeking | ENSCAFG00000031499 | *GLIS3* | 0.00002 |
| Attach & Atn Seeking | ENSCAFG00000003701 | *ATG5* | 0.00002 |
| Attach & Atn Seeking | ENSCAFG00000018858 | *TARS* | 0.00003 |
| Attach & Atn Seeking | ENSCAFG00000019650 | *CAMTA1* | 0.00003 |
| Attach & Atn Seeking | ENSCAFG00000038450 | | 0.00004 |
| Attach & Atn Seeking | ENSCAFG00000001700 | *SND1* | 0.00005 |
| Attach & Atn Seeking | ENSCAFG00000013838 | *MASP1* | 0.00007 |
| Attach & Atn Seeking | ENSCAFG00000008056 | *RYR3* | 0.00011 |
| Attach & Atn Seeking | ENSCAFG00000034522 | | 0.00011 |

| | | | |
|---|---|---|---|
| Attach & Atn Seeking | ENSCAFG00000033275 | | 0.00014 |
| Attach & Atn Seeking | ENSCAFG00000005192 | *ACER3* | 0.00016 |
| Attach & Atn Seeking | ENSCAFG00000009390 | *PPP3CC* | 0.00019 |
| Attach & Atn Seeking | ENSCAFG00000007944 | *PAQR9* | 0.00021 |
| Attach & Atn Seeking | ENSCAFG00000002152 | *IL1R2* | 0.00027 |
| Attach & Atn Seeking | ENSCAFG00000000938 | *MCM9* | 0.00031 |
| Attach & Atn Seeking | ENSCAFG00000031436 | | 0.00032 |
| Attach & Atn Seeking | ENSCAFG00000009088 | *NUP93* | 0.00034 |
| Attach & Atn Seeking | ENSCAFG00000005326 | *UVRAG* | 0.00052 |
| Attach & Atn Seeking | ENSCAFG00000000361 | *TBC1D30* | 0.00055 |
| Attach & Atn Seeking | ENSCAFG00000004795 | *MTRF1* | 0.00062 |
| Attach & Atn Seeking | ENSCAFG00000009207 | *AP3B1* | 0.00069 |
| Attach & Atn Seeking | ENSCAFG00000010984 | *VTI1A* | 0.00070 |
| Attach & Atn Seeking | ENSCAFG00000032548 | *KCNH5* | 0.00097 |
| Attach & Atn Seeking | ENSCAFG00000001889 | *GABRB1* | 0.00114 |
| Attach & Atn Seeking | ENSCAFG00000030012 | | 0.00123 |
| Attach & Atn Seeking | ENSCAFG00000018564 | *GRIA3* | 0.00130 |
| Attach & Atn Seeking | ENSCAFG00000034953 | | 0.00144 |
| Attach & Atn Seeking | ENSCAFG00000037461 | | 0.00146 |
| Attach & Atn Seeking | ENSCAFG00000008525 | *CACNA1D* | 0.00152 |
| Attach & Atn Seeking | ENSCAFG00000013667 | *ARHGEF10* | 0.00164 |
| Attach & Atn Seeking | ENSCAFG00000018267 | *TOP3A* | 0.00167 |
| Attach & Atn Seeking | ENSCAFG00000000522 | *SNCAIP* | 0.00210 |
| Attach & Atn Seeking | ENSCAFG00000001852 | *ADAM22* | 0.00228 |
| Attach & Atn Seeking | ENSCAFG00000034741 | | 0.00240 |
| Attach & Atn Seeking | ENSCAFG00000034840 | | 0.00245 |
| Attach & Atn Seeking | ENSCAFG00000007963 | *BLK* | 0.00246 |
| Attach & Atn Seeking | ENSCAFG00000006266 | *ARHGAP26* | 0.00284 |
| Attach & Atn Seeking | ENSCAFG00000005054 | *ZRANB3* | 0.00293 |

| | | | |
|---|---|---|---|
| Attach & Atn Seeking | ENSCAFG00000033544 | | 0.00304 |
| Attach & Atn Seeking | ENSCAFG00000019042 | *METTL22* | 0.00352 |
| Attach & Atn Seeking | ENSCAFG00000030771 | *LCLAT1* | 0.00361 |
| Attach & Atn Seeking | ENSCAFG00000008568 | *AGPAT5* | 0.00368 |
| Attach & Atn Seeking | ENSCAFG00000011580 | *ENPEP* | 0.00392 |
| Attach & Atn Seeking | ENSCAFG00000004358 | *IQSEC1* | 0.00408 |
| Attach & Atn Seeking | ENSCAFG00000032853 | | 0.00445 |
| Attach & Atn Seeking | ENSCAFG00000003737 | *PARD3* | 0.00466 |
| Attach & Atn Seeking | ENSCAFG00000000935 | *CEP85L* | 0.00481 |
| Attach & Atn Seeking | ENSCAFG00000034873 | | 0.00742 |
| Attach & Atn Seeking | ENSCAFG00000011125 | *RAB3GAP2* | 0.00792 |
| Attach & Atn Seeking | ENSCAFG00000004182 | *ZC3H4* | 0.00818 |
| Attach & Atn Seeking | ENSCAFG00000018882 | *FAM122B* | 0.00850 |
| Attach & Atn Seeking | ENSCAFG00000005766 | *SYT1* | 0.00889 |
| Attach & Atn Seeking | ENSCAFG00000027573 | | 0.00916 |
| Attach & Atn Seeking | ENSCAFG00000017419 | *DACH2* | 0.00964 |
| Attach & Atn Seeking | ENSCAFG00000009756 | *ELOVL2* | 0.00997 |
| Attach & Atn Seeking | ENSCAFG00000004290 | *SAP130* | 0.01004 |
| Attach & Atn Seeking | ENSCAFG00000009609 | *TMEM117* | 0.01041 |
| Attach & Atn Seeking | ENSCAFG00000039830 | | 0.01045 |
| Attach & Atn Seeking | ENSCAFG00000000460 | *LAPTM4B* | 0.01251 |
| Attach & Atn Seeking | ENSCAFG00000006670 | *RAN* | 0.01251 |
| Attach & Atn Seeking | ENSCAFG00000005165 | *MGAT5B* | 0.01268 |
| Attach & Atn Seeking | ENSCAFG00000033697 | | 0.01286 |
| Attach & Atn Seeking | ENSCAFG00000005720 | *CLPB* | 0.01437 |
| Attach & Atn Seeking | ENSCAFG00000003998 | *BBS12* | 0.01455 |
| Attach & Atn Seeking | ENSCAFG00000029926 | | 0.01526 |
| Attach & Atn Seeking | ENSCAFG00000021943 | *RF00026* | 0.01616 |
| Attach & Atn Seeking | ENSCAFG00000003996 | *FGF2* | 0.01624 |

| | | | |
|---|---|---|---|
| Attach & Atn Seeking | ENSCAFG00000000828 | *RPS6KA2* | 0.01665 |
| Attach & Atn Seeking | ENSCAFG00000035717 | | 0.01676 |
| Attach & Atn Seeking | ENSCAFG00000025075 | *DEFB1* | 0.01720 |
| Attach & Atn Seeking | ENSCAFG00000029250 | *CDK13* | 0.01932 |
| Attach & Atn Seeking | ENSCAFG00000003651 | *CSMD2* | 0.01948 |
| Attach & Atn Seeking | ENSCAFG00000039575 | | 0.02044 |
| Attach & Atn Seeking | ENSCAFG00000039541 | | 0.02157 |
| Attach & Atn Seeking | ENSCAFG00000014707 | *CDC42* | 0.02213 |
| Attach & Atn Seeking | ENSCAFG00000025529 | *TPK1* | 0.02607 |
| Attach & Atn Seeking | ENSCAFG00000012059 | *CAMK2D* | 0.02619 |
| Attach & Atn Seeking | ENSCAFG00000037144 | | 0.02925 |
| Attach & Atn Seeking | ENSCAFG00000035644 | | 0.03182 |
| Attach & Atn Seeking | ENSCAFG00000007780 | *HTR1F* | 0.03274 |
| Attach & Atn Seeking | ENSCAFG00000003334 | *ABCA13* | 0.03479 |
| Attach & Atn Seeking | ENSCAFG00000004254 | *FAAH* | 0.03604 |
| Attach & Atn Seeking | ENSCAFG00000001930 | *PGM5* | 0.03707 |
| Attach & Atn Seeking | ENSCAFG00000033154 | | 0.03846 |
| Attach & Atn Seeking | ENSCAFG00000007770 | *LNPEP* | 0.03860 |
| Attach & Atn Seeking | ENSCAFG00000004406 | *FBLN2* | 0.03918 |
| Attach & Atn Seeking | ENSCAFG00000008501 | *PPEF2* | 0.03923 |
| Attach & Atn Seeking | ENSCAFG00000039072 | | 0.03936 |
| Attach & Atn Seeking | ENSCAFG00000009840 | *FAM13A* | 0.03959 |
| Attach & Atn Seeking | ENSCAFG00000035923 | | 0.04325 |
| Chasing | ENSCAFG00000003554 | *ASCC3* | < 0.00001 |
| Chasing | ENSCAFG00000008141 | *FBXW7* | < 0.00001 |
| Chasing | ENSCAFG00000005998 | *LHFPL6* | < 0.00001 |
| Chasing | ENSCAFG00000000670 | *RIMS2* | < 0.00001 |
| Chasing | ENSCAFG00000018720 | *ATG4C* | < 0.00001 |
| Chasing | ENSCAFG00000005849 | *TBC1D5* | < 0.00001 |

| | | | |
|---|---|---|---|
| Chasing | ENSCAFG00000006568 | *ARHGAP22* | < 0.00001 |
| Chasing | ENSCAFG00000014845 | *FAM193A* | < 0.00001 |
| Chasing | ENSCAFG00000008160 | | < 0.00001 |
| Chasing | ENSCAFG00000030067 | | < 0.00001 |
| Chasing | ENSCAFG00000039364 | | < 0.00001 |
| Chasing | ENSCAFG00000002155 | | < 0.00001 |
| Chasing | ENSCAFG00000030939 | | < 0.00001 |
| Chasing | ENSCAFG00000033197 | | < 0.00001 |
| Chasing | ENSCAFG00000035873 | | < 0.00001 |
| Chasing | ENSCAFG00000008886 | *FGF5* | < 0.00001 |
| Chasing | ENSCAFG00000003553 | *CADPS2* | < 0.00001 |
| Chasing | ENSCAFG00000018081 | *SPECC1* | < 0.00001 |
| Chasing | ENSCAFG00000011773 | *C6H7orf50* | < 0.00001 |
| Chasing | ENSCAFG00000010354 | *NT5C2* | < 0.00001 |
| Chasing | ENSCAFG00000032853 | | < 0.00001 |
| Chasing | ENSCAFG00000000770 | | < 0.00001 |
| Chasing | ENSCAFG00000000677 | *LRP12* | < 0.00001 |
| Chasing | ENSCAFG00000034837 | | < 0.00001 |
| Chasing | ENSCAFG00000013779 | | < 0.00001 |
| Chasing | ENSCAFG00000022461 | *RF00100* | < 0.00001 |
| Chasing | ENSCAFG00000038312 | | < 0.00001 |
| Chasing | ENSCAFG00000034827 | | < 0.00001 |
| Chasing | ENSCAFG00000003580 | *GRIK2* | < 0.00001 |
| Chasing | ENSCAFG00000002316 | *MLIP* | < 0.00001 |
| Chasing | ENSCAFG00000018181 | | < 0.00001 |
| Chasing | ENSCAFG00000002083 | | < 0.00001 |
| Chasing | ENSCAFG00000002562 | *COL19A1* | < 0.00001 |
| Chasing | ENSCAFG00000014682 | *HTT* | < 0.00001 |
| Chasing | ENSCAFG00000014403 | *SORCS2* | < 0.00001 |

| | | | |
|---|---|---|---|
| Chasing | ENSCAFG00000014975 | *EIF4G3* | < 0.00001 |
| Chasing | ENSCAFG00000004986 | *GDPD4* | 0.00001 |
| Chasing | ENSCAFG00000026776 | | 0.00001 |
| Chasing | ENSCAFG00000031643 | | 0.00002 |
| Chasing | ENSCAFG00000034549 | | 0.00002 |
| Chasing | ENSCAFG00000021196 | *RF00100* | 0.00002 |
| Chasing | ENSCAFG00000010744 | *NAV1* | 0.00002 |
| Chasing | ENSCAFG00000023072 | *POC1A* | 0.00002 |
| Chasing | ENSCAFG00000037340 | | 0.00002 |
| Chasing | ENSCAFG00000026119 | *RF00026* | 0.00003 |
| Chasing | ENSCAFG00000018570 | *SGIP1* | 0.00003 |
| Chasing | ENSCAFG00000034984 | | 0.00004 |
| Chasing | ENSCAFG00000034717 | | 0.00004 |
| Chasing | ENSCAFG00000038858 | | 0.00004 |
| Chasing | ENSCAFG00000018418 | *RHBDL3* | 0.00005 |
| Chasing | ENSCAFG00000009831 | *PRMT3* | 0.00006 |
| Chasing | ENSCAFG00000039762 | | 0.00006 |
| Chasing | ENSCAFG00000010822 | *ACTN2* | 0.00006 |
| Chasing | ENSCAFG00000006776 | *CIAO1* | 0.00008 |
| Chasing | ENSCAFG00000019237 | *ADCY9* | 0.00008 |
| Chasing | ENSCAFG00000004817 | *ELF1* | 0.00010 |
| Chasing | ENSCAFG00000015345 | *CC2D2A* | 0.00012 |
| Chasing | ENSCAFG00000035608 | | 0.00012 |
| Chasing | ENSCAFG00000035580 | | 0.00014 |
| Chasing | ENSCAFG00000010034 | *RYR2* | 0.00020 |
| Chasing | ENSCAFG00000031408 | *CFAP299* | 0.00022 |
| Chasing | ENSCAFG00000007246 | *CYP7B1* | 0.00026 |
| Chasing | ENSCAFG00000009924 | *TAOK3* | 0.00031 |
| Chasing | ENSCAFG00000011506 | *RRH* | 0.00035 |

| | | | |
|---|---|---|---|
| Chasing | ENSCAFG00000000459 | *TBC1D15* | 0.00047 |
| Chasing | ENSCAFG00000035147 | | 0.00047 |
| Chasing | ENSCAFG00000001801 | *TMC1* | 0.00063 |
| Chasing | ENSCAFG00000040770 | | 0.00065 |
| Chasing | ENSCAFG00000009828 | *SPATA19* | 0.00065 |
| Chasing | ENSCAFG00000031499 | *GLIS3* | 0.00070 |
| Chasing | ENSCAFG00000026661 | | 0.00071 |
| Chasing | ENSCAFG00000006694 | *STX2* | 0.00074 |
| Chasing | ENSCAFG00000019650 | *CAMTA1* | 0.00079 |
| Chasing | ENSCAFG00000007152 | *PTPRG* | 0.00091 |
| Chasing | ENSCAFG00000029135 | *GDNF* | 0.00096 |
| Chasing | ENSCAFG00000033124 | | 0.00102 |
| Chasing | ENSCAFG00000020373 | *USP33* | 0.00107 |
| Chasing | ENSCAFG00000038869 | | 0.00107 |
| Chasing | ENSCAFG00000034423 | | 0.00108 |
| Chasing | ENSCAFG00000015232 | *ZNF182* | 0.00111 |
| Chasing | ENSCAFG00000023764 | | 0.00114 |
| Chasing | ENSCAFG00000000353 | *STXBP5* | 0.00116 |
| Chasing | ENSCAFG00000003176 | *DPY19L2* | 0.00145 |
| Chasing | ENSCAFG00000006293 | *NR3C1* | 0.00158 |
| Chasing | ENSCAFG00000002081 | *CDC37L1* | 0.00178 |
| Chasing | ENSCAFG00000026467 | *RF00003* | 0.00209 |
| Chasing | ENSCAFG00000000011 | *NFATC1* | 0.00230 |
| Chasing | ENSCAFG00000007009 | *TMEM68* | 0.00270 |
| Chasing | ENSCAFG00000006790 | *STARD7* | 0.00282 |
| Chasing | ENSCAFG00000004406 | *FBLN2* | 0.00285 |
| Chasing | ENSCAFG00000006920 | *KCNIP3* | 0.00287 |
| Chasing | ENSCAFG00000034961 | | 0.00339 |
| Chasing | ENSCAFG00000004535 | *NR2C2* | 0.00354 |

| | | | |
|---|---|---|---|
| Chasing | ENSCAFG00000006691 | *SLC46A3* | 0.00354 |
| Chasing | ENSCAFG00000039479 | | 0.00358 |
| Chasing | ENSCAFG00000018858 | *TARS* | 0.00399 |
| Chasing | ENSCAFG00000002941 | *ME1* | 0.00400 |
| Chasing | ENSCAFG00000006266 | *ARHGAP26* | 0.00402 |
| Chasing | ENSCAFG00000012212 | *RAB4A* | 0.00485 |
| Chasing | ENSCAFG00000015800 | *MYO5A* | 0.00502 |
| Chasing | ENSCAFG00000003927 | *ELAVL4* | 0.00521 |
| Chasing | ENSCAFG00000033379 | | 0.00583 |
| Chasing | ENSCAFG00000038024 | | 0.00594 |
| Chasing | ENSCAFG00000016428 | *ARG2* | 0.00636 |
| Chasing | ENSCAFG00000038915 | | 0.00636 |
| Chasing | ENSCAFG00000002300 | *KLHL31* | 0.00637 |
| Chasing | ENSCAFG00000008185 | *ARFIP1* | 0.00690 |
| Chasing | ENSCAFG00000004735 | | 0.00696 |
| Chasing | ENSCAFG00000007657 | *ATOH8* | 0.00698 |
| Chasing | ENSCAFG00000031727 | *STC2* | 0.00822 |
| Chasing | ENSCAFG00000036119 | | 0.00854 |
| Chasing | ENSCAFG00000027959 | *RF01226* | 0.00855 |
| Chasing | ENSCAFG00000039894 | | 0.00879 |
| Chasing | ENSCAFG00000008123 | *TENM3* | 0.01080 |
| Chasing | ENSCAFG00000009984 | *TBR1* | 0.01111 |
| Chasing | ENSCAFG00000037051 | | 0.01123 |
| Chasing | ENSCAFG00000030105 | *IL33* | 0.01176 |
| Chasing | ENSCAFG00000006655 | *ADGRD1* | 0.01240 |
| Chasing | ENSCAFG00000007130 | *NSMAF* | 0.01374 |
| Chasing | ENSCAFG00000010000 | *C27H12orf40* | 0.01385 |
| Chasing | ENSCAFG00000035216 | | 0.01461 |
| Chasing | ENSCAFG00000007822 | *BDP1* | 0.01603 |

| | | | |
|---|---|---|---|
| Chasing | ENSCAFG00000032786 | | 0.01623 |
| Chasing | ENSCAFG00000027923 | *RF01169* | 0.01635 |
| Chasing | ENSCAFG00000009881 | *ALAS1* | 0.01748 |
| Chasing | ENSCAFG00000040639 | | 0.01766 |
| Chasing | ENSCAFG00000011155 | *STARD9* | 0.01921 |
| Chasing | ENSCAFG00000006604 | *VSTM4* | 0.01959 |
| Chasing | ENSCAFG00000001859 | *TEX47* | 0.02006 |
| Chasing | ENSCAFG00000039166 | | 0.02014 |
| Chasing | ENSCAFG00000003663 | *GSN* | 0.02022 |
| Chasing | ENSCAFG00000003778 | *AFG1L* | 0.02123 |
| Chasing | ENSCAFG00000002271 | *PHF14* | 0.02343 |
| Chasing | ENSCAFG00000013247 | *PRR14L* | 0.02502 |
| Chasing | ENSCAFG00000035717 | | 0.02746 |
| Chasing | ENSCAFG00000017167 | *KCNN3* | 0.02921 |
| Chasing | ENSCAFG00000003891 | *SLC26A4* | 0.03018 |
| Chasing | ENSCAFG00000036400 | | 0.03277 |
| Chasing | ENSCAFG00000010114 | *CIT* | 0.03379 |
| Chasing | ENSCAFG00000006612 | *SUCLG2* | 0.03432 |
| Chasing | ENSCAFG00000040008 | | 0.03741 |
| Chasing | ENSCAFG00000014856 | *RNF4* | 0.03780 |
| Chasing | ENSCAFG00000036840 | | 0.03804 |
| Chasing | ENSCAFG00000028798 | | 0.03909 |
| Chasing | ENSCAFG00000013067 | *PAK2* | 0.04067 |
| Chasing | ENSCAFG00000032608 | *LURAP1L* | 0.04324 |
| Chasing | ENSCAFG00000017519 | *TNMD* | 0.04621 |
| Chasing | ENSCAFG00000010735 | *GRAMD1C* | 0.04810 |
| Dog Aggression | ENSCAFG00000009924 | *TAOK3* | < 0.00001 |
| Dog Aggression | ENSCAFG00000023804 | *HERC3* | < 0.00001 |
| Dog Aggression | ENSCAFG00000009911 | *FBXW4* | < 0.00001 |

| | | | |
|---|---|---|---|
| Dog Aggression | ENSCAFG00000002155 | | < 0.00001 |
| Dog Aggression | ENSCAFG00000031429 | *GNA14* | < 0.00001 |
| Dog Aggression | ENSCAFG00000036640 | | < 0.00001 |
| Dog Aggression | ENSCAFG00000021629 | *RF00026* | < 0.00001 |
| Dog Aggression | ENSCAFG00000020112 | *ABCD3* | < 0.00001 |
| Dog Aggression | ENSCAFG00000023460 | *FBRSL1* | < 0.00001 |
| Dog Aggression | ENSCAFG00000001016 | *TRDN* | < 0.00001 |
| Dog Aggression | ENSCAFG00000018100 | *SCAPER* | < 0.00001 |
| Dog Aggression | ENSCAFG00000001010 | *PPP2CA* | < 0.00001 |
| Dog Aggression | ENSCAFG00000018864 | *GPC3* | < 0.00001 |
| Dog Aggression | ENSCAFG00000024350 | *CENPP* | < 0.00001 |
| Dog Aggression | ENSCAFG00000013004 | *CACNA1E* | < 0.00001 |
| Dog Aggression | ENSCAFG00000039364 | | < 0.00001 |
| Dog Aggression | ENSCAFG00000037915 | | < 0.00001 |
| Dog Aggression | ENSCAFG00000008431 | *C30H15orf41* | < 0.00001 |
| Dog Aggression | ENSCAFG00000018811 | | < 0.00001 |
| Dog Aggression | ENSCAFG00000018850 | *HS6ST2* | < 0.00001 |
| Dog Aggression | ENSCAFG00000005998 | *LHFPL6* | < 0.00001 |
| Dog Aggression | ENSCAFG00000004872 | *PTPN4* | 0.00001 |
| Dog Aggression | ENSCAFG00000011936 | *TRPM8* | 0.00001 |
| Dog Aggression | ENSCAFG00000015679 | *RGS7* | 0.00001 |
| Dog Aggression | ENSCAFG00000000678 | *ZFPM2* | 0.00001 |
| Dog Aggression | ENSCAFG00000039974 | | 0.00001 |
| Dog Aggression | ENSCAFG00000014222 | *IFT80* | 0.00001 |
| Dog Aggression | ENSCAFG00000022438 | *RF00026* | 0.00002 |
| Dog Aggression | ENSCAFG00000015803 | *SDCCAG8* | 0.00002 |
| Dog Aggression | ENSCAFG00000006196 | *CPNE4* | 0.00003 |
| Dog Aggression | ENSCAFG00000002364 | *ADGRL3* | 0.00003 |
| Dog Aggression | ENSCAFG00000010941 | *SLC24A4* | 0.00003 |

| | | | |
|---|---|---|---|
| Dog Aggression | ENSCAFG00000009771 | *TTC3* | 0.00004 |
| Dog Aggression | ENSCAFG00000001935 | *AKAP9* | 0.00006 |
| Dog Aggression | ENSCAFG00000013733 | *GATM* | 0.00007 |
| Dog Aggression | ENSCAFG00000031950 | *CARTPT* | 0.00008 |
| Dog Aggression | ENSCAFG00000009842 | *OPCML* | 0.00012 |
| Dog Aggression | ENSCAFG00000006766 | *DLC1* | 0.00012 |
| Dog Aggression | ENSCAFG00000036595 | | 0.00016 |
| Dog Aggression | ENSCAFG00000033134 | | 0.00024 |
| Dog Aggression | ENSCAFG00000000279 | *REPS1* | 0.00024 |
| Dog Aggression | ENSCAFG00000010772 | *GAL* | 0.00027 |
| Dog Aggression | ENSCAFG00000000557 | *CSNK1G3* | 0.00033 |
| Dog Aggression | ENSCAFG00000007822 | *BDP1* | 0.00053 |
| Dog Aggression | ENSCAFG00000018882 | *FAM122B* | 0.00055 |
| Dog Aggression | ENSCAFG00000014569 | *PDCD10* | 0.00069 |
| Dog Aggression | ENSCAFG00000014864 | *CASP12* | 0.00077 |
| Dog Aggression | ENSCAFG00000008557 | *APP* | 0.00101 |
| Dog Aggression | ENSCAFG00000001371 | *SUN2* | 0.00107 |
| Dog Aggression | ENSCAFG00000040355 | | 0.00119 |
| Dog Aggression | ENSCAFG00000015700 | *STX18* | 0.00127 |
| Dog Aggression | ENSCAFG00000034165 | | 0.00135 |
| Dog Aggression | ENSCAFG00000005423 | *STT3B* | 0.00152 |
| Dog Aggression | ENSCAFG00000030515 | *YPEL2* | 0.00178 |
| Dog Aggression | ENSCAFG00000007803 | *GALNTL6* | 0.00191 |
| Dog Aggression | ENSCAFG00000000239 | *PDE7B* | 0.00194 |
| Dog Aggression | ENSCAFG00000014845 | *FAM193A* | 0.00195 |
| Dog Aggression | ENSCAFG00000007364 | *KLKB1* | 0.00195 |
| Dog Aggression | ENSCAFG00000039354 | | 0.00227 |
| Dog Aggression | ENSCAFG00000008931 | | 0.00230 |
| Dog Aggression | ENSCAFG00000033673 | | 0.00235 |

| | | | |
|---|---|---|---|
| Dog Aggression | ENSCAFG00000034987 | | 0.00245 |
| Dog Aggression | ENSCAFG00000009831 | *PRMT3* | 0.00245 |
| Dog Aggression | ENSCAFG00000004731 | *TENM4* | 0.00281 |
| Dog Aggression | ENSCAFG00000018799 | *IGSF1* | 0.00292 |
| Dog Aggression | ENSCAFG00000007857 | *SCRG1* | 0.00311 |
| Dog Aggression | ENSCAFG00000014169 | | 0.00319 |
| Dog Aggression | ENSCAFG00000019194 | *SLC43A2* | 0.00327 |
| Dog Aggression | ENSCAFG00000002390 | *FAM178B* | 0.00328 |
| Dog Aggression | ENSCAFG00000028102 | *RF00088* | 0.00358 |
| Dog Aggression | ENSCAFG00000034449 | | 0.00376 |
| Dog Aggression | ENSCAFG00000022608 | *RF00026* | 0.00383 |
| Dog Aggression | ENSCAFG00000003792 | | 0.00389 |
| Dog Aggression | ENSCAFG00000000079 | *RELCH* | 0.00392 |
| Dog Aggression | ENSCAFG00000028659 | | 0.00407 |
| Dog Aggression | ENSCAFG00000037886 | | 0.00422 |
| Dog Aggression | ENSCAFG00000035815 | | 0.00432 |
| Dog Aggression | ENSCAFG00000005809 | *TM9SF2* | 0.00453 |
| Dog Aggression | ENSCAFG00000009327 | *ADGRG7* | 0.00528 |
| Dog Aggression | ENSCAFG00000027910 | *RF00026* | 0.00595 |
| Dog Aggression | ENSCAFG00000033300 | | 0.00673 |
| Dog Aggression | ENSCAFG00000018538 | *SERBP1* | 0.00759 |
| Dog Aggression | ENSCAFG00000033796 | | 0.00766 |
| Dog Aggression | ENSCAFG00000018841 | *MBNL3* | 0.00811 |
| Dog Aggression | ENSCAFG00000007251 | *KRT5* | 0.00815 |
| Dog Aggression | ENSCAFG00000004282 | *WDFY2* | 0.00887 |
| Dog Aggression | ENSCAFG00000006952 | *DHX37* | 0.00892 |
| Dog Aggression | ENSCAFG00000007235 | *HIPK3* | 0.00942 |
| Dog Aggression | ENSCAFG00000028870 | *RAB6A* | 0.00970 |
| Dog Aggression | ENSCAFG00000000935 | *CEP85L* | 0.01011 |

| | | | |
|---|---|---|---|
| Dog Aggression | ENSCAFG00000020342 | *IFI44* | 0.01048 |
| Dog Aggression | ENSCAFG00000030145 | *AUH* | 0.01129 |
| Dog Aggression | ENSCAFG00000015219 | *ZZEF1* | 0.01424 |
| Dog Aggression | ENSCAFG00000039775 | | 0.01424 |
| Dog Aggression | ENSCAFG00000002730 | | 0.01612 |
| Dog Aggression | ENSCAFG00000040105 | | 0.01635 |
| Dog Aggression | ENSCAFG00000011544 | *AOX4* | 0.01826 |
| Dog Aggression | ENSCAFG00000033999 | | 0.02082 |
| Dog Aggression | ENSCAFG00000023498 | *TLR8* | 0.02203 |
| Dog Aggression | ENSCAFG00000004436 | *RB1* | 0.02278 |
| Dog Aggression | ENSCAFG00000016949 | *ALOXE3* | 0.02333 |
| Dog Aggression | ENSCAFG00000005465 | *DZIP1* | 0.02370 |
| Dog Aggression | ENSCAFG00000010766 | *TESMIN* | 0.02617 |
| Dog Aggression | ENSCAFG00000020012 | *PKD1L2* | 0.02736 |
| Dog Aggression | ENSCAFG00000039115 | | 0.02872 |
| Dog Aggression | ENSCAFG00000030179 | *RFK* | 0.03420 |
| Dog Aggression | ENSCAFG00000015272 | *FGFBP1* | 0.03514 |
| Dog Aggression | ENSCAFG00000015343 | *DCAF6* | 0.03611 |
| Dog Aggression | ENSCAFG00000001355 | *KIAA2026* | 0.03769 |
| Dog Aggression | ENSCAFG00000014856 | *RNF4* | 0.04157 |
| Dog Aggression | ENSCAFG00000038556 | | 0.04206 |
| Dog Aggression | ENSCAFG00000001630 | *MYH9* | 0.04611 |
| Dog Fear | ENSCAFG00000013297 | *IGF2BP2* | < 0.00001 |
| Dog Fear | ENSCAFG00000008827 | *ALMS1* | < 0.00001 |
| Dog Fear | ENSCAFG00000007218 | | < 0.00001 |
| Dog Fear | ENSCAFG00000002667 | | < 0.00001 |
| Dog Fear | ENSCAFG00000009207 | *AP3B1* | < 0.00001 |
| Dog Fear | ENSCAFG00000011430 | *STXBP5L* | < 0.00001 |
| Dog Fear | ENSCAFG00000009242 | *FILIP1L* | < 0.00001 |

| | | | |
|---|---|---|---|
| Dog Fear | ENSCAFG00000005692 | *RPTOR* | < 0.00001 |
| Dog Fear | ENSCAFG00000002257 | *ICA1* | < 0.00001 |
| Dog Fear | ENSCAFG00000012967 | *PCYT1A* | < 0.00001 |
| Dog Fear | ENSCAFG00000029740 | *MSRB3* | < 0.00001 |
| Dog Fear | ENSCAFG00000028659 | | < 0.00001 |
| Dog Fear | ENSCAFG00000009390 | *PPP3CC* | < 0.00001 |
| Dog Fear | ENSCAFG00000000945 | *MAN1A1* | < 0.00001 |
| Dog Fear | ENSCAFG00000040856 | | < 0.00001 |
| Dog Fear | ENSCAFG00000001002 | *SMPDL3A* | < 0.00001 |
| Dog Fear | ENSCAFG00000018100 | *SCAPER* | < 0.00001 |
| Dog Fear | ENSCAFG00000005699 | *DNAJC18* | < 0.00001 |
| Dog Fear | ENSCAFG00000000334 | *ADAMTS2* | < 0.00001 |
| Dog Fear | ENSCAFG00000031408 | *CFAP299* | < 0.00001 |
| Dog Fear | ENSCAFG00000039641 | | < 0.00001 |
| Dog Fear | ENSCAFG00000006592 | *STK32A* | < 0.00001 |
| Dog Fear | ENSCAFG00000006248 | *CHL1* | < 0.00001 |
| Dog Fear | ENSCAFG00000013228 | *CNKSR2* | < 0.00001 |
| Dog Fear | ENSCAFG00000003657 | *NBAS* | < 0.00001 |
| Dog Fear | ENSCAFG00000002848 | *BCKDHB* | < 0.00001 |
| Dog Fear | ENSCAFG00000016811 | *RHBG* | < 0.00001 |
| Dog Fear | ENSCAFG00000037560 | | < 0.00001 |
| Dog Fear | ENSCAFG00000004436 | *RB1* | < 0.00001 |
| Dog Fear | ENSCAFG00000036506 | | < 0.00001 |
| Dog Fear | ENSCAFG00000036994 | | < 0.00001 |
| Dog Fear | ENSCAFG00000004586 | *RSU1* | 0.00001 |
| Dog Fear | ENSCAFG00000012344 | *FASTKD1* | 0.00001 |
| Dog Fear | ENSCAFG00000006743 | *PCMTD1* | 0.00001 |
| Dog Fear | ENSCAFG00000001879 | *NUAK1* | 0.00001 |
| Dog Fear | ENSCAFG00000003373 | *DGKI* | 0.00003 |

| | | | |
|---|---|---|---|
| Dog Fear | ENSCAFG00000021668 | *RF00003* | 0.00004 |
| Dog Fear | ENSCAFG00000035994 | | 0.00005 |
| Dog Fear | ENSCAFG00000017453 | *INTS3* | 0.00005 |
| Dog Fear | ENSCAFG00000000932 | *SLC35F1* | 0.00007 |
| Dog Fear | ENSCAFG00000001105 | *KCNQ3* | 0.00007 |
| Dog Fear | ENSCAFG00000001277 | *ANKS1A* | 0.00014 |
| Dog Fear | ENSCAFG00000010811 | *LRRC28* | 0.00015 |
| Dog Fear | ENSCAFG00000000969 | *HSF2* | 0.00017 |
| Dog Fear | ENSCAFG00000035064 | | 0.00018 |
| Dog Fear | ENSCAFG00000000345 | *SRGAP1* | 0.00020 |
| Dog Fear | ENSCAFG00000009270 | *PEBP4* | 0.00021 |
| Dog Fear | ENSCAFG00000000557 | *CSNK1G3* | 0.00022 |
| Dog Fear | ENSCAFG00000026448 | *RF00100* | 0.00039 |
| Dog Fear | ENSCAFG00000002227 | *C1GALT1* | 0.00044 |
| Dog Fear | ENSCAFG00000008279 | *PTPRJ* | 0.00044 |
| Dog Fear | ENSCAFG00000003323 | *KIDINS220* | 0.00050 |
| Dog Fear | ENSCAFG00000004799 | *GBX1* | 0.00053 |
| Dog Fear | ENSCAFG00000027110 | *RF00026* | 0.00060 |
| Dog Fear | ENSCAFG00000001312 | *CHCHD3* | 0.00066 |
| Dog Fear | ENSCAFG00000015695 | *PRKCH* | 0.00067 |
| Dog Fear | ENSCAFG00000033761 | | 0.00067 |
| Dog Fear | ENSCAFG00000017298 | *MAP2K1* | 0.00068 |
| Dog Fear | ENSCAFG00000035477 | | 0.00071 |
| Dog Fear | ENSCAFG00000003334 | *ABCA13* | 0.00077 |
| Dog Fear | ENSCAFG00000011756 | *PLXNA2* | 0.00089 |
| Dog Fear | ENSCAFG00000033988 | | 0.00108 |
| Dog Fear | ENSCAFG00000015422 | *KIAA0753* | 0.00112 |
| Dog Fear | ENSCAFG00000008739 | *BOLA3* | 0.00118 |
| Dog Fear | ENSCAFG00000030592 | *RF00026* | 0.00118 |

| | | | |
|---|---|---|---|
| Dog Fear | ENSCAFG00000033403 | | 0.00141 |
| Dog Fear | ENSCAFG00000010102 | *SEMA5A* | 0.00173 |
| Dog Fear | ENSCAFG00000015257 | *ANO2* | 0.00198 |
| Dog Fear | ENSCAFG00000032976 | | 0.00203 |
| Dog Fear | ENSCAFG00000015787 | *PPP2R5E* | 0.00224 |
| Dog Fear | ENSCAFG00000013995 | *CABIN1* | 0.00246 |
| Dog Fear | ENSCAFG00000039113 | | 0.00269 |
| Dog Fear | ENSCAFG00000015705 | *MSX1* | 0.00279 |
| Dog Fear | ENSCAFG00000008370 | *RNF13* | 0.00285 |
| Dog Fear | ENSCAFG00000007780 | *HTR1F* | 0.00300 |
| Dog Fear | ENSCAFG00000007691 | *GALNT18* | 0.00389 |
| Dog Fear | ENSCAFG00000001646 | *PRUNE2* | 0.00419 |
| Dog Fear | ENSCAFG00000014670 | | 0.00430 |
| Dog Fear | ENSCAFG00000036543 | | 0.00430 |
| Dog Fear | ENSCAFG00000010127 | *SNRPN* | 0.00461 |
| Dog Fear | ENSCAFG00000011155 | *STARD9* | 0.00473 |
| Dog Fear | ENSCAFG00000000430 | *ESR1* | 0.00478 |
| Dog Fear | ENSCAFG00000000167 | *DMXL1* | 0.00500 |
| Dog Fear | ENSCAFG00000011680 | *ATP11B* | 0.00550 |
| Dog Fear | ENSCAFG00000006264 | *PCLO* | 0.00561 |
| Dog Fear | ENSCAFG00000014302 | *ACTL6B* | 0.00584 |
| Dog Fear | ENSCAFG00000000409 | | 0.00606 |
| Dog Fear | ENSCAFG00000010536 | *CARMIL1* | 0.00623 |
| Dog Fear | ENSCAFG00000007533 | *ANKRD27* | 0.00682 |
| Dog Fear | ENSCAFG00000033910 | | 0.00686 |
| Dog Fear | ENSCAFG00000005061 | *LMO7* | 0.00719 |
| Dog Fear | ENSCAFG00000017847 | | 0.00770 |
| Dog Fear | ENSCAFG00000038217 | | 0.00780 |
| Dog Fear | ENSCAFG00000005814 | *RAB5A* | 0.00800 |

| | | | |
|---|---|---|---|
| Dog Fear | ENSCAFG00000037425 | | 0.00841 |
| Dog Fear | ENSCAFG00000008101 | *PLOD2* | 0.00861 |
| Dog Fear | ENSCAFG00000033944 | | 0.00924 |
| Dog Fear | ENSCAFG00000023549 | *C9orf3* | 0.01006 |
| Dog Fear | ENSCAFG00000001251 | | 0.01007 |
| Dog Fear | ENSCAFG00000034976 | | 0.01189 |
| Dog Fear | ENSCAFG00000040081 | | 0.01195 |
| Dog Fear | ENSCAFG00000039324 | | 0.01335 |
| Dog Fear | ENSCAFG00000004369 | *NOX4* | 0.01358 |
| Dog Fear | ENSCAFG00000017318 | *EBF1* | 0.01465 |
| Dog Fear | ENSCAFG00000028434 | *RF01210* | 0.01672 |
| Dog Fear | ENSCAFG00000033054 | | 0.01714 |
| Dog Fear | ENSCAFG00000010766 | *TESMIN* | 0.01787 |
| Dog Fear | ENSCAFG00000037826 | | 0.01831 |
| Dog Fear | ENSCAFG00000008551 | | 0.01909 |
| Dog Fear | ENSCAFG00000004250 | *RAB10* | 0.02010 |
| Dog Fear | ENSCAFG00000031708 | *ATXN10* | 0.02052 |
| Dog Fear | ENSCAFG00000015499 | *DLG5* | 0.02241 |
| Dog Fear | ENSCAFG00000035634 | | 0.02279 |
| Dog Fear | ENSCAFG00000017932 | *ARID3B* | 0.02497 |
| Dog Fear | ENSCAFG00000010745 | *IGHMBP2* | 0.02594 |
| Dog Fear | ENSCAFG00000015679 | *RGS7* | 0.02623 |
| Dog Fear | ENSCAFG00000023314 | *PRDM13* | 0.02633 |
| Dog Fear | ENSCAFG00000007525 | *SMARCA5* | 0.02636 |
| Dog Fear | ENSCAFG00000014255 | | 0.02848 |
| Dog Fear | ENSCAFG00000035176 | | 0.02858 |
| Dog Fear | ENSCAFG00000006370 | *HGF* | 0.02872 |
| Dog Fear | ENSCAFG00000027709 | *RF00001* | 0.02979 |
| Dog Fear | ENSCAFG00000035441 | | 0.03190 |

| | | | |
|---|---|---|---|
| Dog Fear | ENSCAFG00000020135 | *FNBP1L* | 0.03199 |
| Dog Fear | ENSCAFG00000012408 | *RPS6KA1* | 0.03377 |
| Dog Fear | ENSCAFG00000029744 | | 0.03387 |
| Dog Fear | ENSCAFG00000018996 | *CDH10* | 0.03466 |
| Dog Fear | ENSCAFG00000014153 | *ATP13A1* | 0.03678 |
| Dog Fear | ENSCAFG00000004568 | *CUBN* | 0.03927 |
| Dog Fear | ENSCAFG00000034017 | | 0.03982 |
| Dog Fear | ENSCAFG00000013246 | *LIPH* | 0.04020 |
| Dog Fear | ENSCAFG00000038695 | | 0.04119 |
| Dog Fear | ENSCAFG00000007235 | *HIPK3* | 0.04215 |
| Dog Fear | ENSCAFG00000002238 | *MIOS* | 0.04307 |
| Dog Fear | ENSCAFG00000000361 | *TBC1D30* | 0.04891 |
| Dog Fear | ENSCAFG00000008276 | *HLTF* | 0.04979 |
| Dog Rivalry | ENSCAFG00000014093 | | < 0.00001 |
| Dog Rivalry | ENSCAFG00000039058 | | < 0.00001 |
| Dog Rivalry | ENSCAFG00000008909 | *CNBD1* | < 0.00001 |
| Dog Rivalry | ENSCAFG00000038113 | | < 0.00001 |
| Dog Rivalry | ENSCAFG00000015903 | | < 0.00001 |
| Dog Rivalry | ENSCAFG00000020112 | *ABCD3* | < 0.00001 |
| Dog Rivalry | ENSCAFG00000037915 | | < 0.00001 |
| Dog Rivalry | ENSCAFG00000038128 | | < 0.00001 |
| Dog Rivalry | ENSCAFG00000034875 | | < 0.00001 |
| Dog Rivalry | ENSCAFG00000012545 | *CARF* | < 0.00001 |
| Dog Rivalry | ENSCAFG00000031241 | *LRRC55* | < 0.00001 |
| Dog Rivalry | ENSCAFG00000034281 | | < 0.00001 |
| Dog Rivalry | ENSCAFG00000028659 | | < 0.00001 |
| Dog Rivalry | ENSCAFG00000001630 | *MYH9* | < 0.00001 |
| Dog Rivalry | ENSCAFG00000016111 | *FOXK1* | < 0.00001 |
| Dog Rivalry | ENSCAFG00000005040 | *KLF12* | < 0.00001 |

| | | | |
|---|---|---|---|
| Dog Rivalry | ENSCAFG00000017651 | *DNM2* | < 0.00001 |
| Dog Rivalry | ENSCAFG00000003005 | *CREB5* | < 0.00001 |
| Dog Rivalry | ENSCAFG00000005809 | *TM9SF2* | < 0.00001 |
| Dog Rivalry | ENSCAFG00000002291 | *LYG1* | < 0.00001 |
| Dog Rivalry | ENSCAFG00000017996 | *ACACA* | < 0.00001 |
| Dog Rivalry | ENSCAFG00000001358 | *AGTPBP1* | < 0.00001 |
| Dog Rivalry | ENSCAFG00000019514 | *NPHP4* | 0.00001 |
| Dog Rivalry | ENSCAFG00000039648 | | 0.00001 |
| Dog Rivalry | ENSCAFG00000030938 | | 0.00001 |
| Dog Rivalry | ENSCAFG00000003467 | *FBXL4* | 0.00001 |
| Dog Rivalry | ENSCAFG00000038668 | | 0.00001 |
| Dog Rivalry | ENSCAFG00000021629 | *RF00026* | 0.00001 |
| Dog Rivalry | ENSCAFG00000040355 | | 0.00002 |
| Dog Rivalry | ENSCAFG00000040702 | | 0.00003 |
| Dog Rivalry | ENSCAFG00000017293 | *CEP128* | 0.00003 |
| Dog Rivalry | ENSCAFG00000030539 | | 0.00004 |
| Dog Rivalry | ENSCAFG00000004343 | *EFCC1* | 0.00004 |
| Dog Rivalry | ENSCAFG00000002413 | *DST* | 0.00007 |
| Dog Rivalry | ENSCAFG00000033119 | | 0.00008 |
| Dog Rivalry | ENSCAFG00000004243 | *SLC36A4* | 0.00010 |
| Dog Rivalry | ENSCAFG00000016200 | *TCF12* | 0.00011 |
| Dog Rivalry | ENSCAFG00000004855 | *EPB41L5* | 0.00011 |
| Dog Rivalry | ENSCAFG00000014222 | *IFT80* | 0.00012 |
| Dog Rivalry | ENSCAFG00000000086 | *CDH20* | 0.00015 |
| Dog Rivalry | ENSCAFG00000001646 | *PRUNE2* | 0.00017 |
| Dog Rivalry | ENSCAFG00000031628 | | 0.00026 |
| Dog Rivalry | ENSCAFG00000005549 | *UCP3* | 0.00027 |
| Dog Rivalry | ENSCAFG00000035258 | | 0.00030 |
| Dog Rivalry | ENSCAFG00000009831 | *PRMT3* | 0.00035 |

| | | | |
|---|---|---|---|
| Dog Rivalry | ENSCAFG00000019133 | *MTMR1* | 0.00036 |
| Dog Rivalry | ENSCAFG00000002271 | *PHF14* | 0.00036 |
| Dog Rivalry | ENSCAFG00000005862 | *PLCL2* | 0.00042 |
| Dog Rivalry | ENSCAFG00000012638 | *DPP3* | 0.00046 |
| Dog Rivalry | ENSCAFG00000010677 | *EYA2* | 0.00048 |
| Dog Rivalry | ENSCAFG00000033142 | | 0.00061 |
| Dog Rivalry | ENSCAFG00000036254 | | 0.00073 |
| Dog Rivalry | ENSCAFG00000010905 | *B4GALT4* | 0.00084 |
| Dog Rivalry | ENSCAFG00000007803 | *GALNTL6* | 0.00085 |
| Dog Rivalry | ENSCAFG00000020516 | *MIR137* | 0.00089 |
| Dog Rivalry | ENSCAFG00000002390 | *FAM178B* | 0.00092 |
| Dog Rivalry | ENSCAFG00000002663 | *CD109* | 0.00095 |
| Dog Rivalry | ENSCAFG00000001166 | *KHDRBS3* | 0.00099 |
| Dog Rivalry | ENSCAFG00000025956 | *RF00001* | 0.00101 |
| Dog Rivalry | ENSCAFG00000018890 | *TXNDC11* | 0.00108 |
| Dog Rivalry | ENSCAFG00000011176 | *NR5A2* | 0.00108 |
| Dog Rivalry | ENSCAFG00000009954 | | 0.00134 |
| Dog Rivalry | ENSCAFG00000033155 | | 0.00162 |
| Dog Rivalry | ENSCAFG00000036020 | | 0.00168 |
| Dog Rivalry | ENSCAFG00000035994 | | 0.00194 |
| Dog Rivalry | ENSCAFG00000020206 | *SCAI* | 0.00210 |
| Dog Rivalry | ENSCAFG00000016354 | *STIM2* | 0.00213 |
| Dog Rivalry | ENSCAFG00000034449 | | 0.00221 |
| Dog Rivalry | ENSCAFG00000039491 | | 0.00229 |
| Dog Rivalry | ENSCAFG00000012804 | *MACO1* | 0.00232 |
| Dog Rivalry | ENSCAFG00000013733 | *GATM* | 0.00360 |
| Dog Rivalry | ENSCAFG00000002420 | *MEOX2* | 0.00362 |
| Dog Rivalry | ENSCAFG00000000655 | *TMEM181* | 0.00367 |
| Dog Rivalry | ENSCAFG00000012413 | *RPS6KC1* | 0.00412 |

| | | | |
|---|---|---|---|
| Dog Rivalry | ENSCAFG00000034165 | | 0.00549 |
| Dog Rivalry | ENSCAFG00000015877 | *SH2D4B* | 0.00611 |
| Dog Rivalry | ENSCAFG00000011544 | *AOX4* | 0.00656 |
| Dog Rivalry | ENSCAFG00000024975 | *GRXCR1* | 0.00678 |
| Dog Rivalry | ENSCAFG00000040595 | | 0.00696 |
| Dog Rivalry | ENSCAFG00000005405 | *SEC23B* | 0.00703 |
| Dog Rivalry | ENSCAFG00000010224 | | 0.00727 |
| Dog Rivalry | ENSCAFG00000003420 | *VRK3* | 0.00735 |
| Dog Rivalry | ENSCAFG00000006462 | *UBXN8* | 0.00782 |
| Dog Rivalry | ENSCAFG00000015255 | *SLC35F4* | 0.00865 |
| Dog Rivalry | ENSCAFG00000009999 | *DSTYK* | 0.00896 |
| Dog Rivalry | ENSCAFG00000018011 | *LRRC75A* | 0.00916 |
| Dog Rivalry | ENSCAFG00000023912 | *LYG2* | 0.00973 |
| Dog Rivalry | ENSCAFG00000029987 | *PATE3* | 0.01049 |
| Dog Rivalry | ENSCAFG00000005873 | *RFTN1* | 0.01130 |
| Dog Rivalry | ENSCAFG00000020080 | *PTBP2* | 0.01209 |
| Dog Rivalry | ENSCAFG00000013057 | *MAP3K15* | 0.01215 |
| Dog Rivalry | ENSCAFG00000001733 | *ZNF800* | 0.01243 |
| Dog Rivalry | ENSCAFG00000033172 | | 0.01406 |
| Dog Rivalry | ENSCAFG00000000696 | *RSPO2* | 0.01425 |
| Dog Rivalry | ENSCAFG00000011983 | *HSPBAP1* | 0.01569 |
| Dog Rivalry | ENSCAFG00000003043 | *KIAA1958* | 0.01611 |
| Dog Rivalry | ENSCAFG00000024350 | *CENPP* | 0.01705 |
| Dog Rivalry | ENSCAFG00000037076 | | 0.01832 |
| Dog Rivalry | ENSCAFG00000018204 | *TRPC5* | 0.01968 |
| Dog Rivalry | ENSCAFG00000014892 | | 0.02018 |
| Dog Rivalry | ENSCAFG00000033819 | | 0.02107 |
| Dog Rivalry | ENSCAFG00000035757 | | 0.02130 |
| Dog Rivalry | ENSCAFG00000034506 | | 0.02389 |

| | | | |
|---|---|---|---|
| Dog Rivalry | ENSCAFG00000039974 | | 0.02409 |
| Dog Rivalry | ENSCAFG00000028386 | *RF00026* | 0.02502 |
| Dog Rivalry | ENSCAFG00000000719 | *NUDCD1* | 0.02702 |
| Dog Rivalry | ENSCAFG00000010942 | *CCDC91* | 0.02747 |
| Dog Rivalry | ENSCAFG00000014087 | | 0.03031 |
| Dog Rivalry | ENSCAFG00000005121 | *SFMBT2* | 0.03038 |
| Dog Rivalry | ENSCAFG00000002640 | *RIMS1* | 0.03310 |
| Dog Rivalry | ENSCAFG00000009669 | *MDM4* | 0.03521 |
| Dog Rivalry | ENSCAFG00000023463 | *ITPR1* | 0.03837 |
| Dog Rivalry | ENSCAFG00000008407 | *EIF2A* | 0.04294 |
| Dog Rivalry | ENSCAFG00000008185 | *ARFIP1* | 0.04394 |
| Dog Rivalry | ENSCAFG00000034736 | | 0.04562 |
| Dog Rivalry | ENSCAFG00000028804 | *RNF150* | 0.04953 |
| Energy | ENSCAFG00000003737 | *PARD3* | < 0.00001 |
| Energy | ENSCAFG00000001106 | *LAMA2* | < 0.00001 |
| Energy | ENSCAFG00000009699 | *CHD9* | < 0.00001 |
| Energy | ENSCAFG00000008056 | *RYR3* | < 0.00001 |
| Energy | ENSCAFG00000001700 | *SND1* | < 0.00001 |
| Energy | ENSCAFG00000007520 | *CBR4* | < 0.00001 |
| Energy | ENSCAFG00000018849 | *SNX29* | < 0.00001 |
| Energy | ENSCAFG00000023267 | *PRIM2* | < 0.00001 |
| Energy | ENSCAFG00000007301 | *CWC27* | < 0.00001 |
| Energy | ENSCAFG00000008199 | *FMN1* | < 0.00001 |
| Energy | ENSCAFG00000013221 | | < 0.00001 |
| Energy | ENSCAFG00000000557 | *CSNK1G3* | < 0.00001 |
| Energy | ENSCAFG00000025529 | *TPK1* | < 0.00001 |
| Energy | ENSCAFG00000009971 | *ARMH3* | < 0.00001 |
| Energy | ENSCAFG00000018857 | *GPC4* | < 0.00001 |
| Energy | ENSCAFG00000032608 | *LURAP1L* | < 0.00001 |

| | | | |
|---|---|---|---|
| Energy | ENSCAFG00000000430 | *ESR1* | < 0.00001 |
| Energy | ENSCAFG00000001154 | *MRTFA* | < 0.00001 |
| Energy | ENSCAFG00000038915 | | < 0.00001 |
| Energy | ENSCAFG00000032832 | | < 0.00001 |
| Energy | ENSCAFG00000006625 | *SFSWAP* | < 0.00001 |
| Energy | ENSCAFG00000004589 | | < 0.00001 |
| Energy | ENSCAFG00000008790 | *CTNNBL1* | < 0.00001 |
| Energy | ENSCAFG00000006812 | *TMEM132D* | < 0.00001 |
| Energy | ENSCAFG00000018850 | *HS6ST2* | < 0.00001 |
| Energy | ENSCAFG00000014349 | *TBC1D14* | 0.00001 |
| Energy | ENSCAFG00000018564 | *GRIA3* | 0.00001 |
| Energy | ENSCAFG00000039364 | | 0.00001 |
| Energy | ENSCAFG00000040039 | | 0.00001 |
| Energy | ENSCAFG00000035898 | | 0.00002 |
| Energy | ENSCAFG00000018952 | *ARHGEF6* | 0.00002 |
| Energy | ENSCAFG00000013228 | *CNKSR2* | 0.00002 |
| Energy | ENSCAFG00000039407 | | 0.00002 |
| Energy | ENSCAFG00000002364 | *ADGRL3* | 0.00002 |
| Energy | ENSCAFG00000008211 | *CACNA2D3* | 0.00003 |
| Energy | ENSCAFG00000030512 | | 0.00003 |
| Energy | ENSCAFG00000019051 | | 0.00004 |
| Energy | ENSCAFG00000033823 | | 0.00005 |
| Energy | ENSCAFG00000007586 | *MAST4* | 0.00006 |
| Energy | ENSCAFG00000018024 | *TRAPPC8* | 0.00006 |
| Energy | ENSCAFG00000014257 | *PPP2R2C* | 0.00007 |
| Energy | ENSCAFG00000028405 | *RF00026* | 0.00007 |
| Energy | ENSCAFG00000012360 | *ALS2* | 0.00007 |
| Energy | ENSCAFG00000014916 | *FAM227B* | 0.00007 |
| Energy | ENSCAFG00000011082 | *CAMSAP2* | 0.00008 |

| | | | |
|---|---|---|---|
| Energy | ENSCAFG00000012098 | *PGBD5* | 0.00010 |
| Energy | ENSCAFG00000018853 | *USP26* | 0.00010 |
| Energy | ENSCAFG00000033989 | | 0.00011 |
| Energy | ENSCAFG00000022240 | *RF00410* | 0.00011 |
| Energy | ENSCAFG00000031048 | *ZNF84* | 0.00015 |
| Energy | ENSCAFG00000040529 | | 0.00015 |
| Energy | ENSCAFG00000000279 | *REPS1* | 0.00015 |
| Energy | ENSCAFG00000006764 | *PARG* | 0.00016 |
| Energy | ENSCAFG00000008551 | | 0.00016 |
| Energy | ENSCAFG00000006249 | *NDFIP1* | 0.00019 |
| Energy | ENSCAFG00000028137 | *RF00100* | 0.00021 |
| Energy | ENSCAFG00000004290 | *SAP130* | 0.00022 |
| Energy | ENSCAFG00000003322 | *IFRD1* | 0.00025 |
| Energy | ENSCAFG00000039300 | | 0.00025 |
| Energy | ENSCAFG00000000011 | *NFATC1* | 0.00031 |
| Energy | ENSCAFG00000015604 | *ILDR2* | 0.00032 |
| Energy | ENSCAFG00000035434 | | 0.00033 |
| Energy | ENSCAFG00000016531 | *ADGRA3* | 0.00036 |
| Energy | ENSCAFG00000008684 | *SLC4A5* | 0.00038 |
| Energy | ENSCAFG00000003663 | *GSN* | 0.00039 |
| Energy | ENSCAFG00000036039 | | 0.00048 |
| Energy | ENSCAFG00000029157 | *NOVA1* | 0.00049 |
| Energy | ENSCAFG00000013749 | *CUX1* | 0.00053 |
| Energy | ENSCAFG00000014454 | *ATM* | 0.00055 |
| Energy | ENSCAFG00000001234 | *UHRF1BP1* | 0.00064 |
| Energy | ENSCAFG00000002640 | *RIMS1* | 0.00073 |
| Energy | ENSCAFG00000011500 | *NOL11* | 0.00079 |
| Energy | ENSCAFG00000006670 | *RAN* | 0.00080 |
| Energy | ENSCAFG00000002028 | *VLDLR* | 0.00083 |

| | | | |
|---|---|---|---|
| Energy | ENSCAFG00000008284 | *EXTL3* | 0.00088 |
| Energy | ENSCAFG00000001425 | *GKAP1* | 0.00092 |
| Energy | ENSCAFG00000007750 | *CASP3* | 0.00095 |
| Energy | ENSCAFG00000038450 | | 0.00096 |
| Energy | ENSCAFG00000036209 | | 0.00109 |
| Energy | ENSCAFG00000007949 | *LGI1* | 0.00118 |
| Energy | ENSCAFG00000018858 | *TARS* | 0.00131 |
| Energy | ENSCAFG00000033361 | | 0.00134 |
| Energy | ENSCAFG00000000327 | | 0.00178 |
| Energy | ENSCAFG00000018204 | *TRPC5* | 0.00208 |
| Energy | ENSCAFG00000002045 | | 0.00219 |
| Energy | ENSCAFG00000028957 | *GLT1D1* | 0.00238 |
| Energy | ENSCAFG00000002025 | *GCC2* | 0.00249 |
| Energy | ENSCAFG00000018128 | | 0.00250 |
| Energy | ENSCAFG00000014046 | *RASAL2* | 0.00257 |
| Energy | ENSCAFG00000012544 | *COPA* | 0.00264 |
| Energy | ENSCAFG00000009903 | *MYLIP* | 0.00289 |
| Energy | ENSCAFG00000003161 | *BBS9* | 0.00323 |
| Energy | ENSCAFG00000032743 | | 0.00347 |
| Energy | ENSCAFG00000038201 | | 0.00353 |
| Energy | ENSCAFG00000019082 | *RBFOX1* | 0.00364 |
| Energy | ENSCAFG00000036778 | | 0.00365 |
| Energy | ENSCAFG00000011167 | *PAPSS1* | 0.00405 |
| Energy | ENSCAFG00000030012 | | 0.00435 |
| Energy | ENSCAFG00000002341 | *HMGCLL1* | 0.00443 |
| Energy | ENSCAFG00000009486 | *RPGRIP1L* | 0.00450 |
| Energy | ENSCAFG00000001852 | *ADAM22* | 0.00467 |
| Energy | ENSCAFG00000017349 | *STXBP4* | 0.00487 |
| Energy | ENSCAFG00000002414 | *AGMO* | 0.00493 |

| | | | |
|---|---|---|---|
| Energy | ENSCAFG00000011698 | *TLR7* | 0.00493 |
| Energy | ENSCAFG00000018971 | | 0.00497 |
| Energy | ENSCAFG00000038471 | | 0.00518 |
| Energy | ENSCAFG00000039328 | | 0.00545 |
| Energy | ENSCAFG00000015422 | *KIAA0753* | 0.00555 |
| Energy | ENSCAFG00000039092 | | 0.00562 |
| Energy | ENSCAFG00000008293 | *FZD3* | 0.00574 |
| Energy | ENSCAFG00000008039 | *DIP2B* | 0.00609 |
| Energy | ENSCAFG00000013114 | *RGSL1* | 0.00628 |
| Energy | ENSCAFG00000010028 | *IFITM10* | 0.00683 |
| Energy | ENSCAFG00000036051 | | 0.00696 |
| Energy | ENSCAFG00000039058 | | 0.00866 |
| Energy | ENSCAFG00000038846 | | 0.00889 |
| Energy | ENSCAFG00000004716 | | 0.00903 |
| Energy | ENSCAFG00000013852 | *MATN2* | 0.01017 |
| Energy | ENSCAFG00000011514 | *GTF2I* | 0.01026 |
| Energy | ENSCAFG00000004479 | *ANO10* | 0.01067 |
| Energy | ENSCAFG00000009991 | | 0.01088 |
| Energy | ENSCAFG00000018083 | *TMEM164* | 0.01222 |
| Energy | ENSCAFG00000018996 | *CDH10* | 0.01256 |
| Energy | ENSCAFG00000036869 | | 0.01277 |
| Energy | ENSCAFG00000007907 | *LRBA* | 0.01344 |
| Energy | ENSCAFG00000015135 | *LDB2* | 0.01351 |
| Energy | ENSCAFG00000029142 | *C4H5orf51* | 0.01474 |
| Energy | ENSCAFG00000006236 | *DELE1* | 0.01571 |
| Energy | ENSCAFG00000040375 | | 0.01609 |
| Energy | ENSCAFG00000015351 | *BAIAP2L1* | 0.01723 |
| Energy | ENSCAFG00000014169 | | 0.01850 |
| Energy | ENSCAFG00000011150 | *GALNT17* | 0.02069 |

| | | | |
|---|---|---|---|
| Energy | ENSCAFG00000006229 | *PCDH1* | 0.02161 |
| Energy | ENSCAFG00000004866 | *USP6NL* | 0.02195 |
| Energy | ENSCAFG00000038646 | | 0.02226 |
| Energy | ENSCAFG00000037023 | | 0.02550 |
| Energy | ENSCAFG00000005330 | *SH2D4A* | 0.02565 |
| Energy | ENSCAFG00000018351 | *ENOSF1* | 0.02919 |
| Energy | ENSCAFG00000028579 | *RTN4RL1* | 0.03023 |
| Energy | ENSCAFG00000001906 | *FZD1* | 0.03507 |
| Energy | ENSCAFG00000026050 | *RF00026* | 0.03566 |
| Energy | ENSCAFG00000017848 | | 0.03605 |
| Energy | ENSCAFG00000039266 | | 0.03703 |
| Energy | ENSCAFG00000038167 | | 0.03763 |
| Energy | ENSCAFG00000035032 | | 0.03887 |
| Energy | ENSCAFG00000023825 | | 0.03908 |
| Energy | ENSCAFG00000024967 | *ASIP* | 0.04016 |
| Energy | ENSCAFG00000003370 | *ASAP2* | 0.04098 |
| Energy | ENSCAFG00000038463 | | 0.04505 |
| Energy | ENSCAFG00000037362 | | 0.04664 |
| Energy | ENSCAFG00000037990 | | 0.04740 |
| Energy | ENSCAFG00000035285 | | 0.04960 |
| Energy | ENSCAFG00000001434 | *KDM4C* | 0.04984 |
| Excitability | ENSCAFG00000017252 | *ATRX* | < 0.00001 |
| Excitability | ENSCAFG00000007650 | *NEK1* | < 0.00001 |
| Excitability | ENSCAFG00000005914 | *TMTC2* | < 0.00001 |
| Excitability | ENSCAFG00000006743 | *PCMTD1* | < 0.00001 |
| Excitability | ENSCAFG00000038256 | | < 0.00001 |
| Excitability | ENSCAFG00000006732 | *PXDNL* | < 0.00001 |
| Excitability | ENSCAFG00000007301 | *CWC27* | < 0.00001 |
| Excitability | ENSCAFG00000031499 | *GLIS3* | < 0.00001 |

| | | | |
|---|---|---|---|
| Excitability | ENSCAFG00000000070 | *PHLPP1* | < 0.00001 |
| Excitability | ENSCAFG00000004087 | *MAML2* | < 0.00001 |
| Excitability | ENSCAFG00000006249 | *NDFIP1* | < 0.00001 |
| Excitability | ENSCAFG00000004160 | *TNIP3* | < 0.00001 |
| Excitability | ENSCAFG00000009849 | *SPTBN5* | < 0.00001 |
| Excitability | ENSCAFG00000013607 | *PDE11A* | < 0.00001 |
| Excitability | ENSCAFG00000006764 | *PARG* | < 0.00001 |
| Excitability | ENSCAFG00000039072 | | < 0.00001 |
| Excitability | ENSCAFG00000023079 | *AFF3* | < 0.00001 |
| Excitability | ENSCAFG00000027835 | *RF00026* | < 0.00001 |
| Excitability | ENSCAFG00000027967 | *RF00009* | < 0.00001 |
| Excitability | ENSCAFG00000007007 | *ATP8A2* | < 0.00001 |
| Excitability | ENSCAFG00000013221 | | < 0.00001 |
| Excitability | ENSCAFG00000001314 | *CACNA1I* | 0.00001 |
| Excitability | ENSCAFG00000008352 | *PHF20* | 0.00001 |
| Excitability | ENSCAFG00000033690 | | 0.00001 |
| Excitability | ENSCAFG00000010194 | *ABCC12* | 0.00001 |
| Excitability | ENSCAFG00000003465 | *EGFR* | 0.00001 |
| Excitability | ENSCAFG00000009662 | *MGA* | 0.00001 |
| Excitability | ENSCAFG00000006729 | *OGDHL* | 0.00001 |
| Excitability | ENSCAFG00000001557 | *CCDC171* | 0.00002 |
| Excitability | ENSCAFG00000014916 | *FAM227B* | 0.00002 |
| Excitability | ENSCAFG00000038695 | | 0.00003 |
| Excitability | ENSCAFG00000008293 | *FZD3* | 0.00003 |
| Excitability | ENSCAFG00000007379 | *CAMK4* | 0.00005 |
| Excitability | ENSCAFG00000004535 | *NR2C2* | 0.00005 |
| Excitability | ENSCAFG00000000935 | *CEP85L* | 0.00007 |
| Excitability | ENSCAFG00000006625 | *SFSWAP* | 0.00008 |
| Excitability | ENSCAFG00000007963 | *BLK* | 0.00008 |

| | | | |
|---|---|---|---|
| Excitability | ENSCAFG00000017622 | *PRKCB* | 0.00013 |
| Excitability | ENSCAFG00000010242 | *SLC4A10* | 0.00013 |
| Excitability | ENSCAFG00000024647 | *SERPINB5* | 0.00013 |
| Excitability | ENSCAFG00000033155 | | 0.00019 |
| Excitability | ENSCAFG00000038283 | | 0.00019 |
| Excitability | ENSCAFG00000005373 | *SLC24A3* | 0.00020 |
| Excitability | ENSCAFG00000004589 | | 0.00020 |
| Excitability | ENSCAFG00000038471 | | 0.00021 |
| Excitability | ENSCAFG00000033688 | | 0.00022 |
| Excitability | ENSCAFG00000011212 | *PIK3CA* | 0.00025 |
| Excitability | ENSCAFG00000000756 | *GRAMD4* | 0.00029 |
| Excitability | ENSCAFG00000002155 | | 0.00032 |
| Excitability | ENSCAFG00000000786 | *CSMD3* | 0.00035 |
| Excitability | ENSCAFG00000015932 | *DNAAF4* | 0.00040 |
| Excitability | ENSCAFG00000039585 | | 0.00041 |
| Excitability | ENSCAFG00000005054 | *ZRANB3* | 0.00042 |
| Excitability | ENSCAFG00000037577 | | 0.00044 |
| Excitability | ENSCAFG00000000337 | *PPM1H* | 0.00044 |
| Excitability | ENSCAFG00000019902 | *RAPGEF1* | 0.00051 |
| Excitability | ENSCAFG00000016732 | *SIPA1L1* | 0.00059 |
| Excitability | ENSCAFG00000026032 | *RF00425* | 0.00060 |
| Excitability | ENSCAFG00000011922 | *KPNA1* | 0.00069 |
| Excitability | ENSCAFG00000013228 | *CNKSR2* | 0.00121 |
| Excitability | ENSCAFG00000001930 | *PGM5* | 0.00132 |
| Excitability | ENSCAFG00000036039 | | 0.00146 |
| Excitability | ENSCAFG00000036709 | | 0.00158 |
| Excitability | ENSCAFG00000008761 | *CX3CL1* | 0.00166 |
| Excitability | ENSCAFG00000039058 | | 0.00181 |
| Excitability | ENSCAFG00000004632 | | 0.00195 |

| | | | |
|---|---|---|---|
| Excitability | ENSCAFG00000018100 | *SCAPER* | 0.00197 |
| Excitability | ENSCAFG00000026031 | *RF00026* | 0.00208 |
| Excitability | ENSCAFG00000012890 | *HAT1* | 0.00211 |
| Excitability | ENSCAFG00000031780 | | 0.00248 |
| Excitability | ENSCAFG00000012478 | *PLEKHA5* | 0.00251 |
| Excitability | ENSCAFG00000008350 | *PKIA* | 0.00256 |
| Excitability | ENSCAFG00000020376 | *ZZZ3* | 0.00260 |
| Excitability | ENSCAFG00000035101 | | 0.00286 |
| Excitability | ENSCAFG00000018151 | *SLC47A2* | 0.00314 |
| Excitability | ENSCAFG00000019051 | | 0.00352 |
| Excitability | ENSCAFG00000022264 | *RF00015* | 0.00370 |
| Excitability | ENSCAFG00000035453 | | 0.00375 |
| Excitability | ENSCAFG00000040724 | | 0.00443 |
| Excitability | ENSCAFG00000040631 | | 0.00449 |
| Excitability | ENSCAFG00000002671 | *SP4* | 0.00452 |
| Excitability | ENSCAFG00000015721 | *LIPA* | 0.00473 |
| Excitability | ENSCAFG00000031235 | | 0.00486 |
| Excitability | ENSCAFG00000013838 | *MASP1* | 0.00511 |
| Excitability | ENSCAFG00000018971 | | 0.00553 |
| Excitability | ENSCAFG00000029961 | *APCDD1L* | 0.00554 |
| Excitability | ENSCAFG00000010221 | *SUFU* | 0.00555 |
| Excitability | ENSCAFG00000007361 | *EPB41L4A* | 0.00601 |
| Excitability | ENSCAFG00000015648 | | 0.00697 |
| Excitability | ENSCAFG00000033410 | | 0.00796 |
| Excitability | ENSCAFG00000031045 | | 0.00906 |
| Excitability | ENSCAFG00000031721 | *CPNE1* | 0.00927 |
| Excitability | ENSCAFG00000014454 | *ATM* | 0.01005 |
| Excitability | ENSCAFG00000002045 | | 0.01103 |
| Excitability | ENSCAFG00000029313 | *CLDN1* | 0.01301 |

| | | | |
|---|---|---|---|
| Excitability | ENSCAFG00000013891 | *SST* | 0.01353 |
| Excitability | ENSCAFG00000015943 | *PRTG* | 0.01407 |
| Excitability | ENSCAFG00000005271 | *C22H16orf87* | 0.01413 |
| Excitability | ENSCAFG00000007079 | | 0.01578 |
| Excitability | ENSCAFG00000008058 | *MTMR9* | 0.01686 |
| Excitability | ENSCAFG00000008568 | *AGPAT5* | 0.01740 |
| Excitability | ENSCAFG00000039209 | | 0.01751 |
| Excitability | ENSCAFG00000007970 | *FAM167A* | 0.01755 |
| Excitability | ENSCAFG00000012179 | *UNC45A* | 0.01772 |
| Excitability | ENSCAFG00000005140 | *VHL* | 0.01775 |
| Excitability | ENSCAFG00000018979 | *ZIC3* | 0.01916 |
| Excitability | ENSCAFG00000002364 | *ADGRL3* | 0.02048 |
| Excitability | ENSCAFG00000015859 | *JAKMIP1* | 0.02067 |
| Excitability | ENSCAFG00000007789 | *ARHGAP10* | 0.02103 |
| Excitability | ENSCAFG00000006555 | *PRKDC* | 0.02145 |
| Excitability | ENSCAFG00000007912 | *TRPC4AP* | 0.02300 |
| Excitability | ENSCAFG00000013925 | *SEMA6D* | 0.02603 |
| Excitability | ENSCAFG00000038846 | | 0.02611 |
| Excitability | ENSCAFG00000018123 | *ARSI* | 0.02667 |
| Excitability | ENSCAFG00000032470 | | 0.02682 |
| Excitability | ENSCAFG00000009371 | *SORBS3* | 0.02791 |
| Excitability | ENSCAFG00000010988 | *CFAP410* | 0.03206 |
| Excitability | ENSCAFG00000000125 | *NARS* | 0.03299 |
| Excitability | ENSCAFG00000009367 | *HPS1* | 0.03302 |
| Excitability | ENSCAFG00000017958 | *COL5A3* | 0.03321 |
| Excitability | ENSCAFG00000020310 | *SMPD3* | 0.03369 |
| Excitability | ENSCAFG00000001106 | *LAMA2* | 0.03556 |
| Excitability | ENSCAFG00000036231 | | 0.03617 |
| Excitability | ENSCAFG00000022350 | *RF00026* | 0.03627 |

| | | | |
|---|---|---|---|
| Excitability | ENSCAFG00000029157 | *NOVA1* | 0.03681 |
| Excitability | ENSCAFG00000036595 | | 0.03712 |
| Excitability | ENSCAFG00000033823 | | 0.03863 |
| Excitability | ENSCAFG00000011934 | *ARHGEF12* | 0.03996 |
| Excitability | ENSCAFG00000006050 | *MYO16* | 0.04113 |
| Excitability | ENSCAFG00000020112 | *ABCD3* | 0.04298 |
| Excitability | ENSCAFG00000036099 | | 0.04326 |
| Excitability | ENSCAFG00000034875 | | 0.04335 |
| Excitability | ENSCAFG00000006809 | *NCOA4* | 0.04675 |
| Excitability | ENSCAFG00000003737 | *PARD3* | 0.04921 |
| Nonsocial Fear | ENSCAFG00000005775 | *BIRC6* | < 0.00001 |
| Nonsocial Fear | ENSCAFG00000038400 | | < 0.00001 |
| Nonsocial Fear | ENSCAFG00000009386 | *HPSE2* | < 0.00001 |
| Nonsocial Fear | ENSCAFG00000010026 | *SMARCAD1* | < 0.00001 |
| Nonsocial Fear | ENSCAFG00000030358 | *RF00026* | < 0.00001 |
| Nonsocial Fear | ENSCAFG00000009227 | *WDR41* | < 0.00001 |
| Nonsocial Fear | ENSCAFG00000001460 | *PTPRD* | < 0.00001 |
| Nonsocial Fear | ENSCAFG00000000955 | *TBC1D32* | < 0.00001 |
| Nonsocial Fear | ENSCAFG00000035815 | | < 0.00001 |
| Nonsocial Fear | ENSCAFG00000033300 | | < 0.00001 |
| Nonsocial Fear | ENSCAFG00000034053 | | < 0.00001 |
| Nonsocial Fear | ENSCAFG00000017256 | | < 0.00001 |
| Nonsocial Fear | ENSCAFG00000033142 | | < 0.00001 |
| Nonsocial Fear | ENSCAFG00000034838 | | < 0.00001 |
| Nonsocial Fear | ENSCAFG00000040257 | | < 0.00001 |
| Nonsocial Fear | ENSCAFG00000011680 | *ATP11B* | < 0.00001 |
| Nonsocial Fear | ENSCAFG00000004432 | *RCBTB2* | < 0.00001 |
| Nonsocial Fear | ENSCAFG00000017297 | *GPR174* | < 0.00001 |
| Nonsocial Fear | ENSCAFG00000017327 | *BRWD3* | < 0.00001 |

| | | | |
|---|---|---|---|
| Nonsocial Fear | ENSCAFG00000011936 | *TRPM8* | < 0.00001 |
| Nonsocial Fear | ENSCAFG00000012098 | *PGBD5* | 0.00001 |
| Nonsocial Fear | ENSCAFG00000002160 | *SPTLC1* | 0.00001 |
| Nonsocial Fear | ENSCAFG00000014245 | *PDE1A* | 0.00001 |
| Nonsocial Fear | ENSCAFG00000018635 | *ROR1* | 0.00001 |
| Nonsocial Fear | ENSCAFG00000003161 | *BBS9* | 0.00001 |
| Nonsocial Fear | ENSCAFG00000009764 | *PPHLN1* | 0.00004 |
| Nonsocial Fear | ENSCAFG00000033248 | | 0.00005 |
| Nonsocial Fear | ENSCAFG00000006592 | *STK32A* | 0.00007 |
| Nonsocial Fear | ENSCAFG00000006278 | *ATRN* | 0.00011 |
| Nonsocial Fear | ENSCAFG00000003479 | | 0.00012 |
| Nonsocial Fear | ENSCAFG00000002527 | *LRPPRC* | 0.00013 |
| Nonsocial Fear | ENSCAFG00000029154 | *EFCAB2* | 0.00014 |
| Nonsocial Fear | ENSCAFG00000004568 | *CUBN* | 0.00019 |
| Nonsocial Fear | ENSCAFG00000004377 | *ACBD5* | 0.00021 |
| Nonsocial Fear | ENSCAFG00000007420 | *CBFA2T2* | 0.00029 |
| Nonsocial Fear | ENSCAFG00000017294 | *P2RY10* | 0.00038 |
| Nonsocial Fear | ENSCAFG00000016161 | *FAM114A1* | 0.00053 |
| Nonsocial Fear | ENSCAFG00000011430 | *STXBP5L* | 0.00066 |
| Nonsocial Fear | ENSCAFG00000004186 | | 0.00074 |
| Nonsocial Fear | ENSCAFG00000018580 | | 0.00088 |
| Nonsocial Fear | ENSCAFG00000028102 | *RF00088* | 0.00094 |
| Nonsocial Fear | ENSCAFG00000037837 | | 0.00104 |
| Nonsocial Fear | ENSCAFG00000002667 | | 0.00114 |
| Nonsocial Fear | ENSCAFG00000002390 | *FAM178B* | 0.00116 |
| Nonsocial Fear | ENSCAFG00000034178 | | 0.00124 |
| Nonsocial Fear | ENSCAFG00000001106 | *LAMA2* | 0.00128 |
| Nonsocial Fear | ENSCAFG00000032645 | *RF00026* | 0.00135 |
| Nonsocial Fear | ENSCAFG00000036236 | | 0.00147 |

| | | | |
|---|---|---|---|
| Nonsocial Fear | ENSCAFG00000039648 | | 0.00171 |
| Nonsocial Fear | ENSCAFG00000039818 | | 0.00179 |
| Nonsocial Fear | ENSCAFG00000034343 | | 0.00181 |
| Nonsocial Fear | ENSCAFG00000023549 | *C9orf3* | 0.00215 |
| Nonsocial Fear | ENSCAFG00000006691 | *SLC46A3* | 0.00254 |
| Nonsocial Fear | ENSCAFG00000001362 | *PLXNA4* | 0.00266 |
| Nonsocial Fear | ENSCAFG00000018100 | *SCAPER* | 0.00308 |
| Nonsocial Fear | ENSCAFG00000036778 | | 0.00341 |
| Nonsocial Fear | ENSCAFG00000014569 | *PDCD10* | 0.00346 |
| Nonsocial Fear | ENSCAFG00000030067 | | 0.00351 |
| Nonsocial Fear | ENSCAFG00000016051 | *CACNA1C* | 0.00378 |
| Nonsocial Fear | ENSCAFG00000034666 | | 0.00405 |
| Nonsocial Fear | ENSCAFG00000009417 | *CDYL* | 0.00419 |
| Nonsocial Fear | ENSCAFG00000017252 | *ATRX* | 0.00432 |
| Nonsocial Fear | ENSCAFG00000027810 | *RF00026* | 0.00436 |
| Nonsocial Fear | ENSCAFG00000019384 | *DIRAS1* | 0.00442 |
| Nonsocial Fear | ENSCAFG00000018702 | *EFCAB7* | 0.00460 |
| Nonsocial Fear | ENSCAFG00000005704 | *TMEM173* | 0.00474 |
| Nonsocial Fear | ENSCAFG00000033801 | | 0.00489 |
| Nonsocial Fear | ENSCAFG00000008945 | *SLIT1* | 0.00491 |
| Nonsocial Fear | ENSCAFG00000036586 | | 0.00497 |
| Nonsocial Fear | ENSCAFG00000037340 | | 0.00528 |
| Nonsocial Fear | ENSCAFG00000027105 | *RF00026* | 0.00702 |
| Nonsocial Fear | ENSCAFG00000005720 | *CLPB* | 0.00750 |
| Nonsocial Fear | ENSCAFG00000017645 | *MYO9A* | 0.00768 |
| Nonsocial Fear | ENSCAFG00000019831 | *SYPL2* | 0.00957 |
| Nonsocial Fear | ENSCAFG00000035944 | | 0.01020 |
| Nonsocial Fear | ENSCAFG00000038749 | | 0.01047 |
| Nonsocial Fear | ENSCAFG00000040591 | | 0.01333 |

| | | | |
|---|---|---|---|
| Nonsocial Fear | ENSCAFG00000008882 | *LEKR1* | 0.01452 |
| Nonsocial Fear | ENSCAFG00000009885 | *JARID2* | 0.01536 |
| Nonsocial Fear | ENSCAFG00000035143 | | 0.01594 |
| Nonsocial Fear | ENSCAFG00000016982 | *DOCK2* | 0.01772 |
| Nonsocial Fear | ENSCAFG00000017295 | | 0.01848 |
| Nonsocial Fear | ENSCAFG00000022548 | *RF00001* | 0.01944 |
| Nonsocial Fear | ENSCAFG00000017502 | *ANP32A* | 0.02075 |
| Nonsocial Fear | ENSCAFG00000034943 | | 0.02160 |
| Nonsocial Fear | ENSCAFG00000001091 | *TGFBI* | 0.02235 |
| Nonsocial Fear | ENSCAFG00000037802 | | 0.02346 |
| Nonsocial Fear | ENSCAFG00000036735 | | 0.02370 |
| Nonsocial Fear | ENSCAFG00000009624 | *CWF19L1* | 0.02544 |
| Nonsocial Fear | ENSCAFG00000027110 | *RF00026* | 0.03251 |
| Nonsocial Fear | ENSCAFG00000040668 | | 0.03632 |
| Nonsocial Fear | ENSCAFG00000031017 | *SMIM33* | 0.03717 |
| Nonsocial Fear | ENSCAFG00000014030 | *LARS2* | 0.03836 |
| Nonsocial Fear | ENSCAFG00000007898 | *CLIP1* | 0.03873 |
| Nonsocial Fear | ENSCAFG00000039348 | | 0.03925 |
| Nonsocial Fear | ENSCAFG00000008630 | *KIFC3* | 0.04105 |
| Owner Aggression | ENSCAFG00000033361 | | < 0.00001 |
| Owner Aggression | ENSCAFG00000006730 | *TTC17* | < 0.00001 |
| Owner Aggression | ENSCAFG00000018100 | *SCAPER* | < 0.00001 |
| Owner Aggression | ENSCAFG00000011983 | *HSPBAP1* | < 0.00001 |
| Owner Aggression | ENSCAFG00000018811 | | < 0.00001 |
| Owner Aggression | ENSCAFG00000013779 | | < 0.00001 |
| Owner Aggression | ENSCAFG00000015639 | *MNAT1* | < 0.00001 |
| Owner Aggression | ENSCAFG00000003749 | *SLC7A11* | < 0.00001 |
| Owner Aggression | ENSCAFG00000001646 | *PRUNE2* | < 0.00001 |
| Owner Aggression | ENSCAFG00000038128 | | < 0.00001 |

| | | | |
|---|---|---|---|
| Owner Aggression | ENSCAFG00000020206 | *SCAI* | < 0.00001 |
| Owner Aggression | ENSCAFG00000004157 | *RAB7A* | < 0.00001 |
| Owner Aggression | ENSCAFG00000001630 | *MYH9* | < 0.00001 |
| Owner Aggression | ENSCAFG00000013288 | *ADAMTSL3* | < 0.00001 |
| Owner Aggression | ENSCAFG00000000770 | | < 0.00001 |
| Owner Aggression | ENSCAFG00000039058 | | < 0.00001 |
| Owner Aggression | ENSCAFG00000036018 | | < 0.00001 |
| Owner Aggression | ENSCAFG00000018971 | | < 0.00001 |
| Owner Aggression | ENSCAFG00000001557 | *CCDC171* | < 0.00001 |
| Owner Aggression | ENSCAFG00000035215 | | < 0.00001 |
| Owner Aggression | ENSCAFG00000027188 | *RF00026* | < 0.00001 |
| Owner Aggression | ENSCAFG00000002271 | *PHF14* | < 0.00001 |
| Owner Aggression | ENSCAFG00000034411 | | < 0.00001 |
| Owner Aggression | ENSCAFG00000005121 | *SFMBT2* | < 0.00001 |
| Owner Aggression | ENSCAFG00000014257 | *PPP2R2C* | < 0.00001 |
| Owner Aggression | ENSCAFG00000010120 | *RAD54L2* | < 0.00001 |
| Owner Aggression | ENSCAFG00000016511 | *RFX1* | < 0.00001 |
| Owner Aggression | ENSCAFG00000023938 | *GPR158* | < 0.00001 |
| Owner Aggression | ENSCAFG00000009669 | *MDM4* | 0.00001 |
| Owner Aggression | ENSCAFG00000011940 | *PARP9* | 0.00001 |
| Owner Aggression | ENSCAFG00000013840 | | 0.00003 |
| Owner Aggression | ENSCAFG00000038859 | | 0.00003 |
| Owner Aggression | ENSCAFG00000018799 | *IGSF1* | 0.00003 |
| Owner Aggression | ENSCAFG00000006492 | *B4GALNT4* | 0.00003 |
| Owner Aggression | ENSCAFG00000001700 | *SND1* | 0.00003 |
| Owner Aggression | ENSCAFG00000006989 | *DEFB119* | 0.00004 |
| Owner Aggression | ENSCAFG00000032160 | | 0.00007 |
| Owner Aggression | ENSCAFG00000007420 | *CBFA2T2* | 0.00007 |
| Owner Aggression | ENSCAFG00000007107 | *TTLL9* | 0.00010 |

| | | | |
|---|---|---|---|
| Owner Aggression | ENSCAFG00000033119 | | 0.00014 |
| Owner Aggression | ENSCAFG00000019958 | *NTNG1* | 0.00014 |
| Owner Aggression | ENSCAFG00000003580 | *GRIK2* | 0.00020 |
| Owner Aggression | ENSCAFG00000018777 | *ENOX2* | 0.00024 |
| Owner Aggression | ENSCAFG00000025956 | *RF00001* | 0.00032 |
| Owner Aggression | ENSCAFG00000034449 | | 0.00041 |
| Owner Aggression | ENSCAFG00000015618 | *TULP3* | 0.00046 |
| Owner Aggression | ENSCAFG00000016982 | *DOCK2* | 0.00047 |
| Owner Aggression | ENSCAFG00000006462 | *UBXN8* | 0.00057 |
| Owner Aggression | ENSCAFG00000003705 | *CUL2* | 0.00068 |
| Owner Aggression | ENSCAFG00000003106 | *EHBP1* | 0.00070 |
| Owner Aggression | ENSCAFG00000002949 | *GC* | 0.00070 |
| Owner Aggression | ENSCAFG00000001166 | *KHDRBS3* | 0.00090 |
| Owner Aggression | ENSCAFG00000018752 | *IL7R* | 0.00099 |
| Owner Aggression | ENSCAFG00000004764 | *VWA8* | 0.00115 |
| Owner Aggression | ENSCAFG00000011176 | *NR5A2* | 0.00132 |
| Owner Aggression | ENSCAFG00000010222 | *BARX2* | 0.00140 |
| Owner Aggression | ENSCAFG00000035525 | | 0.00143 |
| Owner Aggression | ENSCAFG00000004671 | *FAM216B* | 0.00144 |
| Owner Aggression | ENSCAFG00000005164 | | 0.00185 |
| Owner Aggression | ENSCAFG00000006573 | *FAM19A4* | 0.00190 |
| Owner Aggression | ENSCAFG00000014579 | *PRDX6* | 0.00228 |
| Owner Aggression | ENSCAFG00000002714 | *EPHA5* | 0.00241 |
| Owner Aggression | ENSCAFG00000028804 | *RNF150* | 0.00258 |
| Owner Aggression | ENSCAFG00000012729 | *CGN* | 0.00272 |
| Owner Aggression | ENSCAFG00000037915 | | 0.00291 |
| Owner Aggression | ENSCAFG00000004343 | *EFCC1* | 0.00329 |
| Owner Aggression | ENSCAFG00000031347 | | 0.00356 |
| Owner Aggression | ENSCAFG00000017676 | *SETBP1* | 0.00398 |

| | | | |
|---|---|---|---|
| Owner Aggression | ENSCAFG00000039028 | | 0.00418 |
| Owner Aggression | ENSCAFG00000035446 | | 0.00513 |
| Owner Aggression | ENSCAFG00000002796 | *CCDC88A* | 0.00518 |
| Owner Aggression | ENSCAFG00000023541 | | 0.00541 |
| Owner Aggression | ENSCAFG00000023322 | *RASSF9* | 0.00624 |
| Owner Aggression | ENSCAFG00000012545 | *CARF* | 0.00646 |
| Owner Aggression | ENSCAFG00000000446 | *PTPRB* | 0.00668 |
| Owner Aggression | ENSCAFG00000023267 | *PRIM2* | 0.00672 |
| Owner Aggression | ENSCAFG00000031408 | *CFAP299* | 0.00714 |
| Owner Aggression | ENSCAFG00000018785 | *ARHGAP36* | 0.00739 |
| Owner Aggression | ENSCAFG00000035968 | | 0.00740 |
| Owner Aggression | ENSCAFG00000005607 | *SPTLC3* | 0.00884 |
| Owner Aggression | ENSCAFG00000031628 | | 0.00966 |
| Owner Aggression | ENSCAFG00000038140 | | 0.00985 |
| Owner Aggression | ENSCAFG00000019882 | *TSC1* | 0.01045 |
| Owner Aggression | ENSCAFG00000033801 | | 0.01087 |
| Owner Aggression | ENSCAFG00000039505 | | 0.01098 |
| Owner Aggression | ENSCAFG00000040675 | | 0.01144 |
| Owner Aggression | ENSCAFG00000029470 | | 0.01163 |
| Owner Aggression | ENSCAFG00000017787 | | 0.01229 |
| Owner Aggression | ENSCAFG00000015877 | *SH2D4B* | 0.01237 |
| Owner Aggression | ENSCAFG00000003323 | *KIDINS220* | 0.01291 |
| Owner Aggression | ENSCAFG00000012804 | *MACO1* | 0.01298 |
| Owner Aggression | ENSCAFG00000004224 | *PLEKHB2* | 0.01333 |
| Owner Aggression | ENSCAFG00000014892 | | 0.01382 |
| Owner Aggression | ENSCAFG00000038826 | | 0.01599 |
| Owner Aggression | ENSCAFG00000000828 | *RPS6KA2* | 0.01674 |
| Owner Aggression | ENSCAFG00000014718 | *CWF19L2* | 0.01685 |
| Owner Aggression | ENSCAFG00000026119 | *RF00026* | 0.01702 |

| | | | |
|---|---|---|---|
| Owner Aggression | ENSCAFG00000027541 | *RF00003* | 0.01762 |
| Owner Aggression | ENSCAFG00000012413 | *RPS6KC1* | 0.01875 |
| Owner Aggression | ENSCAFG00000008237 | *HMBOX1* | 0.01890 |
| Owner Aggression | ENSCAFG00000016111 | *FOXK1* | 0.01931 |
| Owner Aggression | ENSCAFG00000038496 | | 0.02112 |
| Owner Aggression | ENSCAFG00000010857 | *MMAB* | 0.02303 |
| Owner Aggression | ENSCAFG00000037664 | | 0.02539 |
| Owner Aggression | ENSCAFG00000039151 | | 0.02777 |
| Owner Aggression | ENSCAFG00000000086 | *CDH20* | 0.03216 |
| Owner Aggression | ENSCAFG00000014093 | | 0.03278 |
| Owner Aggression | ENSCAFG00000038122 | | 0.03378 |
| Owner Aggression | ENSCAFG00000033917 | | 0.03395 |
| Owner Aggression | ENSCAFG00000011672 | *SLC15A2* | 0.03450 |
| Owner Aggression | ENSCAFG00000013410 | *RGL1* | 0.04037 |
| Owner Aggression | ENSCAFG00000009318 | *CCDC148* | 0.04495 |
| Separation Problems | ENSCAFG00000028659 | | < 0.00001 |
| Separation Problems | ENSCAFG00000035085 | | < 0.00001 |
| Separation Problems | ENSCAFG00000031499 | *GLIS3* | < 0.00001 |
| Separation Problems | ENSCAFG00000013221 | | < 0.00001 |
| Separation Problems | ENSCAFG00000011820 | *PRPF3* | < 0.00001 |
| Separation Problems | ENSCAFG00000001251 | | < 0.00001 |
| Separation Problems | ENSCAFG00000039076 | | < 0.00001 |
| Separation Problems | ENSCAFG00000004087 | *MAML2* | < 0.00001 |
| Separation Problems | ENSCAFG00000015459 | | < 0.00001 |
| Separation Problems | ENSCAFG00000040856 | | < 0.00001 |
| Separation Problems | ENSCAFG00000002096 | *UXS1* | < 0.00001 |
| Separation Problems | ENSCAFG00000039762 | | < 0.00001 |
| Separation Problems | ENSCAFG00000005746 | *NAV3* | < 0.00001 |
| Separation Problems | ENSCAFG00000002667 | | < 0.00001 |

| | | | |
|---|---|---|---|
| Separation Problems | ENSCAFG00000015477 | | < 0.00001 |
| Separation Problems | ENSCAFG00000036551 | | < 0.00001 |
| Separation Problems | ENSCAFG00000000745 | *TBC1D22A* | < 0.00001 |
| Separation Problems | ENSCAFG00000017685 | *RIT2* | < 0.00001 |
| Separation Problems | ENSCAFG00000000756 | *GRAMD4* | < 0.00001 |
| Separation Problems | ENSCAFG00000002413 | *DST* | 0.00001 |
| Separation Problems | ENSCAFG00000000164 | *SMAD4* | 0.00001 |
| Separation Problems | ENSCAFG00000019132 | *RPH3AL* | 0.00001 |
| Separation Problems | ENSCAFG00000012545 | *CARF* | 0.00001 |
| Separation Problems | ENSCAFG00000013926 | *FBXO33* | 0.00002 |
| Separation Problems | ENSCAFG00000009318 | *CCDC148* | 0.00002 |
| Separation Problems | ENSCAFG00000001002 | *SMPDL3A* | 0.00002 |
| Separation Problems | ENSCAFG00000004092 | *MTMR2* | 0.00003 |
| Separation Problems | ENSCAFG00000014975 | *EIF4G3* | 0.00003 |
| Separation Problems | ENSCAFG00000039187 | | 0.00003 |
| Separation Problems | ENSCAFG00000003570 | *FBXW2* | 0.00004 |
| Separation Problems | ENSCAFG00000010120 | *RAD54L2* | 0.00004 |
| Separation Problems | ENSCAFG00000040676 | | 0.00004 |
| Separation Problems | ENSCAFG00000004593 | *CCDC90B* | 0.00007 |
| Separation Problems | ENSCAFG00000000767 | *CELSR1* | 0.00009 |
| Separation Problems | ENSCAFG00000030067 | | 0.00010 |
| Separation Problems | ENSCAFG00000006743 | *PCMTD1* | 0.00016 |
| Separation Problems | ENSCAFG00000008827 | *ALMS1* | 0.00016 |
| Separation Problems | ENSCAFG00000026031 | *RF00026* | 0.00016 |
| Separation Problems | ENSCAFG00000000334 | *ADAMTS2* | 0.00020 |
| Separation Problems | ENSCAFG00000018041 | *WDR20* | 0.00023 |
| Separation Problems | ENSCAFG00000036684 | | 0.00023 |
| Separation Problems | ENSCAFG00000036899 | | 0.00025 |
| Separation Problems | ENSCAFG00000033314 | | 0.00027 |

| | | | |
|---|---|---|---|
| Separation Problems | ENSCAFG00000005270 | *CFAP61* | 0.00030 |
| Separation Problems | ENSCAFG00000018100 | *SCAPER* | 0.00035 |
| Separation Problems | ENSCAFG00000033275 | | 0.00041 |
| Separation Problems | ENSCAFG00000001161 | *OBSCN* | 0.00045 |
| Separation Problems | ENSCAFG00000010377 | | 0.00046 |
| Separation Problems | ENSCAFG00000016354 | *STIM2* | 0.00053 |
| Separation Problems | ENSCAFG00000016568 | *ADAM10* | 0.00053 |
| Separation Problems | ENSCAFG00000017765 | *GRIA1* | 0.00061 |
| Separation Problems | ENSCAFG00000031802 | *ISPD* | 0.00064 |
| Separation Problems | ENSCAFG00000000945 | *MAN1A1* | 0.00070 |
| Separation Problems | ENSCAFG00000025220 | *ACVRL1* | 0.00074 |
| Separation Problems | ENSCAFG00000005720 | *CLPB* | 0.00079 |
| Separation Problems | ENSCAFG00000039072 | | 0.00080 |
| Separation Problems | ENSCAFG00000002762 | *EML6* | 0.00099 |
| Separation Problems | ENSCAFG00000034267 | | 0.00102 |
| Separation Problems | ENSCAFG00000032971 | | 0.00104 |
| Separation Problems | ENSCAFG00000038198 | | 0.00107 |
| Separation Problems | ENSCAFG00000017787 | | 0.00181 |
| Separation Problems | ENSCAFG00000000169 | *MRO* | 0.00201 |
| Separation Problems | ENSCAFG00000004545 | *TRDMT1* | 0.00210 |
| Separation Problems | ENSCAFG00000007568 | *CPA6* | 0.00234 |
| Separation Problems | ENSCAFG00000005450 | *KAT14* | 0.00246 |
| Separation Problems | ENSCAFG00000021196 | *RF00100* | 0.00258 |
| Separation Problems | ENSCAFG00000015993 | *DCP1B* | 0.00258 |
| Separation Problems | ENSCAFG00000006103 | *NEK11* | 0.00308 |
| Separation Problems | ENSCAFG00000022418 | *RF00100* | 0.00329 |
| Separation Problems | ENSCAFG00000038028 | | 0.00331 |
| Separation Problems | ENSCAFG00000035474 | | 0.00335 |
| Separation Problems | ENSCAFG00000013410 | *RGL1* | 0.00339 |

| | | | |
|---|---|---|---|
| Separation Problems | ENSCAFG00000013927 | | 0.00347 |
| Separation Problems | ENSCAFG00000009831 | *PRMT3* | 0.00356 |
| Separation Problems | ENSCAFG00000039158 | | 0.00372 |
| Separation Problems | ENSCAFG00000016511 | *RFX1* | 0.00395 |
| Separation Problems | ENSCAFG00000009799 | *NCAPD3* | 0.00471 |
| Separation Problems | ENSCAFG00000030219 | | 0.00484 |
| Separation Problems | ENSCAFG00000006042 | *SLC23A2* | 0.00501 |
| Separation Problems | ENSCAFG00000023562 | *DMD* | 0.00523 |
| Separation Problems | ENSCAFG00000003752 | | 0.00598 |
| Separation Problems | ENSCAFG00000032668 | *SPOCK1* | 0.00728 |
| Separation Problems | ENSCAFG00000015800 | *MYO5A* | 0.00737 |
| Separation Problems | ENSCAFG00000020112 | *ABCD3* | 0.00739 |
| Separation Problems | ENSCAFG00000016588 | *C20H19orf57* | 0.00749 |
| Separation Problems | ENSCAFG00000008413 | *EPB41L1* | 0.00769 |
| Separation Problems | ENSCAFG00000015742 | | 0.00820 |
| Separation Problems | ENSCAFG00000018156 | *SMG1* | 0.00831 |
| Separation Problems | ENSCAFG00000010026 | *SMARCAD1* | 0.00851 |
| Separation Problems | ENSCAFG00000015686 | | 0.00904 |
| Separation Problems | ENSCAFG00000032250 | *OR4N2* | 0.00989 |
| Separation Problems | ENSCAFG00000013052 | *TANC2* | 0.00997 |
| Separation Problems | ENSCAFG00000039437 | | 0.01024 |
| Separation Problems | ENSCAFG00000006762 | *ST18* | 0.01037 |
| Separation Problems | ENSCAFG00000013503 | *DGKG* | 0.01046 |
| Separation Problems | ENSCAFG00000001532 | *ZDHHC21* | 0.01123 |
| Separation Problems | ENSCAFG00000012301 | *TNS3* | 0.01191 |
| Separation Problems | ENSCAFG00000036504 | | 0.01457 |
| Separation Problems | ENSCAFG00000020187 | *HFM1* | 0.01504 |
| Separation Problems | ENSCAFG00000015862 | *FAM214A* | 0.01585 |
| Separation Problems | ENSCAFG00000002091 | *RCL1* | 0.01894 |

| | | | |
|---|---|---|---|
| Separation Problems | ENSCAFG00000000166 | *ELAC1* | 0.02106 |
| Separation Problems | ENSCAFG00000001312 | *CHCHD3* | 0.02198 |
| Separation Problems | ENSCAFG00000014112 | *KIF15* | 0.02469 |
| Separation Problems | ENSCAFG00000029321 | *WNT5A* | 0.02496 |
| Separation Problems | ENSCAFG00000008805 | | 0.02522 |
| Separation Problems | ENSCAFG00000008340 | *RACGAP1* | 0.02523 |
| Separation Problems | ENSCAFG00000034552 | | 0.02533 |
| Separation Problems | ENSCAFG00000016310 | *IKZF3* | 0.02916 |
| Separation Problems | ENSCAFG00000030017 | | 0.03048 |
| Separation Problems | ENSCAFG00000008846 | *FBXO41* | 0.03164 |
| Separation Problems | ENSCAFG00000014146 | *ZNF385B* | 0.03210 |
| Separation Problems | ENSCAFG00000006520 | *KCNG3* | 0.03213 |
| Separation Problems | ENSCAFG00000035538 | | 0.03398 |
| Separation Problems | ENSCAFG00000011585 | *PCNX2* | 0.03692 |
| Separation Problems | ENSCAFG00000006668 | *KBTBD8* | 0.04178 |
| Separation Problems | ENSCAFG00000030145 | *AUH* | 0.04354 |
| Separation Problems | ENSCAFG00000031998 | *RF00026* | 0.04627 |
| Separation Problems | ENSCAFG00000000557 | *CSNK1G3* | 0.04678 |
| Stranger Aggression | ENSCAFG00000000079 | *RELCH* | < 0.00001 |
| Stranger Aggression | ENSCAFG00000003678 | *CCNY* | < 0.00001 |
| Stranger Aggression | ENSCAFG00000004408 | *CAB39L* | < 0.00001 |
| Stranger Aggression | ENSCAFG00000039115 | | < 0.00001 |
| Stranger Aggression | ENSCAFG00000009971 | *ARMH3* | < 0.00001 |
| Stranger Aggression | ENSCAFG00000004341 | *SETDB2* | < 0.00001 |
| Stranger Aggression | ENSCAFG00000004325 | *KPNA3* | < 0.00001 |
| Stranger Aggression | ENSCAFG00000004345 | *RCBTB1* | < 0.00001 |
| Stranger Aggression | ENSCAFG00000010619 | *BICD1* | < 0.00001 |
| Stranger Aggression | ENSCAFG00000031429 | *GNA14* | < 0.00001 |
| Stranger Aggression | ENSCAFG00000004379 | *FNDC3A* | < 0.00001 |

| | | | |
|---|---|---|---|
| Stranger Aggression | ENSCAFG00000004478 | *LRCH1* | < 0.00001 |
| Stranger Aggression | ENSCAFG00000002257 | *ICA1* | < 0.00001 |
| Stranger Aggression | ENSCAFG00000035101 | | < 0.00001 |
| Stranger Aggression | ENSCAFG00000004436 | *RB1* | < 0.00001 |
| Stranger Aggression | ENSCAFG00000011415 | *MCUB* | < 0.00001 |
| Stranger Aggression | ENSCAFG00000003737 | *PARD3* | < 0.00001 |
| Stranger Aggression | ENSCAFG00000031367 | | < 0.00001 |
| Stranger Aggression | ENSCAFG00000005667 | *NEK10* | < 0.00001 |
| Stranger Aggression | ENSCAFG00000003705 | *CUL2* | < 0.00001 |
| Stranger Aggression | ENSCAFG00000001560 | *KIF6* | < 0.00001 |
| Stranger Aggression | ENSCAFG00000033999 | | < 0.00001 |
| Stranger Aggression | ENSCAFG00000005326 | *UVRAG* | < 0.00001 |
| Stranger Aggression | ENSCAFG00000036142 | | < 0.00001 |
| Stranger Aggression | ENSCAFG00000013410 | *RGL1* | < 0.00001 |
| Stranger Aggression | ENSCAFG00000005954 | *SUMF1* | < 0.00001 |
| Stranger Aggression | ENSCAFG00000000353 | *STXBP5* | < 0.00001 |
| Stranger Aggression | ENSCAFG00000005707 | *OSBPL8* | < 0.00001 |
| Stranger Aggression | ENSCAFG00000037076 | | 0.00001 |
| Stranger Aggression | ENSCAFG00000034987 | | 0.00002 |
| Stranger Aggression | ENSCAFG00000009031 | *CMYA5* | 0.00002 |
| Stranger Aggression | ENSCAFG00000018882 | *FAM122B* | 0.00002 |
| Stranger Aggression | ENSCAFG00000001557 | *CCDC171* | 0.00002 |
| Stranger Aggression | ENSCAFG00000002364 | *ADGRL3* | 0.00003 |
| Stranger Aggression | ENSCAFG00000014030 | *LARS2* | 0.00005 |
| Stranger Aggression | ENSCAFG00000034539 | | 0.00005 |
| Stranger Aggression | ENSCAFG00000005040 | *KLF12* | 0.00006 |
| Stranger Aggression | ENSCAFG00000034005 | | 0.00008 |
| Stranger Aggression | ENSCAFG00000007235 | *HIPK3* | 0.00013 |
| Stranger Aggression | ENSCAFG00000005175 | *SCEL* | 0.00014 |

| | | | |
|---|---|---|---|
| Stranger Aggression | ENSCAFG00000005746 | *NAV3* | 0.00018 |
| Stranger Aggression | ENSCAFG00000020058 | *SNX7* | 0.00023 |
| Stranger Aggression | ENSCAFG00000000637 | *SLC12A2* | 0.00028 |
| Stranger Aggression | ENSCAFG00000040913 | | 0.00032 |
| Stranger Aggression | ENSCAFG00000006103 | *NEK11* | 0.00035 |
| Stranger Aggression | ENSCAFG00000036701 | | 0.00069 |
| Stranger Aggression | ENSCAFG00000006496 | *MITF* | 0.00070 |
| Stranger Aggression | ENSCAFG00000031547 | *TMED3* | 0.00073 |
| Stranger Aggression | ENSCAFG00000027991 | *RF00026* | 0.00083 |
| Stranger Aggression | ENSCAFG00000010984 | *VTI1A* | 0.00091 |
| Stranger Aggression | ENSCAFG00000018920 | *CLEC16A* | 0.00139 |
| Stranger Aggression | ENSCAFG00000019027 | *LDOC1* | 0.00148 |
| Stranger Aggression | ENSCAFG00000038703 | | 0.00151 |
| Stranger Aggression | ENSCAFG00000038544 | | 0.00166 |
| Stranger Aggression | ENSCAFG00000006263 | | 0.00194 |
| Stranger Aggression | ENSCAFG00000014864 | *CASP12* | 0.00214 |
| Stranger Aggression | ENSCAFG00000001312 | *CHCHD3* | 0.00220 |
| Stranger Aggression | ENSCAFG00000024856 | *KPNA4* | 0.00224 |
| Stranger Aggression | ENSCAFG00000005838 | *PCCA* | 0.00238 |
| Stranger Aggression | ENSCAFG00000027303 | *RF00100* | 0.00272 |
| Stranger Aggression | ENSCAFG00000004428 | *CYSLTR2* | 0.00301 |
| Stranger Aggression | ENSCAFG00000004309 | *SPRYD7* | 0.00321 |
| Stranger Aggression | ENSCAFG00000038361 | | 0.00377 |
| Stranger Aggression | ENSCAFG00000000507 | *VPS13B* | 0.00429 |
| Stranger Aggression | ENSCAFG00000034814 | | 0.00442 |
| Stranger Aggression | ENSCAFG00000026003 | *RF00100* | 0.00478 |
| Stranger Aggression | ENSCAFG00000001286 | *FANCC* | 0.00496 |
| Stranger Aggression | ENSCAFG00000004337 | *EBPL* | 0.00560 |
| Stranger Aggression | ENSCAFG00000014952 | | 0.00652 |

| | | | |
|---|---|---|---|
| Stranger Aggression | ENSCAFG00000017571 | *CACNG3* | 0.00657 |
| Stranger Aggression | ENSCAFG00000021438 | *RF00100* | 0.00718 |
| Stranger Aggression | ENSCAFG00000004764 | *VWA8* | 0.00780 |
| Stranger Aggression | ENSCAFG00000000157 | *DCC* | 0.00822 |
| Stranger Aggression | ENSCAFG00000004795 | *MTRF1* | 0.00887 |
| Stranger Aggression | ENSCAFG00000003860 | *MFSD8* | 0.00927 |
| Stranger Aggression | ENSCAFG00000013966 | *CPS1* | 0.01188 |
| Stranger Aggression | ENSCAFG00000036116 | | 0.01218 |
| Stranger Aggression | ENSCAFG00000014191 | *PBX4* | 0.01394 |
| Stranger Aggression | ENSCAFG00000038556 | | 0.01414 |
| Stranger Aggression | ENSCAFG00000020014 | *GCSH* | 0.01624 |
| Stranger Aggression | ENSCAFG00000010942 | *CCDC91* | 0.01837 |
| Stranger Aggression | ENSCAFG00000033721 | | 0.01937 |
| Stranger Aggression | ENSCAFG00000006431 | *LARS* | 0.01953 |
| Stranger Aggression | ENSCAFG00000036595 | | 0.01960 |
| Stranger Aggression | ENSCAFG00000006879 | *MARCH8* | 0.02336 |
| Stranger Aggression | ENSCAFG00000037402 | | 0.02361 |
| Stranger Aggression | ENSCAFG00000015771 | | 0.02432 |
| Stranger Aggression | ENSCAFG00000024350 | *CENPP* | 0.02775 |
| Stranger Aggression | ENSCAFG00000009506 | *ABI3BP* | 0.03373 |
| Stranger Aggression | ENSCAFG00000017685 | *RIT2* | 0.03393 |
| Stranger Aggression | ENSCAFG00000002335 | | 0.03476 |
| Stranger Aggression | ENSCAFG00000038657 | | 0.03561 |
| Stranger Aggression | ENSCAFG00000002397 | *SHB* | 0.03818 |
| Stranger Aggression | ENSCAFG00000035993 | | 0.04308 |
| Stranger Aggression | ENSCAFG00000007943 | *FAM81B* | 0.04779 |
| Stranger Aggression | ENSCAFG00000035622 | | 0.04838 |
| Stranger Fear | ENSCAFG00000003929 | *COG5* | < 0.00001 |
| Stranger Fear | ENSCAFG00000011082 | *CAMSAP2* | < 0.00001 |

| | | | |
|---|---|---|---|
| Stranger Fear | ENSCAFG00000001434 | *KDM4C* | < 0.00001 |
| Stranger Fear | ENSCAFG00000031429 | *GNA14* | < 0.00001 |
| Stranger Fear | ENSCAFG00000002257 | *ICA1* | < 0.00001 |
| Stranger Fear | ENSCAFG00000014785 | *CEP152* | < 0.00001 |
| Stranger Fear | ENSCAFG00000018024 | *TRAPPC8* | < 0.00001 |
| Stranger Fear | ENSCAFG00000023580 | | < 0.00001 |
| Stranger Fear | ENSCAFG00000036261 | | < 0.00001 |
| Stranger Fear | ENSCAFG00000033485 | | < 0.00001 |
| Stranger Fear | ENSCAFG00000000693 | *ANGPT1* | < 0.00001 |
| Stranger Fear | ENSCAFG00000002667 | | 0.00001 |
| Stranger Fear | ENSCAFG00000006462 | *UBXN8* | 0.00001 |
| Stranger Fear | ENSCAFG00000015131 | *CEP126* | 0.00002 |
| Stranger Fear | ENSCAFG00000015395 | | 0.00002 |
| Stranger Fear | ENSCAFG00000034027 | | 0.00002 |
| Stranger Fear | ENSCAFG00000034629 | | 0.00004 |
| Stranger Fear | ENSCAFG00000013221 | | 0.00030 |
| Stranger Fear | ENSCAFG00000004568 | *CUBN* | 0.00033 |
| Stranger Fear | ENSCAFG00000009280 | *PTPRT* | 0.00038 |
| Stranger Fear | ENSCAFG00000023938 | *GPR158* | 0.00040 |
| Stranger Fear | ENSCAFG00000015679 | *RGS7* | 0.00043 |
| Stranger Fear | ENSCAFG00000003153 | *VPS54* | 0.00045 |
| Stranger Fear | ENSCAFG00000010811 | *LRRC28* | 0.00051 |
| Stranger Fear | ENSCAFG00000035111 | | 0.00055 |
| Stranger Fear | ENSCAFG00000016020 | *ACTB* | 0.00056 |
| Stranger Fear | ENSCAFG00000002546 | *CAMKMT* | 0.00058 |
| Stranger Fear | ENSCAFG00000001988 | *DMRT2* | 0.00074 |
| Stranger Fear | ENSCAFG00000032631 | | 0.00083 |
| Stranger Fear | ENSCAFG00000017227 | *GABRG2* | 0.00084 |
| Stranger Fear | ENSCAFG00000038551 | | 0.00102 |

| | | | |
|---|---|---|---|
| Stranger Fear | ENSCAFG00000035409 | | 0.00107 |
| Stranger Fear | ENSCAFG00000014030 | *LARS2* | 0.00141 |
| Stranger Fear | ENSCAFG00000009159 | *TRIO* | 0.00142 |
| Stranger Fear | ENSCAFG00000013671 | *EFL1* | 0.00150 |
| Stranger Fear | ENSCAFG00000004942 | | 0.00150 |
| Stranger Fear | ENSCAFG00000037968 | | 0.00178 |
| Stranger Fear | ENSCAFG00000015967 | *MTHFD1* | 0.00190 |
| Stranger Fear | ENSCAFG00000019296 | *METTL16* | 0.00235 |
| Stranger Fear | ENSCAFG00000006103 | *NEK11* | 0.00267 |
| Stranger Fear | ENSCAFG00000000337 | *PPM1H* | 0.00307 |
| Stranger Fear | ENSCAFG00000035994 | | 0.00374 |
| Stranger Fear | ENSCAFG00000025104 | *OR51F2* | 0.00431 |
| Stranger Fear | ENSCAFG00000016732 | *SIPA1L1* | 0.00482 |
| Stranger Fear | ENSCAFG00000008669 | *TMEM144* | 0.00589 |
| Stranger Fear | ENSCAFG00000010766 | *TESMIN* | 0.00653 |
| Stranger Fear | ENSCAFG00000006326 | *KCTD16* | 0.00698 |
| Stranger Fear | ENSCAFG00000004408 | *CAB39L* | 0.00702 |
| Stranger Fear | ENSCAFG00000010532 | *SPHKAP* | 0.00781 |
| Stranger Fear | ENSCAFG00000023546 | *TCTN2* | 0.00838 |
| Stranger Fear | ENSCAFG00000002552 | *INVS* | 0.00843 |
| Stranger Fear | ENSCAFG00000039722 | | 0.00963 |
| Stranger Fear | ENSCAFG00000009140 | *GALNT13* | 0.01012 |
| Stranger Fear | ENSCAFG00000000086 | *CDH20* | 0.01050 |
| Stranger Fear | ENSCAFG00000009173 | *EXOC2* | 0.01072 |
| Stranger Fear | ENSCAFG00000007652 | *SPSB4* | 0.01161 |
| Stranger Fear | ENSCAFG00000036396 | | 0.01254 |
| Stranger Fear | ENSCAFG00000037907 | | 0.01350 |
| Stranger Fear | ENSCAFG00000029787 | *SAAL1* | 0.01442 |
| Stranger Fear | ENSCAFG00000010619 | *BICD1* | 0.01495 |

| | | | |
|---|---|---|---|
| Stranger Fear | ENSCAFG00000025134 | | 0.01579 |
| Stranger Fear | ENSCAFG00000008738 | *TM9SF3* | 0.01836 |
| Stranger Fear | ENSCAFG00000035988 | | 0.01932 |
| Stranger Fear | ENSCAFG00000035085 | | 0.02068 |
| Stranger Fear | ENSCAFG00000012632 | *OR51S1* | 0.02356 |
| Stranger Fear | ENSCAFG00000038028 | | 0.02372 |
| Stranger Fear | ENSCAFG00000000696 | *RSPO2* | 0.02879 |
| Stranger Fear | ENSCAFG00000033696 | | 0.03380 |
| Stranger Fear | ENSCAFG00000035420 | | 0.03866 |
| Stranger Fear | ENSCAFG00000035101 | | 0.03982 |
| Stranger Fear | ENSCAFG00000033197 | | 0.04245 |
| Stranger Fear | ENSCAFG00000017412 | | 0.04285 |
| Stranger Fear | ENSCAFG00000017314 | *MBTD1* | 0.04357 |
| Stranger Fear | ENSCAFG00000003582 | *VPS41* | 0.04387 |
| Stranger Fear | ENSCAFG00000018267 | *TOP3A* | 0.04389 |
| Stranger Fear | ENSCAFG00000033416 | | 0.04596 |
| Stranger Fear | ENSCAFG00000014618 | *ARPC2* | 0.04626 |
| Stranger Fear | ENSCAFG00000000974 | *DERL1* | 0.04675 |
| Touch Sensitivity | ENSCAFG00000006013 | *CNTN4* | < 0.00001 |
| Touch Sensitivity | ENSCAFG00000033155 | | < 0.00001 |
| Touch Sensitivity | ENSCAFG00000017362 | *NUP210L* | < 0.00001 |
| Touch Sensitivity | ENSCAFG00000002796 | *CCDC88A* | < 0.00001 |
| Touch Sensitivity | ENSCAFG00000005998 | *LHFPL6* | < 0.00001 |
| Touch Sensitivity | ENSCAFG00000006791 | | < 0.00001 |
| Touch Sensitivity | ENSCAFG00000028659 | | < 0.00001 |
| Touch Sensitivity | ENSCAFG00000001754 | *POT1* | < 0.00001 |
| Touch Sensitivity | ENSCAFG00000000696 | *RSPO2* | < 0.00001 |
| Touch Sensitivity | ENSCAFG00000015625 | *FMN2* | < 0.00001 |
| Touch Sensitivity | ENSCAFG00000000090 | *MC4R* | < 0.00001 |

| | | | |
|---|---|---|---|
| Touch Sensitivity | ENSCAFG00000036746 | | < 0.00001 |
| Touch Sensitivity | ENSCAFG00000002762 | *EML6* | < 0.00001 |
| Touch Sensitivity | ENSCAFG00000020187 | *HFM1* | < 0.00001 |
| Touch Sensitivity | ENSCAFG00000035449 | | < 0.00001 |
| Touch Sensitivity | ENSCAFG00000003479 | | < 0.00001 |
| Touch Sensitivity | ENSCAFG00000033127 | | < 0.00001 |
| Touch Sensitivity | ENSCAFG00000003471 | *PNISR* | < 0.00001 |
| Touch Sensitivity | ENSCAFG00000034355 | | < 0.00001 |
| Touch Sensitivity | ENSCAFG00000005720 | *CLPB* | < 0.00001 |
| Touch Sensitivity | ENSCAFG00000017248 | *UBAP2L* | < 0.00001 |
| Touch Sensitivity | ENSCAFG00000025531 | | < 0.00001 |
| Touch Sensitivity | ENSCAFG00000010355 | *ICE1* | < 0.00001 |
| Touch Sensitivity | ENSCAFG00000004454 | *SUCLA2* | 0.00001 |
| Touch Sensitivity | ENSCAFG00000016568 | *ADAM10* | 0.00002 |
| Touch Sensitivity | ENSCAFG00000002661 | *SMC2* | 0.00002 |
| Touch Sensitivity | ENSCAFG00000007822 | *BDP1* | 0.00004 |
| Touch Sensitivity | ENSCAFG00000005916 | *ANKRD28* | 0.00004 |
| Touch Sensitivity | ENSCAFG00000005775 | *BIRC6* | 0.00005 |
| Touch Sensitivity | ENSCAFG00000008185 | *ARFIP1* | 0.00006 |
| Touch Sensitivity | ENSCAFG00000035409 | | 0.00006 |
| Touch Sensitivity | ENSCAFG00000015877 | *SH2D4B* | 0.00006 |
| Touch Sensitivity | ENSCAFG00000011585 | *PCNX2* | 0.00006 |
| Touch Sensitivity | ENSCAFG00000032632 | *WNT7A* | 0.00007 |
| Touch Sensitivity | ENSCAFG00000017645 | *MYO9A* | 0.00007 |
| Touch Sensitivity | ENSCAFG00000020206 | *SCAI* | 0.00014 |
| Touch Sensitivity | ENSCAFG00000020112 | *ABCD3* | 0.00027 |
| Touch Sensitivity | ENSCAFG00000018665 | *POLDIP2* | 0.00044 |
| Touch Sensitivity | ENSCAFG00000017598 | *SHISA6* | 0.00046 |
| Touch Sensitivity | ENSCAFG00000017651 | *DNM2* | 0.00052 |

| | | | |
|---|---|---|---|
| Touch Sensitivity | ENSCAFG00000033639 | | 0.00054 |
| Touch Sensitivity | ENSCAFG00000018663 | *TNFAIP1* | 0.00055 |
| Touch Sensitivity | ENSCAFG00000006520 | *KCNG3* | 0.00081 |
| Touch Sensitivity | ENSCAFG00000014275 | *SOS2* | 0.00096 |
| Touch Sensitivity | ENSCAFG00000033474 | | 0.00098 |
| Touch Sensitivity | ENSCAFG00000040274 | | 0.00104 |
| Touch Sensitivity | ENSCAFG00000030928 | | 0.00111 |
| Touch Sensitivity | ENSCAFG00000007067 | *NUP58* | 0.00112 |
| Touch Sensitivity | ENSCAFG00000034267 | | 0.00113 |
| Touch Sensitivity | ENSCAFG00000018564 | *GRIA3* | 0.00117 |
| Touch Sensitivity | ENSCAFG00000000737 | *CDC42SE2* | 0.00120 |
| Touch Sensitivity | ENSCAFG00000000884 | *WDR27* | 0.00125 |
| Touch Sensitivity | ENSCAFG00000038140 | | 0.00152 |
| Touch Sensitivity | ENSCAFG00000017606 | *UNC79* | 0.00167 |
| Touch Sensitivity | ENSCAFG00000037655 | | 0.00186 |
| Touch Sensitivity | ENSCAFG00000005600 | *TASP1* | 0.00196 |
| Touch Sensitivity | ENSCAFG00000035826 | | 0.00198 |
| Touch Sensitivity | ENSCAFG00000015383 | *TNFSF10* | 0.00210 |
| Touch Sensitivity | ENSCAFG00000018366 | *KIAA1210* | 0.00336 |
| Touch Sensitivity | ENSCAFG00000034987 | | 0.00414 |
| Touch Sensitivity | ENSCAFG00000037214 | | 0.00419 |
| Touch Sensitivity | ENSCAFG00000023460 | *FBRSL1* | 0.00420 |
| Touch Sensitivity | ENSCAFG00000035962 | | 0.00436 |
| Touch Sensitivity | ENSCAFG00000022906 | *RF00026* | 0.00512 |
| Touch Sensitivity | ENSCAFG00000038034 | | 0.00562 |
| Touch Sensitivity | ENSCAFG00000002753 | *GPNMB* | 0.00571 |
| Touch Sensitivity | ENSCAFG00000013945 | *C1QL1* | 0.00578 |
| Touch Sensitivity | ENSCAFG00000035085 | | 0.00630 |
| Touch Sensitivity | ENSCAFG00000018591 | *ARHGAP28* | 0.00654 |

| | | | |
|---|---|---|---|
| Touch Sensitivity | ENSCAFG00000034887 | | 0.00737 |
| Touch Sensitivity | ENSCAFG00000003705 | *CUL2* | 0.00753 |
| Touch Sensitivity | ENSCAFG00000006730 | *TTC17* | 0.00914 |
| Touch Sensitivity | ENSCAFG00000000828 | *RPS6KA2* | 0.00947 |
| Touch Sensitivity | ENSCAFG00000018890 | *TXNDC11* | 0.00980 |
| Touch Sensitivity | ENSCAFG00000004130 | *NEBL* | 0.01026 |
| Touch Sensitivity | ENSCAFG00000037076 | | 0.01026 |
| Touch Sensitivity | ENSCAFG00000039677 | | 0.01030 |
| Touch Sensitivity | ENSCAFG00000000569 | *ZNF608* | 0.01045 |
| Touch Sensitivity | ENSCAFG00000020387 | *ST6GALNAC3* | 0.01129 |
| Touch Sensitivity | ENSCAFG00000031628 | | 0.01133 |
| Touch Sensitivity | ENSCAFG00000003595 | | 0.01171 |
| Touch Sensitivity | ENSCAFG00000000226 | *HBS1L* | 0.01216 |
| Touch Sensitivity | ENSCAFG00000016310 | *IKZF3* | 0.01265 |
| Touch Sensitivity | ENSCAFG00000038398 | | 0.01311 |
| Touch Sensitivity | ENSCAFG00000037289 | | 0.01363 |
| Touch Sensitivity | ENSCAFG00000012361 | *ASCC2* | 0.01512 |
| Touch Sensitivity | ENSCAFG00000033119 | | 0.01799 |
| Touch Sensitivity | ENSCAFG00000024183 | | 0.01857 |
| Touch Sensitivity | ENSCAFG00000001173 | *FAM135B* | 0.01983 |
| Touch Sensitivity | ENSCAFG00000017197 | *ATP8B2* | 0.02656 |
| Touch Sensitivity | ENSCAFG00000005061 | *LMO7* | 0.02672 |
| Touch Sensitivity | ENSCAFG00000009631 | | 0.02709 |
| Touch Sensitivity | ENSCAFG00000002527 | *LRPPRC* | 0.02976 |
| Touch Sensitivity | ENSCAFG00000030884 | *APOPT1* | 0.02984 |
| Touch Sensitivity | ENSCAFG00000000068 | *BCL2* | 0.03004 |
| Touch Sensitivity | ENSCAFG00000010682 | *SP110* | 0.03069 |
| Touch Sensitivity | ENSCAFG00000005403 | *LRP1B* | 0.03076 |
| Touch Sensitivity | ENSCAFG00000040355 | | 0.03209 |

| | | | |
|---|---|---|---|
| Touch Sensitivity | ENSCAFG00000010897 | *ARHGEF38* | 0.03392 |
| Touch Sensitivity | ENSCAFG00000038632 | | 0.03468 |
| Touch Sensitivity | ENSCAFG00000008744 | *RASGRF2* | 0.03563 |
| Touch Sensitivity | ENSCAFG00000008319 | *CHODL* | 0.03563 |
| Touch Sensitivity | ENSCAFG00000018100 | *SCAPER* | 0.03808 |
| Touch Sensitivity | ENSCAFG00000009227 | *WDR41* | 0.04010 |
| Touch Sensitivity | ENSCAFG00000001700 | *SND1* | 0.04072 |
| Touch Sensitivity | ENSCAFG00000009218 | *MORN4* | 0.04075 |
| Touch Sensitivity | ENSCAFG00000032020 | *TRAPPC3L* | 0.04622 |
| Touch Sensitivity | ENSCAFG00000006050 | *MYO16* | 0.04739 |
| Touch Sensitivity | ENSCAFG00000011657 | *EAF2* | 0.04750 |
| Touch Sensitivity | ENSCAFG00000004249 | *ARHGAP21* | 0.04884 |
| Trainability | ENSCAFG00000018564 | *GRIA3* | < 0.00001 |
| Trainability | ENSCAFG00000000090 | *MC4R* | < 0.00001 |
| Trainability | ENSCAFG00000007420 | *CBFA2T2* | < 0.00001 |
| Trainability | ENSCAFG00000014257 | *PPP2R2C* | < 0.00001 |
| Trainability | ENSCAFG00000018858 | *TARS* | < 0.00001 |
| Trainability | ENSCAFG00000000011 | *NFATC1* | < 0.00001 |
| Trainability | ENSCAFG00000000133 | *WDR7* | < 0.00001 |
| Trainability | ENSCAFG00000029157 | *NOVA1* | < 0.00001 |
| Trainability | ENSCAFG00000035434 | | < 0.00001 |
| Trainability | ENSCAFG00000033089 | | < 0.00001 |
| Trainability | ENSCAFG00000000353 | *STXBP5* | < 0.00001 |
| Trainability | ENSCAFG00000037352 | | < 0.00001 |
| Trainability | ENSCAFG00000018849 | *SNX29* | < 0.00001 |
| Trainability | ENSCAFG00000003373 | *DGKI* | < 0.00001 |
| Trainability | ENSCAFG00000019030 | *ABAT* | < 0.00001 |
| Trainability | ENSCAFG00000000012 | *ATP9B* | < 0.00001 |
| Trainability | ENSCAFG00000004545 | *TRDMT1* | < 0.00001 |

| | | | |
|---|---|---|---|
| Trainability | ENSCAFG00000004157 | *RAB7A* | < 0.00001 |
| Trainability | ENSCAFG00000035215 | | < 0.00001 |
| Trainability | ENSCAFG00000033361 | | < 0.00001 |
| Trainability | ENSCAFG00000003580 | *GRIK2* | < 0.00001 |
| Trainability | ENSCAFG00000031001 | *RF00026* | < 0.00001 |
| Trainability | ENSCAFG00000001700 | *SND1* | < 0.00001 |
| Trainability | ENSCAFG00000009912 | *ERG* | < 0.00001 |
| Trainability | ENSCAFG00000008333 | *TMEM161B* | < 0.00001 |
| Trainability | ENSCAFG00000027467 | *RF00026* | < 0.00001 |
| Trainability | ENSCAFG00000038771 | | < 0.00001 |
| Trainability | ENSCAFG00000030656 | | < 0.00001 |
| Trainability | ENSCAFG00000000522 | *SNCAIP* | < 0.00001 |
| Trainability | ENSCAFG00000039318 | | < 0.00001 |
| Trainability | ENSCAFG00000038087 | | < 0.00001 |
| Trainability | ENSCAFG00000034669 | | < 0.00001 |
| Trainability | ENSCAFG00000035520 | | < 0.00001 |
| Trainability | ENSCAFG00000028239 | *RF00003* | < 0.00001 |
| Trainability | ENSCAFG00000035311 | | < 0.00001 |
| Trainability | ENSCAFG00000008211 | *CACNA2D3* | 0.00001 |
| Trainability | ENSCAFG00000039561 | | 0.00001 |
| Trainability | ENSCAFG00000028601 | | 0.00001 |
| Trainability | ENSCAFG00000028075 | | 0.00001 |
| Trainability | ENSCAFG00000018972 | *EFCAB5* | 0.00001 |
| Trainability | ENSCAFG00000012729 | *CGN* | 0.00001 |
| Trainability | ENSCAFG00000015351 | *BAIAP2L1* | 0.00001 |
| Trainability | ENSCAFG00000011725 | *ATRNL1* | 0.00002 |
| Trainability | ENSCAFG00000008074 | | 0.00002 |
| Trainability | ENSCAFG00000032608 | *LURAP1L* | 0.00002 |
| Trainability | ENSCAFG00000033559 | | 0.00002 |

| | | | |
|---|---|---|---|
| Trainability | ENSCAFG00000013852 | *MATN2* | 0.00002 |
| Trainability | ENSCAFG00000000582 | *GRAMD2B* | 0.00002 |
| Trainability | ENSCAFG00000010692 | *USF3* | 0.00002 |
| Trainability | ENSCAFG00000028708 | *SCD5* | 0.00002 |
| Trainability | ENSCAFG00000000807 | *TBXT* | 0.00003 |
| Trainability | ENSCAFG00000006791 | | 0.00003 |
| Trainability | ENSCAFG00000009081 | *EPHA6* | 0.00004 |
| Trainability | ENSCAFG00000036869 | | 0.00007 |
| Trainability | ENSCAFG00000033886 | | 0.00008 |
| Trainability | ENSCAFG00000007837 | *KNTC1* | 0.00010 |
| Trainability | ENSCAFG00000005949 | *HACL1* | 0.00010 |
| Trainability | ENSCAFG00000014096 | | 0.00011 |
| Trainability | ENSCAFG00000034355 | | 0.00019 |
| Trainability | ENSCAFG00000008199 | *FMN1* | 0.00023 |
| Trainability | ENSCAFG00000005056 | *SCN10A* | 0.00027 |
| Trainability | ENSCAFG00000000745 | *TBC1D22A* | 0.00030 |
| Trainability | ENSCAFG00000032743 | | 0.00030 |
| Trainability | ENSCAFG00000027188 | *RF00026* | 0.00031 |
| Trainability | ENSCAFG00000006017 | *EAF1* | 0.00035 |
| Trainability | ENSCAFG00000034432 | | 0.00035 |
| Trainability | ENSCAFG00000009052 | *LIN54* | 0.00045 |
| Trainability | ENSCAFG00000038280 | | 0.00048 |
| Trainability | ENSCAFG00000001116 | *SAMD3* | 0.00068 |
| Trainability | ENSCAFG00000039328 | | 0.00072 |
| Trainability | ENSCAFG00000019042 | *METTL22* | 0.00078 |
| Trainability | ENSCAFG00000005849 | *TBC1D5* | 0.00087 |
| Trainability | ENSCAFG00000011266 | *TDRD1* | 0.00091 |
| Trainability | ENSCAFG00000002959 | | 0.00097 |
| Trainability | ENSCAFG00000015877 | *SH2D4B* | 0.00117 |

| | | | |
|---|---|---|---|
| Trainability | ENSCAFG00000031346 | *RHOU* | 0.00120 |
| Trainability | ENSCAFG00000002364 | *ADGRL3* | 0.00122 |
| Trainability | ENSCAFG00000003651 | *CSMD2* | 0.00132 |
| Trainability | ENSCAFG00000033071 | | 0.00165 |
| Trainability | ENSCAFG00000010827 | *TTC23* | 0.00217 |
| Trainability | ENSCAFG00000014755 | *SAMD7* | 0.00231 |
| Trainability | ENSCAFG00000027730 | *RF00026* | 0.00288 |
| Trainability | ENSCAFG00000030938 | | 0.00322 |
| Trainability | ENSCAFG00000030561 | | 0.00327 |
| Trainability | ENSCAFG00000014358 | *CASK* | 0.00352 |
| Trainability | ENSCAFG00000037144 | | 0.00368 |
| Trainability | ENSCAFG00000006989 | *DEFB119* | 0.00378 |
| Trainability | ENSCAFG00000005326 | *UVRAG* | 0.00399 |
| Trainability | ENSCAFG00000038450 | | 0.00458 |
| Trainability | ENSCAFG00000017514 | *PCDH19* | 0.00460 |
| Trainability | ENSCAFG00000025531 | | 0.00482 |
| Trainability | ENSCAFG00000000855 | *IL5* | 0.00486 |
| Trainability | ENSCAFG00000006046 | *COL6A5* | 0.00507 |
| Trainability | ENSCAFG00000001425 | *GKAP1* | 0.00516 |
| Trainability | ENSCAFG00000035552 | | 0.00600 |
| Trainability | ENSCAFG00000012544 | *COPA* | 0.00633 |
| Trainability | ENSCAFG00000039541 | | 0.00642 |
| Trainability | ENSCAFG00000015742 | | 0.00651 |
| Trainability | ENSCAFG00000013288 | *ADAMTSL3* | 0.00727 |
| Trainability | ENSCAFG00000008275 | *LYSMD3* | 0.00742 |
| Trainability | ENSCAFG00000034093 | | 0.00775 |
| Trainability | ENSCAFG00000001544 | *EIF3D* | 0.00894 |
| Trainability | ENSCAFG00000039816 | | 0.01100 |
| Trainability | ENSCAFG00000009991 | | 0.01154 |

| | | | |
|---|---|---|---|
| Trainability | ENSCAFG00000001927 | *TMEM252* | 0.01223 |
| Trainability | ENSCAFG00000008790 | *CTNNBL1* | 0.01225 |
| Trainability | ENSCAFG00000002025 | *GCC2* | 0.01320 |
| Trainability | ENSCAFG00000039642 | | 0.01363 |
| Trainability | ENSCAFG00000011247 | *LAPTM5* | 0.01408 |
| Trainability | ENSCAFG00000039734 | | 0.01581 |
| Trainability | ENSCAFG00000002023 | *RCAN2* | 0.01643 |
| Trainability | ENSCAFG00000008851 | *HECTD4* | 0.01729 |
| Trainability | ENSCAFG00000016024 | *FBXL18* | 0.01755 |
| Trainability | ENSCAFG00000038274 | | 0.01819 |
| Trainability | ENSCAFG00000010235 | *ARHGAP32* | 0.01851 |
| Trainability | ENSCAFG00000018267 | *TOP3A* | 0.02119 |
| Trainability | ENSCAFG00000011571 | *SOX5* | 0.02210 |
| Trainability | ENSCAFG00000025514 | *C28H10orf53* | 0.02239 |
| Trainability | ENSCAFG00000020330 | *ADGRL2* | 0.02296 |
| Trainability | ENSCAFG00000011962 | *RFC2* | 0.02385 |
| Trainability | ENSCAFG00000005839 | *SATB1* | 0.02594 |
| Trainability | ENSCAFG00000011796 | *MCF2L2* | 0.02601 |
| Trainability | ENSCAFG00000011940 | *PARP9* | 0.02610 |
| Trainability | ENSCAFG00000035103 | | 0.02730 |
| Trainability | ENSCAFG00000010996 | *CCDC15* | 0.02813 |
| Trainability | ENSCAFG00000009595 | *PKD2* | 0.02864 |
| Trainability | ENSCAFG00000005180 | *MRPL33* | 0.02908 |
| Trainability | ENSCAFG00000005090 | *AANAT* | 0.03065 |
| Trainability | ENSCAFG00000013218 | *EHHADH* | 0.03115 |
| Trainability | ENSCAFG00000038230 | | 0.03131 |
| Trainability | ENSCAFG00000016413 | *GNAI1* | 0.03170 |
| Trainability | ENSCAFG00000040090 | | 0.03245 |
| Trainability | ENSCAFG00000023322 | *RASSF9* | 0.03347 |

| | | | |
|---|---|---|---|
| Trainability | ENSCAFG00000014001 | *TOGARAM1* | 0.03395 |
| Trainability | ENSCAFG00000000507 | *VPS13B* | 0.03450 |
| Trainability | ENSCAFG00000010120 | *RAD54L2* | 0.03732 |
| Trainability | ENSCAFG00000017137 | *TENM2* | 0.04007 |
| Trainability | ENSCAFG00000023015 | *SERPINB12* | 0.04145 |
| Trainability | ENSCAFG00000014045 | *LRRC20* | 0.04241 |
| Trainability | ENSCAFG00000014441 | *DNAJB12* | 0.04261 |
| Trainability | ENSCAFG00000034449 | | 0.04618 |
| Trainability | ENSCAFG00000016511 | *RFX1* | 0.04758 |
| Trainability | ENSCAFG00000015716 | *RAC1* | 0.04785 |
| Trainability | ENSCAFG00000000867 | *RAD50* | 0.04893 |

Table S8. Significant Gene Ontology (GO) terms from enrichment analyses mapping SNPs to the nearest gene within 20kb to derive gene-level p values (meta-analysis, Fisher's method).

| *Behavioral Trait* | *GO ID* | *Term* | *p value* |
|---|---|---|---|
| Attach & Atn Seeking | GO:0060047 | heart contraction | 0.00051 |
| Attach & Atn Seeking | GO:0090382 | phagosome maturation | 0.00089 |
| Attach & Atn Seeking | GO:0051491 | positive regulation of filopodium assemb... | 0.00123 |
| Attach & Atn Seeking | GO:0015872 | dopamine transport | 0.00138 |
| Attach & Atn Seeking | GO:0048172 | regulation of short-term neuronal synapt... | 0.00139 |
| Attach & Atn Seeking | GO:0099531 | presynaptic process involved in chemical... | 0.00276 |
| Attach & Atn Seeking | GO:0014065 | phosphatidylinositol 3-kinase signaling | 0.00367 |
| Attach & Atn Seeking | GO:0046621 | negative regulation of organ growth | 0.00416 |
| Attach & Atn Seeking | GO:0060831 | smoothened signaling pathway involved in... | 0.00416 |
| Attach & Atn Seeking | GO:0007040 | lysosome organization | 0.00454 |
| Attach & Atn Seeking | GO:0090162 | establishment of epithelial cell polarit... | 0.00558 |
| Attach & Atn Seeking | GO:0008104 | protein localization | 0.00680 |
| Attach & Atn Seeking | GO:0033173 | calcineurin-NFAT signaling cascade | 0.00847 |
| Attach & Atn Seeking | GO:0042220 | response to cocaine | 0.00852 |
| Attach & Atn Seeking | GO:0031338 | regulation of vesicle fusion | 0.01012 |
| Attach & Atn Seeking | GO:0032088 | negative regulation of NF-kappaB transcr... | 0.01213 |
| Attach & Atn Seeking | GO:0031589 | cell-substrate adhesion | 0.01433 |
| Attach & Atn Seeking | GO:0002093 | auditory receptor cell morphogenesis | 0.01438 |
| Attach & Atn Seeking | GO:0032367 | intracellular cholesterol transport | 0.01438 |
| Attach & Atn Seeking | GO:0048678 | response to axon injury | 0.01638 |
| Attach & Atn Seeking | GO:0007178 | transmembrane receptor protein serine/th... | 0.01641 |
| Attach & Atn Seeking | GO:0045992 | negative regulation of embryonic develop... | 0.01742 |
| Attach & Atn Seeking | GO:0032648 | regulation of interferon-beta production | 0.01889 |
| Attach & Atn Seeking | GO:0021984 | adenohypophysis development | 0.01898 |
| Attach & Atn Seeking | GO:0034311 | diol metabolic process | 0.01898 |
| Attach & Atn Seeking | GO:0008608 | attachment of spindle microtubules to ki... | 0.02086 |

| | | | |
|---|---|---|---|
| Attach & Atn Seeking | GO:0030513 | positive regulation of BMP signaling pat... | 0.02086 |
| Attach & Atn Seeking | GO:0071634 | regulation of transforming growth factor... | 0.02431 |
| Attach & Atn Seeking | GO:1903861 | positive regulation of dendrite extensio... | 0.02431 |
| Attach & Atn Seeking | GO:2000678 | negative regulation of transcription reg... | 0.02431 |
| Attach & Atn Seeking | GO:0060740 | prostate gland epithelium morphogenesis | 0.02469 |
| Attach & Atn Seeking | GO:0031032 | actomyosin structure organization | 0.02473 |
| Attach & Atn Seeking | GO:1903532 | positive regulation of secretion by cell | 0.02611 |
| Attach & Atn Seeking | GO:0051588 | regulation of neurotransmitter transport | 0.02625 |
| Attach & Atn Seeking | GO:0071322 | cellular response to carbohydrate stimul... | 0.02643 |
| Attach & Atn Seeking | GO:0008344 | adult locomotory behavior | 0.02787 |
| Attach & Atn Seeking | GO:0098742 | cell-cell adhesion via plasma-membrane a... | 0.02930 |
| Attach & Atn Seeking | GO:0003198 | epithelial to mesenchymal transition inv... | 0.03035 |
| Attach & Atn Seeking | GO:0090136 | epithelial cell-cell adhesion | 0.03035 |
| Attach & Atn Seeking | GO:0001893 | maternal placenta development | 0.03711 |
| Attach & Atn Seeking | GO:0032506 | cytokinetic process | 0.03711 |
| Attach & Atn Seeking | GO:0050687 | negative regulation of defense response ... | 0.03711 |
| Attach & Atn Seeking | GO:0060390 | regulation of SMAD protein signal transd... | 0.03711 |
| Attach & Atn Seeking | GO:0071467 | cellular response to pH | 0.03711 |
| Attach & Atn Seeking | GO:0042692 | muscle cell differentiation | 0.03784 |
| Attach & Atn Seeking | GO:0010611 | regulation of cardiac muscle hypertrophy | 0.03805 |
| Attach & Atn Seeking | GO:1903363 | negative regulation of cellular protein ... | 0.03809 |
| Attach & Atn Seeking | GO:0090559 | regulation of membrane permeability | 0.03826 |
| Attach & Atn Seeking | GO:0034198 | cellular response to amino acid starvati... | 0.04402 |
| Attach & Atn Seeking | GO:0042073 | intraciliary transport | 0.04402 |
| Attach & Atn Seeking | GO:0048675 | axon extension | 0.04444 |
| Attach & Atn Seeking | GO:0042474 | middle ear morphogenesis | 0.04457 |
| Attach & Atn Seeking | GO:0048488 | synaptic vesicle endocytosis | 0.04457 |
| Attach & Atn Seeking | GO:0048670 | regulation of collateral sprouting | 0.04457 |
| Attach & Atn Seeking | GO:0031175 | neuron projection development | 0.04898 |

| | | | |
|---|---|---|---|
| Attach & Atn Seeking | GO:0044271 | cellular nitrogen compound biosynthetic ... | 0.04912 |
| Chasing | GO:0060047 | heart contraction | 0.00016 |
| Chasing | GO:0050919 | negative chemotaxis | 0.00019 |
| Chasing | GO:0043547 | positive regulation of GTPase activity | 0.00056 |
| Chasing | GO:0008360 | regulation of cell shape | 0.00064 |
| Chasing | GO:0006887 | exocytosis | 0.00077 |
| Chasing | GO:0045197 | establishment or maintenance of epitheli... | 0.00136 |
| Chasing | GO:0008045 | motor neuron axon guidance | 0.00165 |
| Chasing | GO:0035335 | peptidyl-tyrosine dephosphorylation | 0.00270 |
| Chasing | GO:0034755 | iron ion transmembrane transport | 0.00386 |
| Chasing | GO:0060390 | regulation of SMAD protein signal transd... | 0.00386 |
| Chasing | GO:0050730 | regulation of peptidyl-tyrosine phosphor... | 0.00388 |
| Chasing | GO:0010646 | regulation of cell communication | 0.00616 |
| Chasing | GO:0051056 | regulation of small GTPase mediated sign... | 0.00618 |
| Chasing | GO:0006886 | intracellular protein transport | 0.00619 |
| Chasing | GO:0030859 | polarized epithelial cell differentiatio... | 0.00623 |
| Chasing | GO:0045603 | positive regulation of endothelial cell ... | 0.00623 |
| Chasing | GO:0050910 | detection of mechanical stimulus involve... | 0.00623 |
| Chasing | GO:0051299 | centrosome separation | 0.00623 |
| Chasing | GO:0061339 | establishment or maintenance of monopola... | 0.00623 |
| Chasing | GO:0003013 | circulatory system process | 0.00653 |
| Chasing | GO:2001237 | negative regulation of extrinsic apoptot... | 0.00696 |
| Chasing | GO:0001759 | organ induction | 0.00734 |
| Chasing | GO:0071526 | semaphorin-plexin signaling pathway | 0.00734 |
| Chasing | GO:0051279 | regulation of release of sequestered cal... | 0.00735 |
| Chasing | GO:0048813 | dendrite morphogenesis | 0.00739 |
| Chasing | GO:0040011 | locomotion | 0.00820 |
| Chasing | GO:0010506 | regulation of autophagy | 0.00865 |
| Chasing | GO:1903146 | regulation of autophagy of mitochondrion | 0.00898 |

| | | | |
|---|---|---|---|
| Chasing | GO:0010633 | negative regulation of epithelial cell m... | 0.00915 |
| Chasing | GO:0072178 | nephric duct morphogenesis | 0.00916 |
| Chasing | GO:0030324 | lung development | 0.01078 |
| Chasing | GO:0000186 | activation of MAPKK activity | 0.01097 |
| Chasing | GO:0050922 | negative regulation of chemotaxis | 0.01281 |
| Chasing | GO:0048522 | positive regulation of cellular process | 0.01282 |
| Chasing | GO:0000466 | maturation of 5.8S rRNA from tricistroni... | 0.01287 |
| Chasing | GO:0021795 | cerebral cortex cell migration | 0.01287 |
| Chasing | GO:0061154 | endothelial tube morphogenesis | 0.01287 |
| Chasing | GO:0010721 | negative regulation of cell development | 0.01294 |
| Chasing | GO:0010595 | positive regulation of endothelial cell ... | 0.01327 |
| Chasing | GO:0008219 | cell death | 0.01461 |
| Chasing | GO:1903169 | regulation of calcium ion transmembrane ... | 0.01466 |
| Chasing | GO:0007044 | cell-substrate junction assembly | 0.01478 |
| Chasing | GO:0050850 | positive regulation of calcium-mediated ... | 0.01592 |
| Chasing | GO:1901998 | toxin transport | 0.01630 |
| Chasing | GO:0006929 | substrate-dependent cell migration | 0.01741 |
| Chasing | GO:0007413 | axonal fasciculation | 0.01741 |
| Chasing | GO:0036035 | osteoclast development | 0.01741 |
| Chasing | GO:0046931 | pore complex assembly | 0.01741 |
| Chasing | GO:0061097 | regulation of protein tyrosine kinase ac... | 0.01873 |
| Chasing | GO:0034329 | cell junction assembly | 0.01875 |
| Chasing | GO:0072283 | metanephric renal vesicle morphogenesis | 0.01883 |
| Chasing | GO:1902667 | regulation of axon guidance | 0.01883 |
| Chasing | GO:0031345 | negative regulation of cell projection o... | 0.01886 |
| Chasing | GO:2000696 | regulation of epithelial cell differenti... | 0.01887 |
| Chasing | GO:0046636 | negative regulation of alpha-beta T cell... | 0.01889 |
| Chasing | GO:0006475 | internal protein amino acid acetylation | 0.01902 |
| Chasing | GO:0006487 | protein N-linked glycosylation | 0.01929 |

| | | | |
|---|---|---|---|
| Chasing | GO:0010811 | positive regulation of cell-substrate ad... | 0.01976 |
| Chasing | GO:0007052 | mitotic spindle organization | 0.02276 |
| Chasing | GO:0032506 | cytokinetic process | 0.02283 |
| Chasing | GO:0061298 | retina vasculature development in camera... | 0.02283 |
| Chasing | GO:0090162 | establishment of epithelial cell polarit... | 0.02283 |
| Chasing | GO:0097286 | iron ion import | 0.02283 |
| Chasing | GO:0009948 | anterior/posterior axis specification | 0.02322 |
| Chasing | GO:0050772 | positive regulation of axonogenesis | 0.02416 |
| Chasing | GO:0001960 | negative regulation of cytokine-mediated... | 0.02430 |
| Chasing | GO:0001662 | behavioral fear response | 0.02433 |
| Chasing | GO:0007165 | signal transduction | 0.02607 |
| Chasing | GO:0001764 | neuron migration | 0.02821 |
| Chasing | GO:0000132 | establishment of mitotic spindle orienta... | 0.02916 |
| Chasing | GO:0002026 | regulation of the force of heart contrac... | 0.02916 |
| Chasing | GO:0051150 | regulation of smooth muscle cell differe... | 0.02916 |
| Chasing | GO:0019318 | hexose metabolic process | 0.02922 |
| Chasing | GO:0032467 | positive regulation of cytokinesis | 0.02942 |
| Chasing | GO:0051962 | positive regulation of nervous system de... | 0.03072 |
| Chasing | GO:0055074 | calcium ion homeostasis | 0.03099 |
| Chasing | GO:0046677 | response to antibiotic | 0.03133 |
| Chasing | GO:0019637 | organophosphate metabolic process | 0.03136 |
| Chasing | GO:0070972 | protein localization to endoplasmic reti... | 0.03138 |
| Chasing | GO:0046854 | phosphatidylinositol phosphorylation | 0.03186 |
| Chasing | GO:0061098 | positive regulation of protein tyrosine ... | 0.03186 |
| Chasing | GO:0070374 | positive regulation of ERK1 and ERK2 cas... | 0.03229 |
| Chasing | GO:0006906 | vesicle fusion | 0.03267 |
| Chasing | GO:0009593 | detection of chemical stimulus | 0.03277 |
| Chasing | GO:0010469 | regulation of signaling receptor activit... | 0.03385 |
| Chasing | GO:0007169 | transmembrane receptor protein tyrosine ... | 0.03488 |

| | | | |
|---|---|---|---|
| Chasing | GO:0006508 | proteolysis | 0.03502 |
| Chasing | GO:0001755 | neural crest cell migration | 0.03515 |
| Chasing | GO:0007265 | Ras protein signal transduction | 0.03527 |
| Chasing | GO:0048846 | axon extension involved in axon guidance | 0.03563 |
| Chasing | GO:0140053 | mitochondrial gene expression | 0.03565 |
| Chasing | GO:0035767 | endothelial cell chemotaxis | 0.03571 |
| Chasing | GO:0055017 | cardiac muscle tissue growth | 0.03577 |
| Chasing | GO:0090100 | positive regulation of transmembrane rec... | 0.03577 |
| Chasing | GO:1905475 | regulation of protein localization to me... | 0.03579 |
| Chasing | GO:0061614 | pri-miRNA transcription by RNA polymeras... | 0.03580 |
| Chasing | GO:0070665 | positive regulation of leukocyte prolife... | 0.03602 |
| Chasing | GO:0033157 | regulation of intracellular protein tran... | 0.03606 |
| Chasing | GO:0030500 | regulation of bone mineralization | 0.03618 |
| Chasing | GO:0048278 | vesicle docking | 0.03625 |
| Chasing | GO:0030517 | negative regulation of axon extension | 0.03644 |
| Chasing | GO:0031639 | plasminogen activation | 0.03644 |
| Chasing | GO:0007411 | axon guidance | 0.04196 |
| Chasing | GO:0045124 | regulation of bone resorption | 0.04220 |
| Chasing | GO:0045104 | intermediate filament cytoskeleton organ... | 0.04222 |
| Chasing | GO:0002070 | epithelial cell maturation | 0.04223 |
| Chasing | GO:0007183 | SMAD protein complex assembly | 0.04223 |
| Chasing | GO:0007350 | blastoderm segmentation | 0.04223 |
| Chasing | GO:0007512 | adult heart development | 0.04223 |
| Chasing | GO:0032352 | positive regulation of hormone metabolic... | 0.04223 |
| Chasing | GO:0043923 | positive regulation by host of viral tra... | 0.04223 |
| Chasing | GO:0048172 | regulation of short-term neuronal synapt... | 0.04223 |
| Chasing | GO:0051654 | establishment of mitochondrion localizat... | 0.04223 |
| Chasing | GO:0071711 | basement membrane organization | 0.04223 |
| Chasing | GO:0072425 | signal transduction involved in G2 DNA d... | 0.04223 |

| | | | |
|---|---|---|---|
| Chasing | GO:0090148 | membrane fission | 0.04223 |
| Chasing | GO:2000316 | regulation of T-helper 17 type immune re... | 0.04223 |
| Chasing | GO:0045471 | response to ethanol | 0.04226 |
| Chasing | GO:0060065 | uterus development | 0.04466 |
| Chasing | GO:0060602 | branch elongation of an epithelium | 0.04466 |
| Chasing | GO:0010976 | positive regulation of neuron projection... | 0.04516 |
| Chasing | GO:0030218 | erythrocyte differentiation | 0.04845 |
| Chasing | GO:0043113 | receptor clustering | 0.04853 |
| Chasing | GO:0050918 | positive chemotaxis | 0.04853 |
| Dog Aggression | GO:0043393 | regulation of protein binding | 0.00110 |
| Dog Aggression | GO:0000226 | microtubule cytoskeleton organization | 0.00150 |
| Dog Aggression | GO:0030336 | negative regulation of cell migration | 0.00150 |
| Dog Aggression | GO:0045687 | positive regulation of glial cell differ... | 0.00200 |
| Dog Aggression | GO:0018279 | protein N-linked glycosylation via aspar... | 0.00270 |
| Dog Aggression | GO:0072384 | organelle transport along microtubule | 0.00270 |
| Dog Aggression | GO:0010171 | body morphogenesis | 0.00320 |
| Dog Aggression | GO:0034333 | adherens junction assembly | 0.00320 |
| Dog Aggression | GO:0051383 | kinetochore organization | 0.00350 |
| Dog Aggression | GO:0050680 | negative regulation of epithelial cell p... | 0.00390 |
| Dog Aggression | GO:1902905 | positive regulation of supramolecular fi... | 0.00650 |
| Dog Aggression | GO:2001259 | positive regulation of cation channel ac... | 0.00680 |
| Dog Aggression | GO:0043170 | macromolecule metabolic process | 0.00840 |
| Dog Aggression | GO:0035315 | hair cell differentiation | 0.00920 |
| Dog Aggression | GO:0070936 | protein K48-linked ubiquitination | 0.00960 |
| Dog Aggression | GO:0050709 | negative regulation of protein secretion | 0.01170 |
| Dog Aggression | GO:0001541 | ovarian follicle development | 0.01190 |
| Dog Aggression | GO:0045747 | positive regulation of Notch signaling p... | 0.01190 |
| Dog Aggression | GO:0006890 | retrograde vesicle-mediated transport, G... | 0.01340 |
| Dog Aggression | GO:0048713 | regulation of oligodendrocyte differenti... | 0.01340 |

| | | | |
|---|---|---|---|
| Dog Aggression | GO:0007049 | cell cycle | 0.01600 |
| Dog Aggression | GO:0003157 | endocardium development | 0.01610 |
| Dog Aggression | GO:0003176 | aortic valve development | 0.01610 |
| Dog Aggression | GO:0009110 | vitamin biosynthetic process | 0.01610 |
| Dog Aggression | GO:0010832 | negative regulation of myotube different... | 0.01610 |
| Dog Aggression | GO:0045618 | positive regulation of keratinocyte diff... | 0.01610 |
| Dog Aggression | GO:0046596 | regulation of viral entry into host cell | 0.01610 |
| Dog Aggression | GO:0050910 | detection of mechanical stimulus involve... | 0.01610 |
| Dog Aggression | GO:0051497 | negative regulation of stress fiber asse... | 0.01610 |
| Dog Aggression | GO:0051954 | positive regulation of amine transport | 0.01610 |
| Dog Aggression | GO:0090051 | negative regulation of cell migration in... | 0.01610 |
| Dog Aggression | GO:1901018 | positive regulation of potassium ion tra... | 0.01610 |
| Dog Aggression | GO:0007399 | nervous system development | 0.01650 |
| Dog Aggression | GO:0038127 | ERBB signaling pathway | 0.01770 |
| Dog Aggression | GO:0032774 | RNA biosynthetic process | 0.01810 |
| Dog Aggression | GO:0031648 | protein destabilization | 0.01910 |
| Dog Aggression | GO:0045724 | positive regulation of cilium assembly | 0.02000 |
| Dog Aggression | GO:0045737 | positive regulation of cyclin-dependent ... | 0.02000 |
| Dog Aggression | GO:0034138 | toll-like receptor 3 signaling pathway | 0.02120 |
| Dog Aggression | GO:0039529 | RIG-I signaling pathway | 0.02120 |
| Dog Aggression | GO:0006874 | cellular calcium ion homeostasis | 0.02390 |
| Dog Aggression | GO:0030316 | osteoclast differentiation | 0.02390 |
| Dog Aggression | GO:0055010 | ventricular cardiac muscle tissue morpho... | 0.02810 |
| Dog Aggression | GO:0042177 | negative regulation of protein catabolic... | 0.02830 |
| Dog Aggression | GO:0048013 | ephrin receptor signaling pathway | 0.02830 |
| Dog Aggression | GO:0051057 | positive regulation of small GTPase medi... | 0.02840 |
| Dog Aggression | GO:0003015 | heart process | 0.02850 |
| Dog Aggression | GO:0032392 | DNA geometric change | 0.02850 |
| Dog Aggression | GO:1903902 | positive regulation of viral life cycle | 0.02850 |

| | | | |
|---|---|---|---|
| Dog Aggression | GO:0000186 | activation of MAPKK activity | 0.02870 |
| Dog Aggression | GO:0035249 | synaptic transmission, glutamatergic | 0.03360 |
| Dog Aggression | GO:0007188 | adenylate cyclase-modulating G-protein c... | 0.03370 |
| Dog Aggression | GO:0048169 | regulation of long-term neuronal synapti... | 0.03380 |
| Dog Aggression | GO:0070979 | protein K11-linked ubiquitination | 0.03380 |
| Dog Aggression | GO:0007265 | Ras protein signal transduction | 0.03760 |
| Dog Aggression | GO:0045880 | positive regulation of smoothened signal... | 0.03830 |
| Dog Aggression | GO:0007050 | cell cycle arrest | 0.03990 |
| Dog Aggression | GO:0010799 | regulation of peptidyl-threonine phospho... | 0.04100 |
| Dog Aggression | GO:0071826 | ribonucleoprotein complex subunit organi... | 0.04100 |
| Dog Aggression | GO:0060560 | developmental growth involved in morphog... | 0.04110 |
| Dog Aggression | GO:0043297 | apical junction assembly | 0.04120 |
| Dog Aggression | GO:0001502 | cartilage condensation | 0.04130 |
| Dog Aggression | GO:0006029 | proteoglycan metabolic process | 0.04130 |
| Dog Aggression | GO:0060292 | long term synaptic depression | 0.04130 |
| Dog Aggression | GO:1901021 | positive regulation of calcium ion trans... | 0.04130 |
| Dog Aggression | GO:0021987 | cerebral cortex development | 0.04540 |
| Dog Aggression | GO:0007157 | heterophilic cell-cell adhesion via plas... | 0.04560 |
| Dog Aggression | GO:0007030 | Golgi organization | 0.04670 |
| Dog Aggression | GO:0006468 | protein phosphorylation | 0.04800 |
| Dog Aggression | GO:0007093 | mitotic cell cycle checkpoint | 0.04930 |
| Dog Aggression | GO:0048593 | camera-type eye morphogenesis | 0.04930 |
| Dog Aggression | GO:0046887 | positive regulation of hormone secretion | 0.04940 |
| Dog Aggression | GO:0033962 | cytoplasmic mRNA processing body assembl... | 0.04950 |
| Dog Aggression | GO:0060045 | positive regulation of cardiac muscle ce... | 0.04950 |
| Dog Aggression | GO:0030048 | actin filament-based movement | 0.04960 |
| Dog Aggression | GO:0030177 | positive regulation of Wnt signaling pat... | 0.04960 |
| Dog Aggression | GO:0042035 | regulation of cytokine biosynthetic proc... | 0.04960 |
| Dog Aggression | GO:0051402 | neuron apoptotic process | 0.04970 |

| | | | |
|---|---|---|---|
| Dog Fear | GO:0006082 | organic acid metabolic process | 0.00110 |
| Dog Fear | GO:0034138 | toll-like receptor 3 signaling pathway | 0.00170 |
| Dog Fear | GO:0050775 | positive regulation of dendrite morphoge... | 0.00250 |
| Dog Fear | GO:0016042 | lipid catabolic process | 0.00330 |
| Dog Fear | GO:0035556 | intracellular signal transduction | 0.00350 |
| Dog Fear | GO:0048013 | ephrin receptor signaling pathway | 0.00370 |
| Dog Fear | GO:0072521 | purine-containing compound metabolic pro... | 0.00420 |
| Dog Fear | GO:0045022 | early endosome to late endosome transpor... | 0.00450 |
| Dog Fear | GO:0060390 | regulation of SMAD protein signal transd... | 0.00460 |
| Dog Fear | GO:0010639 | negative regulation of organelle organiz... | 0.00760 |
| Dog Fear | GO:0035265 | organ growth | 0.00760 |
| Dog Fear | GO:0045923 | positive regulation of fatty acid metabo... | 0.00770 |
| Dog Fear | GO:0042531 | positive regulation of tyrosine phosphor... | 0.00800 |
| Dog Fear | GO:0016358 | dendrite development | 0.00920 |
| Dog Fear | GO:0014068 | positive regulation of phosphatidylinosi... | 0.00950 |
| Dog Fear | GO:0035418 | protein localization to synapse | 0.00970 |
| Dog Fear | GO:0050658 | RNA transport | 0.01250 |
| Dog Fear | GO:2001224 | positive regulation of neuron migration | 0.01250 |
| Dog Fear | GO:0048870 | cell motility | 0.01450 |
| Dog Fear | GO:0051098 | regulation of binding | 0.01460 |
| Dog Fear | GO:0010324 | membrane invagination | 0.01480 |
| Dog Fear | GO:0034248 | regulation of cellular amide metabolic p... | 0.01480 |
| Dog Fear | GO:0090100 | positive regulation of transmembrane rec... | 0.01480 |
| Dog Fear | GO:0045727 | positive regulation of translation | 0.01500 |
| Dog Fear | GO:0032502 | developmental process | 0.01610 |
| Dog Fear | GO:0003351 | epithelial cilium movement | 0.01650 |
| Dog Fear | GO:0032594 | protein transport within lipid bilayer | 0.01650 |
| Dog Fear | GO:0048820 | hair follicle maturation | 0.01650 |
| Dog Fear | GO:0060548 | negative regulation of cell death | 0.01720 |

| | | | |
|---|---|---|---|
| Dog Fear | GO:0048538 | thymus development | 0.01750 |
| Dog Fear | GO:0031397 | negative regulation of protein ubiquitin... | 0.01760 |
| Dog Fear | GO:0042177 | negative regulation of protein catabolic... | 0.02090 |
| Dog Fear | GO:0014009 | glial cell proliferation | 0.02110 |
| Dog Fear | GO:0032770 | positive regulation of monooxygenase act... | 0.02110 |
| Dog Fear | GO:0070129 | regulation of mitochondrial translation | 0.02110 |
| Dog Fear | GO:2000001 | regulation of DNA damage checkpoint | 0.02110 |
| Dog Fear | GO:0009063 | cellular amino acid catabolic process | 0.02390 |
| Dog Fear | GO:0003015 | heart process | 0.02400 |
| Dog Fear | GO:0051209 | release of sequestered calcium ion into ... | 0.02430 |
| Dog Fear | GO:0032436 | positive regulation of proteasomal ubiqu... | 0.02440 |
| Dog Fear | GO:2001259 | positive regulation of cation channel ac... | 0.02440 |
| Dog Fear | GO:0006636 | unsaturated fatty acid biosynthetic proc... | 0.02650 |
| Dog Fear | GO:0051043 | regulation of membrane protein ectodomai... | 0.02650 |
| Dog Fear | GO:0060999 | positive regulation of dendritic spine d... | 0.02650 |
| Dog Fear | GO:0010506 | regulation of autophagy | 0.02790 |
| Dog Fear | GO:0001822 | kidney development | 0.02840 |
| Dog Fear | GO:0007612 | learning | 0.02850 |
| Dog Fear | GO:0042130 | negative regulation of T cell proliferat... | 0.02880 |
| Dog Fear | GO:0030324 | lung development | 0.02940 |
| Dog Fear | GO:0007018 | microtubule-based movement | 0.03060 |
| Dog Fear | GO:0009056 | catabolic process | 0.03230 |
| Dog Fear | GO:0042059 | negative regulation of epidermal growth ... | 0.03240 |
| Dog Fear | GO:0051216 | cartilage development | 0.03270 |
| Dog Fear | GO:0043161 | proteasome-mediated ubiquitin-dependent ... | 0.03330 |
| Dog Fear | GO:0010611 | regulation of cardiac muscle hypertrophy | 0.03450 |
| Dog Fear | GO:0034333 | adherens junction assembly | 0.03450 |
| Dog Fear | GO:0051224 | negative regulation of protein transport | 0.03450 |
| Dog Fear | GO:0099518 | vesicle cytoskeletal trafficking | 0.03450 |

| | | | |
|---|---|---|---|
| Dog Fear | GO:0006900 | vesicle budding from membrane | 0.03460 |
| Dog Fear | GO:0046834 | lipid phosphorylation | 0.03460 |
| Dog Fear | GO:0048016 | inositol phosphate-mediated signaling | 0.03460 |
| Dog Fear | GO:0110020 | regulation of actomyosin structure organ... | 0.03460 |
| Dog Fear | GO:0032088 | negative regulation of NF-kappaB transcr... | 0.03600 |
| Dog Fear | GO:0001541 | ovarian follicle development | 0.03730 |
| Dog Fear | GO:0001775 | cell activation | 0.03850 |
| Dog Fear | GO:0046887 | positive regulation of hormone secretion | 0.03880 |
| Dog Fear | GO:0046320 | regulation of fatty acid oxidation | 0.03900 |
| Dog Fear | GO:0071168 | protein localization to chromatin | 0.03900 |
| Dog Fear | GO:0097120 | receptor localization to synapse | 0.03900 |
| Dog Fear | GO:1900087 | positive regulation of G1/S transition o... | 0.03900 |
| Dog Fear | GO:0030048 | actin filament-based movement | 0.03910 |
| Dog Fear | GO:0090150 | establishment of protein localization to... | 0.04160 |
| Dog Fear | GO:0009966 | regulation of signal transduction | 0.04200 |
| Dog Fear | GO:0006470 | protein dephosphorylation | 0.04210 |
| Dog Fear | GO:0009880 | embryonic pattern specification | 0.04600 |
| Dog Fear | GO:0014059 | regulation of dopamine secretion | 0.04620 |
| Dog Fear | GO:2001023 | regulation of response to drug | 0.04630 |
| Dog Fear | GO:0006399 | tRNA metabolic process | 0.04670 |
| Dog Fear | GO:0050852 | T cell receptor signaling pathway | 0.04790 |
| Dog Rivalry | GO:0060669 | embryonic placenta morphogenesis | 0.00170 |
| Dog Rivalry | GO:0021801 | cerebral cortex radial glia guided migra... | 0.00180 |
| Dog Rivalry | GO:0071826 | ribonucleoprotein complex subunit organi... | 0.00180 |
| Dog Rivalry | GO:0042177 | negative regulation of protein catabolic... | 0.00240 |
| Dog Rivalry | GO:0060074 | synapse maturation | 0.00240 |
| Dog Rivalry | GO:0048477 | oogenesis | 0.00320 |
| Dog Rivalry | GO:0015807 | L-amino acid transport | 0.00420 |
| Dog Rivalry | GO:0050807 | regulation of synapse organization | 0.00470 |

| | | | |
|---|---|---|---|
| Dog Rivalry | GO:0001541 | ovarian follicle development | 0.00530 |
| Dog Rivalry | GO:0051443 | positive regulation of ubiquitin-protein... | 0.00540 |
| Dog Rivalry | GO:0006471 | protein ADP-ribosylation | 0.00680 |
| Dog Rivalry | GO:1900026 | positive regulation of substrate adhesio... | 0.00680 |
| Dog Rivalry | GO:0051496 | positive regulation of stress fiber asse... | 0.00870 |
| Dog Rivalry | GO:0031532 | actin cytoskeleton reorganization | 0.00880 |
| Dog Rivalry | GO:0032012 | regulation of ARF protein signal transdu... | 0.00950 |
| Dog Rivalry | GO:0050855 | regulation of B cell receptor signaling ... | 0.00950 |
| Dog Rivalry | GO:0022408 | negative regulation of cell-cell adhesio... | 0.01020 |
| Dog Rivalry | GO:0006984 | ER-nucleus signaling pathway | 0.01040 |
| Dog Rivalry | GO:0031397 | negative regulation of protein ubiquitin... | 0.01120 |
| Dog Rivalry | GO:0042100 | B cell proliferation | 0.01260 |
| Dog Rivalry | GO:1902894 | negative regulation of pri-miRNA transcr... | 0.01260 |
| Dog Rivalry | GO:0007140 | male meiotic nuclear division | 0.01610 |
| Dog Rivalry | GO:0006896 | Golgi to vacuole transport | 0.01620 |
| Dog Rivalry | GO:0050680 | negative regulation of epithelial cell p... | 0.01920 |
| Dog Rivalry | GO:0032092 | positive regulation of protein binding | 0.01930 |
| Dog Rivalry | GO:0021587 | cerebellum morphogenesis | 0.01960 |
| Dog Rivalry | GO:0051293 | establishment of spindle localization | 0.01970 |
| Dog Rivalry | GO:0006974 | cellular response to DNA damage stimulus | 0.01980 |
| Dog Rivalry | GO:0043534 | blood vessel endothelial cell migration | 0.01980 |
| Dog Rivalry | GO:0030036 | actin cytoskeleton organization | 0.01990 |
| Dog Rivalry | GO:0043046 | DNA methylation involved in gamete gener... | 0.02030 |
| Dog Rivalry | GO:0010464 | regulation of mesenchymal cell prolifera... | 0.02870 |
| Dog Rivalry | GO:0044782 | cilium organization | 0.02870 |
| Dog Rivalry | GO:0030001 | metal ion transport | 0.02880 |
| Dog Rivalry | GO:0035561 | regulation of chromatin binding | 0.03020 |
| Dog Rivalry | GO:0090114 | COPII-coated vesicle budding | 0.03020 |
| Dog Rivalry | GO:0090305 | nucleic acid phosphodiester bond hydroly... | 0.03100 |

| | | | |
|---|---|---|---|
| Dog Rivalry | GO:0035023 | regulation of Rho protein signal transdu... | 0.03270 |
| Dog Rivalry | GO:0030316 | osteoclast differentiation | 0.03590 |
| Dog Rivalry | GO:0033043 | regulation of organelle organization | 0.03680 |
| Dog Rivalry | GO:1903046 | meiotic cell cycle process | 0.03810 |
| Dog Rivalry | GO:0090101 | negative regulation of transmembrane rec... | 0.03880 |
| Dog Rivalry | GO:0015804 | neutral amino acid transport | 0.03940 |
| Dog Rivalry | GO:0010001 | glial cell differentiation | 0.04210 |
| Dog Rivalry | GO:0010837 | regulation of keratinocyte proliferation | 0.04210 |
| Dog Rivalry | GO:0045892 | negative regulation of transcription, DN... | 0.04610 |
| Dog Rivalry | GO:0021511 | spinal cord patterning | 0.04650 |
| Dog Rivalry | GO:0044036 | cell wall macromolecule metabolic proces... | 0.04650 |
| Dog Rivalry | GO:0030031 | cell projection assembly | 0.04660 |
| Dog Rivalry | GO:1902850 | microtubule cytoskeleton organization in... | 0.04660 |
| Dog Rivalry | GO:1904894 | positive regulation of STAT cascade | 0.04660 |
| Dog Rivalry | GO:0034654 | nucleobase-containing compound biosynthe... | 0.04800 |
| Dog Rivalry | GO:0009225 | nucleotide-sugar metabolic process | 0.04880 |
| Dog Rivalry | GO:0010955 | negative regulation of protein processin... | 0.04880 |
| Dog Rivalry | GO:0070830 | bicellular tight junction assembly | 0.04880 |
| Dog Rivalry | GO:2000352 | negative regulation of endothelial cell ... | 0.04880 |
| Energy | GO:0051570 | regulation of histone H3-K9 methylation | 0.00110 |
| Energy | GO:0001764 | neuron migration | 0.00130 |
| Energy | GO:0098742 | cell-cell adhesion via plasma-membrane a... | 0.00170 |
| Energy | GO:0006622 | protein targeting to lysosome | 0.00230 |
| Energy | GO:0007099 | centriole replication | 0.00230 |
| Energy | GO:0048172 | regulation of short-term neuronal synapt... | 0.00230 |
| Energy | GO:2000209 | regulation of anoikis | 0.00230 |
| Energy | GO:0006886 | intracellular protein transport | 0.00240 |
| Energy | GO:0048710 | regulation of astrocyte differentiation | 0.00300 |
| Energy | GO:0070509 | calcium ion import | 0.00300 |

| | | | |
|---|---|---|---|
| Energy | GO:1905515 | non-motile cilium assembly | 0.00320 |
| Energy | GO:0016445 | somatic diversification of immunoglobuli... | 0.00380 |
| Energy | GO:0086002 | cardiac muscle cell action potential inv... | 0.00480 |
| Energy | GO:0000413 | protein peptidyl-prolyl isomerization | 0.00500 |
| Energy | GO:0030036 | actin cytoskeleton organization | 0.00540 |
| Energy | GO:0006906 | vesicle fusion | 0.00620 |
| Energy | GO:0031648 | protein destabilization | 0.00650 |
| Energy | GO:0031061 | negative regulation of histone methylati... | 0.00670 |
| Energy | GO:0070076 | histone lysine demethylation | 0.00670 |
| Energy | GO:0022408 | negative regulation of cell-cell adhesio... | 0.00780 |
| Energy | GO:0001662 | behavioral fear response | 0.00800 |
| Energy | GO:0010954 | positive regulation of protein processin... | 0.00890 |
| Energy | GO:0010951 | negative regulation of endopeptidase act... | 0.00910 |
| Energy | GO:0007528 | neuromuscular junction development | 0.00990 |
| Energy | GO:0031102 | neuron projection regeneration | 0.01100 |
| Energy | GO:2000114 | regulation of establishment of cell pola... | 0.01160 |
| Energy | GO:0086019 | cell-cell signaling involved in cardiac ... | 0.01470 |
| Energy | GO:0090023 | positive regulation of neutrophil chemot... | 0.01470 |
| Energy | GO:0021549 | cerebellum development | 0.01500 |
| Energy | GO:2000573 | positive regulation of DNA biosynthetic ... | 0.01500 |
| Energy | GO:0006890 | retrograde vesicle-mediated transport, G... | 0.01830 |
| Energy | GO:0017158 | regulation of calcium ion-dependent exoc... | 0.02000 |
| Energy | GO:0010457 | centriole-centriole cohesion | 0.02050 |
| Energy | GO:0032012 | regulation of ARF protein signal transdu... | 0.02050 |
| Energy | GO:2000637 | positive regulation of gene silencing by... | 0.02050 |
| Energy | GO:0046328 | regulation of JNK cascade | 0.02080 |
| Energy | GO:0031023 | microtubule organizing center organizati... | 0.02100 |
| Energy | GO:1905475 | regulation of protein localization to me... | 0.02120 |
| Energy | GO:0032774 | RNA biosynthetic process | 0.02320 |

| | | | |
|---|---|---|---|
| Energy | GO:0007173 | epidermal growth factor receptor signali... | 0.02360 |
| Energy | GO:0003254 | regulation of membrane depolarization | 0.02700 |
| Energy | GO:0045879 | negative regulation of smoothened signal... | 0.02700 |
| Energy | GO:0051571 | positive regulation of histone H3-K4 met... | 0.02700 |
| Energy | GO:0055026 | negative regulation of cardiac muscle ti... | 0.02700 |
| Energy | GO:0060122 | inner ear receptor cell stereocilium org... | 0.02700 |
| Energy | GO:0086012 | membrane depolarization during cardiac m... | 0.02700 |
| Energy | GO:0090140 | regulation of mitochondrial fission | 0.02700 |
| Energy | GO:0045732 | positive regulation of protein catabolic... | 0.02850 |
| Energy | GO:0007030 | Golgi organization | 0.03070 |
| Energy | GO:0086091 | regulation of heart rate by cardiac cond... | 0.03110 |
| Energy | GO:0048168 | regulation of neuronal synaptic plastici... | 0.03360 |
| Energy | GO:0032205 | negative regulation of telomere maintena... | 0.03380 |
| Energy | GO:0045190 | isotype switching | 0.03390 |
| Energy | GO:0021587 | cerebellum morphogenesis | 0.03400 |
| Energy | GO:0043405 | regulation of MAP kinase activity | 0.03410 |
| Energy | GO:0018023 | peptidyl-lysine trimethylation | 0.03430 |
| Energy | GO:0045910 | negative regulation of DNA recombination | 0.03430 |
| Energy | GO:0071634 | regulation of transforming growth factor... | 0.03430 |
| Energy | GO:1903861 | positive regulation of dendrite extensio... | 0.03430 |
| Energy | GO:0042177 | negative regulation of protein catabolic... | 0.03790 |
| Energy | GO:0050770 | regulation of axonogenesis | 0.03790 |
| Energy | GO:0007368 | determination of left/right symmetry | 0.03990 |
| Energy | GO:0035249 | synaptic transmission, glutamatergic | 0.04240 |
| Energy | GO:0003198 | epithelial to mesenchymal transition inv... | 0.04260 |
| Energy | GO:0017001 | antibiotic catabolic process | 0.04260 |
| Energy | GO:0030903 | notochord development | 0.04260 |
| Energy | GO:0039694 | viral RNA genome replication | 0.04260 |
| Energy | GO:0045686 | negative regulation of glial cell differ... | 0.04260 |

| | | | |
|---|---|---|---|
| Energy | GO:0062009 | secondary palate development | 0.04260 |
| Energy | GO:2000826 | regulation of heart morphogenesis | 0.04260 |
| Energy | GO:0045022 | early endosome to late endosome transpor... | 0.04400 |
| Energy | GO:0048791 | calcium ion-regulated exocytosis of neur... | 0.04400 |
| Energy | GO:0043409 | negative regulation of MAPK cascade | 0.04910 |
| Energy | GO:0051090 | regulation of DNA binding transcription ... | 0.04920 |
| Excitability | GO:0051291 | protein heterooligomerization | 0.00100 |
| Excitability | GO:0018105 | peptidyl-serine phosphorylation | 0.00130 |
| Excitability | GO:0031032 | actomyosin structure organization | 0.00190 |
| Excitability | GO:0010507 | negative regulation of autophagy | 0.00290 |
| Excitability | GO:0030540 | female genitalia development | 0.00290 |
| Excitability | GO:0010996 | response to auditory stimulus | 0.00430 |
| Excitability | GO:0032970 | regulation of actin filament-based proce... | 0.00430 |
| Excitability | GO:0030890 | positive regulation of B cell proliferat... | 0.00480 |
| Excitability | GO:0071622 | regulation of granulocyte chemotaxis | 0.00490 |
| Excitability | GO:0048286 | lung alveolus development | 0.00600 |
| Excitability | GO:0060042 | retina morphogenesis in camera-type eye | 0.00610 |
| Excitability | GO:0071345 | cellular response to cytokine stimulus | 0.00760 |
| Excitability | GO:0010758 | regulation of macrophage chemotaxis | 0.00830 |
| Excitability | GO:0050891 | multicellular organismal water homeostas... | 0.00830 |
| Excitability | GO:0090407 | organophosphate biosynthetic process | 0.00880 |
| Excitability | GO:0050869 | negative regulation of B cell activation | 0.01110 |
| Excitability | GO:0030888 | regulation of B cell proliferation | 0.01230 |
| Excitability | GO:0006282 | regulation of DNA repair | 0.01250 |
| Excitability | GO:0021885 | forebrain cell migration | 0.01250 |
| Excitability | GO:0031952 | regulation of protein autophosphorylatio... | 0.01250 |
| Excitability | GO:0042698 | ovulation cycle | 0.01250 |
| Excitability | GO:0061099 | negative regulation of protein tyrosine ... | 0.01430 |
| Excitability | GO:0071901 | negative regulation of protein serine/th... | 0.01440 |

| | | | |
|---|---|---|---|
| Excitability | GO:0044271 | cellular nitrogen compound biosynthetic ... | 0.01470 |
| Excitability | GO:0007605 | sensory perception of sound | 0.01490 |
| Excitability | GO:0007166 | cell surface receptor signaling pathway | 0.01500 |
| Excitability | GO:0006644 | phospholipid metabolic process | 0.01550 |
| Excitability | GO:0007030 | Golgi organization | 0.01760 |
| Excitability | GO:0045923 | positive regulation of fatty acid metabo... | 0.01810 |
| Excitability | GO:0043967 | histone H4 acetylation | 0.01830 |
| Excitability | GO:0023052 | signaling | 0.02180 |
| Excitability | GO:0008045 | motor neuron axon guidance | 0.02240 |
| Excitability | GO:0032728 | positive regulation of interferon-beta p... | 0.02240 |
| Excitability | GO:0051281 | positive regulation of release of seques... | 0.02240 |
| Excitability | GO:0002761 | regulation of myeloid leukocyte differen... | 0.02380 |
| Excitability | GO:0071402 | cellular response to lipoprotein particl... | 0.02380 |
| Excitability | GO:0010447 | response to acidic pH | 0.02420 |
| Excitability | GO:0032367 | intracellular cholesterol transport | 0.02420 |
| Excitability | GO:0046839 | phospholipid dephosphorylation | 0.02420 |
| Excitability | GO:0070296 | sbackground-color:#f2f2f2;oplasmic reticulum calcium ion trans... | 0.02420 |
| Excitability | GO:0072425 | signal transduction involved in G2 DNA d... | 0.02420 |
| Excitability | GO:1901532 | regulation of hematopoietic progenitor c... | 0.02420 |
| Excitability | GO:1903514 | release of sequestered calcium ion into ... | 0.02420 |
| Excitability | GO:2000052 | positive regulation of non-canonical Wnt... | 0.02420 |
| Excitability | GO:2000095 | regulation of Wnt signaling pathway, pla... | 0.02420 |
| Excitability | GO:0043124 | negative regulation of I-kappaB kinase/N... | 0.02550 |
| Excitability | GO:0006468 | protein phosphorylation | 0.02650 |
| Excitability | GO:0046718 | viral entry into host cell | 0.02730 |
| Excitability | GO:0070498 | interleukin-1-mediated signaling pathway | 0.02740 |
| Excitability | GO:0018130 | heterocycle biosynthetic process | 0.02750 |
| Excitability | GO:0007173 | epidermal growth factor receptor signali... | 0.02970 |
| Excitability | GO:1901566 | organonitrogen compound biosynthetic pro... | 0.03050 |

| | | | |
|---|---|---|---|
| Excitability | GO:0046777 | protein autophosphorylation | 0.03060 |
| Excitability | GO:0006336 | DNA replication-independent nucleosome a... | 0.03170 |
| Excitability | GO:0030033 | microvillus assembly | 0.03170 |
| Excitability | GO:0034311 | diol metabolic process | 0.03170 |
| Excitability | GO:0060428 | lung epithelium development | 0.03170 |
| Excitability | GO:1905939 | regulation of gonad development | 0.03170 |
| Excitability | GO:0043627 | response to estrogen | 0.03290 |
| Excitability | GO:0050850 | positive regulation of calcium-mediated ... | 0.03290 |
| Excitability | GO:0008104 | protein localization | 0.03550 |
| Excitability | GO:0046464 | acylglycerol catabolic process | 0.03780 |
| Excitability | GO:0050672 | negative regulation of lymphocyte prolif... | 0.03790 |
| Excitability | GO:0051250 | negative regulation of lymphocyte activa... | 0.03790 |
| Excitability | GO:2000736 | regulation of stem cell differentiation | 0.03790 |
| Excitability | GO:0007210 | serotonin receptor signaling pathway | 0.03800 |
| Excitability | GO:0031110 | regulation of microtubule polymerization... | 0.03800 |
| Excitability | GO:0048168 | regulation of neuronal synaptic plastici... | 0.03800 |
| Excitability | GO:0016447 | somatic recombination of immunoglobulin ... | 0.03810 |
| Excitability | GO:0051196 | regulation of coenzyme metabolic process | 0.03810 |
| Excitability | GO:0007165 | signal transduction | 0.03820 |
| Excitability | GO:0043277 | apoptotic cell clearance | 0.03910 |
| Excitability | GO:0032722 | positive regulation of chemokine product... | 0.03920 |
| Excitability | GO:0042098 | T cell proliferation | 0.04010 |
| Excitability | GO:0010812 | negative regulation of cell-substrate ad... | 0.04020 |
| Excitability | GO:0043030 | regulation of macrophage activation | 0.04020 |
| Excitability | GO:0046653 | tetrahydrofolate metabolic process | 0.04020 |
| Excitability | GO:0050908 | detection of light stimulus involved in ... | 0.04020 |
| Excitability | GO:0050922 | negative regulation of chemotaxis | 0.04020 |
| Excitability | GO:0051446 | positive regulation of meiotic cell cycl... | 0.04020 |
| Excitability | GO:0045893 | positive regulation of transcription, DN... | 0.04480 |

| | | | |
|---|---|---|---|
| Excitability | GO:0006790 | sulfur compound metabolic process | 0.04530 |
| Excitability | GO:0016572 | histone phosphorylation | 0.04580 |
| Excitability | GO:0070884 | regulation of calcineurin-NFAT signaling... | 0.04580 |
| Excitability | GO:0009749 | response to glucose | 0.04600 |
| Excitability | GO:0042177 | negative regulation of protein catabolic... | 0.04600 |
| Excitability | GO:0045087 | innate immune response | 0.04680 |
| Excitability | GO:0001843 | neural tube closure | 0.04870 |
| Excitability | GO:0018107 | peptidyl-threonine phosphorylation | 0.04900 |
| Excitability | GO:0002385 | mucosal immune response | 0.04980 |
| Excitability | GO:0003009 | skeletal muscle contraction | 0.04980 |
| Excitability | GO:0010613 | positive regulation of cardiac muscle hy... | 0.04980 |
| Excitability | GO:0017001 | antibiotic catabolic process | 0.04980 |
| Excitability | GO:0019433 | triglyceride catabolic process | 0.04980 |
| Excitability | GO:0032727 | positive regulation of interferon-alpha ... | 0.04980 |
| Excitability | GO:0034383 | low-density lipoprotein particle clearan... | 0.04980 |
| Excitability | GO:0045663 | positive regulation of myoblast differen... | 0.04980 |
| Excitability | GO:0045686 | negative regulation of glial cell differ... | 0.04980 |
| Excitability | GO:0046173 | polyol biosynthetic process | 0.04980 |
| Excitability | GO:2001024 | negative regulation of response to drug | 0.04980 |
| Excitability | GO:0060411 | cardiac septum morphogenesis | 0.04990 |
| Nonsocial Fear | GO:0045773 | positive regulation of axon extension | 0.00180 |
| Nonsocial Fear | GO:0010575 | positive regulation of vascular endothel... | 0.00270 |
| Nonsocial Fear | GO:0034220 | ion transmembrane transport | 0.00280 |
| Nonsocial Fear | GO:0031290 | retinal ganglion cell axon guidance | 0.00540 |
| Nonsocial Fear | GO:0042307 | positive regulation of protein import in... | 0.00690 |
| Nonsocial Fear | GO:0050919 | negative chemotaxis | 0.00690 |
| Nonsocial Fear | GO:0051293 | establishment of spindle localization | 0.00770 |
| Nonsocial Fear | GO:0003007 | heart morphogenesis | 0.00980 |
| Nonsocial Fear | GO:0033522 | histone H2A ubiquitination | 0.01130 |

| | | | |
|---|---|---|---|
| Nonsocial Fear | GO:0046031 | ADP metabolic process | 0.01130 |
| Nonsocial Fear | GO:0035235 | ionotropic glutamate receptor signaling ... | 0.01170 |
| Nonsocial Fear | GO:0006914 | autophagy | 0.01360 |
| Nonsocial Fear | GO:0016239 | positive regulation of macroautophagy | 0.01550 |
| Nonsocial Fear | GO:0071345 | cellular response to cytokine stimulus | 0.01660 |
| Nonsocial Fear | GO:1903506 | regulation of nucleic acid-templated tra... | 0.01660 |
| Nonsocial Fear | GO:0060047 | heart contraction | 0.02030 |
| Nonsocial Fear | GO:0007411 | axon guidance | 0.02050 |
| Nonsocial Fear | GO:0071453 | cellular response to oxygen levels | 0.02550 |
| Nonsocial Fear | GO:0032608 | interferon-beta production | 0.02850 |
| Nonsocial Fear | GO:0032785 | negative regulation of DNA-templated tra... | 0.02850 |
| Nonsocial Fear | GO:0086001 | cardiac muscle cell action potential | 0.02850 |
| Nonsocial Fear | GO:0009218 | pyrimidine ribonucleotide metabolic proc... | 0.02860 |
| Nonsocial Fear | GO:0010657 | muscle cell apoptotic process | 0.02860 |
| Nonsocial Fear | GO:0021988 | olfactory lobe development | 0.02860 |
| Nonsocial Fear | GO:0006811 | ion transport | 0.02870 |
| Nonsocial Fear | GO:0015749 | monosaccharide transmembrane transport | 0.02870 |
| Nonsocial Fear | GO:0044419 | interspecies interaction between organis... | 0.02870 |
| Nonsocial Fear | GO:0046546 | development of primary male sexual chara... | 0.02870 |
| Nonsocial Fear | GO:0065009 | regulation of molecular function | 0.02940 |
| Nonsocial Fear | GO:0002070 | epithelial cell maturation | 0.03160 |
| Nonsocial Fear | GO:0006027 | glycosaminoglycan catabolic process | 0.03160 |
| Nonsocial Fear | GO:0016998 | cell wall macromolecule catabolic proces... | 0.03160 |
| Nonsocial Fear | GO:0032352 | positive regulation of hormone metabolic... | 0.03160 |
| Nonsocial Fear | GO:0034143 | regulation of toll-like receptor 4 signa... | 0.03160 |
| Nonsocial Fear | GO:0007600 | sensory perception | 0.03170 |
| Nonsocial Fear | GO:0009886 | post-embryonic animal morphogenesis | 0.03790 |
| Nonsocial Fear | GO:0032438 | melanosome organization | 0.03790 |
| Nonsocial Fear | GO:0043248 | proteasome assembly | 0.03790 |

| | | | |
|---|---|---|---|
| Nonsocial Fear | GO:0048820 | hair follicle maturation | 0.03790 |
| Nonsocial Fear | GO:0060575 | intestinal epithelial cell differentiati... | 0.03790 |
| Nonsocial Fear | GO:0070932 | histone H3 deacetylation | 0.03790 |
| Nonsocial Fear | GO:0035023 | regulation of Rho protein signal transdu... | 0.03950 |
| Nonsocial Fear | GO:0098609 | cell-cell adhesion | 0.04000 |
| Nonsocial Fear | GO:0007032 | endosome organization | 0.04090 |
| Nonsocial Fear | GO:0006793 | phosphorus metabolic process | 0.04350 |
| Nonsocial Fear | GO:0002230 | positive regulation of defense response ... | 0.04460 |
| Nonsocial Fear | GO:0034505 | tooth mineralization | 0.04460 |
| Nonsocial Fear | GO:0045777 | positive regulation of blood pressure | 0.04460 |
| Nonsocial Fear | GO:0046033 | AMP metabolic process | 0.04460 |
| Nonsocial Fear | GO:0071985 | multivesicular body sorting pathway | 0.04460 |
| Nonsocial Fear | GO:1903861 | positive regulation of dendrite extensio... | 0.04460 |
| Nonsocial Fear | GO:0042733 | embryonic digit morphogenesis | 0.04480 |
| Nonsocial Fear | GO:0007507 | heart development | 0.04830 |
| Owner Aggression | GO:0010506 | regulation of autophagy | 0.00011 |
| Owner Aggression | GO:0006359 | regulation of transcription by RNA polym... | 0.00166 |
| Owner Aggression | GO:0032781 | positive regulation of ATPase activity | 0.00218 |
| Owner Aggression | GO:0018210 | peptidyl-threonine modification | 0.00237 |
| Owner Aggression | GO:0048813 | dendrite morphogenesis | 0.00297 |
| Owner Aggression | GO:0032008 | positive regulation of TOR signaling | 0.00435 |
| Owner Aggression | GO:1901021 | positive regulation of calcium ion trans... | 0.00447 |
| Owner Aggression | GO:0032535 | regulation of cellular component size | 0.00579 |
| Owner Aggression | GO:1900087 | positive regulation of G1/S transition o... | 0.00585 |
| Owner Aggression | GO:0050798 | activated T cell proliferation | 0.00752 |
| Owner Aggression | GO:0046323 | glucose import | 0.00886 |
| Owner Aggression | GO:0035235 | ionotropic glutamate receptor signaling ... | 0.00940 |
| Owner Aggression | GO:0034220 | ion transmembrane transport | 0.01080 |
| Owner Aggression | GO:0033674 | positive regulation of kinase activity | 0.01212 |

| | | | |
|---|---|---|---|
| Owner Aggression | GO:0009435 | NAD biosynthetic process | 0.01214 |
| Owner Aggression | GO:0035459 | cargo loading into vesicle | 0.01214 |
| Owner Aggression | GO:1901018 | positive regulation of potassium ion tra... | 0.01214 |
| Owner Aggression | GO:0030010 | establishment of cell polarity | 0.01237 |
| Owner Aggression | GO:0050806 | positive regulation of synaptic transmis... | 0.01385 |
| Owner Aggression | GO:0032956 | regulation of actin cytoskeleton organiz... | 0.01406 |
| Owner Aggression | GO:0050766 | positive regulation of phagocytosis | 0.01412 |
| Owner Aggression | GO:0097352 | autophagosome maturation | 0.01412 |
| Owner Aggression | GO:0030030 | cell projection organization | 0.01414 |
| Owner Aggression | GO:0006821 | chloride transport | 0.01417 |
| Owner Aggression | GO:0051303 | establishment of chromosome localization | 0.01448 |
| Owner Aggression | GO:0042100 | B cell proliferation | 0.01596 |
| Owner Aggression | GO:1901216 | positive regulation of neuron death | 0.01604 |
| Owner Aggression | GO:0006308 | DNA catabolic process | 0.01607 |
| Owner Aggression | GO:0050650 | chondroitin sulfate proteoglycan biosynt... | 0.01607 |
| Owner Aggression | GO:2000251 | positive regulation of actin cytoskeleto... | 0.01607 |
| Owner Aggression | GO:0007595 | lactation | 0.01696 |
| Owner Aggression | GO:0050770 | regulation of axonogenesis | 0.01912 |
| Owner Aggression | GO:0007140 | male meiotic nuclear division | 0.02059 |
| Owner Aggression | GO:0072520 | seminiferous tubule development | 0.02062 |
| Owner Aggression | GO:0051293 | establishment of spindle localization | 0.02340 |
| Owner Aggression | GO:0042471 | ear morphogenesis | 0.02342 |
| Owner Aggression | GO:0080134 | regulation of response to stress | 0.02351 |
| Owner Aggression | GO:0051017 | actin filament bundle assembly | 0.02353 |
| Owner Aggression | GO:0043279 | response to alkaloid | 0.02579 |
| Owner Aggression | GO:0007413 | axonal fasciculation | 0.02581 |
| Owner Aggression | GO:0042430 | indole-containing compound metabolic pro... | 0.02581 |
| Owner Aggression | GO:0050821 | protein stabilization | 0.02894 |
| Owner Aggression | GO:0010975 | regulation of neuron projection developm... | 0.03083 |

| | | | |
|---|---|---|---|
| Owner Aggression | GO:0006493 | protein O-linked glycosylation | 0.03117 |
| Owner Aggression | GO:0043484 | regulation of RNA splicing | 0.03121 |
| Owner Aggression | GO:0010738 | regulation of protein kinase A signaling | 0.03164 |
| Owner Aggression | GO:0010765 | positive regulation of sodium ion transp... | 0.03164 |
| Owner Aggression | GO:0010954 | positive regulation of protein processin... | 0.03164 |
| Owner Aggression | GO:0060292 | long term synaptic depression | 0.03164 |
| Owner Aggression | GO:0006874 | cellular calcium ion homeostasis | 0.03168 |
| Owner Aggression | GO:0006974 | cellular response to DNA damage stimulus | 0.03186 |
| Owner Aggression | GO:0030001 | metal ion transport | 0.03355 |
| Owner Aggression | GO:0055078 | sodium ion homeostasis | 0.03808 |
| Owner Aggression | GO:1903432 | regulation of TORC1 signaling | 0.03808 |
| Owner Aggression | GO:0018209 | peptidyl-serine modification | 0.03823 |
| Owner Aggression | GO:0019439 | aromatic compound catabolic process | 0.04097 |
| Owner Aggression | GO:0043124 | negative regulation of I-kappaB kinase/N... | 0.04117 |
| Owner Aggression | GO:0090102 | cochlea development | 0.04117 |
| Owner Aggression | GO:2000649 | regulation of sodium ion transmembrane t... | 0.04117 |
| Owner Aggression | GO:0060079 | excitatory postsynaptic potential | 0.04250 |
| Owner Aggression | GO:0001843 | neural tube closure | 0.04418 |
| Owner Aggression | GO:0051452 | intracellular pH reduction | 0.04514 |
| Owner Aggression | GO:1903046 | meiotic cell cycle process | 0.04537 |
| Owner Aggression | GO:0097193 | intrinsic apoptotic signaling pathway | 0.04609 |
| Owner Aggression | GO:0031098 | stress-activated protein kinase signalin... | 0.04701 |
| Separation Problems | GO:0062009 | secondary palate development | 0.00100 |
| Separation Problems | GO:0051099 | positive regulation of binding | 0.00130 |
| Separation Problems | GO:0090162 | establishment of epithelial cell polarit... | 0.00140 |
| Separation Problems | GO:0042176 | regulation of protein catabolic process | 0.00160 |
| Separation Problems | GO:0034329 | cell junction assembly | 0.00400 |
| Separation Problems | GO:0010927 | cellular component assembly involved in ... | 0.00460 |
| Separation Problems | GO:0031498 | chromatin disassembly | 0.00500 |

| | | | |
|---|---|---|---|
| Separation Problems | GO:0090177 | establishment of planar polarity involve... | 0.00500 |
| Separation Problems | GO:0006833 | water transport | 0.00660 |
| Separation Problems | GO:0030866 | cortical actin cytoskeleton organization | 0.00670 |
| Separation Problems | GO:0051304 | chromosome separation | 0.00770 |
| Separation Problems | GO:0001919 | regulation of receptor recycling | 0.01090 |
| Separation Problems | GO:1905276 | regulation of epithelial tube formation | 0.01090 |
| Separation Problems | GO:0010977 | negative regulation of neuron projection... | 0.01220 |
| Separation Problems | GO:2000725 | regulation of cardiac muscle cell differ... | 0.01260 |
| Separation Problems | GO:1901021 | positive regulation of calcium ion trans... | 0.01350 |
| Separation Problems | GO:0007411 | axon guidance | 0.01450 |
| Separation Problems | GO:0035561 | regulation of chromatin binding | 0.01640 |
| Separation Problems | GO:0046834 | lipid phosphorylation | 0.01840 |
| Separation Problems | GO:0010611 | regulation of cardiac muscle hypertrophy | 0.01850 |
| Separation Problems | GO:0051960 | regulation of nervous system development | 0.01870 |
| Separation Problems | GO:0001756 | somitogenesis | 0.02250 |
| Separation Problems | GO:0060071 | Wnt signaling pathway, planar cell polar... | 0.02310 |
| Separation Problems | GO:0040020 | regulation of meiotic nuclear division | 0.02320 |
| Separation Problems | GO:0003203 | endocardial cushion morphogenesis | 0.02530 |
| Separation Problems | GO:0010922 | positive regulation of phosphatase activ... | 0.02530 |
| Separation Problems | GO:0055008 | cardiac muscle tissue morphogenesis | 0.02530 |
| Separation Problems | GO:0060317 | cardiac epithelial to mesenchymal transi... | 0.02530 |
| Separation Problems | GO:0000186 | activation of MAPKK activity | 0.02640 |
| Separation Problems | GO:0001702 | gastrulation with mouth forming second | 0.02710 |
| Separation Problems | GO:0061311 | cell surface receptor signaling pathway ... | 0.02710 |
| Separation Problems | GO:0043280 | positive regulation of cysteine-type end... | 0.02720 |
| Separation Problems | GO:0051154 | negative regulation of striated muscle c... | 0.03290 |
| Separation Problems | GO:0060047 | heart contraction | 0.03300 |
| Separation Problems | GO:0034220 | ion transmembrane transport | 0.03380 |
| Separation Problems | GO:0060255 | regulation of macromolecule metabolic pr... | 0.03430 |

| | | | |
|---|---|---|---|
| Separation Problems | GO:0001947 | heart looping | 0.03570 |
| Separation Problems | GO:0060548 | negative regulation of cell death | 0.03580 |
| Separation Problems | GO:0008608 | attachment of spindle microtubules to ki... | 0.03590 |
| Separation Problems | GO:0000305 | response to oxygen radical | 0.03690 |
| Separation Problems | GO:0046546 | development of primary male sexual chara... | 0.03690 |
| Separation Problems | GO:0002757 | immune response-activating signal transd... | 0.03700 |
| Separation Problems | GO:0048017 | inositol lipid-mediated signaling | 0.03700 |
| Separation Problems | GO:0015749 | monosaccharide transmembrane transport | 0.03710 |
| Separation Problems | GO:0031032 | actomyosin structure organization | 0.04040 |
| Separation Problems | GO:0048013 | ephrin receptor signaling pathway | 0.04080 |
| Separation Problems | GO:0034332 | adherens junction organization | 0.04130 |
| Separation Problems | GO:0043044 | ATP-dependent chromatin remodeling | 0.04130 |
| Separation Problems | GO:1903725 | regulation of phospholipid metabolic pro... | 0.04140 |
| Separation Problems | GO:0071346 | cellular response to interferon-gamma | 0.04520 |
| Separation Problems | GO:0032467 | positive regulation of cytokinesis | 0.04600 |
| Separation Problems | GO:0045197 | establishment or maintenance of epitheli... | 0.04600 |
| Separation Problems | GO:0050922 | negative regulation of chemotaxis | 0.04600 |
| Separation Problems | GO:2001259 | positive regulation of cation channel ac... | 0.04600 |
| Separation Problems | GO:0043966 | histone H3 acetylation | 0.04660 |
| Stranger Aggression | GO:0015813 | L-glutamate transmembrane transport | 0.00120 |
| Stranger Aggression | GO:0060047 | heart contraction | 0.00220 |
| Stranger Aggression | GO:0070498 | interleukin-1-mediated signaling pathway | 0.00330 |
| Stranger Aggression | GO:0046112 | nucleobase biosynthetic process | 0.00450 |
| Stranger Aggression | GO:0050881 | musculoskeletal movement | 0.00450 |
| Stranger Aggression | GO:0070296 | sbackground-color:#f2f2f2;oplasmic reticulum calcium ion trans... | 0.00450 |
| Stranger Aggression | GO:1903514 | release of sequestered calcium ion into ... | 0.00450 |
| Stranger Aggression | GO:0033043 | regulation of organelle organization | 0.00560 |
| Stranger Aggression | GO:0009070 | serine family amino acid biosynthetic pr... | 0.00610 |
| Stranger Aggression | GO:0072111 | cell proliferation involved in kidney de... | 0.00610 |

| | | | |
|---|---|---|---|
| Stranger Aggression | GO:0072178 | nephric duct morphogenesis | 0.00610 |
| Stranger Aggression | GO:0097479 | synaptic vesicle localization | 0.00730 |
| Stranger Aggression | GO:0006896 | Golgi to vacuole transport | 0.00790 |
| Stranger Aggression | GO:0007026 | negative regulation of microtubule depol... | 0.00790 |
| Stranger Aggression | GO:0046653 | tetrahydrofolate metabolic process | 0.00790 |
| Stranger Aggression | GO:0051383 | kinetochore organization | 0.00790 |
| Stranger Aggression | GO:0070233 | negative regulation of T cell apoptotic ... | 0.00790 |
| Stranger Aggression | GO:0071322 | cellular response to carbohydrate stimul... | 0.01200 |
| Stranger Aggression | GO:1901998 | toxin transport | 0.01300 |
| Stranger Aggression | GO:0001556 | oocyte maturation | 0.01510 |
| Stranger Aggression | GO:1903432 | regulation of TORC1 signaling | 0.01510 |
| Stranger Aggression | GO:0043123 | positive regulation of I-kappaB kinase/N... | 0.01650 |
| Stranger Aggression | GO:0006623 | protein targeting to vacuole | 0.01730 |
| Stranger Aggression | GO:0010464 | regulation of mesenchymal cell prolifera... | 0.01740 |
| Stranger Aggression | GO:0051146 | striated muscle cell differentiation | 0.01760 |
| Stranger Aggression | GO:0071773 | cellular response to BMP stimulus | 0.01760 |
| Stranger Aggression | GO:0002089 | lens morphogenesis in camera-type eye | 0.01810 |
| Stranger Aggression | GO:0097150 | neuronal stem cell population maintenanc... | 0.01810 |
| Stranger Aggression | GO:2000242 | negative regulation of reproductive proc... | 0.01900 |
| Stranger Aggression | GO:0007041 | lysosomal transport | 0.02110 |
| Stranger Aggression | GO:0006730 | one-carbon metabolic process | 0.02140 |
| Stranger Aggression | GO:0006890 | retrograde vesicle-mediated transport, G... | 0.02140 |
| Stranger Aggression | GO:0050974 | detection of mechanical stimulus involve... | 0.02380 |
| Stranger Aggression | GO:0086065 | cell communication involved in cardiac c... | 0.02380 |
| Stranger Aggression | GO:0002699 | positive regulation of immune effector p... | 0.02390 |
| Stranger Aggression | GO:0070306 | lens fiber cell differentiation | 0.02500 |
| Stranger Aggression | GO:0006418 | tRNA aminoacylation for protein translat... | 0.02650 |
| Stranger Aggression | GO:0021952 | central nervous system projection neuron... | 0.02890 |
| Stranger Aggression | GO:0045879 | negative regulation of smoothened signal... | 0.02890 |

| | | | |
|---|---|---|---|
| Stranger Aggression | GO:0040013 | negative regulation of locomotion | 0.03070 |
| Stranger Aggression | GO:0030520 | intracellular estrogen receptor signalin... | 0.03110 |
| Stranger Aggression | GO:0042130 | negative regulation of T cell proliferat... | 0.03230 |
| Stranger Aggression | GO:0032388 | positive regulation of intracellular tra... | 0.03280 |
| Stranger Aggression | GO:0001958 | endochondral ossification | 0.03310 |
| Stranger Aggression | GO:0008608 | attachment of spindle microtubules to ki... | 0.03310 |
| Stranger Aggression | GO:0021987 | cerebral cortex development | 0.03520 |
| Stranger Aggression | GO:0009799 | specification of symmetry | 0.03580 |
| Stranger Aggression | GO:0061326 | renal tubule development | 0.03580 |
| Stranger Aggression | GO:0050865 | regulation of cell activation | 0.03600 |
| Stranger Aggression | GO:0042177 | negative regulation of protein catabolic... | 0.03780 |
| Stranger Aggression | GO:1903169 | regulation of calcium ion transmembrane ... | 0.03830 |
| Stranger Aggression | GO:0034332 | adherens junction organization | 0.03860 |
| Stranger Aggression | GO:0032088 | negative regulation of NF-kappaB transcr... | 0.03880 |
| Stranger Aggression | GO:0032008 | positive regulation of TOR signaling | 0.04250 |
| Stranger Aggression | GO:2001259 | positive regulation of cation channel ac... | 0.04250 |
| Stranger Aggression | GO:0031175 | neuron projection development | 0.04600 |
| Stranger Aggression | GO:0060627 | regulation of vesicle-mediated transport | 0.04610 |
| Stranger Aggression | GO:0001755 | neural crest cell migration | 0.04760 |
| Stranger Aggression | GO:0002224 | toll-like receptor signaling pathway | 0.04760 |
| Stranger Aggression | GO:0042398 | cellular modified amino acid biosyntheti... | 0.04760 |
| Stranger Aggression | GO:0045880 | positive regulation of smoothened signal... | 0.04760 |
| Stranger Aggression | GO:2000378 | negative regulation of reactive oxygen s... | 0.04760 |
| Stranger Aggression | GO:0003176 | aortic valve development | 0.04770 |
| Stranger Aggression | GO:0006622 | protein targeting to lysosome | 0.04770 |
| Stranger Aggression | GO:0007252 | I-kappaB phosphorylation | 0.04770 |
| Stranger Aggression | GO:0030859 | polarized epithelial cell differentiatio... | 0.04770 |
| Stranger Aggression | GO:0045117 | azole transport | 0.04770 |
| Stranger Aggression | GO:0046596 | regulation of viral entry into host cell | 0.04770 |

| | | | |
|---|---|---|---|
| Stranger Aggression | GO:0048172 | regulation of short-term neuronal synapt... | 0.04770 |
| Stranger Aggression | GO:0050667 | homocysteine metabolic process | 0.04770 |
| Stranger Aggression | GO:0060272 | embryonic skeletal joint morphogenesis | 0.04770 |
| Stranger Aggression | GO:0070633 | transepithelial transport | 0.04770 |
| Stranger Aggression | GO:0071712 | ER-associated misfolded protein cataboli... | 0.04770 |
| Stranger Aggression | GO:0072393 | microtubule anchoring at microtubule org... | 0.04770 |
| Stranger Aggression | GO:1905809 | negative regulation of synapse organizat... | 0.04770 |
| Stranger Aggression | GO:2000052 | positive regulation of non-canonical Wnt... | 0.04770 |
| Stranger Aggression | GO:2000095 | regulation of Wnt signaling pathway, pla... | 0.04770 |
| Stranger Aggression | GO:0031532 | actin cytoskeleton reorganization | 0.04980 |
| Stranger Fear | GO:0051897 | positive regulation of protein kinase B ... | 0.00068 |
| Stranger Fear | GO:0050852 | T cell receptor signaling pathway | 0.00148 |
| Stranger Fear | GO:0010165 | response to X-ray | 0.00223 |
| Stranger Fear | GO:0032092 | positive regulation of protein binding | 0.00268 |
| Stranger Fear | GO:0035335 | peptidyl-tyrosine dephosphorylation | 0.00268 |
| Stranger Fear | GO:0006656 | phosphatidylcholine biosynthetic process | 0.00332 |
| Stranger Fear | GO:2001240 | negative regulation of extrinsic apoptot... | 0.00404 |
| Stranger Fear | GO:0000281 | mitotic cytokinesis | 0.00427 |
| Stranger Fear | GO:1901701 | cellular response to oxygen-containing c... | 0.00547 |
| Stranger Fear | GO:0007026 | negative regulation of microtubule depol... | 0.00581 |
| Stranger Fear | GO:0001780 | neutrophil homeostasis | 0.00737 |
| Stranger Fear | GO:0040019 | positive regulation of embryonic develop... | 0.00773 |
| Stranger Fear | GO:0018345 | protein palmitoylation | 0.00916 |
| Stranger Fear | GO:0042059 | negative regulation of epidermal growth ... | 0.00916 |
| Stranger Fear | GO:2000272 | negative regulation of signaling recepto... | 0.00916 |
| Stranger Fear | GO:0003015 | heart process | 0.00955 |
| Stranger Fear | GO:0043523 | regulation of neuron apoptotic process | 0.01085 |
| Stranger Fear | GO:0043388 | positive regulation of DNA binding | 0.01113 |
| Stranger Fear | GO:0071168 | protein localization to chromatin | 0.01118 |

| | | | |
|---|---|---|---|
| Stranger Fear | GO:1900087 | positive regulation of G1/S transition o... | 0.01118 |
| Stranger Fear | GO:0007569 | cell aging | 0.01125 |
| Stranger Fear | GO:1900182 | positive regulation of protein localizat... | 0.01332 |
| Stranger Fear | GO:0030540 | female genitalia development | 0.01344 |
| Stranger Fear | GO:0035556 | intracellular signal transduction | 0.01346 |
| Stranger Fear | GO:0034121 | regulation of toll-like receptor signali... | 0.01405 |
| Stranger Fear | GO:0071773 | cellular response to BMP stimulus | 0.01423 |
| Stranger Fear | GO:0042147 | retrograde transport, endosome to Golgi | 0.01541 |
| Stranger Fear | GO:0050673 | epithelial cell proliferation | 0.01567 |
| Stranger Fear | GO:0000186 | activation of MAPKK activity | 0.01654 |
| Stranger Fear | GO:0009225 | nucleotide-sugar metabolic process | 0.01868 |
| Stranger Fear | GO:2000036 | regulation of stem cell population maint... | 0.01868 |
| Stranger Fear | GO:0008156 | negative regulation of DNA replication | 0.01930 |
| Stranger Fear | GO:0043627 | response to estrogen | 0.02167 |
| Stranger Fear | GO:0071242 | cellular response to ammonium ion | 0.02167 |
| Stranger Fear | GO:0001843 | neural tube closure | 0.02303 |
| Stranger Fear | GO:0008277 | regulation of G-protein coupled receptor... | 0.02455 |
| Stranger Fear | GO:0030520 | intracellular estrogen receptor signalin... | 0.02497 |
| Stranger Fear | GO:0008015 | blood circulation | 0.02515 |
| Stranger Fear | GO:0070198 | protein localization to chromosome, telo... | 0.02519 |
| Stranger Fear | GO:1903533 | regulation of protein targeting | 0.02520 |
| Stranger Fear | GO:0030004 | cellular monovalent inorganic cation hom... | 0.02523 |
| Stranger Fear | GO:0007030 | Golgi organization | 0.02605 |
| Stranger Fear | GO:1901990 | regulation of mitotic cell cycle phase t... | 0.03093 |
| Stranger Fear | GO:0032091 | negative regulation of protein binding | 0.03101 |
| Stranger Fear | GO:0043524 | negative regulation of neuron apoptotic ... | 0.03114 |
| Stranger Fear | GO:0043370 | regulation of CD4-positive, alpha-beta T... | 0.03197 |
| Stranger Fear | GO:1904591 | positive regulation of protein import | 0.03198 |
| Stranger Fear | GO:0061213 | positive regulation of mesonephros devel... | 0.03203 |

| | | | |
|---|---|---|---|
| Stranger Fear | GO:1990748 | cellular detoxification | 0.03207 |
| Stranger Fear | GO:0032467 | positive regulation of cytokinesis | 0.03209 |
| Stranger Fear | GO:0046039 | GTP metabolic process | 0.03209 |
| Stranger Fear | GO:0015749 | monosaccharide transmembrane transport | 0.03218 |
| Stranger Fear | GO:0009607 | response to biotic stimulus | 0.03258 |
| Stranger Fear | GO:0048863 | stem cell differentiation | 0.03550 |
| Stranger Fear | GO:0033146 | regulation of intracellular estrogen rec... | 0.03605 |
| Stranger Fear | GO:2000241 | regulation of reproductive process | 0.03894 |
| Stranger Fear | GO:0006978 | DNA damage response, signal transduction... | 0.03896 |
| Stranger Fear | GO:0007379 | segment specification | 0.03896 |
| Stranger Fear | GO:0030859 | polarized epithelial cell differentiatio... | 0.03896 |
| Stranger Fear | GO:0032656 | regulation of interleukin-13 production | 0.03896 |
| Stranger Fear | GO:0032754 | positive regulation of interleukin-5 pro... | 0.03896 |
| Stranger Fear | GO:0046112 | nucleobase biosynthetic process | 0.03896 |
| Stranger Fear | GO:0050667 | homocysteine metabolic process | 0.03896 |
| Stranger Fear | GO:0061339 | establishment or maintenance of monopola... | 0.03896 |
| Stranger Fear | GO:0070633 | transepithelial transport | 0.03896 |
| Stranger Fear | GO:0071712 | ER-associated misfolded protein cataboli... | 0.03896 |
| Stranger Fear | GO:1901018 | positive regulation of potassium ion tra... | 0.03896 |
| Stranger Fear | GO:1903747 | regulation of establishment of protein l... | 0.03896 |
| Stranger Fear | GO:2000052 | positive regulation of non-canonical Wnt... | 0.03896 |
| Stranger Fear | GO:2000095 | regulation of Wnt signaling pathway, pla... | 0.03896 |
| Stranger Fear | GO:0060986 | endocrine hormone secretion | 0.04025 |
| Stranger Fear | GO:0030097 | hemopoiesis | 0.04148 |
| Stranger Fear | GO:1905515 | non-motile cilium assembly | 0.04153 |
| Stranger Fear | GO:0006182 | cGMP biosynthetic process | 0.04469 |
| Stranger Fear | GO:0008272 | sulfate transport | 0.04663 |
| Stranger Fear | GO:0009070 | serine family amino acid biosynthetic pr... | 0.04663 |
| Stranger Fear | GO:0033158 | regulation of protein import into nucleu... | 0.04663 |

| | | | |
|---|---|---|---|
| Stranger Fear | GO:0051354 | negative regulation of oxidoreductase ac... | 0.04663 |
| Stranger Fear | GO:0072337 | modified amino acid transport | 0.04663 |
| Stranger Fear | GO:0090026 | positive regulation of monocyte chemotax... | 0.04663 |
| Stranger Fear | GO:0035116 | embryonic hindlimb morphogenesis | 0.04936 |
| Touch Sensitivity | GO:0090503 | RNA phosphodiester bond hydrolysis, exon... | 0.00038 |
| Touch Sensitivity | GO:0021819 | layer formation in cerebral cortex | 0.00062 |
| Touch Sensitivity | GO:0035518 | histone H2A monoubiquitination | 0.00091 |
| Touch Sensitivity | GO:0097484 | dendrite extension | 0.00286 |
| Touch Sensitivity | GO:0097352 | autophagosome maturation | 0.00570 |
| Touch Sensitivity | GO:0034143 | regulation of toll-like receptor 4 signa... | 0.00589 |
| Touch Sensitivity | GO:2001224 | positive regulation of neuron migration | 0.00589 |
| Touch Sensitivity | GO:0090263 | positive regulation of canonical Wnt sig... | 0.00668 |
| Touch Sensitivity | GO:0051271 | negative regulation of cellular componen... | 0.00780 |
| Touch Sensitivity | GO:0006359 | regulation of transcription by RNA polym... | 0.00786 |
| Touch Sensitivity | GO:0050650 | chondroitin sulfate proteoglycan biosynt... | 0.00786 |
| Touch Sensitivity | GO:0070665 | positive regulation of leukocyte prolife... | 0.00881 |
| Touch Sensitivity | GO:0008625 | extrinsic apoptotic signaling pathway vi... | 0.00994 |
| Touch Sensitivity | GO:0045109 | intermediate filament organization | 0.01018 |
| Touch Sensitivity | GO:1903861 | positive regulation of dendrite extensio... | 0.01018 |
| Touch Sensitivity | GO:0062009 | secondary palate development | 0.01285 |
| Touch Sensitivity | GO:0090305 | nucleic acid phosphodiester bond hydroly... | 0.01369 |
| Touch Sensitivity | GO:0098930 | axonal transport | 0.01423 |
| Touch Sensitivity | GO:0007264 | small GTPase mediated signal transductio... | 0.01573 |
| Touch Sensitivity | GO:0030947 | regulation of vascular endothelial growt... | 0.01589 |
| Touch Sensitivity | GO:0090162 | establishment of epithelial cell polarit... | 0.01589 |
| Touch Sensitivity | GO:0048011 | neurotrophin TRK receptor signaling path... | 0.01929 |
| Touch Sensitivity | GO:0021766 | hippocampus development | 0.02008 |
| Touch Sensitivity | GO:0006998 | nuclear envelope organization | 0.02018 |
| Touch Sensitivity | GO:0070201 | regulation of establishment of protein l... | 0.02112 |

| | | | |
|---|---|---|---|
| Touch Sensitivity | GO:0090090 | negative regulation of canonical Wnt sig... | 0.02261 |
| Touch Sensitivity | GO:0016242 | negative regulation of macroautophagy | 0.02307 |
| Touch Sensitivity | GO:0034122 | negative regulation of toll-like recepto... | 0.02307 |
| Touch Sensitivity | GO:0036342 | post-anal tail morphogenesis | 0.02307 |
| Touch Sensitivity | GO:0048339 | paraxial mesoderm development | 0.02307 |
| Touch Sensitivity | GO:0006099 | tricarboxylic acid cycle | 0.02721 |
| Touch Sensitivity | GO:0035235 | ionotropic glutamate receptor signaling ... | 0.02721 |
| Touch Sensitivity | GO:0042176 | regulation of protein catabolic process | 0.02818 |
| Touch Sensitivity | GO:0042634 | regulation of hair cycle | 0.02835 |
| Touch Sensitivity | GO:0009792 | embryo development ending in birth or eg... | 0.02880 |
| Touch Sensitivity | GO:0000578 | embryonic axis specification | 0.03173 |
| Touch Sensitivity | GO:0018279 | protein N-linked glycosylation via aspar... | 0.03173 |
| Touch Sensitivity | GO:0030488 | tRNA methylation | 0.03173 |
| Touch Sensitivity | GO:0034976 | response to endoplasmic reticulum stress | 0.03175 |
| Touch Sensitivity | GO:0000278 | mitotic cell cycle | 0.03527 |
| Touch Sensitivity | GO:0050767 | regulation of neurogenesis | 0.03585 |
| Touch Sensitivity | GO:0046580 | negative regulation of Ras protein signa... | 0.03625 |
| Touch Sensitivity | GO:0042733 | embryonic digit morphogenesis | 0.03631 |
| Touch Sensitivity | GO:0007219 | Notch signaling pathway | 0.03654 |
| Touch Sensitivity | GO:0048643 | positive regulation of skeletal muscle t... | 0.03661 |
| Touch Sensitivity | GO:1902042 | negative regulation of extrinsic apoptot... | 0.03661 |
| Touch Sensitivity | GO:0030888 | regulation of B cell proliferation | 0.03672 |
| Touch Sensitivity | GO:0010927 | cellular component assembly involved in ... | 0.03675 |
| Touch Sensitivity | GO:0099175 | regulation of postsynapse organization | 0.03680 |
| Touch Sensitivity | GO:1901215 | negative regulation of neuron death | 0.03692 |
| Touch Sensitivity | GO:0099173 | postsynapse organization | 0.03908 |
| Touch Sensitivity | GO:0002757 | immune response-activating signal transd... | 0.03916 |
| Touch Sensitivity | GO:0070304 | positive regulation of stress-activated ... | 0.03925 |
| Touch Sensitivity | GO:0000018 | regulation of DNA recombination | 0.03929 |

| | | | |
|---|---|---|---|
| Touch Sensitivity | GO:0010657 | muscle cell apoptotic process | 0.03929 |
| Touch Sensitivity | GO:0033036 | macromolecule localization | 0.03930 |
| Touch Sensitivity | GO:0006986 | response to unfolded protein | 0.03938 |
| Touch Sensitivity | GO:0071222 | cellular response to lipopolysaccharide | 0.04211 |
| Touch Sensitivity | GO:0090287 | regulation of cellular response to growt... | 0.04574 |
| Touch Sensitivity | GO:0006310 | DNA recombination | 0.04579 |
| Touch Sensitivity | GO:0001952 | regulation of cell-matrix adhesion | 0.04585 |
| Touch Sensitivity | GO:0043551 | regulation of phosphatidylinositol 3-kin... | 0.04594 |
| Touch Sensitivity | GO:0047496 | vesicle transport along microtubule | 0.04609 |
| Touch Sensitivity | GO:0021549 | cerebellum development | 0.04614 |
| Touch Sensitivity | GO:0031032 | actomyosin structure organization | 0.04768 |
| Trainability | GO:0051099 | positive regulation of binding | 0.00180 |
| Trainability | GO:0046395 | carboxylic acid catabolic process | 0.00320 |
| Trainability | GO:0086019 | cell-cell signaling involved in cardiac ... | 0.00340 |
| Trainability | GO:0043552 | positive regulation of phosphatidylinosi... | 0.00450 |
| Trainability | GO:0046475 | glycerophospholipid catabolic process | 0.00480 |
| Trainability | GO:0006623 | protein targeting to vacuole | 0.00540 |
| Trainability | GO:0006956 | complement activation | 0.00690 |
| Trainability | GO:0031116 | positive regulation of microtubule polym... | 0.00690 |
| Trainability | GO:0051893 | regulation of focal adhesion assembly | 0.00750 |
| Trainability | GO:0032956 | regulation of actin cytoskeleton organiz... | 0.00760 |
| Trainability | GO:0050919 | negative chemotaxis | 0.00850 |
| Trainability | GO:0048024 | regulation of mRNA splicing, via spliceo... | 0.00930 |
| Trainability | GO:0001662 | behavioral fear response | 0.01190 |
| Trainability | GO:0010765 | positive regulation of sodium ion transp... | 0.01240 |
| Trainability | GO:1901021 | positive regulation of calcium ion trans... | 0.01240 |
| Trainability | GO:0006886 | intracellular protein transport | 0.01260 |
| Trainability | GO:0034260 | negative regulation of GTPase activity | 0.01460 |
| Trainability | GO:0006508 | proteolysis | 0.01480 |

| | | | |
|---|---|---|---|
| Trainability | GO:0050731 | positive regulation of peptidyl-tyrosine... | 0.01510 |
| Trainability | GO:0031334 | positive regulation of protein complex a... | 0.01890 |
| Trainability | GO:2000573 | positive regulation of DNA biosynthetic ... | 0.01950 |
| Trainability | GO:0021915 | neural tube development | 0.02010 |
| Trainability | GO:0032801 | receptor catabolic process | 0.02030 |
| Trainability | GO:0001764 | neuron migration | 0.02230 |
| Trainability | GO:0007029 | endoplasmic reticulum organization | 0.02510 |
| Trainability | GO:0050892 | intestinal absorption | 0.02510 |
| Trainability | GO:0031023 | microtubule organizing center organizati... | 0.02520 |
| Trainability | GO:0014706 | striated muscle tissue development | 0.02540 |
| Trainability | GO:0072347 | response to anesthetic | 0.02540 |
| Trainability | GO:0032885 | regulation of polysaccharide biosyntheti... | 0.02550 |
| Trainability | GO:2000249 | regulation of actin cytoskeleton reorgan... | 0.02640 |
| Trainability | GO:0042219 | cellular modified amino acid catabolic p... | 0.02650 |
| Trainability | GO:0045652 | regulation of megakaryocyte differentiat... | 0.02650 |
| Trainability | GO:0048172 | regulation of short-term neuronal synapt... | 0.02650 |
| Trainability | GO:2000104 | negative regulation of DNA-dependent DNA... | 0.02650 |
| Trainability | GO:0046907 | intracellular transport | 0.03440 |
| Trainability | GO:0015909 | long-chain fatty acid transport | 0.03470 |
| Trainability | GO:0035065 | regulation of histone acetylation | 0.03470 |
| Trainability | GO:0048841 | regulation of axon extension involved in... | 0.03470 |
| Trainability | GO:0060055 | angiogenesis involved in wound healing | 0.03470 |
| Trainability | GO:0090140 | regulation of mitochondrial fission | 0.03470 |
| Trainability | GO:0007276 | gamete generation | 0.03600 |
| Trainability | GO:0045724 | positive regulation of cilium assembly | 0.03680 |
| Trainability | GO:0050850 | positive regulation of calcium-mediated ... | 0.03680 |
| Trainability | GO:0001974 | blood vessel remodeling | 0.03890 |
| Trainability | GO:0014033 | neural crest cell differentiation | 0.04040 |
| Trainability | GO:0034613 | cellular protein localization | 0.04040 |

| | | | |
|---|---|---|---|
| Trainability | GO:0035137 | hindlimb morphogenesis | 0.04040 |
| Trainability | GO:0009063 | cellular amino acid catabolic process | 0.04050 |
| Trainability | GO:0090559 | regulation of membrane permeability | 0.04070 |
| Trainability | GO:0007156 | homophilic cell adhesion via plasma memb... | 0.04190 |
| Trainability | GO:0006506 | GPI anchor biosynthetic process | 0.04390 |
| Trainability | GO:0006910 | phagocytosis, recognition | 0.04390 |
| Trainability | GO:0007031 | peroxisome organization | 0.04390 |
| Trainability | GO:0009048 | dosage compensation by inactivation of X... | 0.04390 |
| Trainability | GO:0017157 | regulation of exocytosis | 0.04390 |
| Trainability | GO:0035461 | vitamin transmembrane transport | 0.04390 |
| Trainability | GO:0036037 | CD8-positive, alpha-beta T cell activati... | 0.04390 |
| Trainability | GO:0097320 | plasma membrane tubulation | 0.04390 |
| Trainability | GO:0098911 | regulation of ventricular cardiac muscle... | 0.04390 |
| Trainability | GO:0030073 | insulin secretion | 0.04410 |
| Trainability | GO:0051966 | regulation of synaptic transmission, glu... | 0.04440 |
| Trainability | GO:0097191 | extrinsic apoptotic signaling pathway | 0.04440 |
| Trainability | GO:0051091 | positive regulation of DNA binding trans... | 0.04920 |
